# A metabolic labeling-based chemoproteomic platform unravels the physiological roles of choline metabolites

**DOI:** 10.1101/2022.03.31.486572

**Authors:** Aditi Dixit, Gregor P. Jose, Chitra Shanbhag, Nitin Tagad, Jeet Kalia

**Author notes:** Correspondence may be addressed to: Dr. Jeet Kalia.

## Abstract

Choline is an essential nutrient for mammalian cells. Our understanding of the cellular functions of choline and its metabolites, independent of their roles as choline lipid metabolism intermediates, remains limited. In addition to fundamental cellular physiology, this knowledge has implications for cancer biology because elevated choline metabolite levels are a hallmark of cancer. Here, we establish the mammalian choline metabolite-interacting proteome by utilizing a photocrosslinkable choline probe. To design this probe, we performed metabolic labeling experiments with structurally diverse choline analogs that resulted in the serendipitous discovery of a choline lipid headgroup remodeling mechanism involving sequential dealkylation and methylation steps. We demonstrate that phosphocholine inhibits the binding of one of the proteins identified, the attractive anticancer target, p32, to its endogenous ligands and to the promising p32-targeting anticancer agent, Lyp-1. Our results reveal that choline metabolites play vital roles in cellular physiology by serving as modulators of protein function.

## INTRODUCTION

Metabolites serve as precursors, intermediates or end-products of cellular metabolic pathways. Additionally, metabolites interact with cellular proteins and allosterically modulate protein function. These metabolite-protein interactions regulate vital physiological processes including energy production, cellular communication, and protein and nucleic acid biosynthesis.^[1]^ Therefore, the identification of metabolite-interacting proteins and the delineation of the influence of these interactions on protein function is imperative.

The physiological roles of choline metabolites, apart from serving as intermediates in the dominant eukaryotic choline lipid biosynthetic pathway, the Kennedy pathway^[2]^ (Figure 1a), are poorly understood. This pathway starts with the cellular uptake of the nutrient, choline, followed by its sequential conversion into phosphocholine, cytidine 5’-diphospho (CDP) choline and finally to the lipids, phosphatidylcholine (PC) and sphingomyelin (SM) that are catabolized to glycerophosphocholine (GPC). Notably, the cellular levels of total water-soluble choline-containing metabolites (referred to as tCho and comprising choline, phosphocholine, CDP choline and GPC) are elevated in a variety of cancers including brain, prostate, breast and ovarian cancer.^[3]^ In epithelial ovarian cancer cells, for example, the tCho levels are as high as 7 mM and phosphocholine levels are enhanced 3-8 fold as compared to those in the immortalized non-tumoral cell variants.^[4]^ Phosphocholine levels are particularly elevated in cancer cells with concentrations as high as 9 mM reported in the breast cancer cell line, MDA-MB-231.^[5]^ Indeed, elevated tCho and phosphocholine levels have emerged as prominent hallmarks of cancer and non-invasive approaches for the measurement of their cellular levels are being actively pursued for diagnosis and monitoring disease progression.^[3a, 3b, 4]^ Nevertheless, the roles of choline metabolites in cancer pathophysiology outside of serving as precursors for the biosynthesis of choline lipids to meet the enormous membrane biogenesis requirements of rapidly proliferating cancer cells, are poorly understood. Identification of the cellular choline metabolite-interacting proteome followed by the characterization of the influence of those interactions on protein function is vital for unraveling the extended physiological roles of these metabolites.

**Figure 1.**
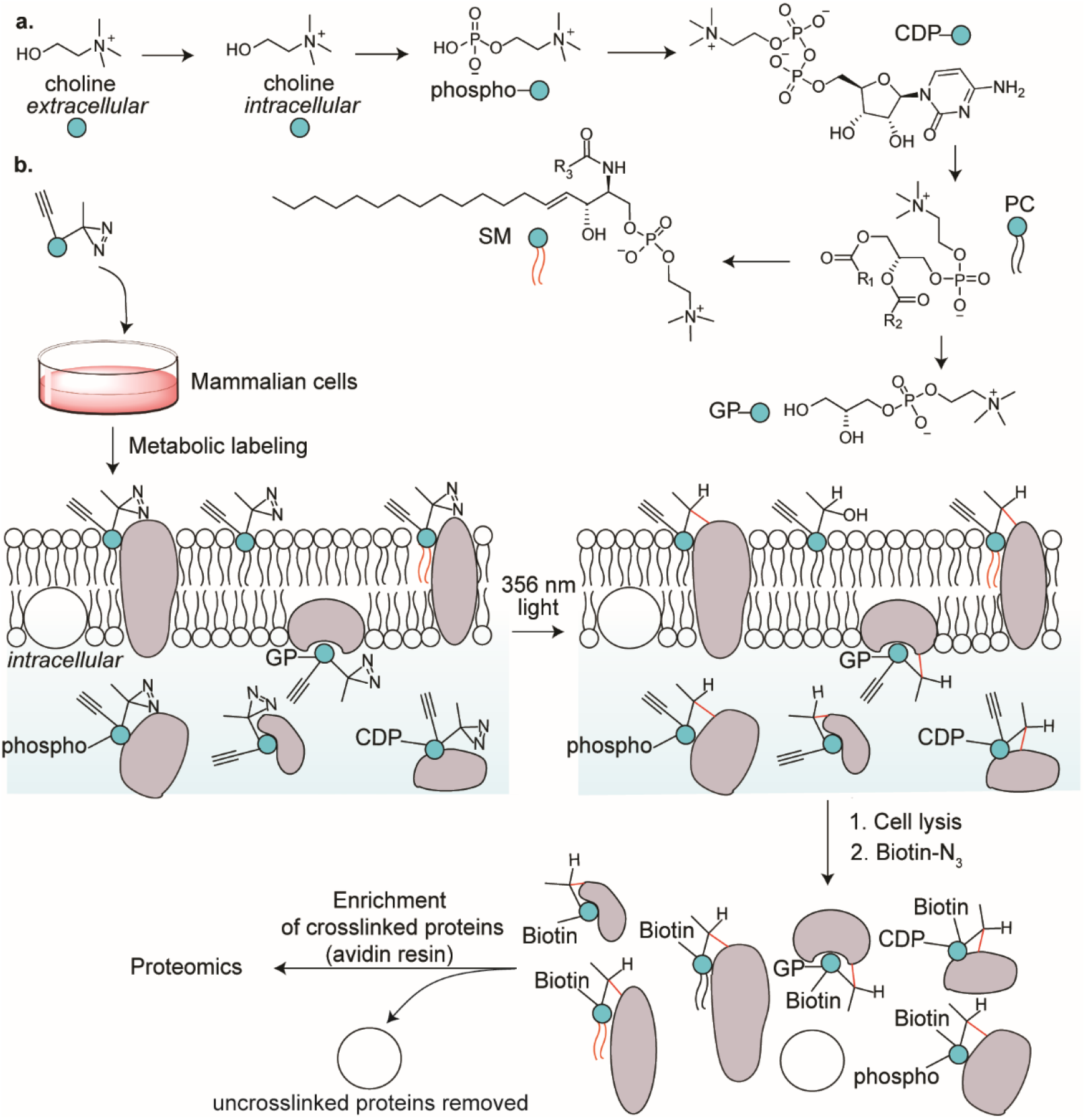
(a) The metabolism of choline in mammalian cells, and (b) our strategy of leveraging it to identify the choline metabolite-interacting proteome of mammalian cells. R_1_, R_2_ and R_3_ represent alkyl groups and the abbreviations CDP, PC, SM and GP denote cytidine 5’-diphospho, phosphatidylcholine, sphingomyelin and glycerophospho, respectively.

A powerful approach for the discovery of metabolite-interacting proteomes involves the cellular administration of bifunctional metabolite analogs appended with a photocrosslinkable group such as diazirine, and a chemical reporter such as alkyne.^[6]^ UV-irradiation of these cells leads to crosslinking of the metabolite analog to its interacting proteins via diazirine chemistry and the resultant crosslinked proteins are biotinylated via bioorthogonal reactions such as alkyne-azide cycloaddition. Subsequently, the proteins are isolated via affinity chromatography and identified by mass spectrometry. This approach enables the discovery of metabolite-interacting proteins in the native cellular milieu and is adept at capturing transient, low affinity interactions^[7]^ typically present in protein-metabolite complexes.^[8]^

Herein, we report the discovery of the mammalian choline metabolite-interacting proteome by deploying a rationally designed alkynyl diazirine choline analog capable of metabolically labeling choline metabolites (Figure 1b). The probe design was guided by our metabolic labeling studies focused on interrogating the promiscuity of the Kennedy pathway with respect to accepting choline analogs as precursors. Functional studies on selected protein hits revealed that choline metabolites profoundly modulate protein function. Additionally, our metabolic labeling studies resulted in the unexpected discovery of a choline lipid headgroup remodeling mechanism.

## RESULTS

### The choline lipid biosynthetic machinery is highly promiscuous

A critical prerequisite for the success of our approach was the amenability of choline metabolites to undergo metabolic labeling with a choline analog appended with both the diazirine and the alkyne/azido groups. Such a compound would be structurally substantially more elaborate than native choline. Therefore, the evaluation of the propensity of structurally diverse choline analogs to metabolically label choline metabolites was warranted. Towards this goal, we performed metabolic labeling studies on nine choline analogs (compounds **1-9**, Figure 2).

**Figure 2.**
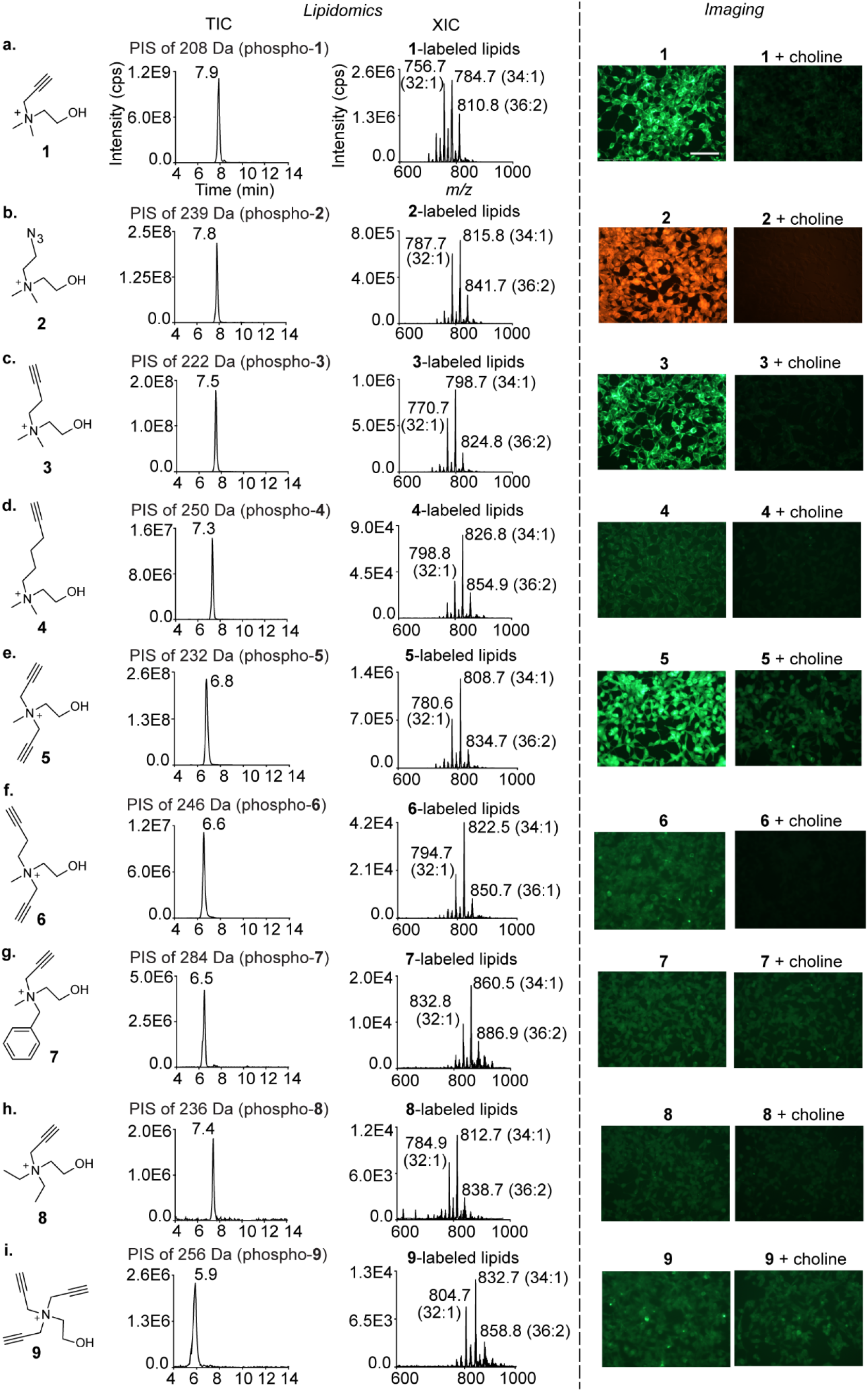
Characterization of the metabolic labeling of mammalian choline lipids with choline analogs via lipidomics (left panel) and cellular imaging (right panel). The lipidomics experiments were performed on the lipids extracted from analog (2 mM)-administered HEK293 cells and subjecting them to LC-MS/MS on a HILIC column in the precursor ion scan (PIS) mode. The resultant Total Ion Chromatograms (TICs) are depicted on the left with the retention time values indicated adjacent to the peaks. The *m/z* values of the daughter ions are denoted above the TICs. The Extracted Ion Chromatograms (XICs) with the three most abundant lipids annotated are depicted on the right. The XICs were processed via the LipidView software to annotate all the labeled lipids detected (Figures S2-S10). The imaging experiments involved treating analog (2 mM)-administered HEK293 cells (left) with either 5-azido fluorescein (for analogs **1**, **3-9**) or rhodamine alkyne (for analog **2**) under click chemistry conditions. In parallel, competition experiments involving the co-administration of equimolar native choline along with the analogs (2 mM of both) were also performed (right). The scale bar represents 100 μm.

Since PC and SM are the end-products of the Kennedy pathway (Figure 1a), the characterization of their metabolic labeling with choline analogs is also a readout of the metabolic labeling status of the precursor choline metabolites. Therefore, we focused on characterizing the metabolic labeling of these lipids via lipidomics and cellular imaging. Our lipidomics experiments entailed subjecting lipids extracted from choline analog-administered HEK293 cells to LC-MS/MS in the precursor ion scan (PIS) mode^[9]^ (Figure S1, supporting information). Our imaging experiments involved treating these cells with azido/alkynyl fluorescent dyes under click chemistry conditions as reported previously for imaging choline lipids metabolically labeled with the close structural mimics of choline, propargyl choline^[10]^ (**1**) and azidoethyl choline (**2**).^[11]^

A major insight from the results of these experiments was that the mammalian choline lipid biosynthetic machinery is remarkably flexible with respect to accepting structurally diverse choline analogs. Indeed, all of the nine analogs tested successfully labeled cellular choline lipids as demonstrated by the lipidomics analyses depicted in the left panel of Figure 2. Even the analogs that are structurally substantially different from native choline—hexynyl choline (**4**), the dialkynyl compound **6**, benzyl propargyl choline (**7**), and the tri-substituted derivatives **8** and **9**—yielded prominent total ion chromatogram (TIC) peaks for the labeled lipids. Importantly, the labeling efficiency dropped with increasing substituent chain lengths as demonstrated by a consistent attenuation of TIC peak intensities going from analog **1** to **4** (Figures 2a-d). This trend was also observed in the imaging results and was prominently highlighted by the dramatically lower fluorescence intensity observed for hexynyl choline (**4**)-labeled cells (Figure 2d) as compared to butynyl choline (**3**)-labeled cells (Figure 2c).

Notably, cells metabolically labeled with our most elaborate analogs (**4** and **6-9**) did not yield intense fluorescence signals (Figures 2d and f-i, right panel) despite confirmation of metabolic labeling via lipidomics (the corresponding left panels). These results demonstrate the higher sensitivity of our PIS lipidomics workflow as compared to our version of click chemistry-based imaging for characterizing metabolic labeling. In fact, fluorescence imaging as a means to characterize metabolic labeling was compatible with only those analogs (**1**, **2, 3** and **5**) that yielded TIC peak intensities of the order of E8 or above for the labeled lipids.

The results of this survey of choline analogs provided us a structural framework within which to design our desired bifunctional choline probes. The observation that extending the chain length of the alkyl substituents leads to reduced metabolic labeling efficiency motivated us to develop bifunctional choline analogs bearing the diazirine and the alkyne/azido groups on different alkyl substituents on the choline nitrogen, rather than on the same alkyl substituent as that would entail extending its length. The feasibility of this strategy was supported by the successful metabolic labeling of choline lipids by analogs **5, 6** and **7** wherein two *N*-methyl groups of choline were replaced with longer alkyl groups (Figures 2e-g).

### Metabolic labeling studies on β-diazirine cholines uncover a choline lipid headgroup remodeling mechanism

We began our metabolic labeling studies on photocrosslinkable choline analogs with the β-diazirine compounds **10** and **11** and discovered that both of them metabolically label HEK293 choline lipids (Figures 3a,b). Surprisingly however, in-gel fluorescence experiments wherein the lysates of UV-irradiated cells metabolically labeled with **11** were treated with a fluorescent azide under click chemistry conditions yielded negligible intensity for fluorescently labeled proteins (Figure 3b, bottom panel). ^1^H-NMR and UV spectroscopy experiments on **11** ruled out lack of UV-reactivity and chemical instability as possible reasons for these poor photocrosslinking yields (Figures S14 and S15).

**Figure 3.**
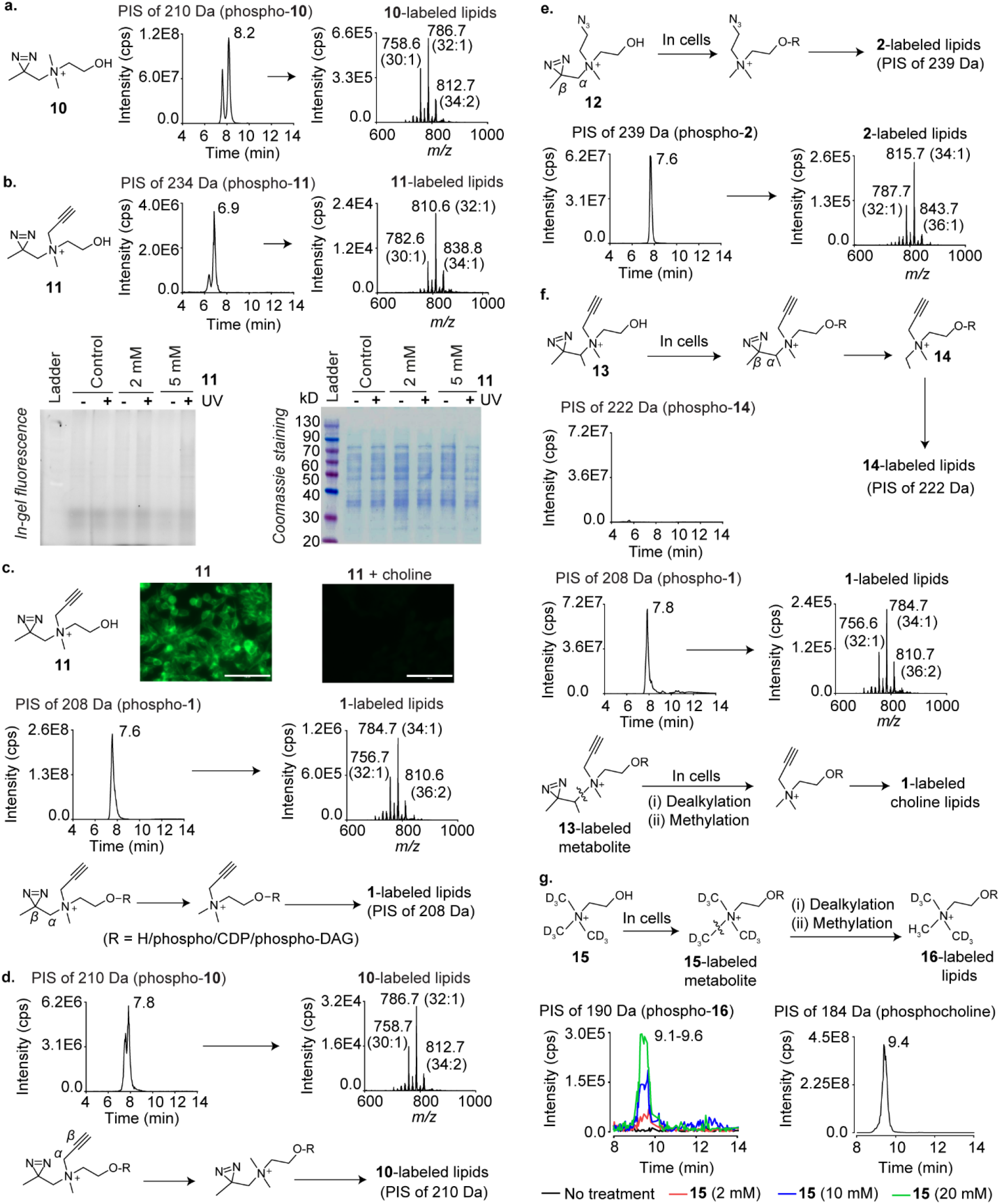
Metabolic labeling studies on β-diazirine choline analogs. a) PIS lipidomics to detect **10**-labeled lipids present in lipids extracted from **10**-administered HEK293 cells. b) Top panel: PIS lipidomics to detect **11**-labeled lipids present in lipids extracted from **11**-administered cells. Bottom panel: In-gel fluorescence experiments on cells metabolically labeled with **11**. c) Top panel: Imaging experiments on **11** (2 mM)-administered cells (left), and cells administered with both **11** and choline (2 mM each; right). The scale bars represent 100 μm. c) Middle panel: PIS lipidomics to detect **1**-labeled lipids present in lipids extracted from **11**-administered cells. Bottom panel: The C_α_–C_β_ bond cleavage hypothesis to explain the formation of **1**-labeled lipids in **11**-administered cells. DAG is an abbreviation for diacylglycerol. d) Top panel: PIS lipidomics to detect **10**-labeled lipids present in lipids extracted from **11**-administered cells. Bottom panel: The C_α_–C_β_ bond cleavage hypothesis to explain the formation of **10**-labeled lipids in **11**-administered cells. e) Top panel: The expected formation of **2**-labeled lipids in **12**-administered cells. Bottom panel: PIS lipidomics to detect **2**-labeled lipids present in lipids extracted from **12**-administered cells. f) Top panel: The expected formation of **14**-labeled lipids in **13**-administered cells. Second panel from the top: PIS lipidomics to detect **14**-labeled lipids present in lipids extracted from **13**-administered cells. Third panel from the top: PIS lipidomics to detect **1**-labeled lipids present in lipids extracted from **13**-administered cells. Bottom panel: A hypothesis invoking N–C bond cleavage followed by methylation to explain the formation of **1**-labeled choline lipids in **13**-administered cells. g) Top panel: A scheme depicting the expected generation of D6 labeled (**16**-labeled) choline lipids due to N–C bond cleavage followed by methylation in D9 choline (**15**)-administered cells. Bottom left panel: PIS lipidomics to detect **16**-labeled lipids present in lipids extracted from **15**-administered cells. Bottom right panel: PIS lipidomics to detect native choline lipids present in lipids extracted from cells cultured in standard media devoid of non-natural choline analogs. The LipidView analyses of all XICs depicting the individual labeled lipids are provided in the supporting information section. All analogs were administered at 2 mM concentrations unless otherwise stated.

Notably, although **11**-labeled lipids yielded a TIC peak intensity similar to those obtained for **7**, **8** and **9**-labeled lipids (~E6; Figure 3b and Figures 2g-i), cellular imaging experiments with **11** yielded intense fluorescence signals (Figure 3c, top panel), in contrast to the low signals obtained with **7**, **8**, and **9** (Figures 2g-i). In fact, the imaging results for **11** were reminiscent of those obtained with **1** (Figure 2a) engendering the hypothesis that **11**-administered cells form **1**-labeled choline lipids. Consistent with this hypothesis, PIS lipidomics experiments on lipids isolated from **11**-administered cells yielded a high-intensity TIC peak for **1**-labeled lipids (~E8; Figure 3c, middle panel). Cells administered with **11** also formed alkyne-devoid **10**-labeled lipids (Figure 3d). A possible explanation for these observations is that **11**-labeled choline metabolites form their **1** and **10**-labeled counterparts by undergoing C_α_–C_β_ bond cleavage within cells (α and β carbons implicated in each case are annotated in the schemes in bottom panels of Figures 3c,d).

The TIC peak intensity for **10**-labeled lipids was similar to that obtained for **11**-labeled lipids (~E6), and two orders of magnitude lower than that for **1**-labeled lipids (comparing the TICs in Figures 3d, b and c). These results demonstrated that **1**-labeled lipids are the dominant metabolically labeled choline lipid species formed in **11**-administered cells. Semi-quantitative experiments on HEK293 cells cultured in the presence of **11** for different time periods (4 h, 8 h, 16 h and 24 h) revealed that this trend persisted at all these time points (Figure S16). Metabolic labeling experiments with **11** on HeLa and MDA-MB-231 cells yielded similar results (Figure S18) demonstrating the generality of this phenomenon across mammalian cell lines. The formation of large amounts of **1**-labeled lipids accounts for the high fluorescence intensity observed in the imaging experiments on **11**-administered cells (Figure 3c, top panel), whereas the formation of small amounts of **11**-labeled choline metabolites explains the low intensity of fluorescently labeled crosslinked proteins obtained in the in-gel fluorescence experiment (Figure 3b, bottom panel). The problem of low protein crosslinking levels is likely aggravated due to competition for protein-binding sites between the sets of photocrosslinkable choline metabolites labeled with **11** and those labeled with **10**.

Two other bifunctional β-diazirine cholines, the azido diazirine **12** (Figure 3e and S21), and compound **S26** (Figure S22), a homolog of **11** containing one additional carbon atom on the diazirine-containing alkyl chain, were also processed by mammalian cells in a fashion similar to **11**, yielding diazirine-devoid azido/alkynyl choline lipids as the dominant metabolically labeled choline lipid species. Interestingly however, metabolic labeling experiments with the branched alkynyl β-diazirine choline compound **13** did not yield the expected major C_α_–C_β_ bond cleavage product, ethyl propargyl choline (**14**)-labeled lipids (top two panels of Figure 3f) and instead, a prominent TIC peak for **1**-labeled lipids was obtained (Figure 3f, third panel). The formation of **1**-labeled lipids in cells administered with both unbranched and branched alkynyl β-diazirine cholines supports a mechanism involving the cleavage of the N–C bonds between the choline nitrogen and their *N*-alkyl substituents followed by methylation (depicted for **13** in Figure 3f, bottom panel), over the C_α_–C_β_ bond cleavage mechanism proposed above.

Intrigued by the above findings on β-diazirine cholines, we investigated whether native choline metabolites are also processed in a similar fashion in mammalian cells by culturing HEK293 cells in the presence of deuterated (D9) choline. If D9 choline is processed similarly, D6 choline lipids should be formed (removal of the CD_3_ group due to N–C bond cleavage followed by the incorporation of the −CH_3_ group; Figure 3g, top panel). Consistent with this expectation, PIS analysis yielded dose-dependent signals for D6 choline lipids at the same retention time as that for native (H9) choline lipids (Figure 3g, bottom panel), establishing that native choline metabolites are also susceptible to sequential dealkylation and methylation in mammalian cells. The discovery of this novel choline lipid headgroup remodeling mechanism has important implications for fundamental lipid biology.

### γ-diazirine cholines are ideal metabolic labeling probes for choline lipids

The valuable insights on lipid biology that emerged from our metabolic labeling studies on β-diazirine cholines notwithstanding, these compounds are not suitable for choline metabolite-interactome mapping due to the low cellular photocrosslinking yields they afford (Figure 3b, bottom panel). Consequently, we next explored γ-diazirine cholines **17** and **18** and observed that they demonstrated robust metabolic labeling (Figures 4a,b). Notably, the TIC peak intensity for **18**-labeled lipids is more than 4-fold higher than that obtained for the analogous alkynyl β-diazirine choline compound (**11**, Figure 3b). Moreover, as expected from an alkynyl choline analog that yields a TIC peak intensity of <E8, a low fluorescence intensity was obtained upon subjecting cells metabolically labeled with **18** to our imaging conditions (Figure S25a) suggesting that this compound, unlike **11**, does not form large amounts of **1**-labeled lipids. Indeed, our lipidomics experiments demonstrated that lipids isolated from **18**-administered cells yielded a small TIC peak (~E6) for **1**-labeled lipids (Figure 4c) as compared to an ~E8 intensity TIC peak obtained for **1**-labeled lipids formed in **11**-administered cells (Figure 3c, middle panel). Furthermore, no detectable amount of choline lipids labeled with the alkyne-devoid diazirine (**17**) was observed (Figure 4d). In fact, the desired **18**-labeled choline lipids were the dominant metabolically labeled choline lipids formed in three different **18**-administered mammalian cell lines (HEK293, HeLa and MDA-MB-231; Figures 4e,f). These results are in stark contrast to those obtained with **11** that primarily forms **1**-labeled lipids (Figures S16 and S18) and unequivocally establish **18** as an ideal probe for metabolically labeling choline metabolites with the diazirine and the alkyne groups.

**Figure 4:**
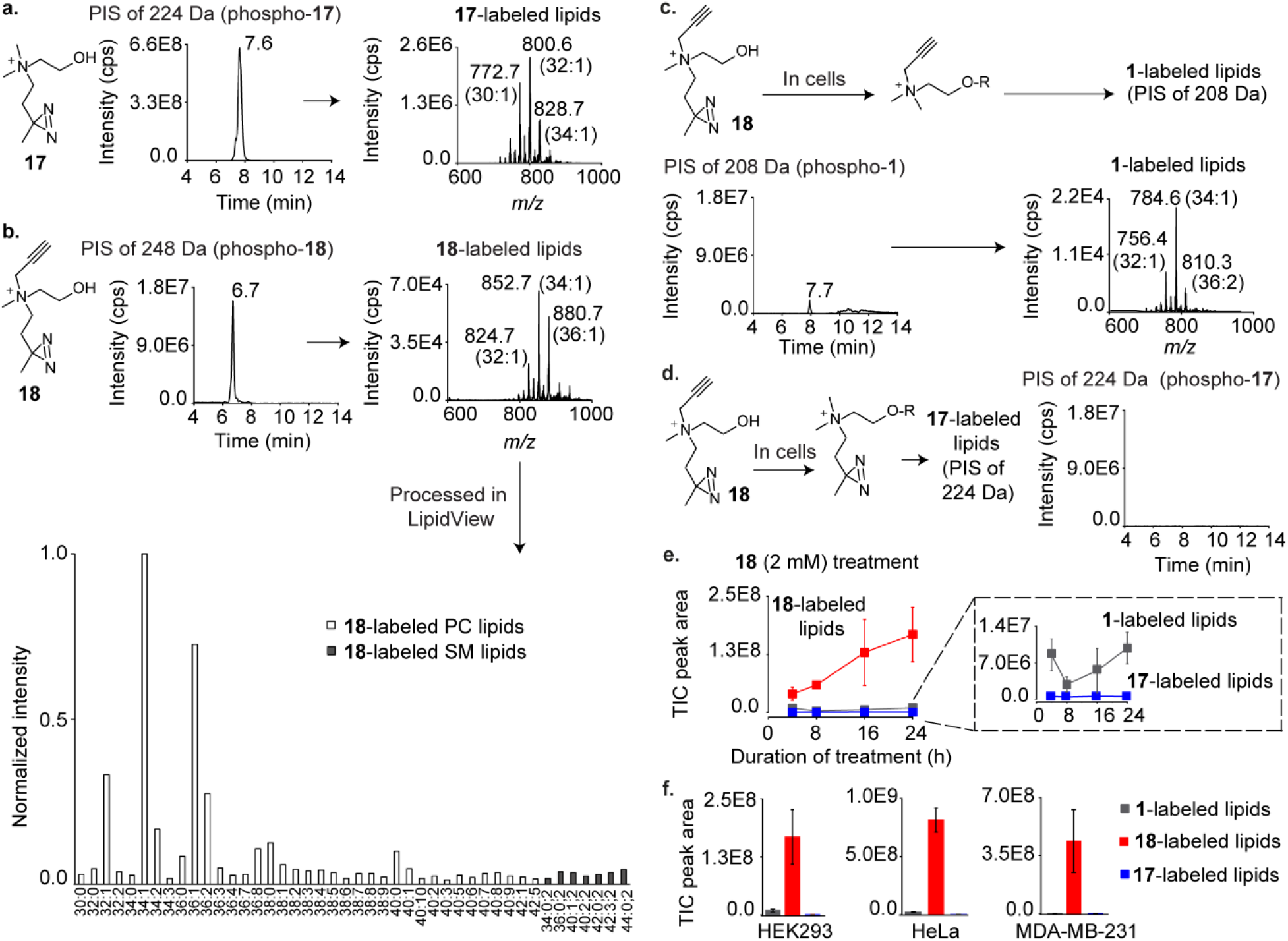
Metabolic labeling studies on γ-diazirine choline analogs. a) PIS lipidomics to detect **17**-labeled lipids present in lipids extracted from **17**-administered HEK293 cells. b) PIS lipidomics (along with LipidView analysis) to detect **18**-labeled lipids present in lipids extracted from **18**-administered cells. c) Top panel: The expected formation of **1**-labeled lipids in **18**-administered cells. Bottom panel: PIS lipidomics to detect **1**-labeled lipids present in lipids extracted from **18**-administered cells. d) Left: The expected formation of **17**-labeled lipids in **18**-administered cells. Right: PIS lipidomics to detect **17**-labeled lipids present in lipids extracted from **18**-administered cells. e) Time course plot for TIC peak areas corresponding to **1**, **17**, and **18**-labeled lipids present in lipids extracted from cells cultured in the presence of **18** for 4, 8, 16, and 24 h. f) The TIC peak areas for **1**, **17**, and **18**-labeled lipids present in lipids extracted from HEK293, HeLa, and MDA-MB-231 cells cultured in the presence of **18** for 24 h. Each data point in the plots depicted in e) and f) is an average of three biological replicates and the error bars represent the standard error values (representative TICs are depicted in Figures S26-S28). The LipidView analyses of the XICs depicted in a) and c) are provided in the supporting information section. All analogs were administered at 2 mM concentrations.

### Discovery of the mammalian choline metabolite-interacting proteome

In-gel fluorescence experiments with **18** yielded a multitude of fluorescently labeled proteins that demonstrated a dose-dependent (Figure 5a) and UV exposure time-dependent (Figure 5b) increase in fluorescence intensities. In contrast, neither of the four bifunctional β-diazirine choline analogs, **11**, **12**, **13** and **S26** yielded fluorescently labeled proteins (Figure 5c). Employing the optimal UV-irradiation and photocrosslinking conditions determined from these in-gel experiments to enrich crosslinked proteins via biotinylation yielded a dose-dependent enrichment of proteins (Figure 5d), demonstrating that compound **18** is an ideal probe for enriching the crosslinked proteome.

**Figure 5.**
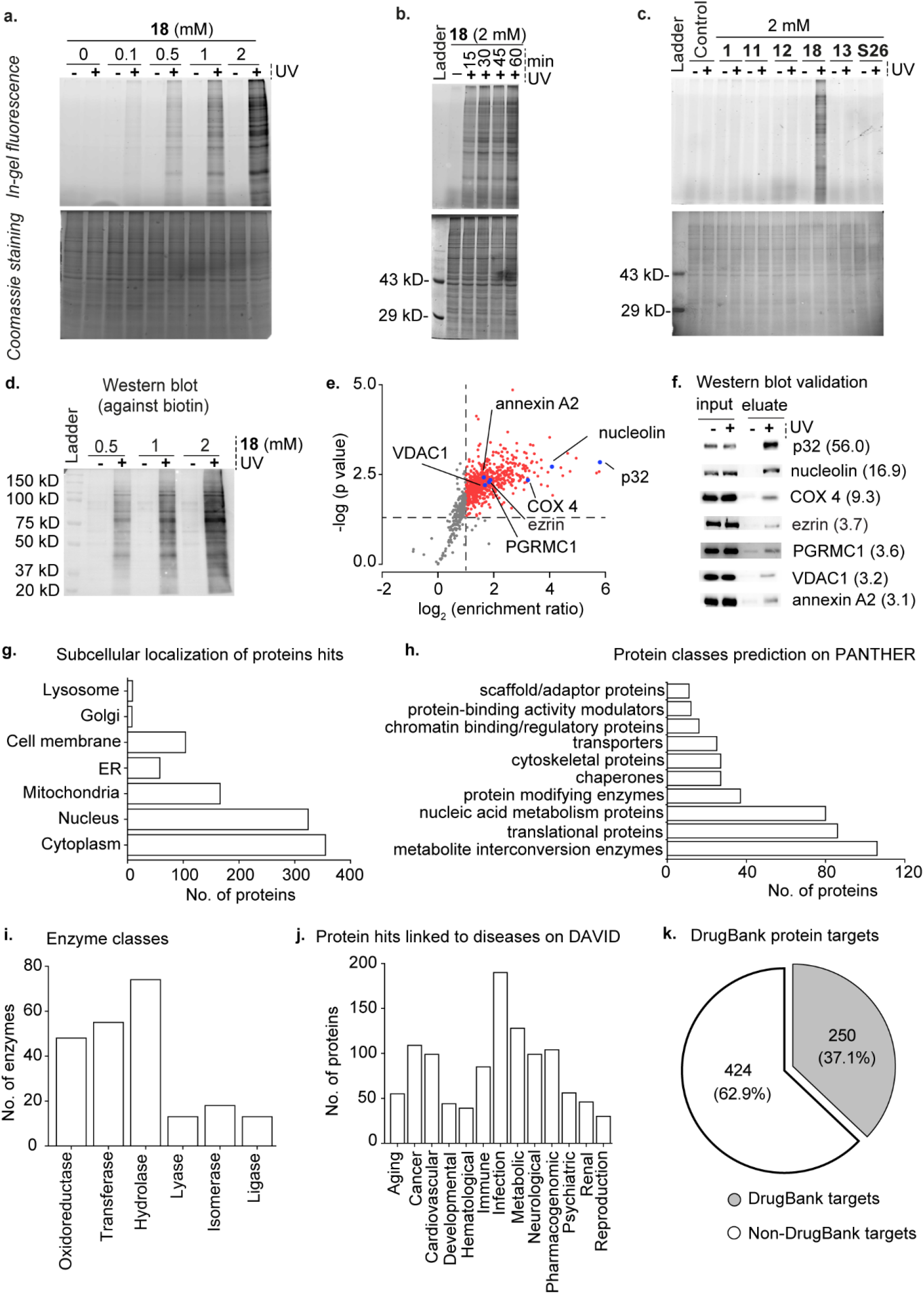
Identification of the choline metabolite-interacting proteome of HEK293 cells by using the γ-diazirine choline analog **18**. Panels a) to c) represent data obtained from in-gel fluorescence experiments with the fluorimager-captured images depicted on the top and the same gels subsequently stained with coomassie on the bottom. a) In-gel fluorescence experiments on HEK293 cells metabolically labeled with different concentrations of **18**. b) Evaluation of UV exposure time-dependence on the photocrosslinking efficiency in cells metabolically labeled with **18**. c) Comparison of photocrosslinking efficiency in cells metabolically labeled with the alkynyl diazirine choline probes **11**, **13**, **S26** and **18**, and the azido diazirine choline probe, **12**. Propargyl choline (**1**) was used a negative control. d) Western blot characterization of the dose-dependence of the avidin-enrichment of biotinylated UV-crosslinked proteins obtained from cells metabolically labeled with **18**. e) A plot depicting the results of our sixplex TMT analysis (three +UV and three −UV samples). The averaged enrichment ratio of the 902 proteins detected is plotted against their respective p-values and the 674 high-confidence protein hits that yielded an enrichment ratio of ≥ 2.00 (dotted line parallel to the *y*-axis) and p-value of ≤ 0.05 (dotted line parallel to the *x*-axis) are depicted as either red or blue dots. Only these high-confidence protein hits were considered for the analyses depicted in Figures 5g-k. f) Western blot validation of the enrichment of the seven proteins marked by blue dots in Figure 5e blotted by using primary antibodies against the individual proteins with their enrichment ratios depicted within parentheses. g) The subcellular distribution of our protein hits as per the UniProt database. h) The protein hits classified according to their biochemical functions as per the PANTHER database. i) The classification of the enzyme protein hits according to the UniProt database. j) The protein hits genetically linked to various disease classes as per the DAVID database. k) Pie-chart depicting the classification of the protein hits with respect to their annotation as drug targets according to the DrugBank database. The lists of all the proteins in the above analyses are provided in Table S3.

With optimal metabolic labeling, photocrosslinking and biotinylation conditions determined, we executed our chemoproteomic workflow depicted in Figure 1b. Specifically, we treated the cell lysate of UV-irradiated HEK293 cells cultured in the presence of **18** (2 mM) with azido biotin under click-chemistry conditions and applied the proteins extracted from this reaction mixture onto an avidin column. We performed these “+UV” experiments on three cell cultures along with the “−UV” experiments on three different cell cultures, and subjected the avidin resin containing the captured proteins obtained from these six independent sets of treatments to sixplex tandem mass tag (TMT) proteomics.^[12]^ This analysis yielded a total of 674 proteins with a minimum enrichment ratio of 2.0 and p value ≤ 0.05 which we annotated as “high-confidence hits” (depicted as red and blue dots in the plot shown in Figure 5e; the complete protein list is provided in Sheet 1 of Table S3).

To evaluate the veracity of this proteomics data, we performed Western blotting-based validation experiments on seven of these protein hits (p32, nucleolin, COX 4, ezrin, PGRMC1, VDAC1, and annexin A2) possessing a range of enrichment values (depicted as blue dots in Figure 5e). These experiments involved the UV-irradiation of HEK293 cells metabolically labeled with **18** followed by treatment with azido biotin under click chemistry conditions and enrichment on avidin resin. The proteins captured on the resin were eluted and probed for each of the seven proteins by Western blotting. The results of these experiments (Figure 5f) revealed UV-dependent enrichment of each of these proteins thereby validating the results of the proteomics studies.

### Choline metabolites interact with a diverse range of cellular proteins

A survey of the choline metabolite and lipid-binding proteins identified by our chemoproteomic approach revealed that they are expressed in a variety of cellular organelles and participate in diverse metabolic pathways. Whereas a majority of these proteins (355 out of 674) are expressed in the cytoplasm, 324 are nuclear and 166 are mitochondrial proteins (Figure 5g and Sheet 2 of Table S3). Analysis of our hits on the UniProt database revealed that the majority (73%) of these proteins are soluble proteins (Sheet 3 of Table S3). This conclusion was supported by in-gel fluorescence experiments that yielded a substantially higher fluorescence intensity in the soluble protein fraction as compared to the membrane protein fraction (Figure S29).

Although cellular proteins that interact with water-soluble choline metabolites have not been profiled previously, several proteins have been reported to bind to choline lipids, many of which are also present in our list of protein hits. These proteins include ADP/ATP translocase, the cytochrome c oxidase complex proteins COX 2, 4I1, 5A, 5B, 7A2, 7C, and subunits 1 and 2 of the cytochrome bc1 complex whose structures depict PC lipids bound to them.^[13]^ Additionally, the chaperone, HSP-90 beta that has been previously reported to bind to PC lipids was also identified in our study.^[14]^ Moreover, 60 of our protein hits were identified by Yao and co-workers to bind to PC lipids in HeLa cells (Sheet 4 of Table S3).^[15]^ Comparison of our protein hits with the PC-interacting yeast mitochondrial proteins identified by de Kroon and co-workers^[16]^ revealed that the mammalian homologs of 10 out of the 47 proteins reported in that study were present in our list of identified proteins (Sheet 5 of the Table S3).

Subjecting our protein hits to the PANTHER classification system^[17]^ resulted in their categorization into sixteen functional classes (Sheet 6 of Table S3; the ten most abundant classes are depicted in Figure 5h). Interestingly, a large number of proteins (106) belong to the metabolite interconversion enzymes category suggesting that choline metabolites may serve as allosteric modulators of these enzymes and regulate the biosynthesis of other metabolites. Proteins belonging to the translational protein and the nucleic acid metabolism categories contribute 86 and 80 proteins respectively to our list (Figure 5h and Sheet 6 of Table S3), several of which are implicated in disease states including the proliferating cell nuclear antigen (PCNA) that is involved in hypothyroidism and is targeted by the drug, liothyronine.^[18]^ Our list contains enzymes belonging to all of the six major enzyme subclasses (Figure 5i and Sheet 7 of Table S3), and is enriched in enzymes involved in key cellular metabolic pathways. Indeed, our list includes seven of the ten glycolytic enzymes, seven of the eight Kreb’s cycle enzymes, and proteins belonging to all five enzyme complexes of the oxidative phosphorylation (OXPHOS) pathway.

### Choline metabolite-interacting proteins play important roles in diseases and are attractive drug targets

Analysis on the DAVID resource^[19]^ revealed that 67% of our protein hits are associated with diseases, including 190 with infectious diseases, 109 with cancer, 128 with metabolic disorders, and 99 with neurological disorders (Figure 5j, Sheet 8 of Table S3). Remarkably, ~55% of the genes encoding for our protein hits are overexpressed in at least five different types of cancer according to the ONCOMINE database^[20]^ (Sheet 9 of Table S3). Moreover, several of our protein hits are known anticancer drug targets including PARP1, DNA topoisomerase 2-alpha, GART and isocitrate dehydrogenase 2 that are targeted by the drugs olaparib,^[21]^ doxorubicin,^[22]^ pemetrexed^[23]^ and enasidenib,^[24]^ respectively (Sheet 10 of Table S3). DrugBank database^[25]^ analysis revealed that in addition to cancer, the proteins identified in our study are targets for the therapy of several other diseases as well, with 37.1% (a total of 250 proteins) of them known to serve as drug targets (Figure 5k and Sheet 10 of Table S3). This percentage is much higher than the percentage of proteins among the entire human proteome pool that are known drug targets (12%),^[25–26]^ suggesting that choline metabolite-binding proteins are attractive targets for drug development.

### Phosphocholine profoundly modulates the binding of p32 with its interacting partners

To evaluate the importance of protein-choline metabolite interactions on protein function, we studied the effect of choline metabolites on the binding of our most enriched protein hit, p32 (enrichment ratio of 56; Figure 5e and Sheet 1 of Table S3), with a variety of its interacting partners. Also known as HABP1, p33 and gC1qR, p32 is a multifunctional protein that plays vital roles in a diverse array of physiological processes and disease states, especially cancer.^[27]^ p32 is expressed at disparate cellular locations including the mitochondrial matrix, the plasma membrane, the nucleus, the endoplasmic reticulum and the cytosol, and it performs diverse functions including facilitating mitochondrial translation,^[28]^ orchestrating cell attachment and migration by binding to extracellular matrix (ECM) components such as hyaluronic acid (HA)^[29]^ at the plasma membrane, and contributing to HIV infectivity by binding to the HIV Tat protein within the nucleus.^[30]^ p32 is overexpressed in malignant tumors found in various organs including lungs, breast and ovary,^[27a]^ and p32 expressed on the surface of cancer cells plays an important role in cell migration.^[31]^ The levels of plasma membrane-expressed p32 are dramatically enhanced in cancer cells and they correlate with the degree of cancer progression suggesting that cell surface p32 can serve both as a diagnostic marker for the disease as well as a target for anticancer drugs.^[27a]^ In fact, the promising anticancer and tumor imaging agent, the cyclic peptide Lyp-1, is recruited to cancer cells via its specific binding to cell surface-expressed p32.^[32]^ Moreover, the treatment of various cancer cell lines with antibodies against p32 inhibits cell migration and attenuates tumorigenesis in xenograft mouse models, demonstrating the tremendous potential of p32-binding agents for cancer therapy development.^[33]^ Indeed, small molecule libraries are actively being screened as p32 ligands to identify attractive anticancer lead compounds.^[34]^

Our studies on p32 began with investigating its binding to individual choline metabolites by performing drug affinity responsive target stability (DARTS) experiments.^[35]^ These experiments involved the treatment of recombinantly-produced p32 with the protease cocktail, pronase,^[36]^ in the presence of choline metabolites followed by SDS-PAGE analysis to evaluate whether they protect p32 from proteolysis. These experiments revealed that whereas choline, CDP choline, PC, SM and GPC, do not protect p32 from protease digestion (Figures 6a,c-f) pronase-mediated cleavage of p32 was substantially attenuated in the presence of 25 and 50 mM phosphocholine (Figure 6b; left panel). Identical experiments on the myoglobin protein demonstrated negligible phosphocholine-mediated protection against protease cleavage (Figure 6b; right panel) establishing that the protective effect of phosphocholine towards the proteolytic digestion of p32 is due to the specific binding of phosphocholine to p32.

**Figure 6.**
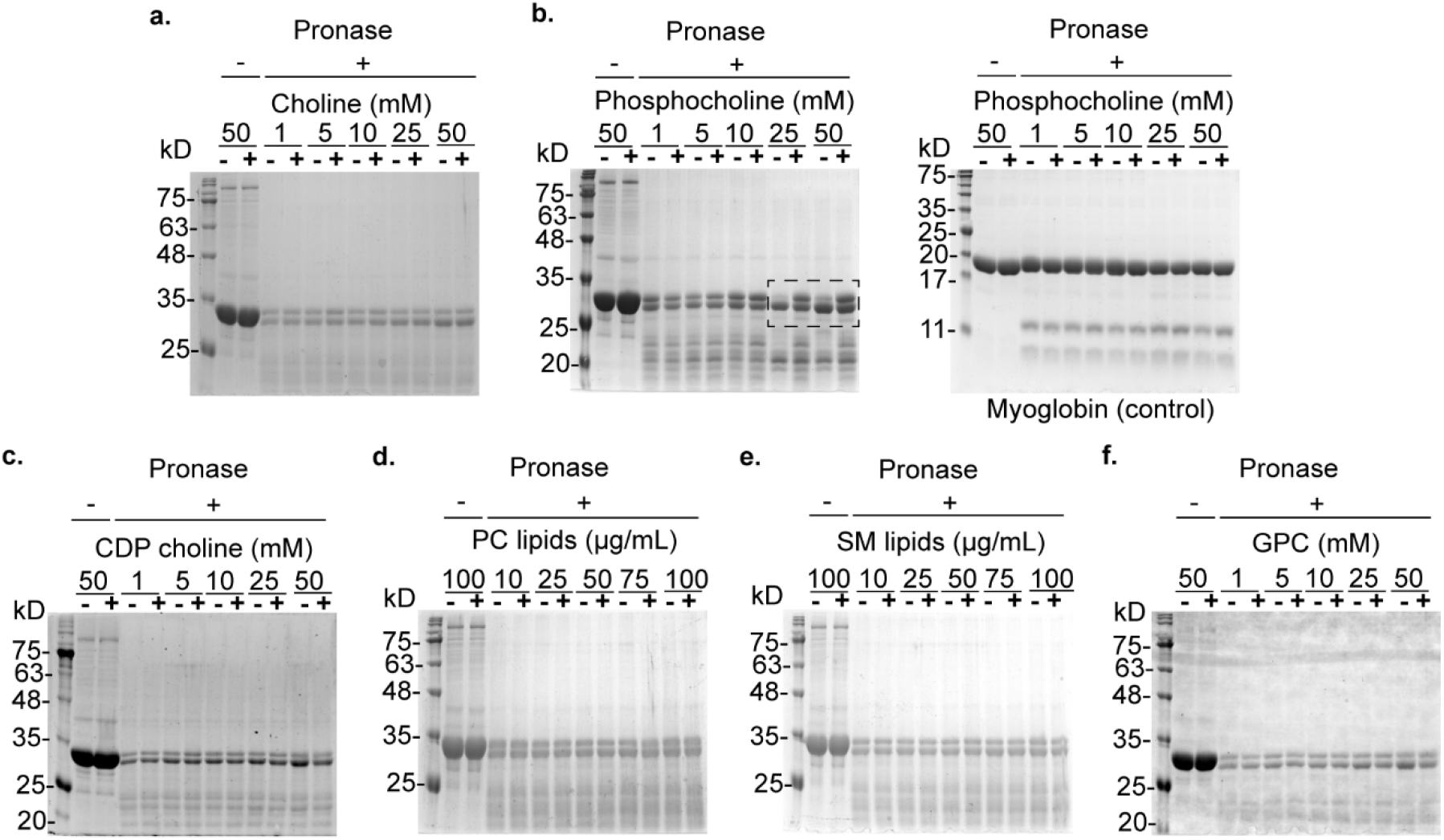
DARTS analysis of the binding of choline metabolites to p32. All images depicted are those of coomassie-stained gels. Pronase was added at a dilution of 1:300 (*w/w*) relative to p32 and proteolysis reactions were performed for 30 min. Experiments with choline (a), phosphocholine (b), CDP choline (c) and GPC (f) were performed with 1-50 mM concentrations of these metabolites (lanes marked “+”), whereas those with PC (d) and SM (e) were performed with DMSO-solubilized lipids (10-100 μg/mL). The metabolite-devoid negative controls (lanes marked “–”) for experiments with choline, CDP choline and GPC contained NaCl instead of the choline metabolites, and those for phosphocholine (used in its calcium salt form) contained CaCl_2_ instead of phosphocholine. The “–” lanes for the PC and SM experiments contained DMSO (vehicle). The enhanced proteolytic efficiency of pronase observed in the presence of 25 and 50 mM CaCl_2_ (the “–” lanes within the dotted rectangle in b) as compared to the other “–” lanes in the same gel corresponding to lower CaCl_2_ levels, is expected as pronase activity is catalyzed by Ca^2+^.^36^

We employed ELISA to characterize the effect of choline metabolites on the binding of p32 to its known interacting partners. These experiments entailed incubating biotinylated recombinant p32 on immobilized p32-interacting partners in the presence of physiologically relevant concentrations of the choline metabolites followed by estimation of the bound p32. Considering that phosphocholine is present in concentrations as high as 9 mM in mammalian cells,^[5]^ we employed 5 mM phosphocholine in these experiments. Other choline metabolites were used at 1 mM concentrations as they are present at lower concentrations in mammalian cells with GPC levels reaching 0.6 mM in MDA-MB-231 breast cancer cells and 1 mM in the PC-3 metastatic prostate cancer cells.^[3c, 5]^

We first focused on the p32-HA complex that is formed on the surface of cancer cells and plays an important role in tumor cell adhesion.^[27a, 29, 37]^ Our ELISA experiments revealed that phosphocholine inhibits the binding between p32 and HA in a competitive fashion (Figure 7a, left panel), whereas choline, CDP choline, GPC, PC and SM demonstrated minimal effects (Figure7a, middle and right panels). We next studied another interaction that occurs on cancer cell surfaces, the binding of p32 to the tumor homing peptide, Lyp-1, leading to the peptide’s cellular internalization, enabling it to trigger apoptosis.^[32]^ The formation of this complex was profoundly inhibited specifically by phosphocholine (Figure 7b).

**Figure 7.**
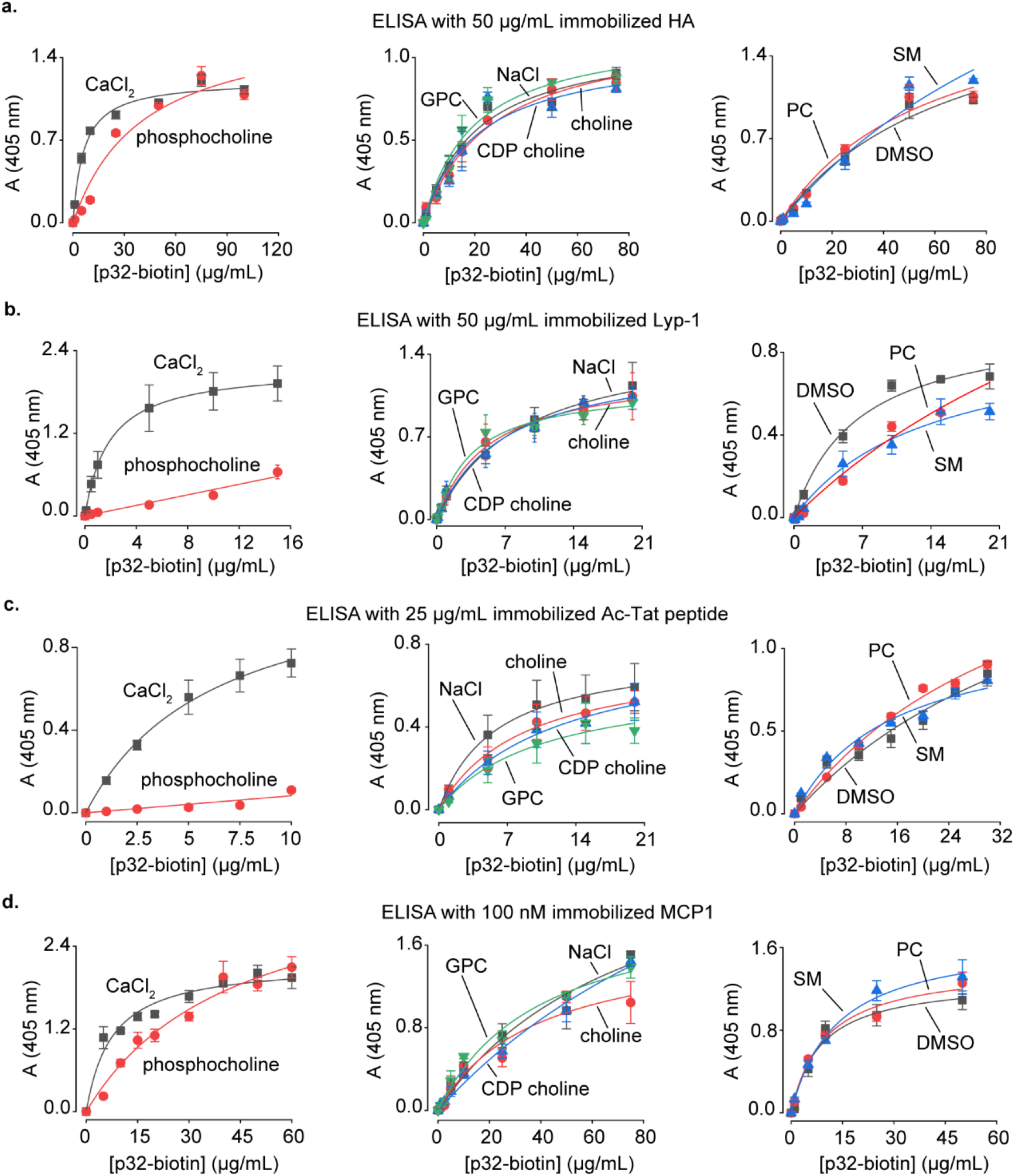
Studies on the modulation of binding of p32 with its interacting partners by choline metabolites. a-d) Characterization of the effect of choline metabolites on the binding of p32 to HA (a), Lyp-1 (b), acetylated-Tat peptide (c), and MCP1 (d) via ELISA. The metabolite/salt concentrations employed in the ELISAs were as follows: 1 mM choline/CDP choline/GPC/NaCl, 5 mM phosphocholine/CaCl_2_, and 100 μg/mL PC/SM.

In addition to its important role in cancer, p32 regulates viral infectivity in mammalian cells by binding to the acetylated-Tat protein of HIV-1 resulting in its recruitment to the HIV-1 promoter.^[30, 38]^ An 18 amino acids-long peptide spanning residues 36-53 of the Tat protein acetylated at Lys41, Lys50 and Lys51 recapitulates the binding of the acetylated-Tat protein to p32.^[30]^ ELISA on this acetylated-Tat peptide revealed that phosphocholine potently inhibits its binding to p32 whereas GPC, CDP choline, choline, PC and SM demonstrate negligible to moderate inhibitory effects (Figure 7c).

Another p32-interacting mammalian protein is the chemokine, monocyte chemoattractant protein-1 (MCP1). p32 serves as a pro-inflammatory agent by binding to MCP1 and slowing down its cellular degradation leading to an increase in its cellular levels.^[39]^ Our ELISA experiments revealed that phosphocholine inhibits the binding of p32 with MCP1 in a competitive fashion (Figure 7d).

Taken together, our studies on p32 establish that phosphocholine inhibits the binding of p32 to a range of its interacting partners, both endogenous and synthetic.

### Choline metabolites modulate the activity of cellular enzymes

We also characterized the effect of choline metabolites on the function of five of our enzyme hits: lactate dehydrogenase (LDH), citrate synthase, glyceraldehyde 3-phosphate dehydrogenase (GAPDH), malic enzyme and glutamic dehydrogenase (Sheet 1 of Table S3). LDH is overexpressed in almost all types of cancer and LDH inhibitors are promising anticancer agents.^[40]^ This enzyme catalyzes the conversion of pyruvate to lactate by employing NADH as a coenzyme (Figure S31a). At constant NADH concentrations, we observed that the *K*_m_ value for pyruvate was reduced ~5-fold in the presence of 1 mM choline, whereas it was halved in the presence of the same concentrations of CDP choline and GPC (Figure S31b,c). The presence of these three metabolites also caused an attenuation of the *K*_m_ values of the enzyme in assays wherein the concentration of pyruvate was kept constant whereas that of NADH was altered, with the largest effect seen with choline (a ~4-fold reduction, Figure S31b,d). Our experiments on citrate synthase revealed a ~2.6-fold attenuation of the substrate (oxaloacetate) *K*_m_ value at constant acetyl CoA concentrations in the presence of PC (Figure S32b,d). Assays on malic enzyme revealed a ~2 fold-attenuation of the *K*_m_ values of NADP^+^ in the presence of GPC, phosphocholine and PC, and a similar attenuation of the *K*_m_ value of malate in presence of GPC (Figure S34). GAPDH (Figure S33) and glutamate dehydrogenase (Figure S35) were negligibly modulated by choline metabolites.

## DISCUSSION

The goal of this study was to shed light on the functions of choline metabolites outside of their well-known roles as intermediates for choline lipid biosynthesis. Specifically, we focused on investigating the functional significance of choline metabolite-protein interactions by employing a sequential three-tier work flow consisting of the following components: a) metabolic labeling studies to identify a photocrosslinkable choline analog for labeling choline metabolites in mammalian cells, b) establishing a chemoproteomic platform utilizing this analog for the discovery of the mammalian choline metabolite-interacting proteome, and c) studies focused on characterizing the effect of choline metabolites on the function of proteins identified via this platform.

Our metabolic labeling studies provided valuable insights on lipid biology. Several of our choline analogs that successfully label choline lipids are structurally considerably more elaborate than native choline (Figure 2), thereby demonstrating that the mammalian choline lipid biosynthetic machinery is remarkably promiscuous. These findings add to the rapidly expanding toolkit of lipidic metabolic labeling probes that include non-natural fatty acids for labeling lipidic hydrophobic tails,^[15, 41]^ sphingosine derivatives for labeling sphingolipids,^[42]^ an azido *myo*-inositol derivative for labeling inositol lipids,^[43]^ and alkynols for labeling phosphatidic acid.^[44]^ Another significant outcome of these studies is the discovery that choline lipid headgroups are susceptible to chemical remodeling via sequential dealkylation and methylation in mammalian cells (Figures 3). This discovery will engender studies focused on elucidating the biochemical basis of this mechanism, and its roles in lipid homeostasis and disease pathophysiology.

Interestingly our ideal chemoproteomic probe, compound **18**, has been previously investigated for metabolically labeling choline lipids in mammalian cells.^[15]^ Based on imaging experiments, this previous study concluded that **18** is incapable of labeling choline lipids in HeLa cells. Consistent with this observation, our imaging experiments on HEK293 cells administered with **18** failed to yield fluorescence signals above background (Figure S25a). Yet, our lipidomics experiments convincingly demonstrated that **18** metabolically labels choline lipids in HEK293, HeLa and MDA-MB-231 cells (Figure 4f). These results establish the superiority of sensitive lipidomics approaches over standard imaging techniques for characterizing metabolic labeling.

Our functional studies on six selected protein hits revealed that four of them are modulated by choline metabolites. In particular, the results of our studies on the p32 protein were extremely insightful. These studies revealed that phosphocholine is a potent and general modulator of the binding of p32 with a variety of its interacting partners. This finding has major implications in cancer biology not only because p32 plays important roles in cancer progression and is an attractive target for anticancer drugs,^[32a, 34]^ but also because cancer is accompanied by a marked elevation in the cellular phosphocholine levels.^[3–5]^ In addition to cancer physiology, our studies on p32 implicate phosphocholine in mechanisms underlying HIV-1 infectivity and the inflammation response. The binding of p32 to HIV-1 acetylated-Tat is important for regulating HIV-1 infectivity,^[30]^ whereas its binding to MCP1 regulates the inflammation response.^[39]^ Therefore, our discovery that phosphocholine inhibits both of these binding interactions (Figures 7c,d) suggests that it plays important roles in these two processes.

In summary, our studies have provided novel insights on choline lipid biology and established the importance of choline metabolite-protein interactions in protein function. We anticipate that our findings will encourage studies focused on interrogating the functional significance of the interactions of choline metabolites with the other protein hits that we have identified.

## MATERIALS AND METHODS

Please see the supporting information section for organic synthetic schemes and procedures, compound characterization data, LipidView data, and protocols for metabolic labeling, cellular imaging, photocrosslinking, lipidomics, proteomics, Western blotting experiments, in-gel fluorescence, biotinylation, the recombinant production and biotinylation of p32, ELISA, DARTS assays and enzyme assays. Table S3 is uploaded as a separate file.

## Supporting information

SI_Choline

## AUTHOR CONTRIBUTIONS

J.K. conceived of the research and wrote the paper with inputs from A.D. A.D., C.S. and N.T. synthesized the choline probes. A.D. and G.P.J. performed the imaging, lipidomics, in-gel fluorescence and biotinylation experiments. A.D. analyzed the lipidomics data, performed the recombinant expression and biotinylation of p32, ELISA, the DARTS assays, and the enzyme assays.

## ACKNOWLEDGEMENTS

This work was supported by the DBT/Wellcome Trust India Alliance Fellowship [grant number IA/I/14/2/501551] awarded to J.K., and funds from IISER Pune and IISER Bhopal. G.P.J. thanks SERB for the SERB-National Postdoctoral Fellowship [PDF/2016/003750] that he secured under J.K.’s supervision. A.D. and N.T. thank IISER Pune and CSIR respectively for their graduate fellowships. We thank Drs. Sanjeev Shukla and Nagaraj Balasubramanian for providing access to the EVOS cell imaging system, Dr. Ankur Gupta for providing access to a UV-Visible spectrophotometer, and Dr. Himanshu Kumar for providing access to a Zeiss Axio Vert.A1 microscope and a Biorad microtiter plate reader. We thank Prof. Tambet Teesalu (University of Tartu, Estonia) for sharing the plasmid encoding the *N*-terminal histidine-tagged p32. The TMT proteomics experiments were outsourced to the Thermo Fisher Scientific Center for Multiplexed Proteomics at the Harvard Medical School (https://tcmp.hms.harvard.edu/).

