## Supplementary material for "A metabolic labeling-based chemoproteomic platform unravels the physiological roles of choline metabolites": SI_Choline

* Correspondence may be addressed to:

### Table of Contents


| Sub-section No. | Contents | Page No. | Table, figures, and schemes in the sub-section |
|---|---|---|---|
| 1. | General information | 3 | - |
| 2 | Synthetic schemes and procedures (Schemes S1-S16) | 4-33 | Schemes S1-S16 |
| 3. | General procedure for cell culture | 33-34 | - |
| 4. | Procedure for the characterization of metabolic labeling by cellular imaging | 34 | - |
| 5. | Procedure for the characterization of metabolic labeling by lipidomics | 35-40 | Figure S1, Table S1 |
| 6. | LipidView analysis of XICs obtained for choline lipids extracted from HEK293 cells cultured in the presence of compounds **1-11** | 40-44 | Figures S2-S13 |
| 7. | Photoactivation and stability studies on **11** | 45-46 | Figures S14, S15 |
| 8. | Metabolic labeling of HEK293, HeLa, and MDA-MB-231 cells with **11** | 47-53 | Figures S16-S20 |
| 9. | Metabolic labeling studies on the β-diazirine compounds **12**, **S26** and **13** | 53-55 | Figures S21-S23 |
| 10. | Metabolic labeling studies on D9 choline **15** | 55 | - |
| 11. | Metabolic labeling studies on the γ-diazirine analogs **17** and **18** | 56-59 | Figures S24-S28 |
| 12. | In-gel fluorescence experiments | 59-62 | Figure S29 |
| 13. | Biotinylation and affinity purification of crosslinked proteins | 62-64 | - |
| 14. | Sample preparation for quantitative proteomics (TMT) and target validation by Western blotting | 64-65 | Table S2 |
| 15. | Quantitative proteomics (TMT) experiments | 65-67 | - |
| 16. | Analysis of the high-confidence proteins obtained from proteomics studies | 67-68 | - |
| 17. | Characterizing the binding of p32 to choline metabolites and lipids | 68-71 | Figure S30 |
| 18. | Investigations on the effects of choline metabolites on the function of enzymes | 72-80 | Figures S31-S35 |
| 19. | References | 81-82 | - |
| 20. | $^{1}H$ and $^{13}C$ NMR spectra | 83-127 | - |


*Note:* Table S3 is not a part of this document and has been uploaded separately as an excel file. Table S3 contains spreadsheets depicting the raw proteomics data and lists of protein hits classified with respect to parameters including expression and functional profiles, involvement in diseases and roles as drug targets.

### 1. General information

All organic synthesis reactions were performed in oven-dried glassware. Chemical reagents for organic synthesis were purchased either from Sigma-Aldrich or Alfa Aesar and were used without purification. Distilled water was used for reaction work-ups, and ultrapure Type 1 water was used for buffer preparations and LC-MS experiments. The progress of chemical reactions was monitored by thin layer chromatography (TLC) using 0.25 mm Merck precoated (60 F254) silica gel plates. For visualization of TLC spots, UV light of wavelength 254 nm (for UV active compounds) and 365 nm (for fluorescent compounds) was used. In some cases, TLC spots were visualized using ninhydrin and phosphomolybdic acid (PMA) staining solutions. Purifications were performed using column chromatography on silica gel (100-200 mesh). All reactions, work-ups and column purifications involving diazirine compounds were performed under minimal exposure to light. $^{1}H$ and $^{13}C$ NMR spectra were recorded on Jeol-400, Bruker-400 or Bruker-500 spectrometers. Chemical shifts are reported as parts per million (δ) relative to tetramethylsilane (TMS) as internal standard and coupling constants (*J* values) in Hertz (Hz). Multiplicities are indicated as follows: s (singlet), d (doublet), t (triplet), q (quartet), dd (doublet of doublet), bs (broad singlet), td (triplet of a doublet) and m (multiplet). High-resolution mass spectra (HRMS) were recorded on HRMS-ESI-Q-Time of Flight LC/MS (Synapt G2, Waters). Lipidomics was performed on a triple quadrupole mass spectrometer (QTRAP® 4500 system from AB SCIEX) equipped with an Exion LC. All mass spectra (TIC and XIC) were analysed on the Analyst software and lipid species were identified by using the LipidView software. All UV exposure experiments were done by irradiating the sample with the UV light emitted from 5 × 8W UV lamps ($\lambda_{max}$ = 356 nm, Sankyo-Denki F8T5BLB lamp) and passed through a Magnaflux black light filter (part #519227). D9 choline chloride was purchased from Cambridge Isotope Laboratories, Lyp-1 peptide from Anaspec, MCP1 from R&D systems, and avidin-agarose from egg white and the sodium salt of hyaluronic acid (HA) from Merck. Acetylated Tat (Ac-Tat) peptide (aa 36 to 53 with the lysine residues at 41, 50 and 51 positions acetylated) was purchased from Grey Matter Research Foundation Pvt. Ltd. Details of antibodies used for western blots and ELISA experiments are provided in Table S2. Recombinant lactate dehydrogenase, citrate synthase, GAPDH, malic enzyme, and glutamate dehydrogenase were purchased from Sigma Aldrich. The enzyme assays were performed on a Perkin Elmer Lambda25 spectrophotometer.

### 2. Synthetic schemes and procedures

**Scheme S1.** Synthesis of propargyl choline **1**

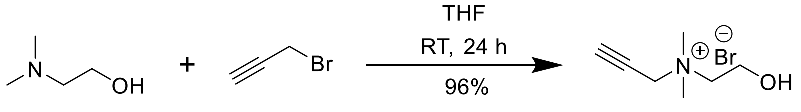


**1**

#### Propargyl choline (1)

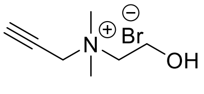

**1**

To an ice-cold solution of 2-dimethylaminoethanol (2 g, 22.4 mmol, 1 eq.) in THF (6 mL), 80% (*w*/*w*) propargyl bromide solution in toluene (2.89 mL, 26.9 mmol, 1.2 eq.) was added dropwise and the resultant mixture was stirred at 0 °C for 15 min. Subsequently, the mixture was warmed to room temperature and stirred for 24 h. The light-brown precipitate thus formed was collected by filtration and washed with cold THF (100 mL) to give **1** as a white solid (4.52 g, 96%).

$^1$H NMR (400 MHz, DMSO-d6) δ 5.34 (t, *J* = 5.0 Hz, 1H), 4.48 (d, *J* = 2.4 Hz, 2H), 4.07 (t, *J* = 2.5 Hz, 1H), 3.85 (q, *J* = 4.8 Hz, 2H), 3.49 – 3.47 (m, 2H), 3.16 (s, 6H).

$^{13}$C NMR (100 MHz, DMSO-d6) δ 82.9, 72.6, 64.7, 54.9, 54.5, 50.6.

HRMS (ESI): $C_7H_{14}NO^+$; Calculated mass, $[M]^+$ 128.1070; Observed mass, $[M]^+$ 128.1.

#### Azido choline 2

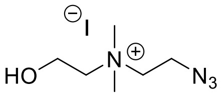

**2**

Azido choline **2** was synthesized according to a previously reported procedure.[1]

**Scheme S2.** Synthesis of butynyl choline **3**

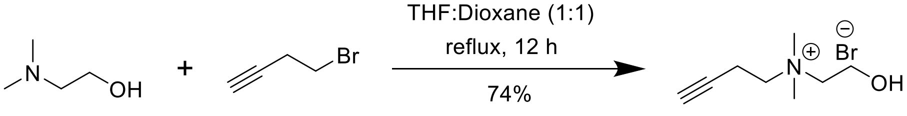


**3**

#### Butynyl choline (3)

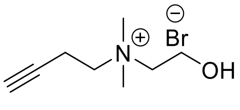

**3**

To a solution of 2-dimethylaminoethanol (0.5 mL, 4.97 mmol, 1 eq.) in a mixture of THF and dioxane (4 mL of a 1:1 solution), 4-bromobutyne (0.7 mL, 7.45 mmol, 1.5 eq.) was added and the reaction mixture was refluxed for 12 h. Subsequently, the solvent was evaporated under reduced pressure and the resultant brown residue was dissolved in ethanol (5 mL) and stored at -20 °C overnight. The precipitate thus obtained was collected, washed with cold ethanol (50 mL), and vacuum-dried to give dirty-white crystals of **3** (0.82 g, 74 %).

$^{1}$H NMR (400 MHz, DMSO-d6) δ 5.30 (t, *J* = 4.8 Hz, 1H), 3.82 (bs, 2H), 3.56 (t, *J* = 7.6 Hz, 2H), 3.43 (t, *J* = 4.8 Hz, 2H), 3.10 (s, 6H), 2.77 (t, *J* = 6.4 Hz, 2H).

$^{13}$C NMR (100 MHz, DMSO-d6) δ 79.9, 74.2, 65.5, 62.5, 55.4, 51.5, 13.5.

HMRS (ESI): $C_8H_{16}NO^+$; Calculated mass, $[M]^+$ 142.1226; Observed mass, $[M]^+$ 142.1232.

**Scheme S3.** Synthesis of hexynyl choline **4**

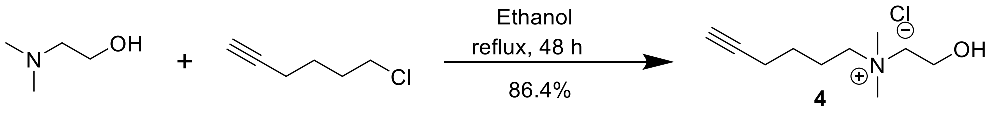


**Hexynyl choline (4)**

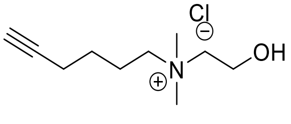

**4**

To a solution of 2-dimethylaminoethanol (0.5 mL, 4.12 mmol, 1 eq.) in ethanol (11 mL), 6-chlorohexyne (2 mL, 16.5 mmol, 4 eq.) was added and the resulting mixture was refluxed for 48 h. Subsequently, the solvent was removed under reduced pressure to give a heterogeneous mixture of a white solid and a yellow viscous oil. This mixture was dissolved in acetone (2 mL) and incubated in a dry ice-acetone cooling mixture at -78 °C for 1 h. A white precipitate was observed to form. The yellow solution was decanted and the precipitate was triturated with ice-cold acetone (3 × 2 mL) to give **4** as a white solid (0.88 g, 86.4%).

$^{1}$H NMR (400 MHz, DMSO-d6) δ 5.51 (t, *J* = 5.2 Hz, 1H), 3.81 (bs, 2H), 3.40 – 3.34 (m, 2H), 3.07 (s, 6H), 2.86 (t, *J* = 2.4 Hz, 1H), 2.23 (td, *J* = 6.8 Hz, 2.4 Hz, 2H), 1.80 – 1.72 (m, 2H), 1.49 – 1.42 (m, 2H).

$^{13}$C NMR (100 MHz, DMSO-d6) δ 83.8, 71.8, 64.6, 63.5, 54.8, 50.9, 24.9, 22.5, 21.1, 17.3.

HRMS (ESI): $C_{10}H_{20}NO^{+}$; Calculated mass, $[M]^+$ 170.1539; Observed mass, $[M]^+$ 170.1544.

**Scheme S4**. Synthesis of dipropargyl choline **5**

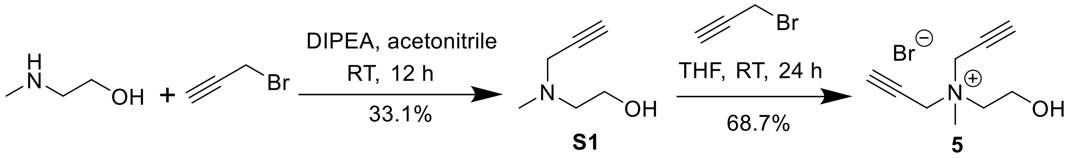


**2-(methyl(prop-2-yn-1-yl)amino)ethan-1-ol (S1)**

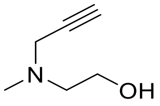

**S1**

To an ice-cold solution of acetonitrile (5 mL) containing *N*-methylaminoethanol (0.5 g, 6.65 mmol, 1.0 eq.) and *N*,*N*-diisopropylethylamine (1.0 mL, 6.0 mmol, 0.9 eq.), 80% (*w*/*w*) propargyl bromide solution in toluene (0.65 mL, 6.0 mmol, 0.9 eq.) was added dropwise. The resultant reaction mixture was stirred at 0 °C for 30 min and then allowed to warm to room temperature and stirred for 12 h. Subsequently, the solvent was removed under reduced pressure and the residue was dissolved in dichloromethane (25 mL) and washed with saturated sodium bicarbonate solution (25 mL). The aqueous layer was further washed with dichloromethane (3 × 25 mL). The organic layers were pooled, dried by adding anhydrous sodium sulfate, filtered and concentrated under reduced pressure and subjected to silica gel column chromatography (10% methanol in ethyl acetate) to afford **S1** as a yellow-colored oil (0.25 g, 33.1%).
$^1$H NMR (400 MHz, $CDCl_3$) δ 3.61 (t, *J* = 5.2 Hz, 2H), 3.39 (d, *J* = 2.4 Hz, 2H), 2.63 (t, *J* = 5.2 Hz, 2H), 2.35 (s, 3H), 2.23 (t, *J* = 2.4 Hz, 1H).
$^{13}$C NMR (100 MHz, $CDCl_3$) δ 78.4, 73.4, 58.7, 57.0, 45.8, 41.3.
HRMS (ESI): $C_6H_{12}NO$; Calculated mass, $[M+H]^+$ 114.0913; Observed mass, $[M+H]^+$ 114.0924.

**Dipropargyl choline (5)**

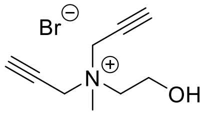

**5**

An ice-cold solution of **S1** (50.5 mg, 0.45 mmol, 1 eq.) in THF (1 mL) was treated with 80% (*w*/*w*) propargyl bromide in toluene (59 µL, 0.54 mmol, 1.2 eq.). The resultant solution was stirred under ice-cold conditions for 30 min and then at room temperature for 24 h. A yellow oil was observed to settle down at the bottom of the round-bottomed flask which upon trituration with ice-cold chloroform (8 × 2 mL), followed by drying under reduced pressure gave **5** as a pale-yellow solid (71.2 mg, 68.7%).

$^{1}$H NMR (400 MHz, DMSO-d6) δ 5.44 (m, 1H), 4.49 (t, *J* = 2.4 Hz, 4H), 4.12 (t, *J* = 2.4 Hz, 2H), 3.87 (bs, 2H), 3.54 (t, *J* = 5.2 Hz, 2H), 3.18 (s, 3H).

$^{13}$C NMR (100 MHz, DMSO-d6) δ 83.6, 71.9, 62.8, 54.7, 52.4, 48.3.

HRMS (ESI): $C_9H_{14}NO^+$; Calculated mass, $[M]^+$ 152.1070; Observed $[M]^+$ 152.1077.

**Scheme S5.** Synthesis of propargyl butynyl choline **6**

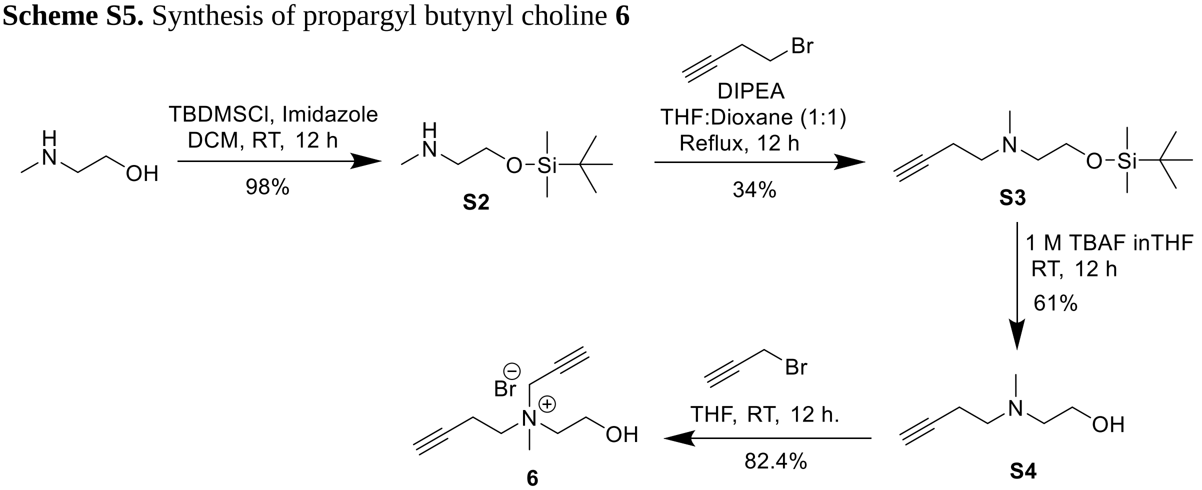


**2-((*t*-butyldimethylsilyl)oxy)-*N*-methylethan-1-amine (S2)**

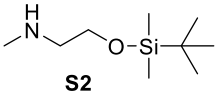

**S2**

To an ice-cold solution of *N*-methylaminoethanol (3 g, 39.9 mmol, 1 eq.) and imidazole (2.85 g, 41.9 mmol, 1.05 eq.) in dichloromethane (30 mL), was added an ice-cold solution of solution of *t*-butyldimethylsilylchoride (6.33 g, 42 mmol, 1.05 eq.) in dichloromethane (20 mL) under nitrogen atmosphere. Before mixing these two solutions, nitrogen gas was bubbled through each of them for 30 min. The resultant solution was allowed to warm to room temperature and stirred for 12 h. The reaction mixture was subsequently washed with 50 mL of water and the organic layer was concentrated under vacuum to give **S2** as a white solid (7.43 g, 98%).

$^{1}$H NMR (400 MHz, $CDCl_3$) δ 3.76 (t, *J* = 5.2 Hz, 2H), 2.74 (t, *J* = 5.2 Hz, 2H), 2.49 (s, 3H), 0.89 (s, 9H), 0.06 (s, 6H).

$^{13}$C NMR (125 MHz, $CDCl_3$) δ 59.7, 51.4, 34.1, 26.0, 18.4, -5.3.

HRMS (ESI): $C_9H_{23}NOSi$; Calculated mass, $[M+H]^+$ 190.1622; Observed $[M+H]^+$ 190.1634.

**N-(2-((t-butyldimethylsilyl)oxy)ethyl)-N-methylbut-3-yn-1-amine (S3)**

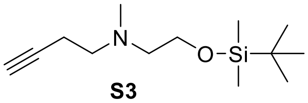


**S3**

To a solution of **S2** (1 g, 5.3 mmol, 1 eq.) and *N,N*-diisopropylethylamine (0.9 mL, 5.3 mmol, 1 eq.) in a mixture of THF and dioxane (15 mL of a 1:1 solution), 4-bromobutyne (0.5 mL, 5.3 mmol, 1 eq.) was added dropwise. After refluxing the resultant mixture for 12 h, the solvent was removed under reduced pressure and the residue was dissolved in dichloromethane (15 mL) and washed with saturated solution of sodium bicarbonate (2 × 20 mL). The organic extracts were combined, dried by adding anhydrous sodium sulfate and concentrated under reduced pressure. The resultant residue was subjected to silica-gel column chromatography (15% ethyl acetate in hexane) to yield **S3** as a yellow oil (436.5 mg, 34%).

$^1$H NMR (400 MHz, $CDCl_3$) δ 3.71 (t, *J* = 6.4 Hz, 2H), 2.67 (t, *J* = 7.5 Hz, 2H), 2.57 (t, *J* = 6.4 Hz, 2H), 2.37 – 2.33 (m, 2H), 2.32 (s, 3H), 1.96 (s, 1H), 0.89 (s, 9H), 0.06 (s, 6H).

$^{13}$C NMR (125 MHz, $CDCl_3$) δ 83.1, 69.0, 61.8, 59.3, 56.8, 42.9, 26.1, 18.5, 17.2, -5.2.

HRMS (ESI): $C_{13}H_{27}NOSi$; Calculated mass, $[M+H]^+$ 242.1935; Observed $[M+H]^+$ 242.1956.

**2-(but-3-yn-1-yl(methyl)amino)ethan-1-ol (S4)**

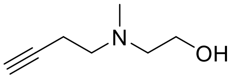


**S4**

To a round-bottomed flask containing **S3** (800 mg, 3.31 mmol, 1 eq.) in THF (30 mL), tetrabutylammonium fluoride (4.96 mL of a 1 M solution in THF, 4.96 mmol, 1.5 eq.) was added. After stirring the mixture at room temperature for 12 h, the solvent was removed under reduced pressure and the residue was subjected to using silica gel column chromatography (5% methanol in ethyl acetate) to yield **S4** as a yellow liquid (255.0 mg, 61%).

$^1$H NMR (400 MHz, $CDCl_3$) δ 3.58 (t, *J* = 5.2 Hz, 2H), 2.64 (t, *J* = 7.2 Hz, 2H), 2.57 (t, *J* = 5.2 Hz, 2H), 2.38 – 2.34 (m, 2H), 2.30 (s, 3H), 1.98 (t, *J* = 2.4 Hz, 1H).

$^{13}$C NMR (125 MHz, $CDCl_3$) δ 82.7, 69.4, 58.54, 58.50, 55.9, 41.5, 17.4.

HRMS (ESI): $C_7H_{13}NO$; Calculated mass, $[M+Na]^+$ 150.0889; Observed $[M+Na]^+$ 150.0912.

**Propargyl butynyl choline (6)**

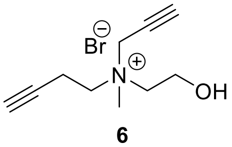


**6**

To an ice-cold solution of **S4** (255 mg, 2 mmol, 1 eq.) in THF (3 mL), 80% (*w/w*) propargyl bromide in toluene (260 µL, 2.4 mmol, 1.2 eq.) was added and the resultant mixture was stirred under ice-cold conditions for 15 min and then at room temperature for 12 h. A brown precipitate was observed to settle down at the bottom of the round-bottomed flask which upon trituration with ice-cold chloroform (15 × 2 mL) followed by drying under vacuum gave **6** as a viscous yellow liquid (406.4 mg, 82.4%).

$^1$H NMR (400 MHz, DMSO-d6) δ 5.39 (t, *J* = 4.8 Hz, 1H), 4.50 (s, 2H), 4.10 (s, 1H), 3.88 – 3.81 (m, 2H), 3.67 – 3.57 (m, 2H), 3.52 – 3.49 (m, 2H), 3.14 (s, 3H), 2.80 (t, *J* = 6.7 Hz, 2H).

$^{13}$C NMR (100 MHz, DMSO-d6) δ 83.4, 79.3, 74.2, 72.2, 62.9, 59.7, 54.9, 52.7, 48.6, 13.1.

HRMS (ESI): $C_{10}H_{16}NO^+$; Calculated mass, $[M]^+$ 166.1226; Observed mass $[M]^+$ 166.1234.

**Scheme S6.** Synthesis of benzyl propargyl choline **7**

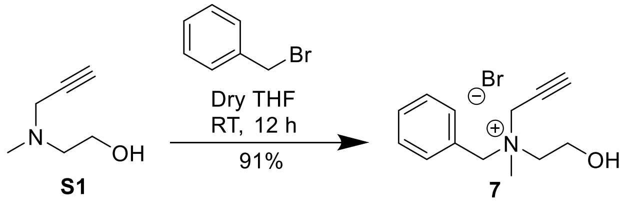


**Benzyl propargyl choline (7)**

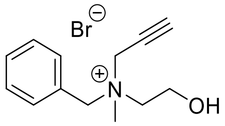


**7**

A round-bottomed flask charged with **S1** (0.3 g, 2.6 mmol, 1 eq.) was purged with nitrogen gas and anhydrous THF (4 mL) was added. This solution was stirred at 0 °C and 80% (*w*/*w*) propargyl bromide in toluene (1.57 mL, 6.6 mmol, 5.0 eq.) was added dropwise under nitrogen atmosphere. After 15 min, this reaction mixture was warmed to room temperature and stirred for another 24 h. The resulting white precipitate was filtered, washed with diethyl ether (10 × 5 mL), and dried under reduced pressure to afford **7** as a white solid (91%).

$^1$H NMR (400 MHz, DMSO-d6) δ 7.64 – 7.50 (m, 5H), 5.44 (t, *J* = 4.8 Hz, 1H), 4.70 (q, *J* = 12.8 Hz, 2H), 4.32 (t, *J* = 2.4 Hz, 2H), 4.17 (t, *J* = 2.4 Hz, 1H), 3.96 – 3.92 (m, 2H), 3.50 – 3.43 (m, 2H), 3.06 (s, 3H).

$^{13}$C NMR (100 MHz, DMSO-d6) δ 133.1, 130.5, 129.1, 127.4, 83.7, 72.6, 65.2, 62.6, 54.7, 51.4, 47.5.

HRMS (ESI): $C_{13}H_{18}NO^+$; Calculated mass, $[M]^+$ 204.1383; Observed $[M]^+$ 204.1390.

**Scheme S7.** Synthesis of diethyl propargyl choline **8**

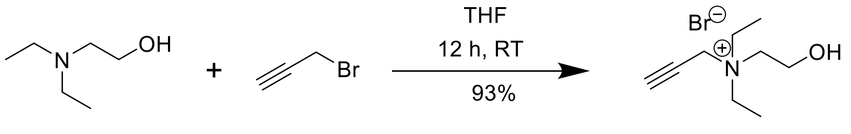


**8**

**Diethyl propargyl choline (8)**

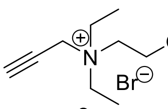

**8**

To an ice-cold solution of 2-diethylaminoethanol (1.13 mL, 8.55 mmol, 1 eq.) in THF (10 mL), 80% (*w*/*w*) propargyl bromide solution in toluene (0.73 mL, 8.53 mmol, 1 eq.) was added dropwise. The reaction mixture was stirred at 0 °C for 30 min and then at room temperature for 12 h to yield a precipitate. This precipitate was collected, washed with ice-cold THF (10 × 10 mL) and then vacuum dried to yield **8** as a dirty-white solid (1.24 g, 93%).

$^{1}$H NMR (400 MHz, DMSO-d6) δ 5.34 (t, *J* = 5.2 Hz, 1H), 4.41 (d, *J* = 2.4 Hz, 2H), 4.03 (t, *J* = 2.8 Hz, 1H), 3.84 – 3.80 (m, 2H), 3.46 – 3.38 (m, 6H), 1.24 (t, *J* = 7.2 Hz, 6H).

$^{13}$C NMR (100 MHz, DMSO-d6) δ 82.5, 72.2, 59.0, 54.5, 48.6, 7.6.

HRMS (ESI): $C_9H_{18}NO^+$, Calculated mass, $[M]^+$ 156.1383; Observed mass, $[M]^+$ 156.1398.

**Scheme S8.** Synthesis of tripropargyl choline **9**

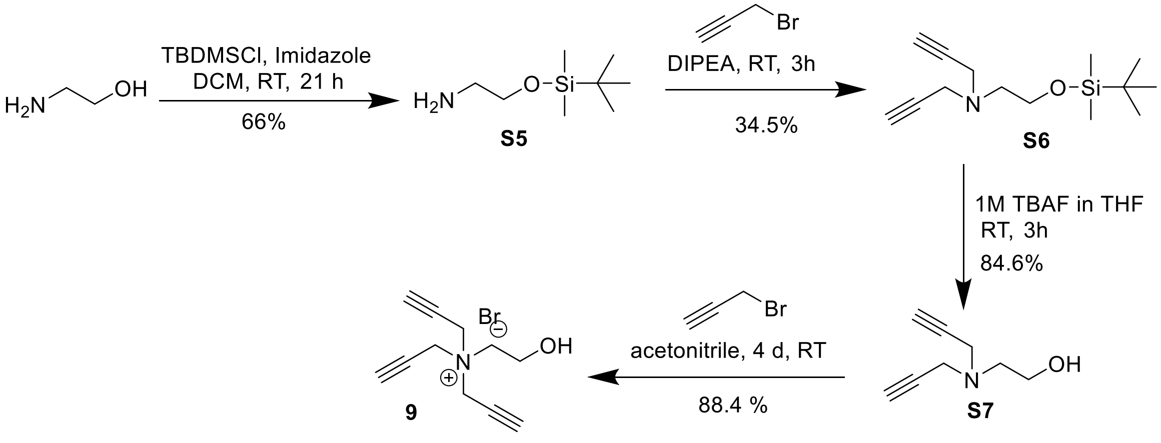


**2-((*t*-butyldimethylsilyl)oxy)ethan-1-amine (S5)**

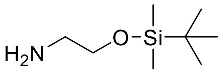

**S5**

To a solution of aminoethanol (4.8 mL, 240 mmol, 1 eq.) in dichloromethane (120 mL), imidazole (17.99 g, 264 mmol, 1.1 eq.) was added. Nitrogen gas was bubbled into this mixture for 30 min and subsequently, the solution was stirred under ice-cold conditions for 15 min. A solution of *t*-butyldimethylsilylchoride (39.78 g, 264 mmol, 1.1 eq.) in dichloromethane (120 mL) was prepared and was also subjected to nitrogen gas treatment as described above. This solution was transferred into the flask containing aminoethanol and imidazole in dichloromethane by using a gas-tight syringe. The resultant turbid mixture was stirred at room temperature for 21 h following which it was washed with water (250 mL). Subsequently, the organic layer was dried by adding anhydrous sodium sulfate, and vacuum dried to afford **S5** as a yellow liquid (15.72 g, 65.6%).

$^{1}$H NMR (500 MHz, $CDCl_3$) δ 3.65 (t, *J* = 5.25 Hz, 2H), 2.94 (s, 2H), 2.81 (t, *J* = 5.25 Hz, 2H), 0.89 (s, 9H), 0.06 (s, 6H).

$^{13}$C NMR (125 MHz, $CDCl_3$) δ 64.5, 44.1, 26.1, 18.5, -5.2.

HRMS (ESI): $C_8H_{21}NOSi$, Calculated mass, $[M+H]^+$ 176.1465; Observed mass, $[M+H]^+$ 176.1470.

**_N_-(2-((_t_-butyldimethylsilyl)oxy)ethyl)-_N_-(prop-2-yn-1-yl)prop-2-yn-1-amine (S6)**

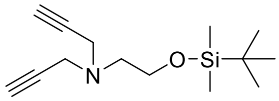


**S6**

To a round-bottomed flask containing **S5** (7.6 g, 40.8 mmol, 1 eq.) dissolved in dichloromethane (50 mL), *N*,*N*-diisopropylethylamine (4.3 mL, 24.5 mmol, 0.6 eq.) was added and the flask was cooled to 0 °C. To this mixture, a solution of 80% *w*/*w* propargyl bromide in toluene (2.64 mL, 24.5 mmol, 0.6 eq.) in dichloromethane (30 mL) was added dropwise and the resultant mixture was stirred at 0 °C for 30 min and then allowed to warm up to room temperature and stirred for 4 h. The orange solution thus obtained was washed with saturated sodium bicarbonate solution (2 × 80 mL), dried by adding anhydrous sodium sulfate, concentrated under reduced pressure and the residue was subjected to silica gel column chromatography (10% ethyl acetate in hexane) to yield **S6** as a yellow liquid (6.16 g, 34.5%).

$^{1}$H NMR (500 MHz, $CDCl_3$) δ 3.75 (t, *J* = 6.05 Hz, 2H), 3.50 (d, *J* = 2.3 Hz, 4H), 2.70 (t, *J* = 6.05 Hz, 2H), 2.21 (t, *J* = 2.3 Hz, 2H), 0.89 (s, 9H), 0.06 (s, 6H).

$^{13}$C NMR (125 MHz, $CDCl_3$) δ 79.2, 73.0, 62.4, 55.2, 43.2, 26.1, 18.4, -5.2.

HRMS (ESI): $C_{14}H_{25}NOSi$, Calculated mass, $[M+H]^+$ 252.1778; Observed mass, $[M+H]^+$ 252.1795.

**2-(di(prop-2-yn-1-yl)amino)ethan-1-ol (S7)**

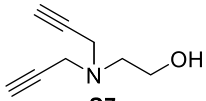


**S7**

To a solution of **S6** (541.1 mg, 2.15 mmol, 1 eq.) in THF (45 mL), tetrabutylammonium fluoride (3.4 mL of a 1 M solution in THF, 3.4 mmol, 1.5 eq.) was added and the resultant orange-colored reaction mixture was stirred at room temperature for 3 h. After removing the solvent under reduced pressure, the residue was subjected to using silica gel column chromatography (5% methanol in ethyl acetate) to yield **S7** as a yellow liquid (250 mg, 84.6%).

$^{1}$H NMR (500 MHz, $CDCl_3$) δ 3.64 (t, *J* = 5.25 Hz, 2H), 3.47 (d, *J* = 2.3 Hz, 4H), 2.74 (t, *J* = 5.2 Hz, 2H), 2.24 (t, *J* = 2.25 Hz, 2H).

$^{13}$C NMR (125 MHz, $CDCl_3$) δ 78.7, 73.4, 58.8, 54.6, 42.3.

HRMS (ESI): $C_8H_{11}NO$; Calculated mass, $[M+Na]^+$ 160.0733 ; Observed $[M+Na]^+$ 160.0753.

**Tripropargyl choline (9)**

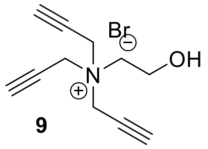


9

To an ice-cold solution of **S7** (180.2 mg, 1.31 mmol, 1 eq.) in acetonitrile (3 mL), 80% *w/w* solution of propargyl bromide in toluene (0.42 mL, 3.93 mmol, 3 eq.) was added dropwise. Stirring this mixture for 4 d at room temperature yielded a white precipitate that was collected and triturated with ice-cold chloroform (10 × 5 mL) which upon drying under reduced pressure gave compound **9** as a white solid (507 mg, 88.4%).

$^{1}$H NMR (500 MHz, DMSO-d6) δ 5.48 (t, *J* = 4.95 Hz, 1H), 4.54 (d, *J* = 2.35 Hz, 6H), 4.16 (t, *J* = 2.3 Hz, 3H), 3.93 – 3.90 (m, 2H), 3.61 (t, *J* = 4.95 Hz, 2H).

$^{13}$C NMR (125 MHz, DMSO-d6) δ 84.1, 71.3, 61.2, 54.6, 50.6.

HRMS (ESI): $C_{11}H_{14}NO^+$; Calculated mass, $[M]^+$ 176.1070; Observed $[M]^+$ 176.1084.

**Scheme S9.** Synthesis of diazirine choline **10**

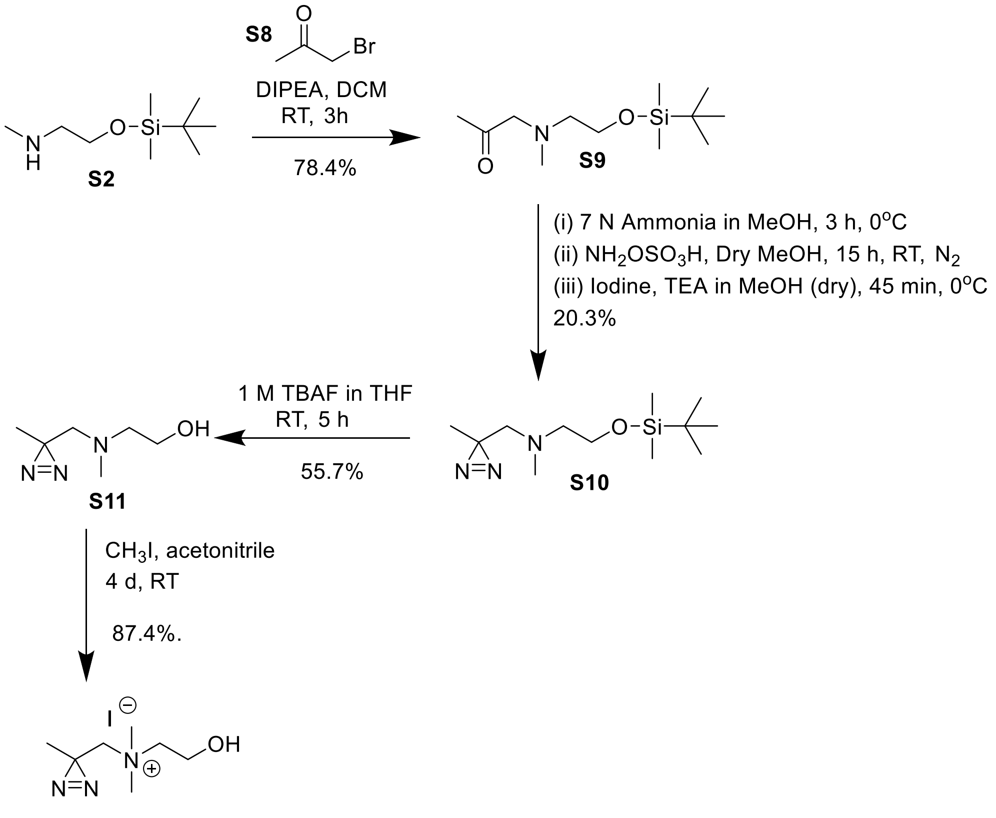


**1-((2-((*t*-butyldimethylsilyl)oxy)ethyl)(methyl)amino)propan-2-one (S9)**

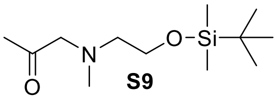


Compound **S8** was synthesized according to a previously reported procedure.[2] **S2** (1.35 g, 7.13 mmol, 1 eq.) and *N*,*N*-diisopropylethylamine (0.46 g, 3.56 mmol, 0.5 eq.) were added to dichloromethane (13 mL) and the mixture was cooled to 0 °C. Subsequently, **S8** (0.488 g, 3.56 mmol, 0.5 eq.) was added dropwise to this solution and the reaction mixture was allowed to warm to room temperature, stirred for 3 h and then washed with saturated sodium bicarbonate solution (2 × 20 mL). The organic extracts were combined, dried over anhydrous sodium sulfate and concentrated under reduced pressure. The resultant residue was subjected to silica gel column chromatography (10% ethyl acetate in hexane) to yield **S9** as a clear yellow liquid (685.6 mg, 78.4%).

$^1$H NMR (500 MHz, $CDCl_3$) δ 3.73 (t, *J* = 5.95 Hz, 2H), 3.30 (s, 2H), 2.58 (t, *J* = 5.95 Hz, 2H), 2.32 (s, 3H), 2.12 (s, 3H), 0.87 (s, 9H), 0.04 (s, 6H).

$^{13}C$ NMR (125 MHz, $CDCl_3$) δ 208.1, 68.3, 61.8, 59.7, 43.7, 27.6, 26.0, 18.4, -5.3.

HRMS (ESI): $C_{12}H_{27}NO_2Si$; Calculated mass, $[M+H]^+$ 246.1884; Observed $[M+H]^+$ 246.1874 .

**2-((*t*-butyldimethylsilyl)oxy)-*N*-methyl-*N*-((3-methyl-3H-diazirin-3-yl)methyl)ethan-1-amine (S10)**

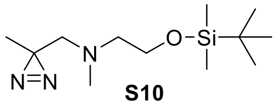


**S10**

A round-bottomed flask charged with **S9** (648.7 mg, 2.64 mmol, 1 eq.) was subjected to high-vacuum for 30 min and cooled to 0 °C. To this flask, an ice-cold solution of 7 N $NH_3$ in methanol (15.5 mL, 710 mmol, 268.9 eq.) was added and the mixture was stirred at 0 °C for 3 h. A solution of hydroxylamine-*O*-sulfonic acid (343.7 mg, 3.04 mmol, 1.2 eq.) in anhydrous methanol (15 mL) that was bubbled with nitrogen gas for 30 min, was added dropwise to the aforementioned solution by using a gas-tight syringe under nitrogen atmosphere. The flask was wrapped with Al-foil and stirred at room temperature for 15 h, resulting in the formation of a heterogeneous mixture of a white precipitate and a clear yellow liquid. Subsequently, the excess ammonia was expelled from the reaction mixture by purging it with nitrogen gas for 2 h. The resultant mixture was then filtered through celite and the retentate was washed with diethyl ether (100 mL). The filtrate was reduced to dryness, subjected to high vacuum for 30 min, purged with nitrogen gas, dissolved in anhydrous methanol (3 mL) and cooled to 0 °C followed by addition of anhydrous triethylamine (552.5µL, 3.96 mmol, 1.5 eq.). After 10 min, iodine beads (737.9 mg, 2.9 mmol, 1.1 eq.) were added and the resultant reaction mixture was stirred at 0 °C for 45 min. Subsequently, the excess iodine was quenched by adding a solution of 10% *w/v* sodium thiosulfate (20 mL) and the mixture was washed with ethyl acetate (1 × 20 mL). The organic layer was separated, dried by adding anhydrous sodium sulfate, concentrated and subjected to silica-gel column chromatography (10% ethyl acetate in hexane) to afford 197.1 mg of **S10** as a yellow liquid (20.3%).

$^{1}H$ NMR (500 MHz, $CDCl_3$) δ 3.69 (t, $J$ = 6.2 Hz, 2H), 2.53 (t, $J$ = 6.2 Hz, 2H), 2.29 (s, 3H), 2.28 (s, 2H), 1.04 (s, 3H), 0.89 (s, 9H), 0.06 (s, 6H).

$^{13}C$ NMR (125 MHz, $CDCl_3$) δ 62.2, 61.5, 59.7, 43.3, 26.1, 25.2, 18.6, 18.4, -5.2.

HRMS (ESI): $C_{12}H_{27}N_3OSi$; Calculated mass, $[M+H]^+$ 258.1996; Observed $[M+H]^+$ 258.2016.

**2-(methyl((3-methyl-3H-diazirin-3-yl)methyl)amino)ethan-1-ol (S11)**

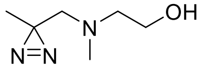


**S11**

A mixture of **S10** (630.4 mg, 2.44 mmol, 1 eq.) and tetrabutylammonium fluoride (3.86 mL of a 1 M solution in THF, 3.86 mmol, 1.5 eq.) in anhydrous THF (44 mL) was stirred at room temperature for 5 h. Subsequently, the solvent was removed under reduced pressure and the residue was subjected to silica-gel column chromatography (2% methanol in ethyl acetate) to yield **S11** as a yellow liquid (195.4 mg, 55.7%).

$^1$H NMR (500 MHz, $CDCl_3$) δ 3.59 (t, *J* = 5.3 Hz, 2H), 2.55 (t, *J* = 5.3 Hz, 2H), 2.31(s, 2H), 2.29 (s, 3H), 1.06 (s, 3H).

$^{13}$C NMR (125 MHz, $CDCl_3$) δ 61.7, 59.0, 58.5, 42.0, 24.9, 18.6.

HRMS (ESI): $C_6H_{13}N_3O$; Calculated mass, $[M+Na]^+$ 166.0951; Observed $[M+Na]^+$ 166.0945.

**Diazirine choline (10)**

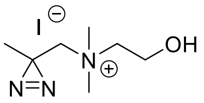


**10**

To a round-bottomed flask containing acetonitrile (6 mL), **S11** (195.4 mg, 1.36 mmol, 1 eq.) and methyl iodide (254.8 µL, 4.09 mmol, 3 eq.) were added and the resultant mixture was stirred at room temperature for 4 d. The brown precipitate thus obtained was triturated with ice-cold chloroform (10 × 2 mL) and, vacuum dried to afford **10** as a deep-yellow solid (339.3 mg, 87%).

$^1$H NMR (400 MHz, DMSO-d6) δ 5.29 (t, *J* = 4.88 Hz, 1H), 3.83 – 3.79 (m, 2H), 3.59 (s, 2H), 3.51 – 3.48 (m, 2H), 3.20 (s, 6H), 1.28 (s, 3H).

$^{13}$C NMR (100 MHz, DMSO-d6) δ 70.0, 66.2, 54.8, 51.5, 21.1, 21.0.

HRMS (ESI): $C_7H_{16}N_3O^+$; Calculated mass $[M]^+$ 158.1288; Observed Mass $[M]^+$ 158.1293.

**Scheme S10.** Synthesis of propargyl diazirine choline **(11)**

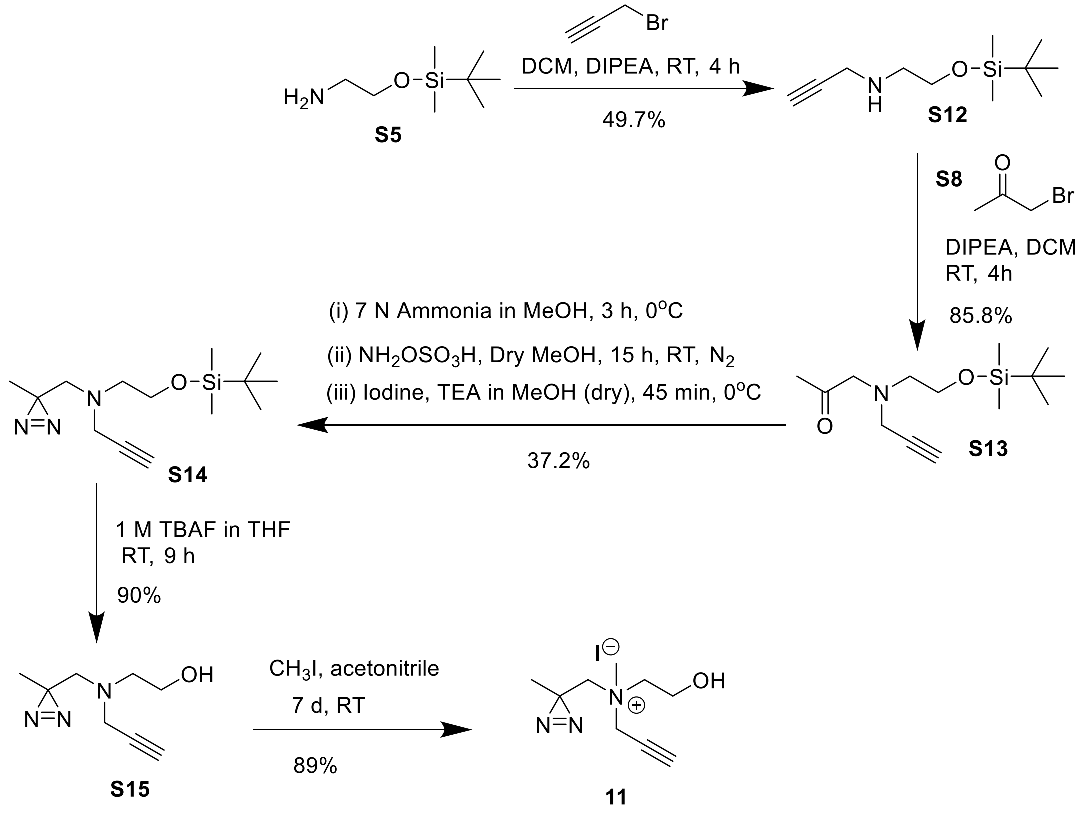


***N*-(2-((*t*-butyldimethylsilyl) oxy)ethyl)prop-2-yn-1-amine (S12)**

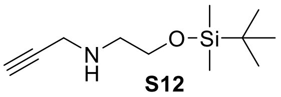


To a round-bottomed flask containing **S5** (15 g, 85.54 mmol, 1 eq.) in dichloromethane (100 mL), *N*,*N*-diisopropylethylamine (7.45 mL, 42.77 mmol, 0.5 eq.) was added and after cooling this solution to 0 °C, a solution of 80% *w*/*w* propargyl bromide in toluene (4.61 mL, 42.77 mmol, 0.5 eq.) in dichloromethane (50 mL) was added dropwise. After stirring the mixture at 0 °C for 30 min, it was warmed up to room temperature and stirred for 4 h. Subsequently, the resulting orange solution was washed with saturated sodium bicarbonate solution (2 × 150 mL), dried by adding anhydrous sodium sulfate, concentrated under reduced pressure and the crude residue was subjected to silica gel column chromatography (30% ethyl acetate in hexane) to yield **S12** as a yellow liquid (4.54 g, 49.7%).

$^1$H NMR (400 MHz, $CDCl_3$) δ 3.73 (t, *J* = 5.2 Hz, 2H), 3.45 (d, *J* = 2.4 Hz, 2H), 2.79 (t, *J* = 5.2 Hz, 2H), 2.20 (t, *J* = 2.4 Hz, 1H), 0.89 (s, 9H), 0.06 (s, 6H).

$^{13}$C NMR (100 MHz, $CDCl_3$) δ 82.3, 71.4, 62.4, 50.7, 38.3, 26.1, 18.4, -5.2.

HRMS (ESI): $C_{11}H_{23}NOSi$; Calculated mass, $[M]^+$ 213.1549; Observed mass, $[M]^+$ 213.1549.

**1-((2-((*t*-butyldimethylsilyl)oxy)ethyl)(prop-2-yn-1-yl)amino)propan-2-one (S13)**

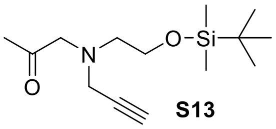


To an ice-cold solution of *N*,*N*-diisopropylethylamine (1.85 mL, 10.59 mmol, 0.5 eq.) and **S12** (4.52 g, 21.18 mmol, 1 eq.) in dichloromethane (51 mL), **S8** (1.45 g, 10.59 mmol, 0.5 eq.) was added dropwise. This mixture was subsequently warmed to room temperature and stirred for 4 h followed by washing with saturated sodium bicarbonate solution (1 × 50 mL). The aqueous layer was given another wash with dichloromethane (1 × 50 mL) and the organic extracts were combined, dried by adding anhydrous sodium sulfate, concentrated under reduced pressure, and subjected to silica gel column chromatography (30% ethyl acetate in hexane) to afford **S13** as a dark-yellow liquid (0.53 g, 90%).

$^{1}$H NMR (400 MHz, $CDCl_3$) δ 3.75 (t, *J* = 6.0 Hz, 2H), 3.52 (d, *J* = 2.4 Hz, 2H), 3.49 (s, 2H), 2.68 (t, *J* = 6.0 Hz, 2H), 2.20 (t, *J* = 2.4 Hz, 1H), 2.14 (s, 3H), 0.88 (s, 9H), 0.05 (s, 6H).

$^{13}$C NMR (100 MHz, $CDCl_3$) δ 207.4, 78.8, 73.3, 64.5, 62.5, 56.4, 44.2, 27.9, 26.1, 18.5, -5.2.

HRMS (ESI): $C_{14}H_{27}NO_2Si$; Calculated mass, $[M]^+$ 269.1811; Observed mass, $[M]^+$ 269.1811.

***N*-(2-((*t*-butyldimethylsilyl)oxy)ethyl)-*N*-((3-methyl-3H-diazirin-3-yl)methyl)prop-2-yn-1-amine (S14)**

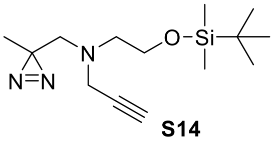


A round-bottomed flask containing **S13** (4.14 g, 15.36 mmol, 1 eq.) was subjected to high-vacuum for 30 min and then cooled on an ice bath. Subsequently, an ice-cold solution of 7 N $NH_3$ in methanol (109 mL, 768 mmol, 50 eq.) was added to this flask and the resulting mixture was stirred at 0 °C for 3 h after which, a solution of hydroxylamine-*O*-sulfonic acid (1.99 g, 17.66 mmol, 1.15 eq.) in anhydrous methanol (88 mL) that had been bubbled with nitrogen gas for 30 min, was added to this reaction mixture by using a gas-tight syringe. The resulting solution was warmed to room temperature and stirred for 15 h, resulting in the formation of a heterogeneous mixture containing a white precipitate and a pale-yellow liquid. Subsequently, ammonia was expelled from the reaction mixture by purging with nitrogen gas for 2 h and the mixture was filtered through a celite plug and washed with diethyl ether (100 mL). The filtrate was reduced to dryness, subjected to high vacuum for 30 min and purged with nitrogen gas. Subsequently, anhydrous methanol (18 mL) and anhydrous triethylamine (3.21 mL, 23.04 mmol, 1.5 eq.) were added and the mixture was cooled to 0 °C. After 10 min, iodine beads (4.29 g, 16.89 mmol, 1.1 eq.)

were added and the reaction mixture was stirred for 45 min under ice-cold conditions. The excess iodine was then quenched with a 10% *w/v* sodium thiosulfate solution (75 mL) and the solution was washed with ethyl acetate (3 × 75 mL). The organic layers were combined, dried by adding anhydrous sodium sulfate, concentrated and subjected to silica gel column chromatography (2% ethyl acetate in hexane) to afford **S14** (1.61 g, 37.2%) as a yellow liquid.

$^1$H NMR (400 MHz, $CDCl_3$) δ 3.70 (t, *J* = 6.0 Hz, 2H), 3.51 (d, *J* = 2.4 Hz, 2H), 2.64 (t, *J* = 6.0 Hz, 2H), 2.44 (s, 2H), 2.15 (t, *J* = 2.4 Hz, 1H), 1.05 (s, 3H), 0.90 (s, 9H), 0.06 (s, 6H).

$^{13}$C NMR (100 MHz, $CDCl_3$) δ 78.5, 73.2, 62.0, 58.2, 55.5, 43.4, 26.1, 24.9, 18.5, 18.4, -5.2.

HRMS (ESI): $C_{14}H_{27}N_3OSi$; Calculated mass, $[M]^+$ 281.1923; Observed mass, $[M]^+$ 281.1923.

**2-(((3-methyl-3H-diazirin-3-yl)methyl)(prop-2-yn-1-yl)amino)ethan-1-ol (S15)**

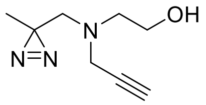


**S15**

**S14** (1.59 g, 5.65 mmol, 1 eq.) and tetrabutylammonium fluoride (8.47 mL of a 1 M solution in THF, 8.47 mmol, 1.5 eq.) were dissolved in anhydrous THF (70 mL) and stirred at room temperature for 9 h. Subsequently, the solvent was evaporated under reduced pressure and the residue was subjected to silica gel column chromatography (20% ethyl acetate in hexane) to afford **S15** (0.85 g, 90%) as a yellow oil.

$^1$H NMR (400 MHz, $CDCl_3$) δ 3.59 (t, *J* = 5.2 Hz, 2H), 3.50 (d, *J* = 2.4 Hz, 2H), 2.69 (t, *J* = 5.2 Hz, 2H), 2.49 (s, 2H), 2.18 (t, *J* = 2.4 Hz, 1H), 1.07 (s, 3H).

$^{13}$C NMR (100 MHz, $CDCl_3$) δ 73.7, 58.6, 57.1, 55.2, 42.1, 24.6, 18.7.

HRMS (ESI): $C_8H_{13}N_3O$; Calculated mass, $[M-N_2+H]^+$ 140.1131; Observed mass, $[M-N_2+H]^+$ 139.9892.

**Propargyl diazirine choline (11)**

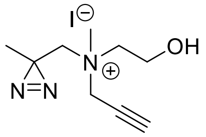


**11**

To a solution of **S15** (0.44 g, 2.62 mmol, 1 eq.) in acetonitrile (4 mL), methyl iodide (0.25 mL, 4.01 mmol, 1.5 eq.) was added and the mixture was stirred at room temperature for 7 d. Each day, methyl iodide (0.25 mL) was added (total of 1.75 mL, 28.1 mmol, 10.5 eq.) to the reaction mixture. Subsequently, the excess solvent was evaporated under reduced pressure and the dark brown residue was triturated with ice-cold chloroform (4 × 5 mL) and vacuum dried to afford **11** as an orange-brown solid (0.72 g, 89%).

$^{1}$H NMR (400 MHz, DMSO-d6) δ 5.37 (t, *J* = 4.8 Hz, 1H), 4.64 – 4.53 (m, 2H), 4.09 (t, *J* = 2.4 Hz, 1H), 3.83 (q, *J* = 5.2 Hz, 2H), 3.83 – 3.56 (m, 4H), 3.24 (s, 3H), 1.29 (s, 3H).

$^{13}$C NMR (100 MHz, DMSO-d6) δ 83.7, 72.0, 68.1, 64.1, 54.7, 52.9, 49.2, 20.8, 20.7.

HRMS (ESI): $C_9H_{16}N_3O^+$; Calculated mass, $[M-N_2]^+$ 154.1288; Observed mass, $[M-N_2]^+$ 154.1239.

**Scheme S11.** Synthesis of azido diazirine choline **12**

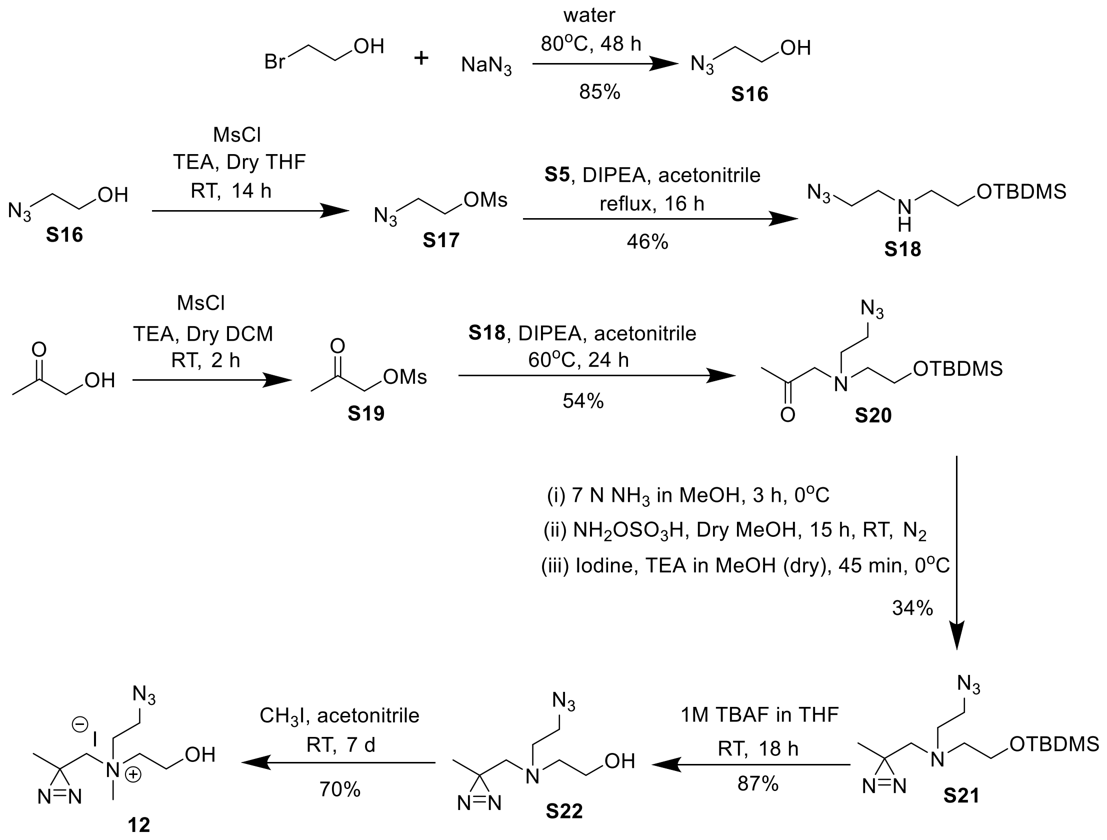


**2-azidoethanol (S16)**

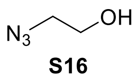


Sodium azide (11.45 g, 176.5 mmol, 1 eq.) was slowly added to an ice-cold solution of 2-bromoethanol (5 mL, 70.5 mmol, 2.5 eq.) in water (10 mL). The solution was then warmed up to room temperature and then heated at 80 °C for 48 h. Subsequently, the reaction mixture was washed with diethyl ether (10 × 10 mL), the organic layers were pooled, dried by adding anhydrous sodium sulfate and concentrated to afford pure **S16** as a colorless oil (85%).

$^{1}$H NMR (400 MHz, $CDCl_3$) δ 3.77 (t, *J* = 5.2 Hz, 2H), 3.44 (t, *J* = 5.2 Hz, 2H), 2.05 (s, 1H).

$^{13}$C NMR (100 MHz, $CDCl_3$) δ 61.6, 53.6.

HRMS (ESI): $C_2H_5N_3O$; Calculated mass for $[M]^+$ 87.0436; Observed mass for $[M]^+$ 87.0761.

**2-azido-*N*-(2-((*t*-butyldimethylsilyl)oxy)ethyl)ethan-1-amine (S18)**

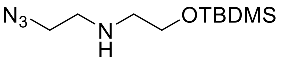


**S18**

**S16** (1.98 g, 22.8 mmol, 1 eq.) was dissolved in dry THF (60 mL) under nitrogen atmosphere and dry triethylamine (6.35 mL, 45.6 mmol, 2 eq.) was added. The resultant mixture was cooled on an ice bath and methanesulfonyl chloride (3.5 mL, 45.6 mmol, 2 eq.) was added. The mixture was stirred at room temperature for 14 h. The reaction mixture was then added to 1 N NaOH (100 mL) and the resultant mixture was washed with dichloromethane (3 × 100 mL), the organic layers were combined, dried by adding anhydrous sodium sulfate, and concentrated under reduced pressure. This concentrated mesylate (**S17**) was treated with a solution of **S5** (4.0 g, 22.8 mmol, 1 eq.) and *N,N*-diisopropylethylamine (4 mL, 22.8 mmol, 1 eq.) in acetonitrile (95 mL). The resultant yellow solution was refluxed for 16 h and then the solvent was evaporated under reduced pressure. The concentrated residue was dissolved in ethyl acetate (100 mL), washed with 1 N NaOH (1 ×100 mL) and the organic layer was dried by adding anhydrous sodium sulfate, concentrated and subjected to silica gel column chromatography (50% ethyl acetate in dichloromethane) to afford **S18** (46.2%) as a pale-yellow oil.

$^{1}$H NMR (400 MHz, $CDCl_3$) δ 3.72 (t, *J* = 5.2 Hz, 2H), 3.42 (t, *J* = 5.6 Hz, 2H), 2.82 (t, *J* = 5.6 Hz, 2H), 2.73 (t, *J* = 5.2 Hz, 2H), 0.90 (s, 9H), 0.06 (s, 6H).

$^{13}$C NMR (100 MHz, $CDCl_3$) δ 62.5, 51.8, 51.6, 48.6, 26.1, 18.5, -5.1.

HRMS (ESI): $C_{10}H_{24}N_4OSi$; Calculated mass $[M+H]^+$ 245.1792; Observed mass $[M+H]^+$ 245.1803.

**1-((2-azidoethyl)(2-((*t*-butyldimethylsilyl)oxy)ethyl)amino)propan-2-one (S20)**

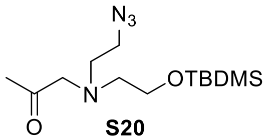


**S20**

A round-bottomed flask containing hydroxyacetone (1.0 mL, 14.76 mmol, 1 eq.) was subjected to high vacuum, flushed with nitrogen gas and dry dichloromethane (67 mL) and dry triethylamine (4.1 mL, 29.46 mmol, 2 eq.) were added. The flask was then cooled to 0 °C and methanesulfonyl chloride (2.3 mL,

29.46 mmol, 2 eq.) was added dropwise under nitrogen atmosphere. The resultant reaction mixture was stirred for 2 h at 0 °C and then washed with 1 N NaOH (60 mL). The organic layer was dried by adding anhydrous sodium sulfate and concentrated under reduced pressure. This concentrated mesylate (**S19)** was added to a solution of **S18** (3.6 g, 14.7 mmol, 1 eq.) and *N,N*-diisopropylethylamine (2.8 mL, 16.2 mmol, 1.1 eq.) in acetonitrile (70 mL) and the resultant solution was stirred for 24 h at 60 °C. Subsequently, the solvent was evaporated under reduced pressure and saturated sodium bicarbonate solution (70 mL) was added to the concentrated residue. The mixture was transferred to a separating funnel and washed with dichloromethane (3 × 70 mL). The organic layers were pooled, dried by adding anhydrous sodium sulfate, concentrated and subjected to silica gel column chromatography (15% ethyl acetate in hexane) to afford **S20** as a yellow oil (2.30 g, 54%).

$^{1}$H NMR (400 MHz, $CDCl_3$) δ 3.71(t, *J* = 5.6 Hz, 2H), 3.49 (s, 2H), 3.30 (t, *J* = 6.0 Hz, 2H), 2.85 (t, *J* = 6.0 Hz, 2H), 2.76 (t, *J* = 5.6 Hz, 2H), 2.14 (s, 3H), 0.88 (s, 9H), 0.05 (s, 6H).

$^{13}$C NMR (100 MHz, $CDCl_3$) δ 208.7, 65.6, 62.4, 56.9, 54.9, 49.9, 27.5, 26.0, 18.4, -5.3.

HRMS (ESI): $C_{13}H_{28}N_4O_2Si$; Calculated mass $[M+H]^+$ 301.2054; Observed mass $[M+H]^+$ 301.2063.

**2-azido-*N*-(2-((*t*-butyldimethylsilyl)oxy)ethyl)-*N*-((3-methyl-3H-diazirin-3-yl)methyl)ethan-1-amine (S21)**

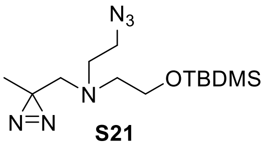


**S21**

A round-bottomed flask containing **S20** (2.12 g, 7.06 mmol, 1 eq.) was subjected to high-vacuum for 30 min and then cooled to 0 °C. To this mixture, an ice-cold solution of 7 N $NH_3$ in methanol (50 mL, 353 mmol, 50 eq.) was added and the mixture was stirred at 0 °C for 3 h. Subsequently, a solution of hydroxylamine-*O*-sulfonic acid (1.19 g, 10.59 mmol, 1.5 eq) in anhydrous methanol (41 mL) that had been bubbled with nitrogen gas for 30 min, was added dropwise via a syringe into the aforementioned mixture and the resultant solution was stirred for 15 h at room temperature. Excess ammonia was expelled by purging the mixture with nitrogen gas for 2 h after which a heterogeneous mixture containing a white precipitate and a yellow liquid was observed. The precipitate was filtered through celite and washed with diethyl ether (100 mL) and the filtrate was concentrated under reduced pressure, kept under high vacuum for 30 min, and purged with nitrogen gas for 30 min. Subsequently, dry methanol (8 mL) and dry triethylamine (1.5 mL, 10.59 mmol, 1.5 eq.) were added and the mixture was cooled to 0 °C. To this flask, iodine beads (1.97 g, 7.77 mmol, 1.1 eq.) were added and the resultant deep-brown colored mixture was stirred for 45 min under ice-cold conditions. Excess iodine was quenched by adding 10% *w/v* sodium thiosulfate solution (20 mL) and the resultant mixture was washed with ethyl acetate (3 × 20 mL). The organic layers were combined, dried by adding anhydrous sodium sulfate, concentrated under reduced pressure to yield a residue that was subjected to silica gel column chromatography (2% ethyl acetate in hexane) to afford **S21** as a yellow liquid (0.77 g, 34%).

$^{1}$H NMR (400 MHz, $CDCl_3$) δ3.69 (t, *J* = 6.0 Hz, 2H), 3.28 (t, *J* = 6.4 Hz, 2H), 2.77 (t, *J* = 6.0 Hz, 2H), 2.70 (t, *J* = 6.0 Hz, 2H), 2.47 (s, 2H), 1.06 (s, 3H), 0.89 (s, 9H), 0.06 (s, 6H).

$^{13}$C NMR (100 MHz, $CDCl_3$) δ 61.6, 59.4, 56.4, 54.5, 49.5, 26.0, 25.5, 18.4, 18.3, -5.3.

HRMS (ESI): $C_{13}H_{28}N_6OSi$; Calculated mass $[M+H]^+$ 313.2166; Observed mass $[M+H]^+$ 313.2176.

**2-((2-azidoethyl)((3-methyl-3H-diazirin-3-yl)methyl)amino)ethan-1-ol (S22)**

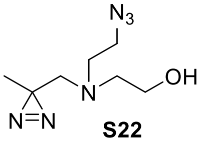


**S22**

**S21** (242 mg, 0.77 mmol, 1 eq.) and tetrabutylammonium fluoride (1.2 mL of a 1 M solution in THF, 1.2 mmol, 1.5 eq.) were added to anhydrous THF (10 mL) and stirred for 18 h at room temperature. Subsequently, the solvent was evaporated under reduced pressure and the residue was subjected to silica gel column chromatography (20% ethyl acetate in dichloromethane) to yield **S22** as a yellow liquid (0.13 g, 87%).

$^{1}$H NMR (400 MHz, $CDCl_3$) δ 3.61 (q, *J* = 5.2 Hz, 2H), 3.36 (t, *J* = 6.0 Hz, 2H), 2.72 (t, *J* = 6.0 Hz, 2H), 2.68 (t, *J* = 5.2 Hz, 2H), 2.49 (s, 2H), 1.09 (s, 3H).

$^{13}$C NMR (100 MHz, $CDCl_3$) δ 59.3, 58.7, 56.6, 53.6, 49.4, 25.2, 18.5.

HRMS (ESI): $C_7H_{14}N_6O$; Calculated mass $[M]^+$ 198.1229; Observed mass $[M]^+$ 198.1308.

**Azido diazirine choline (12)**

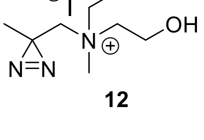


**12**

To a solution of **S22** (3.27 g, 16.5 mmol, 1 eq.) in acetonitrile (12 mL), methyl iodide (1.96 mL, 31.5 mmol, 1.9 eq.) was added and the mixture was stirred at room temperature for 7 d. Each day, methyl iodide (1.96 mL) was added (total of 13.72 mL, 0.22 mol, 13.3 eq.) to the reaction mixture. Subsequently, the solvent was evaporated under reduced pressure and the crude residue was dissolved in water (20 mL) and washed with diethyl ether (10 × 20 mL). The organic layers were combined, dried by adding anhydrous sodium sulfate, and concentrated under reduced pressure to afford **12** as a viscous yellow liquid (70%).

$^{1}$H NMR (400 MHz, DMSO-d6) δ 5.33 (t, *J* = 4.4 Hz, 1H), 3.89 (t, *J* = 6.0 Hz, 2H), 3.78 – 3.83 (m, 2H), 3.68 – 3.65 (m, 4H), 3.58 (t, *J* = 5.2 Hz, 2H), 3.24 (s, 3H), 1.28 (s, 3H).

$^{13}$C NMR (100 MHz, DMSO-d6) δ 68.7, 63.9, 60.9, 54.7, 49.5, 44.0, 21.1, 20.9.

HMRS (ESI): $C_{20}H_{11}N_3O_5$; Calculated mass [M]$^+$ 373.0699; Observed mass [M+H]$^+$ 374.0778.

**Scheme S12.** Synthesis of propargyl ethyl diazirine choline **S26**

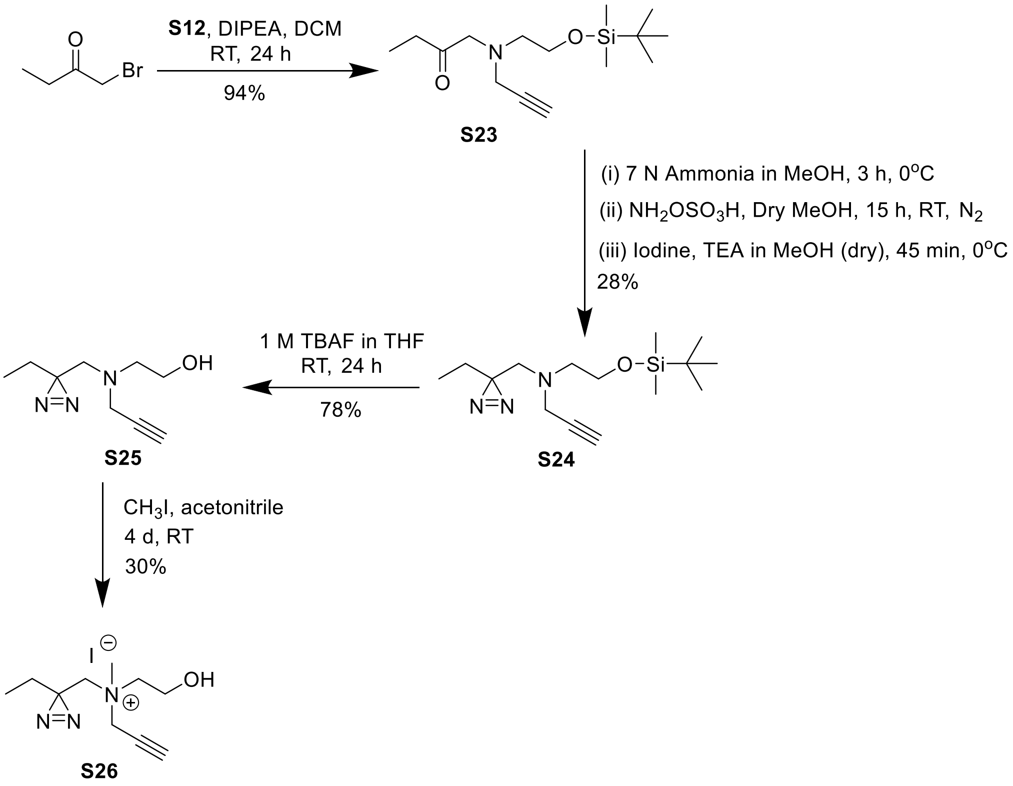


**1-((2-((*t*-butyldimethylsilyl)oxy)ethyl)(prop-2-yn-1-yl)amino)butan-2-one (S23)**

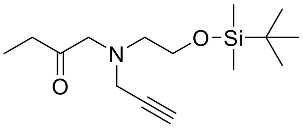

**S23**

To a round-bottomed flask containing an ice-cold solution of **S12** (3.63 g, 17.88 mmol, 1 eq.) in dichloromethane (15 mL), *N*,*N*-diisopropylethylamine (3.80 mL, 21.84 mmol, 1.2 eq.) was added. To this flask, bromobutan-2-one (3 g, 19.86 mmol, 1.1 eq.) was added dropwise and the resultant mixture was stirred at 0 °C for 30 min and then at room temperature for 12 h. The reaction mixture was further diluted with dichloromethane (50 mL) and washed with a saturated solution of sodium bicarbonate (1 × 50 mL).

The aqueous layer was washed with dichloromethane (3 × 50 mL) and all organic layers were combined, dried by adding anhydrous sodium sulfate and concentrated under reduced pressure. The resultant residue was subjected to silica gel column chromatography (8% ethyl acetate in hexane) to give **S23** as a colorless oil (4.70 g, 94%).

$^1$H NMR (400 MHz, $CDCl_3$) δ 3.75 (t, *J* = 6.0 Hz, 2H), 3.52 (d, *J* = 2.4 Hz, 2H), 3.48 (s, 2H), 2.68 (t, *J* = 6.0 Hz, 2H), 2.47 (q, *J* = 7.6 Hz, 2H), 2.20 (t, *J* = 2.4 Hz, 1H), 1.06 (t, *J* = 7.6 Hz, 3H), 0.88 (s, 9H), 0.05 (s, 6H).

$^{13}$C NMR (100 MHz, $CDCl_3$) δ 210.1, 78.9, 73.3, 63.4, 62.4, 56.3, 44.1, 33.6, 26.0, 18.4, 7.9, -5.3.

HRMS (ESI): $C_{15}H_{30}NO_2Si$; Calculated for $[M+H]^+$ 284.2046; Observed for $[M+H]^+$, 284.2051

**N-(2-((t-butyldimethylsilyl)oxy)ethyl)-N-((3-ethyl-3H-diazirin-3-yl)methyl)prop-2-yn-1-amine (S24)**

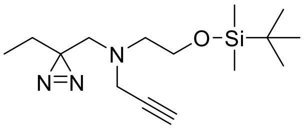


**S24**

A round-bottomed flask charged with **S23** (0.5 g, 1.76 mmol, 1 eq.) was kept under high vacuum for 30 min and cooled to 0 °C. Subsequently, an ice-cold solution of 7 N $NH_3$ in methanol (12.5 mL, 87.5 mmol, 50 eq.) was added and the mixture was stirred at 0 °C for 3 h. To this mixture, a solution of hydroxylamine-*O*-sulfonic acid (0.39 g, 3.26 mmol, 1.9 eq.) in anhydrous methanol (5 mL) that had been bubbled with nitrogen gas for 30 min, was added dropwise via a syringe and the reaction mixture was stirred at room temperature for 15 h. Subsequently, excess ammonia was removed by bubbling nitrogen gas for 2 h, and the heterogeneous mixture containing a white precipitate and a yellow solution thus obtained, was filtered on celite and washed with diethyl ether (100 mL). The filtrate was evaporated to dryness, subjected to high vacuum for 30 min and purged with nitrogen gas. Subsequently, dry methanol (15 mL) and dry triethylamine (0.37 mL, 2.65 mmol, 1.5 eq.) were added and the solution was cooled to 0 °C followed by addition of iodine (0.49 g, 1.94 mmol, 1.1 eq.) in small portions and the mixture was stirred at 0 °C for 45 min. Subsequently, ethyl acetate (50 mL) was added and the mixture was washed with 1 N HCl (1 × 50 mL), and a saturated solution of sodium bicarbonate (1 × 50 mL). The organic layer was dried by adding anhydrous sodium sulfate and concentrated under reduced pressure to give a residue which was subjected to silica gel column chromatography (2% ethyl acetate in hexane) to give **S24** as a pale-yellow oil (0.11 g, 21%).

$^1$H NMR (400 MHz, $CDCl_3$) δ 3.68 (t, *J* = 6.0 Hz, 2H), 3.50 (d, *J* = 2.4 Hz, 2H), 2.63 (t, *J* = 6.0 Hz, 2H), 2.45 (s, 2H), 2.15 (t, *J* = 2.4 Hz, 1H), 1.48 (q, *J* = 8.0 Hz, 2H), 0.89 (s, 9H), 0.69 (t, *J* = 6.0 Hz, 3H), 0.06 (s, 6H).

$^{13}$C NMR (100 MHz, $CDCl_3$) δ 78.4, 73.0, 61.8, 56.7, 55.4, 43.2, 28.5, 25.9, 24.3, 18.3, 8.0, -5.4.

HRMS (ESI): $C_{15}H_{30}N_3OSi$; Calculated mass $[M+Na]^+$ 296.2158; Observed mass $[M+Na]^+$ 296.2152.

**2-(((3-ethyl-3H-diazirin-3-yl)methyl)(prop-2-yn-1-yl)amino)ethan-1-ol (S25)**

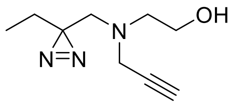

**S25**

**S24** (1.2 g, 4.06 mmol, 1 eq.) and tetrabutylammonium fluoride (6.1 mL of a 1 M solution in THF, 6.09 mmol, 1.5 eq.) was added to dry THF (5 mL) and stirred for 24 h at room temperature. Subsequently, the solvent was removed under reduced pressure and the concentrated residue was subjected to silica gel column chromatography (5% ethyl acetate in dichloromethane) to give **S25** as a colorless oil (0.58 g, 78%).

$^1$H NMR (400 MHz, $CDCl_3$) δ 3.58 (t, $J$ = 5.2 Hz, 2H), 3.48 (d, $J$ = 2.4 Hz, 2H), 2.68 (t, $J$ = 5.2 Hz, 2H), 2.50 (s, 2H), 2.18 (t, $J$ = 2.4 Hz, 1H), 1.51 (q, $J$ = 7.6 Hz, 2H), 0.70 (t, $J$ = 7.6 Hz, 3H).

$^{13}$C NMR (100 MHz, $CDCl_3$) δ 73.7, 58.7, 55.7, 55.2, 42.2, 28.4, 24.6, 8.2, 0.1.

HRMS (ESI): $C_9H_{15}N_3O$; Calculated mass $[M]^+$ 182.1215; Observed mass $[M]^+$ 182.1289.

**Propargyl ethyl diazirine choline (S26)**

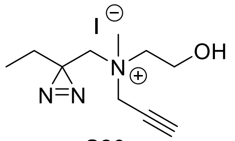

**S26**

To a solution of **S25** (0.56 g, 3.10 mmol, 1 eq.) in acetonitrile (3 mL), methyl iodide (1.15 mL, 18.4 mmol, 6 eq.) was added and the mixture was stirred at room temperature for 4 d. Each day, methyl iodide (1.15 mL) was added (total of 4.6 mL, 74 mmol, 24 eq.) to the reaction mixture. Subsequently, the solvent was removed by evaporation under reduced pressure and the resultant residue was subjected to reverse phase flash column chromatography by using a Redisep Rf 43 g C18 column with water-acetonitrile as the mobile phase (gradient: 0-20 min (100% water), 20-30 min (100-0% water), 30-35 min (0% water), 35-40 min (0-100% water); flow rate = 5 mL/min) to give pure column fractions containing **S26**. These fractions were concentrated under reduced pressure to afford **S26** as a colorless liquid (0.3 g, 30%).

$^1$H NMR (400 MHz, $D_2O$) δ 4.51 (s, 2H), 4.03 – 4.01 (m, 2H), 3.73 (t, $J$ = 4.8 Hz, 2H), 3.69 (d, $J$ = 5.2 Hz, 2H), 3.35 (s, 3H), 1.85 (q, $J$ = 7.6 Hz, 2H), 0.65 (t, $J$ = 7.6 Hz, 3H).

$^{13}$C NMR (100 MHz, DMSO-d6) δ 83.7, 71.9, 70.0, 64.1, 54.7, 52.8, 49.2, 25.2, 24.7, 7.7.

HRMS (ESI): $C_{10}H_{18}N_3O$; Calculated mass $[M]^+$ 196.1444; Observed mass $[M]^+$ 196.1454.

**Scheme 13.** Synthesis of propargyl methyl diazirine choline **13**

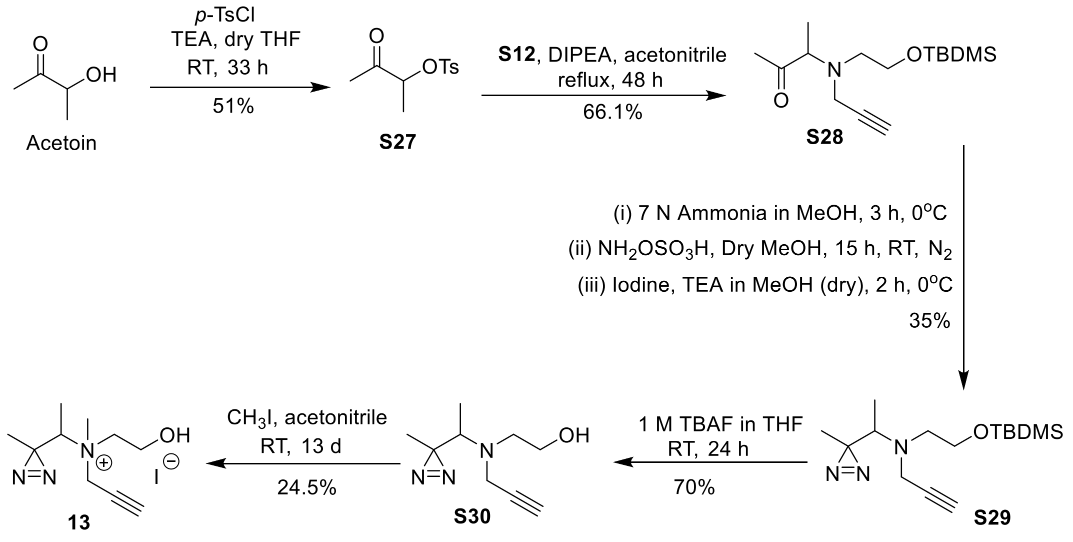


#### 3-oxobutan-2-yl 4-methylbenzenesulfonate (S27)

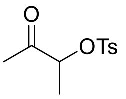


**S27**

A round-bottomed flask containing a mixture of acetoin (6 g, 68 mmol, 1 eq.) and *p*-toluenesulfonyl chloride was flushed with nitrogen gas and cooled in an ice bath. Dry THF (70 mL) was added and the mixture was stirred on the ice bath for 15 min. Dry triethylamine (18.9 mL, 136 mmol, 2 eq.) was then added and the resultant mixture was allowed to stir at 0 °C for another 10 min and then at room temperature for 33 h. Subsequently, the solvent was removed under reduced pressure and the concentrated residue was dissolved in ethyl acetate (100 mL). This solution was washed with 1 N HCl (1 × 100 mL), saturated solution of sodium bicarbonate (1 × 100 mL), and brine (1 × 100 mL). The organic layer was concentrated under reduced pressure and the residue was subjected to silica gel column chromatography (10% ethyl acetate in hexane) to afford **S27** as a yellow liquid (16.5 g, 51%).

$^1$H-NMR (400Hz, $CDCl_3$) δ 7.81 (d, *J* = 8.4 Hz, 2H), 7.36 (d, *J* = 8.8 Hz, 2H), 4.75 (q, *J* = 6.8 Hz, 1H), 2.46 (s, 3H), 2.21 (s, 3H), 1.34 (d, *J* = 6.8 Hz, 3H).

$^{13}$C NMR (100 MHz, $CDCl_3$) δ 205.4, 145.5, 133.3, 130.2, 128.0, 81.0, 25.7, 21.8, 17.4.

HRMS (ESI): $C_{11}H_{14}O_4S$; Calculated mass, $[M+H]^+$ 243.0686; Observed mass, $[M+H]^+$ 243.0695.

**3-((2-((*t*-butyldimethylsilyl)oxy)ethyl)(prop-2-yn-1-yl)amino)butan-2-one (S28)**

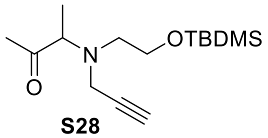


**S28**

*N*,*N*-diisopropylethylamine (3.16 mL, 18.15 mmol, 1.1 eq.) was added to a solution of **S12** (4 g, 16.5 mmol, 1 eq.) in acetonitrile (40 mL). To this flask, **S27** (3.52 g, 16.5 mmol, 1 eq.) was added dropwise and the mixture was refluxed for 48 h at room temperature. Subsequently, the solvent was removed under reduced pressure and the concentrated residue was dissolved in dichloromethane (50 mL) and washed with saturated sodium bicarbonate solution (50 mL). The aqueous layer was further washed with dichloromethane (2 × 50 mL). All the organic layers were pooled, dried by adding anhydrous sodium sulfate, and concentrated under reduced pressure. The concentrated residue was subjected to silica gel column chromatography (5% ethyl acetate in hexane) to afford **S28** as a dark yellow liquid (4.67 g, 66.1%).

$^{1}$H-NMR (400Hz, $CDCl_3$) δ 3.70 (t, *J* = 6.4 Hz, 2H), 3.54 – 3.42 (m, 3H), 2.71 – 2.5 (m, 2H), 2.23 (s, 3H), 2.19 (t, *J* = 2.4 Hz, 1H), 1.18 (d, *J* = 7.2 Hz, 3H), 0.89 (s, 9H), 0.05 (s, 6H).

$^{13}$C NMR (100 MHz, $CDCl_3$) δ 211.5, 79.8, 72.8, 66.7, 62.1, 52.9, 40.6, 26.7, 25.9, 18.3, 11.6, -5.4.

HRMS (ESI): $C_{15}H_{29}NO_2Si$; Calculated mass, $[M+H]^+$ 284.2040, Observed mass, $[M+H]^+$ 284.2047.

***N*-(2-((*t*-butyldimethylsilyl)oxy)ethyl)-*N*-(1-(3-methyl-3H-diazirin-3-yl)ethyl)prop-2-yn-1-amine (S29)**

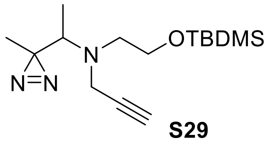


**S29**

A round-bottomed flask charged with **S28** (2.84 g, 10 mmol, 1 eq.) was kept under high vacuum for 30 min and cooled to 0 °C. Subsequently, an ice-cold solution of 7 N ammonia in methanol (72 mL, 500 mmol, 50 eq.) was added and the mixture was stirred at 0 °C for 3 h. To this flask, a solution of hydroxylamine-*O*-sulfonic acid (1.69 g, 15 mmol, 1.5 eq.) in dry methanol (75 mL) that had been bubbled with nitrogen gas for 30 min, was added dropwise via a syringe and the resultant mixture was stirred for 15 h at room temperature. Subsequently, excess ammonia was purged out by bubbling nitrogen gas for 2 h, and the heterogeneous mixture of a white precipitate and a yellow liquid thus obtained was filtered on celite and washed with diethyl ether (100 mL). The filtrate was evaporated to dryness, kept under high vacuum for 30 min, and was purged with nitrogen gas. To this flask, dry methanol (12 mL) and dry triethylamine (2.1 mL, 15 mmol, 1.5 eq.) were added and the mixture was cooled to 0 °C. After 10 min, iodine (2.79 g, mmol, 1.1 eq.) was added in small portions and the reaction mixture was stirred for 2 h at 0 °C. The reaction was quenched by adding a solution of 10% *w*/*v* sodium thiosulfate (50 mL) and turbid solution thus obtained was washed with ethyl acetate (3 × 50 mL). The organic layers were combined, dried by adding anhydrous sodium sulfate and concentrated under reduced pressure. The resultant residue was subjected to silica gel column chromatography (2% ethyl acetate in hexane) to yield **S29** as a yellow liquid (0.95 g, 35%).

$^{1}$H-NMR (400Hz, $CDCl_3$) δ3.75 (td, *J* = 6.2 Hz, 2.0 Hz, 2H). 3.65 (dd, *J* = 17.6 Hz, 2.4 Hz, 1H), 3.48 (dd, *J* = 17.2 Hz, 2 Hz, 1H), 2.81 – 2.77 (m, 2H), 2.48 (q, *J* = 6.8 Hz, 1H), 2.14 (t, *J* = 2.4 Hz, 1H), 1.00 (s, 3H), 0.90 (s, 9H), 0.88 (d, *J* = 6.8 Hz, 3H), 0.08 (s, 6H).

$^{13}$C NMR (100 MHz, $CDCl_3$) δ 79.9, 72.7, 62.4, 59.8, 52.1, 40.3, 28.7, 26.1, 18.5, 17.2, 12.6, -5.2.

HRMS (ESI): $C_{15}H_{29}N_3OSi$; Calculated mass, $[M\text{-}N_2+H]^+$ 268.2152; Observed mass, $[M\text{-}N_2+H]^+$ 268.2101.

**2-((1-(3-methyl-3H-diazirin-3-yl)ethyl)(prop-2-yn-1-yl)amino)ethan-1-ol (S30)**

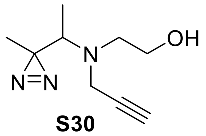


**S30**

To a solution of **S29** (950 mg, 3.21 mmol, 1 eq.) in dry THF (30 mL), tetrabutylammonium fluoride (4.8 mL of a 1 M solution in THF, 4.8 mmol, 1.5 eq.) was added and the mixture was stirred at room temperature for 24 h to give an orange-colored solution. After removing the solvent under reduced pressure, the residue was subjected to silica gel column chromatography (10% ethyl acetate in dichloromethane) to yield **S30** (583 mg, 69%) as a yellow liquid.

$^{1}$H-NMR (400Hz, $CDCl_3$) δ 3.66 (q, *J* = 5.2 Hz, 2H), 3.55 (dd, *J* = 17.6 Hz, 2.4 Hz, 1H), 3.45 (dd, *J* = 17.8 Hz, 2.0 Hz, 1H), 2.86 (t, *J* = 5.2 Hz, 2H), 2.63 (q, *J* = 6.8 Hz, 1H), 2.38 (t, *J* = 5.6 Hz, 1H), 2.19 (t, *J* = 2.4 Hz, 1H), 1.03 (s, 3H), 0.89 (d, *J* = 6.8 Hz, 3H).

$^{13}$C NMR (100 MHz, $CDCl_3$) δ 79.3, 73.3, 59.8, 58.9, 51.6, 38.8, 28.2, 17.7, 12.6.

HRMS (ESI): $C_9H_{15}N_3O$; Calculated mass, $[M+Na]^+$ 204.1107; Observed mass, $[M+Na]^+$ 204.1590.

**Propargyl methyl diazirine choline (13)**

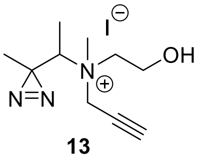


**13**

To a solution of **S30** (0.39 g, 2.15 mmol, 1 eq.) in acetonitrile (10 mL), methyl iodide (0.27 mL, 4.34 mmol, 2 eq.) was added and the mixture was stirred at room temperature for 13 d. Each day, methyl iodide (0.27 mL) was added (total of 3.51 mL, 56.4 mmol, 26 eq.) to the reaction mixture. Subsequently, the solvent was evaporated under reduced pressure and the residue was subjected to reverse phase flash column chromatography by using a Redisep Rf 43 g C18 column with water-acetonitrile as the mobile phase (gradient: 0-20 min (100% water), 20-30 min (100-0% water), 30-35 min (0% water), 35-40 min (0-100% water); flow rate = 5 mL/min) to isolate **13**. The fractions containing **13** were concentrated under reduced pressure to yield a colorless oil (0.14 g, 24.5%).

$^1$H NMR (400 MHz, $D_2O$) δ 4.73 – 4.38 (m, 2H), 4.12 – 4.05 (m, 2H), 3.90 – 3.77 (m, 2H), 3.70 – 3.54 (m, 1H), 3.29 – 3.23 (m, 4H), 1.49 (d, $J$ = 6.8 Hz, 3H), 1.28 (s, 3H).

$^{13}$C NMR (100 MHz, DMSO-d6) δ 83.7, 83.6, 73.1, 73.0, 72.3, 72.1, 61.8, 61.7, 54.9, 54.6, 52.8, 51.5, 51.4, 46.6, 24.4, 24.3, 19.23, 19.22, 11.2, 11.0.

HRMS (ESI): $C_{10}H_{18}N_3O^+$; Calculated mass, $[M]^+$ 196.1444; Observed mass, $[M]^+$ 196.1450.

*Note:* Compound **13** was purified as a racemic mixture and the H atoms on the $CH_2$ group between the quaternary nitrogen atom and the triple bonded carbon atom of this compound are diastereotopic resulting in the appearance of sets of two closely spaced peaks in the $^1$H and $^{13}$C NMR spectra.

**Scheme S14. Synthesis of γ-diazirine choline (17)**

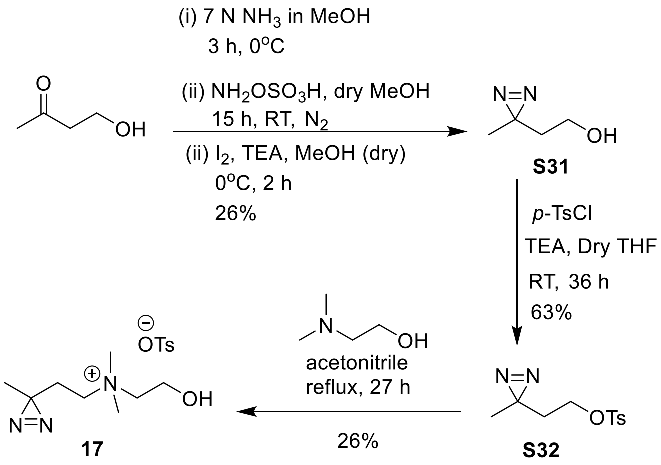


### 2-(3-methyl-3H-diazirin-3-yl)ethan-1-ol (S31)

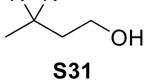

**S31**

A two-necked round-bottomed flask charged with 4-hydroxy-2-butanone (6.13 g, 69.57 mmol, 1 eq.) was kept under high vacuum for 30 min and then cooled to 0 °C. Subsequently, an ice-cold solution of 7

N $NH_3$ in methanol (495 mL, 3.47 mol, 50 eq.) was added and the mixture was stirred at 0 °C for 3 h. To this mixture, a solution of hydroxylamine-*O*-sulfonic acid (11.81 g, 104.50 mmol, 1.5 eq.) in anhydrous methanol (80 mL) that was bubbled with nitrogen gas for 30 min, was added dropwise by using a gas-tight syringe and the resultant reaction mixture was warmed to room temperature and stirred for another 15 h. Subsequently, excess ammonia was purged out by bubbling nitrogen gas for 2 h, and the heterogeneous mixture of a white precipitate and a pale-yellow liquid thus obtained was filtered on celite and washed with diethyl ether (100 mL). The filtrate was evaporated to dryness, kept under high vacuum for 30 min and purged with nitrogen gas. To this flask, dry methanol (60 mL) and dry triethylamine (14.56 mL, 104.3 mmol, 1.5 eq.) were added and the mixture was cooled to 0 °C. Subsequently, iodine (19.44 g, 76.62 mmol, 1.1 eq.) was added in small portions and the reaction mixture was stirred for 2 h at 0 °C followed by the addition of ethyl acetate (100 mL) and the mixture was washed with 1 N HCl (1 × 100 mL), and saturated sodium thiosulfate solution (1 × 100 mL). The organic layer was dried over anhydrous sodium sulfate, concentrated under reduced pressure and the residue was subjected to purification by silica gel column chromatography (30% ethyl acetate in hexane) to give **S31** as a yellow oil (1.83 g, 26%).

$^1$H NMR (400 MHz, $CDCl_3$) δ 3.53 (t, *J* = 6.4 Hz, 2H), 1.63 (t, *J* = 6.4 Hz, 2H), 1.06 (s, 3H).

$^{13}$C NMR (100 MHz, $CDCl_3$) δ 57.6, 37.0, 24.3, 20.2.

HRMS (ESI): $C_4H_8N_2O$; Calculated mass, $[M+H]^+$ 101.0709; Observed mass, $[M+H]^+$ 101.0711.

**2-(3-methyl-3H-diazirin-3-yl)ethyl 4-methylbenzenesulfonate (S32)**

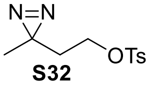


**S32**

A round-bottomed flask containing a mixture of **S31** (1.6 g, 15.98 mmol, 1 eq.) and *p*-toluenesulfonyl chloride (9.14 g, 47.94 mmol, 3 eq.) was flushed with nitrogen gas and cooled to 0 °C. Subsequently, anhydrous THF (30 mL) was added and the mixture was stirred for 15 min. Dry triethylamine (6.68 mL, 47.94 mmol, 3 eq.) was then added and the reaction n mixture was allowed to stir at 0 °C for another 10 min and then at room temperature for 36 h. Subsequently, THF was removed under reduced pressure and ethyl acetate (50 mL) was added to the concentrated residue. This solution was washed with 1 N HCl (1 × 50 mL), saturated solution of sodium bicarbonate (1 × 50 mL), and concentrated under reduced pressure after being dried by adding anhydrous sodium sulfate. The product was isolated from the concentrated residue using silica gel column chromatography (10% ethyl acetate in hexane) to give **S32** (2.58 g, 63%) as a pale-yellow oil.

$^{1}H$ NMR (400 MHz, $CDCl_3$) δ 7.83 (d, *J* = 8.4 Hz, 2H), 7.37 (d, *J* = 8.4 Hz, 2H), 3.95 (t, *J* = 6.4 Hz, 2H), 2.46 (s, 3H), 1.67 (t, *J* = 6.4 Hz, 2H), 1.00 (s, 3H).
$^{13}C$ NMR (100 MHz, $CDCl_3$) δ 145.5, 133.3, 130.4, 128.4, 65.5, 34.7, 23.8, 22.1, 20.3.
HRMS (ESI): $C_{11}H_{14}N_2O_3S$; Calculated mass, $[M+H]^+$ 255.0798; Observed mass, $[M+H]^+$ 255.0801.

**γ-diazirine choline (17)**

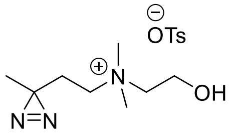


**17**

An ice-cold solution of **S32** (80 mg, 0.31 mmol, 1 eq.) in acetonitrile (1 mL) was added to an ice-cold solution of *N*,*N*-dimethylaminoethanol (23.4 mg, 0.26 mmol, 3 eq.) dissolved in acetonitrile (1 mL) and after 10 min of stirring, the reaction mixture was refluxed for 27 h followed by removal of the solvent under reduced pressure. The remaining crude residue was triturated with ice-cold diethyl ether (5 × 2 mL) to yield 98 mg of **17** (26%) as dirty-white solid.
$^{1}H$ NMR (500 MHz, $CD_3OD$) δ 7.72 (d, *J* = 8.2 Hz, 2H), 7.25 (d, *J* = 7.9 Hz, 2H), 3.96 – 3.94 (m, 2H), 3.49 – 3.46 (m, 2H), 3.43 – 3.41 (m, 2H), 3.12 (s, 6H), 2.37 (s, 3H), 1.87 – 1.83 (m, 2H), 1.08 (s, 3H).
$^{13}C$ NMR (175 MHz, $CD_3OD$) δ 143.6, 141.7, 129.8, 127.0, 66.5, 61.4, 56.8, 52.3, 29.3, 24.4, 21.3, 19.3.
HRMS (ESI): $C_8H_{18}N_3O^+$; Calculated mass, $[M]^+$ 172.1444; Observed mass, $[M]^+$ 172.1444.

**Scheme S15.** Synthesis of propargyl γ-diazirine choline **18**

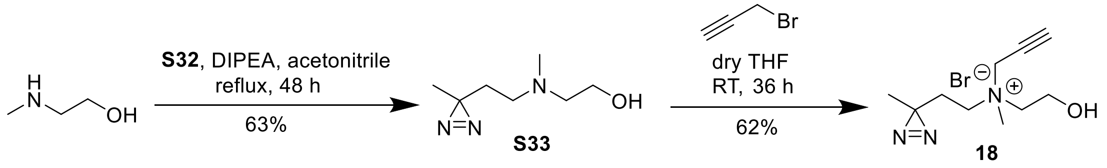


**2-(methyl(2-(3-methyl-3H-diazirin-3-yl)ethyl)amino)ethan-1-ol (S33)**

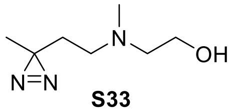


**S33**

To an ice-cold mixture of *N*-methylaminoethanol (0.44 g, 5.87 mmol, 1.5 eq.) and *N*,*N*-diisopropylethylamine (0.56 g, 4.31 mmol, 1.1 eq.) in acetonitrile (5 mL), a solution of **S32** in acetonitrile (2 mL) was added dropwise. The resultant mixture was stirred under ice-cold conditions for 30 min and then refluxed for 48 h. After this, the solvent was removed under reduced pressure and the residue was dissolved in dichloromethane (50 mL) and washed with saturated solution of sodium bicarbonate (1 ×50 mL). The resultant aqueous layer was further washed with dichloromethane (3 × 50 mL). The organic layers were pooled, dried by adding anhydrous sodium sulfate, and concentrated under reduced pressure. The concentrated residue was subjected to silica gel column chromatography (1% triethylamine in ethyl acetate) to afford **S33** as a pale-yellow liquid (0.39 g, 63%).

$^1$H NMR (500 MHz, $CDCl_3$) δ 3.56 (t, *J* = 4.0 Hz, 2H), 2.46 (t, *J* = 4.0 Hz, 2H), 2.27 (t, *J* = 6.0 Hz, 2H), 2.18 (s, 3H), 1.55 (t, *J* = 6.0 Hz, 2H), 1.02 (s, 3H).

$^{13}$C NMR (100 MHz, $CDCl_3$) δ 58.8, 58.5, 52.3, 41.2, 31.9, 25.0, 20.3.

HRMS (ESI): $C_4H_{15}N_3O^+$; Calculated mass, $[M+H]^+$ 158.1288; Observed mass, $[M+H]^+$ 158.1297.

**Propargyl γ-diazirine choline (18)**

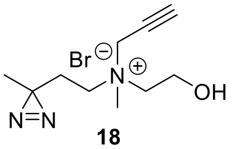

**18**

To an ice-cold solution of **S33** (368.3 mg, 2.34 mmol, 1.0 eq.) in dry THF (20 mL), a solution of 80% *w*/*w* propargyl bromide in toluene (1.31 mL, 11.71 mmol, 5.0 eq.) was added dropwise. The mixture was stirred at 0 °C for 30 min and then at room temperature for 36 h. The yellow precipitate thus obtained was isolated by decantation followed by trituration with THF (5 × 10 mL), diethyl ether (5 × 25 mL), and ethyl acetate (7 × 25 mL) and dried under vacuum to give **18** as a dirty-white solid (0.65 g, 62%).

$^1$H NMR (400 MHz, DMSO-d6) δ 5.37 (t, *J* = 4.8 Hz, 1H), 4.42 (d, *J* = 2.4 Hz, 2H), 4.10 (t, *J* = 2.4 Hz, 1H), 3.83 – 3.82 (m, 2H), 3.52 – 3.48 (m, 2H), 3.44 – 3.42 (m, 2H), 3.08 (s, 3H), 1.83 – 1.79 (m, 2H), 1.06 (s, 3H).

$^{13}$C NMR (100 MHz, DMSO-d6) δ 83.3, 72.1, 62.8, 56.8, 54.7, 52.3, 48.6, 27.5, 24.0, 19.0.

HRMS (ESI):$C_{10}H_{18}N_3O^+$, Calculated mass, $[M]^+$ 196.1444; Observed mass, $[M]^+$ 196.1454.

**Scheme S16.** Synthesis of rhodamine alkyne **19**

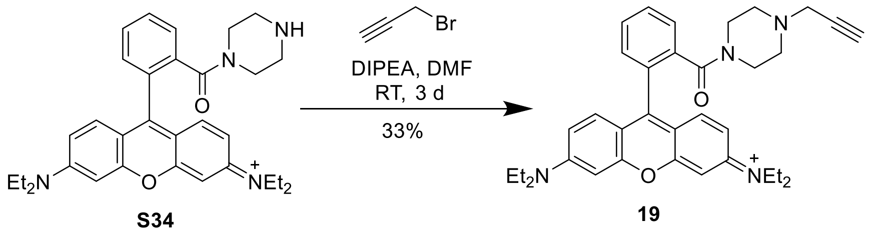


**Rhodamine alkyne (19)**

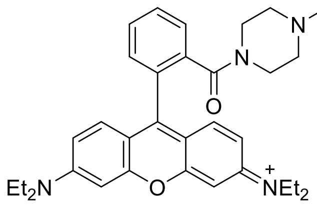


**19**

Rhodamine B piperazine amide (**S34)** was synthesized according to a previously reported method.[3] To an Al-foil wrapped round-bottomed flask charged with **S34** (100 mg, 0.18 mmol, 1 eq.) in DMF (1 mL), a solution of 80% *w/w* propargyl bromide in toluene (75 µL, 0.69 mmol, 3.75 eq.) and *N,N*-diisopropylethylamine (47 µL, 0.27 mmol, 1.5 eq.) in DMF (1 mL) was added dropwise and the mixture was stirred at room temperature for 3 d. Subsequently, the solvent was removed under reduced pressure and the concentrated residue was dissolved in ethyl acetate (50 mL) and washed with saturated sodium bicarbonate solution (1 × 50 mL). The aqueous layer was washed with ethyl acetate (3 × 50 mL) and 1:3 *v/v* solution of isopropanol and dichloromethane (15 × 50 mL). The organic layers were combined, dried by adding anhydrous sodium sulfate and concentrated under reduced pressure. The residue was subjected to silica gel column chromatography (10% methanol in dichloromethane) to afford 35.1 mg of **19** as a deep purple solid (33%).

$^1$H NMR (400 MHz, $CDCl_3$) δ 7.69 – 7.67 (m, 2H), 7.56 – 7.54 (m, 1H), 7.38 – 7.35 (m, 1H), 7.28 (s, 1H), 7.27 (s, 1H), 7.01 – 6.98 (m, 2H), 6.84 (d, *J* = 2.4 Hz, 2H), 3.70 – 3.6 (m, 8H), 3.45 – 3.35 (m, 4H), 3.25 (d, *J* = 2.4 Hz, 2H), 2.38 – 2.35 (m, 4H), 2.26 (t, *J* = 2.4 Hz, 1H), 1.34 (t, *J* = 7.2 Hz, 12H).

$^{13}$C NMR (100 MHz, $CDCl_3$) δ 167.6, 157.9, 155.8, 135.6, 132.3, 130.4, 130.3, 130.1, 127.8, 114.3, 113.9, 96.6, 74.0, 51.7, 51.1, 47.5, 46.7, 46.3, 41.7, 12.81.

HRMS (ESI): $C_{35}H_{41}N_4O_2^+$; Calculated mass, $[M]^+$ 549.3224; Observed mass, $[M+H]^+$ 549.3249.

**5-Azidofluorescein** (**20**)

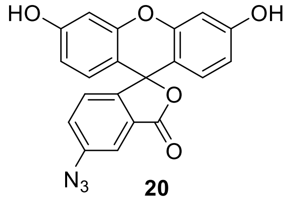


**20**

5-Azidofluorescein (**20**) was synthesized according to a previously reported method.[4]

### 3. General procedure for cell culture

Mammalian cells (HEK293/HeLa/MDA-MB-231) were cultured in DMEM medium supplemented with 10% FBS, 100 units/mL penicillin, and 100 µg/mL streptomycin (complete media) in a humidified $CO_2$

incubator maintaining 5% $CO_2$ levels and a temperature of 37 °C. All the treatments and incubations of mammalian cells performed in a $CO_2$ incubator were done in an incubator maintaining 5% $CO_2$ levels, and a temperature of 37 °C.

### 4. Procedure for the characterization of metabolic labeling by cellular imaging

A. <u>Imaging metabolically labeled choline lipids</u>: 0.2 million HEK293 cells were seeded in 1 mL of complete media in a 24-well plate. After a 12 h incubation in a $CO_2$ incubator, 10 µL of 200 mM stock solutions of choline analogs in PBS were administered into each well (final concentration: 2 mM). The same volume of PBS devoid of choline analogs was added in the control wells. After a further 24 h incubation in a $CO_2$ incubator, cells were washed with PBS (2 × 500 µL) at room temperature and fixed by incubating them in formaldehyde (250 µL of a 3.7% *w/v* solution in PBS) for 2 h at 4 °C. The fixed cells were washed with TBS (2 × 500 µL) and then incubated in a click reaction cocktail (250 µL) for 30 min at room temperature under dark. The click reaction cocktail was made in 0.1 M Tris-HCl buffer at pH 8.5 and contained 1 mM $CuSO_4.5H_2O$ (added from a stock of 500 mM in water), 100 mM L-ascorbic acid (added from a stock of 500 mM in water) and 20 µM 5-azidofluorescein (added from a stock of 10 mM in DMSO) for imaging alkynyl choline analogs or 20 µM rhodamine alkyne (added from a stock of 1.25 mM in DMSO) for imaging cells treated with azido choline analogs. After completion of this 30 min-long incubation, the cells were washed with TBS (1 × 500 µL), 0.5 M NaCl (1 × 500 µL) and then again with TBS (1 × 500 µL). Finally, 250 µL TBS was added to each well and the cells were imaged using the EVOS-M7000 cell imaging system. All images were acquired by using a 40× objective either under the GFP channel (for analogs **1**, **3**-**9**, **11**, and **18**) or the RFP channel (for analog **2**) and the representative images are provided in Figure 2 (for analogs **1**-**9**), Figure 3c (for analog **11**) and Figure S25a (for analog **18**). Each experiment was performed three times by administering the cells with the aforementioned choline analogs into three separate wells and performing the imaging experiment as described above.

B. <u>Choline competition</u>: 0.2 million HEK293 cells were seeded in 1 mL of complete media in a 24-well plate. After 12 h of incubation in a $CO_2$ incubator, choline chloride (10 µL of a 200 mM solution in PBS) and the choline analog (10 µL of a 200 mM solution in PBS) were added. After a further 24 h incubation in a $CO_2$ incubator, cells were treated in the same way as mentioned in sub-section 4A for cellular imaging. All images were acquired by illuminating the cells with the same exposure that was used to acquire images for the corresponding choline analog treatments without any co-administration with choline. Each experiment was performed in triplicate.

### 5. Procedure for the characterization of metabolic labeling by lipidomics

A. Extraction of lipids from HEK293/HeLa/MDA-MB-231 cells: For all lipidomics experiments, 3 million cells in 10 mL of complete medium were seeded into two 10 cm sterile culture dishes (for each treatment) and incubated for 24 h in a $CO_2$ incubator. Subsequently, 317 µL of 65 mM solutions of choline analogs in PBS (final concentration of each analog in each flask was 2 mM) were administered and after a 24 h-long incubation in a $CO_2$ incubator, the adhered cells were washed with ice-cold PBS (2 × 5 mL), resuspended in ice-cold PBS (5 mL), pooled in a centrifuge tube and centrifuged at 1,000 *g* for 5 min at room temperature. The supernatant was then aspirated off and the lipids were extracted from the resultant pellet using modified Bligh Dyer method.[5] Briefly, the cell pellet was treated with a mixture of chloroform and methanol (1 mL of a 1:2 *v*/*v* solution), vortexed for 1 min, followed by addition of chloroform (0.5 mL) and 1 M NaCl (0.5 mL), and again vortexed for 1 min. Phase separation was achieved by centrifuging the mixture at 3,220 *g* at room temperature for 30 min. The lower lipid-enriched organic layer was separated and dried using a benchtop vacuum concentrator (Centrivap Labconco) at room temperature. For compound parameter optimization via direct infusion method, the dried lipid samples were redissolved in chloroform (1 mL) and 40 µL of this lipid stock solution was added to a mixture of chloroform/methanol/300 mM ammonium acetate in water such that the final composition was 300/665/35 (*v*/*v*/*v*) and the final volume of the lipid mixture was 1.2 mL. For all LC-MS/MS analysis, the dried lipids were dissolved in a mixture of chloroform and methanol (200 µL of a 1:1 *v*/*v* solution). For all control experiments, the procedure described above was followed without administration of any choline derivatives.

B. HPLC method details: Lipids extracted from cells were fractionated on a 2.6 µm Kinetex HILIC column (I.D. 100 × 2.1 mm, 100 Å, Phenomenex). Total Ion Chromatograms (TIC) were generated using the Precursor Ion Scan (PIS) modes for various non-natural choline head groups (see Figure S1 for an explanation of the PIS mode) on the HILIC column using a binary gradient. The elution protocol used is summarized below:

Mobile Phase A: 95:5 Acetonitrile: Water (both acetonitrile and water contained 10 mM ammonium acetate)

Mobile Phase B: 1:1 Acetonitrile: Water (both acetonitrile and water contained 10 mM ammonium acetate)

Gradient:

0 to 1 min: 100% A

1 to 5 min: 100% A to 90% A

5 to 13 min: 90% A to 70% A

13 to 25 min: 70% A to 100% A

25 min to 39 min: 100% A (for column re-equilibration)

Flow rate: 0.5 mL/min

Column oven temperature: 40 °C

Sample volume: 10 µL of lipid solution extracted from cells.

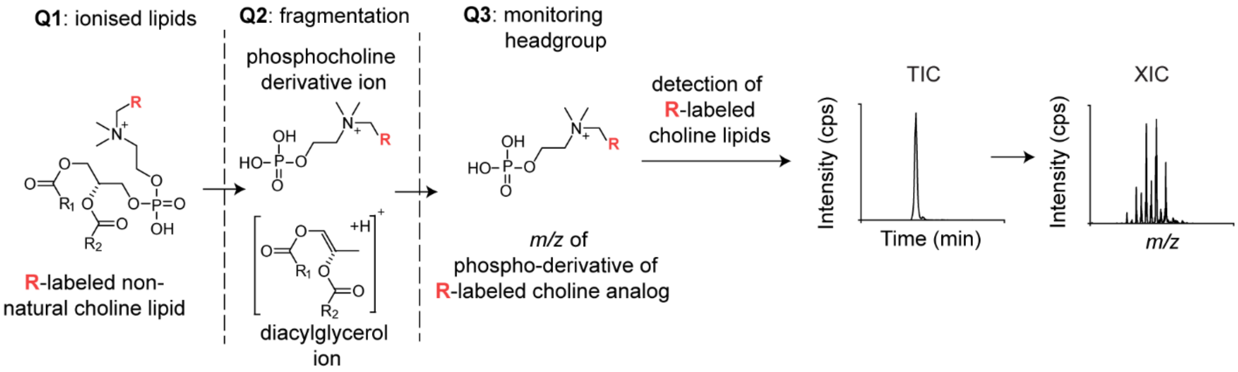


**Figure S1.** A diagrammatic representation for the use of the PIS mode to detect non-natural choline lipids. A representative ionized non-natural choline lipid with **R**-labeled phosphocholine as its headgroup and $R_1$ and $R_2$ as its hydrophobic tails, travels from the first quadrupole (Q1) of the QTRAP4500 instrument to the second quadrupole (Q2) where it is fragmented by collision activated dissociation (CAD) into the diacylglycerol ion and the ion of the phospho-derivative of the **R**-labeled choline analog. The third quadrupole (Q3) scans for the *m/z* of the fragment ion corresponding to the phospho-derivative of **R**-labeled choline analog. All parent ions that yield this fragment ion in the Q3 are detected, yielding a peak on the total ion chromatogram (TIC) which is further processed by the Analyst software to generate the extracted ion chromatogram (XIC) that shows the *m/z* values for all the **R**-labeled choline lipids. To account for the loss of $N_2$ incurred during the fragmentation of diazirine-containing lipids while performing MS/MS analysis, the PIS scans performed for all lipids labeled with diazirine derivatives reported in this study employed the daughter ion *m/z* value corresponding to 28 Da less than that for the phospho-derivative of the compound.

C. Mass-spectrometry details: Lipids eluted from the HILIC column were analyzed and detected on an AB Sciex 4500 QTRAP mass spectrometer equipped with an ESI source heated to 400 °C. The scan speed was maintained at 200 Da/sec. The electrospray capillary was held at +5500 volts, curtain gas was set at 35 (arbitrary units), CAD gas was maintained at the medium level and the two ion source gases GS1 and GS2 were set at 50 and 60 (arbitrary units) respectively. The PIS mode (schematically described in Figure S1) was used for detecting all the non-natural or native choline lipids to generate the TICs and all TICs depicted in the main text (Figures 2-4) and in the sub-sections below (Figures S17-S23, S26-28) are representative of at least three biological replicates. The TIC peaks were processed on the Analyst software to generate the Extracted Ion Chromatograms (XICs). The compound parameters (declustering potential, entrance potential, collision energy, and collision cell exit potential) were optimized by ramping each of them against the intensity by the direct infusion method. Native choline lipids were detected by employing a PIS of 184.1 Da (*m/z* for phosphocholine) and the compound parameters were optimized separately. For characterizing metabolic labeling with alkynyl/azido-substituted choline analogs **1-9** and **14**, and with the D6 choline analog **16**, the PIS masses of their corresponding phosphoryl derivatives, and the optimized compound parameters for the PIS of 184.1 Da (*m/z* for phosphocholine) were used. For all diazirine analogs, the PIS mass used was 28 Da less than that of the fragment mass of their phosphoryl derivatives to account for the $N_2$ loss that diazirines characteristically undergo during fragmentation. Compound parameters were separately optimized for detecting the β-diazirine **11**-labeled choline lipids by using a PIS of 234.1 Da and the same parameters were used in all PIS experiments for detecting choline lipids labeled with other β-diazirine analogs **10**, **12**, **13**, **S26**, **S35**, and **S36**. Compound parameters were separately optimized for detecting γ-diazirine **18-**labeled choline lipids by using a PIS of 248.1 Da and the same parameters were used to detect **17**-labeled choline lipids. Each TIC was background-subtracted with the corresponding blank chromatogram. The details of compound parameters for the detection of metabolic incorporation of each analog using PIS mode are listed in Table S1 below.

**Table S1**. Compound parameters used for the PIS of lipids labeled with all the choline analogs studied

<table>
<tr><th>S. No.</th><th>Head group</th><th>PIS mode mass (Da)</th><th>Declustering Potential (volts)</th><th>Entrance Potential (volts)</th><th>Collision Energy (volts)</th><th>Collision cell Exit Potential (volts)</th></tr>
<tr><td colspan="7">For native, D6 and diazirine-devoid alkynl/azido-substituted choline lipids</td></tr>
<tr><td>1.</td><td>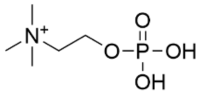<br>phosphocholine (native lipids)</td><td>184.1</td><td rowspan="3">110.0</td><td rowspan="3">7.8</td><td rowspan="3">30.9</td><td rowspan="3">5.2</td></tr>
<tr><td>2.</td><td>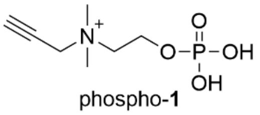<br>phospho-**1**</td><td>208.1</td></tr>
<tr><td>3.</td><td>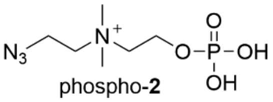<br>phospho-**2**</td><td>239.1</td></tr>
</table>

| | | | | | | |
|---|---|---|---|---|---|---|
| 4. | 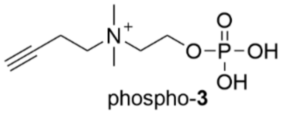 phospho-**3** | 222.1 | | | | |
| 5. | 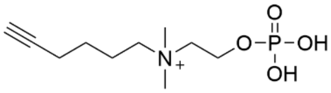 phospho-**4** | 250.1 | | | | |
| 6. | 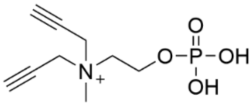 phospho-**5** | 232.1 | | | | |
| 7. | 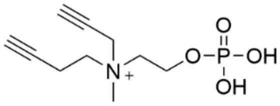 phospho-**6** | 246.1 | | | | |
| 8. | 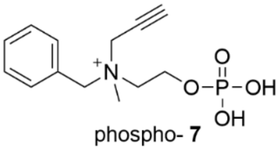 phospho- **7** | 284.1 | 110.0 | 7.8 | 30.9 | 5.2 |
| 9. | 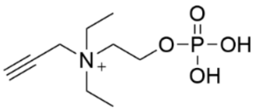 phospho-**8** | 236.1 | | | | |
| 10. | 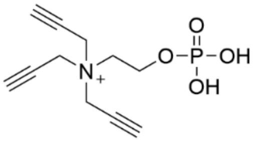 phospho-**9** | 256.1 | | | | |
| 11. | 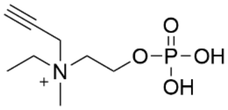 phospho-**14** | 222.1 | | | | |
| 12. | 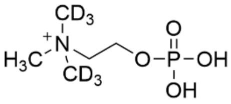 phospho-**16** | 190.1 | | | | |

| | | | | | | |
|---|---|---|---|---|---|---|
| For β-diazirine choline lipids | | | | | | |
| 13. | 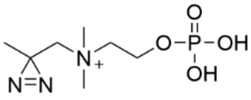 phospho-**10** | 210.1 (loss of $N_2$) | 169.3 | 5.0 | 53.6 | 1.1 |
| 14. | 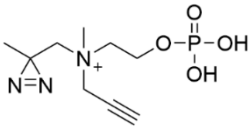 phospho-**11** | 234.1 (loss of $N_2$) | | | | |
| 15. | 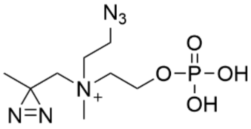 phospho-**12** | 265.1 (loss of $N_2$) | | | | |
| 16. | 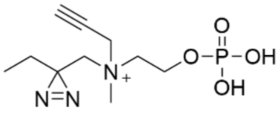 phospho-**S26** | 248.1 (loss of $N_2$) | | | | |
| 17. | 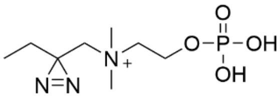 phospho-**S35** | 224.1 (loss of $N_2$) | | | | |
| 18. | 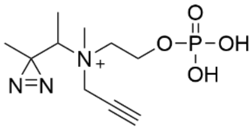 phospho-**13** | 248.1 (loss of $N_2$) | | | | |
| 19. | 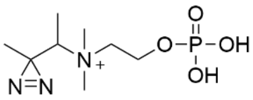 phospho-**S36** | 224.1 (loss of $N_2$) | | | | |
| For γ-diazirine choline lipids | | | | | | |
| 20. | 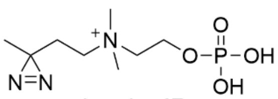 phospho-**17** | 224.1 (loss of $N_2$) | 100.0 | 2.9 | 43.0 | 15.0 |

| | | | | | | |
|---|---|---|---|---|---|---|
| 21. | 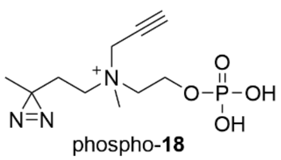 phospho-**18** | 248.1 (loss of $N_2$) | 100.0 | 2.9 | 43.0 | 15.0 |

D. Identification of non-natural lipids using the LipidView software: To identify the choline lipids labeled with non-natural headgroups **1-18**, the corresponding XIC was processed using the LipidView (AB Sciex, version 1.2) software[6] by manually modifying its database. Peaks were corrected for isotopic overlap and were processed by setting a mass tolerance of 0.5 Da, with a minimum intensity of 0.1% and a minimum signal-to-noise ratio of 10.[7, 8] The subsequent phospholipid species were then verified by Lipid Maps structure database (LMSD)[9] and the ones absent from this platform were excluded from the analysis. Each PC lipid is annotated by using two numbers; for example, 34:2 where 34 is the sum of carbon atoms in the two fatty acyl chains and 2 is the total number of double bonds present in the two fatty acyl chains of the corresponding PC lipid. Each SM lipid species is annotated by using three numbers; for example, 40:1;2 where 40 is the sum of the carbon atoms in sphingosine back bone and the fatty acyl chain, 1 is the total number of double bond(s) present in the sphingosine backbone and the fatty acyl chain, and 2 is the sum of the hydroxyl groups present on the sphingosine backbone including the phosphorylated hydroxyl group at C-1 position of the sphingosine backbone.

### 6. LipidView analysis of XICs obtained for choline lipids extracted from HEK293 cells cultured in the presence of compounds 1-11

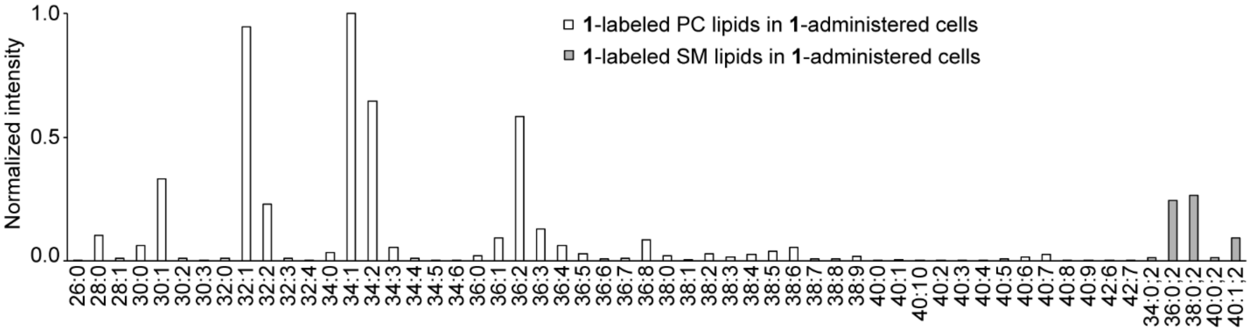


**Figure S2.** A bar graph obtained from the LipidView software denoting all **1**-labeled PC and SM lipids identified from the XIC depicted in Figure 2a of the main text. The *y*-axis has been normalized to the most intense peak *i.e.*, lipid 34:1 of the XIC.

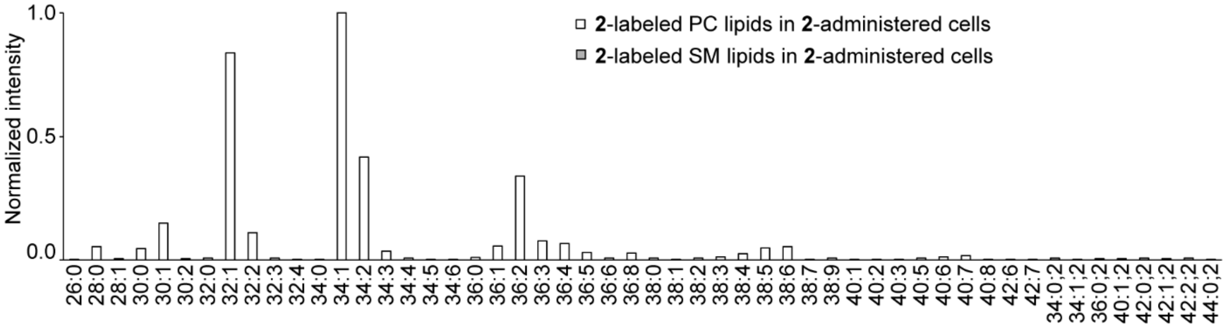


**Figure S3.** A bar graph obtained from the LipidView software denoting all **2**-labeled PC and SM lipids identified from the XIC depicted in Figure 2b of the main text. The *y*-axis has been normalized to the most intense peak *i.e.*, lipid 34:1 of the XIC.

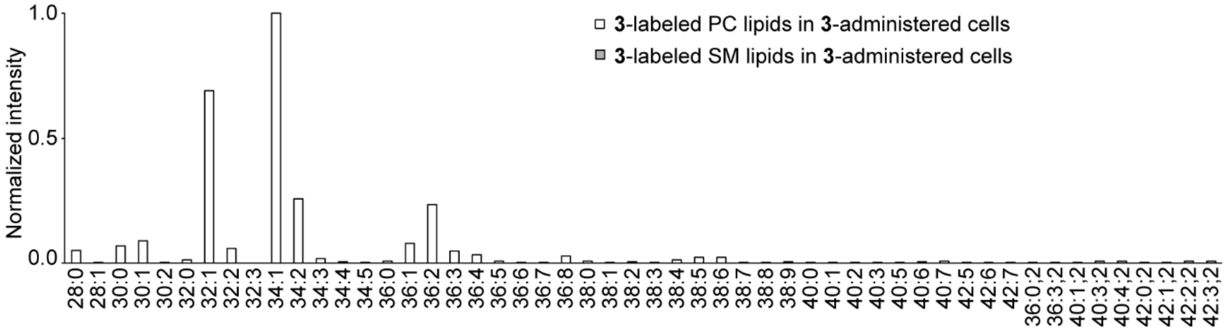


**Figure S4.** A bar graph obtained from the LipidView software denoting all **3**-labeled PC and SM lipids identified from the XIC depicted in Figure 2c of the main text. The *y*-axis has been normalized to the most intense peak *i.e.*, lipid 34:1 of the XIC.

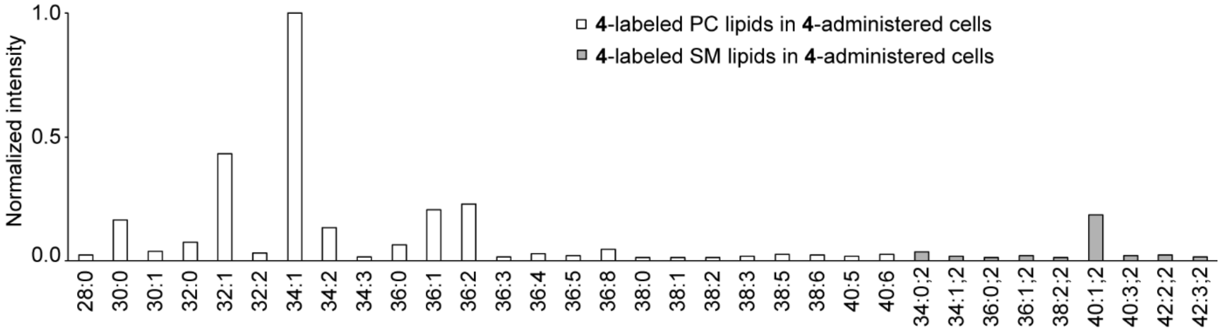


**Figure S5.** A bar graph obtained from the LipidView software denoting all **4**-labeled PC and SM lipids identified from the XIC depicted in Figure 2d of the main text. The *y*-axis has been normalized to the most intense peak *i.e.*, lipid 34:1 of the XIC.

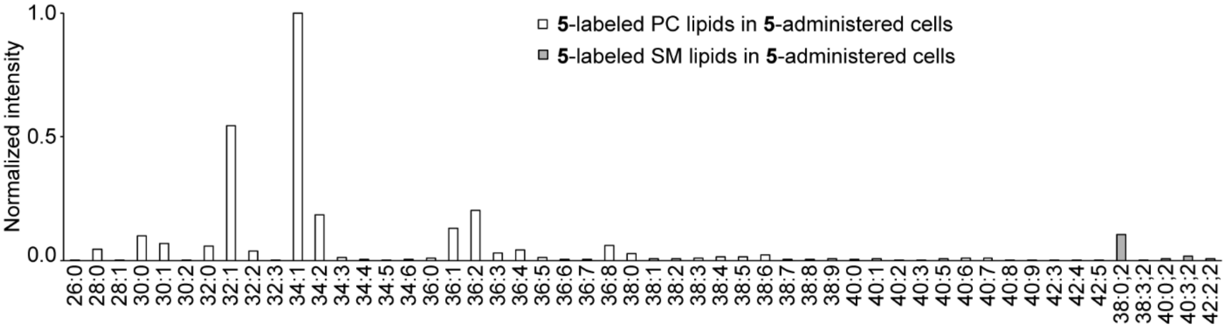


**Figure S6.** A bar graph obtained from the LipidView software denoting all **5**-labeled PC and SM lipids identified from the XIC depicted in Figure 2e of the main text. The *y*-axis has been normalized to the most intense peak *i.e.*, lipid 34:1 of the XIC.

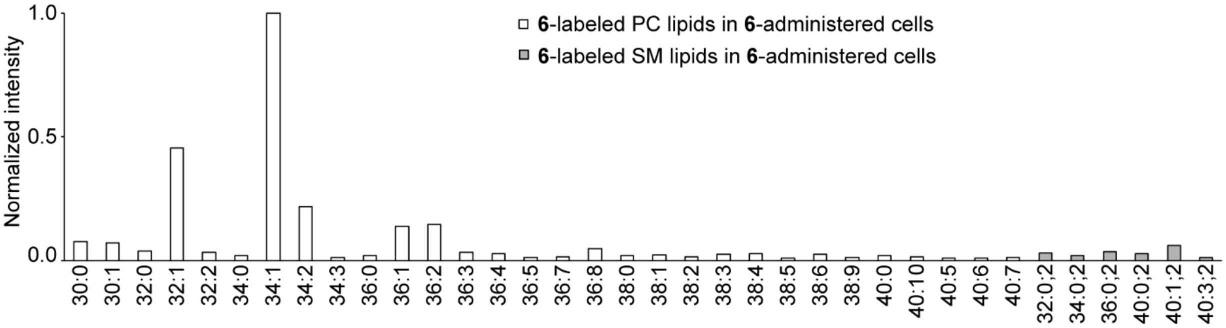


**Figure S7.** A bar graph obtained from the LipidView software denoting all **6**-labeled PC and SM lipids identified from the XIC depicted in Figure 2f of the main text. The *y*-axis has been normalized to the most intense peak *i.e.*, lipid 34:1 of the XIC.

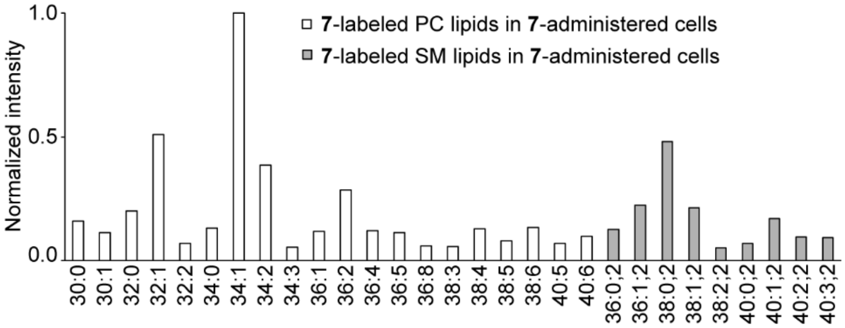


**Figure S8.** A bar graph obtained from the LipidView software denoting all **7**-labeled PC and SM lipids identified from the XIC depicted in Figure 2g of the main text. The *y*-axis has been normalized to the most intense peak *i.e.*, lipid 34:1 of the XIC.

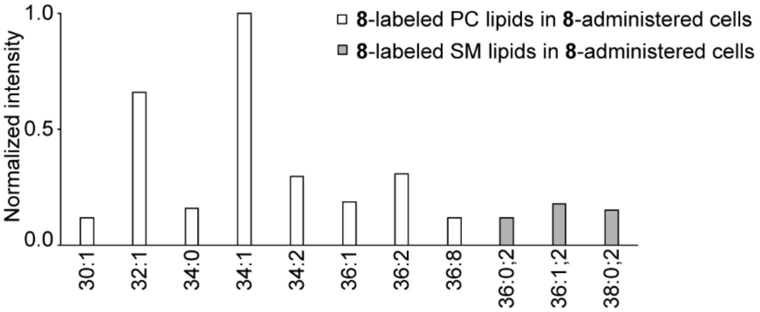


**Figure S9.** A bar graph obtained from the LipidView software denoting all **8**-labeled PC and SM lipids identified from the XIC depicted in Figure 2h of the main text. The *y*-axis has been normalized to the most intense peak *i.e.*, lipid 34:1 of the XIC.

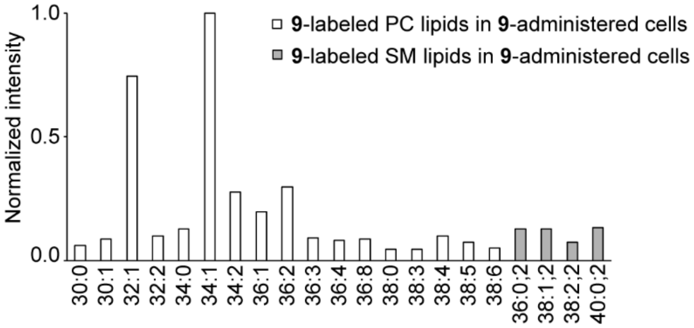


**Figure S10.** A bar graph obtained from the LipidView software denoting all **9**-labeled PC and SM lipids identified from the XIC depicted in Figure 2i of the main text. The *y*-axis has been normalized to the most intense peak *i.e.*, lipid 34:1 of the XIC.

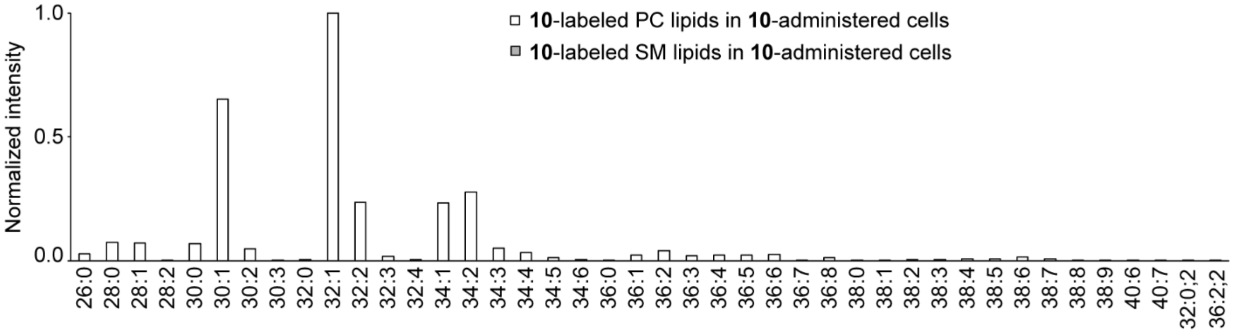


**Figure S11.** A bar graph obtained from the LipidView software denoting all **10**-labeled PC and SM lipids identified from the XIC depicted in Figure 3a of the main text. The *y*-axis has been normalized to the most intense peak *i.e.*, lipid 32:1 of the XIC.

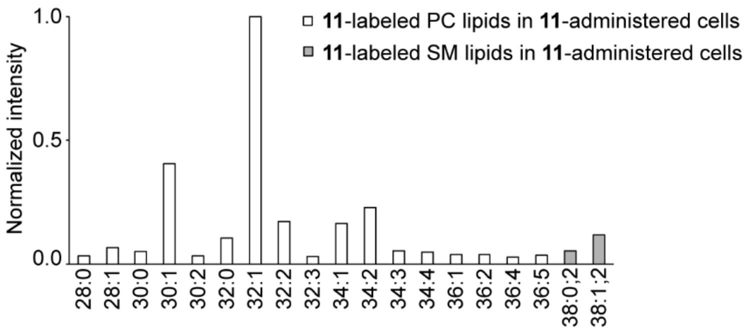


**Figure S12.** A bar graph obtained from the LipidView software denoting all **11**-labeled PC and SM lipids identified from the XIC in Figure 3b of the main text. The *y*-axis has been normalized to the most intense peak *i.e.*, lipid 32:1 of the XIC.

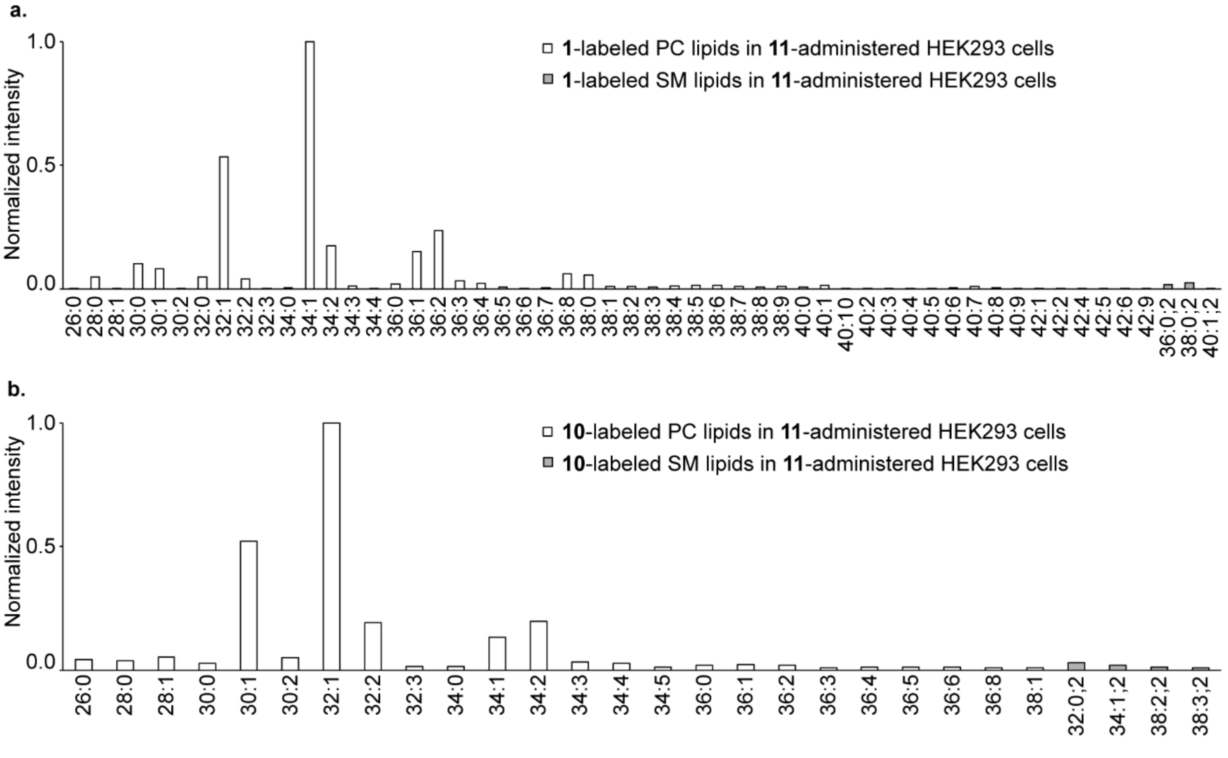


**Figure S13**. a) LipidView analysis of **1**-labeled PC and SM lipids obtained from the HEK293 cells cultured for 24 h in the presence of **11** (2 mM). These lipids were identified from the XIC depicted in Figure 3c (middle panel) of the main text. The *y*-axis has been normalized to the most intense peak *i.e.*, lipid 34:1 of the XIC. b) LipidView analysis of **10**-labeled PC and SM lipids obtained from the HEK293 cells cultured for 24 h in the presence of **11** (2 mM). These lipids were identified from the XIC depicted in Figure 3d (top panel) of the main text. The *y*-axis has been normalized to the most intense peak *i.e.*, lipid 32:1 of the XIC.

7. **Photoactivation and stability studies on 11**

A. UV-mediated photoactivation of **11**: Aliquots of **11** (3.2 mM, 600 µL) in $D_2O$ were irradiated for 5 min, 15 min, and 30 min on ice with 356 nm UV as described in the general information sub-section and then subjected to $^1H$ NMR (400 MHz) analysis. A sample that was not exposed to any UV irradiation served as control. The NMR traces depicted in Figure S14 show UV exposure time-dependent decrease in the peaks corresponding to **11**.

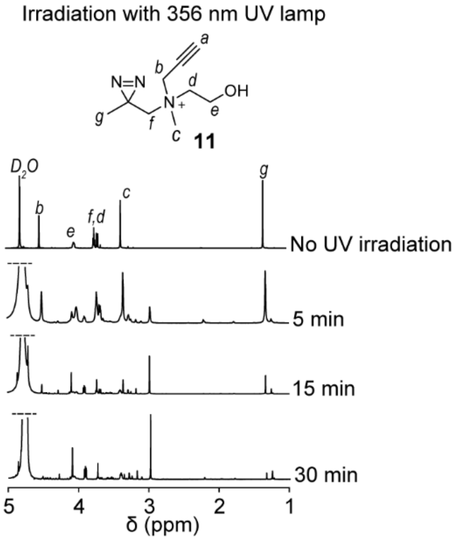


**Figure S14**. $^1H$-NMR characterization of the time-course of UV-mediated uncaging of **11**. $^1H$ NMR traces of solutions of **11** (3.2 mM) in $D_2O$ that were either not irradiated or irradiated for 5 min, 15 min, and 30 min with UV light (356 nm) on ice. The NMR traces show almost complete photolysis of **11** within 30 min as demonstrated by a substantial attenuation of the peak for the protons on the carbon atoms labeled "*g*" in the structure of **11**, as well as those for other peaks.

B. Thermal stability of **11**: Three samples of **11** incubated at -20 °C, 25 °C and 37 °C for 12 h under dark were dissolved in water to yield 4.1 mM solutions of **11**, and subjected to UV-Visible spectroscopy at room temperature. The characteristic peak for the diazirine group at 329 nm was obtained in each of the three samples as shown in the UV absorption spectra in Figure S15a and the peaks were of similar intensities, establishing the thermal stability of **11**.

C. Stability of **11** in complete media: To a 600 µL solution of **11** (22 mM) in $D_2O$ was added 100 µL of complete media containing DMEM with 10% FBS (final conc. of **11** was 18.9 mM) and the resultant solution was incubated at 37 °C for 12 h under dark. In the control sample, 100 µL of complete media was replaced by 100 µL of $D_2O$. The resultant solution was subjected to $^1H$ NMR (400 MHz) spectroscopy that yielded identical NMR traces for both the samples as depicted in Figure S15b demonstrating the stability of **11** in complete media.

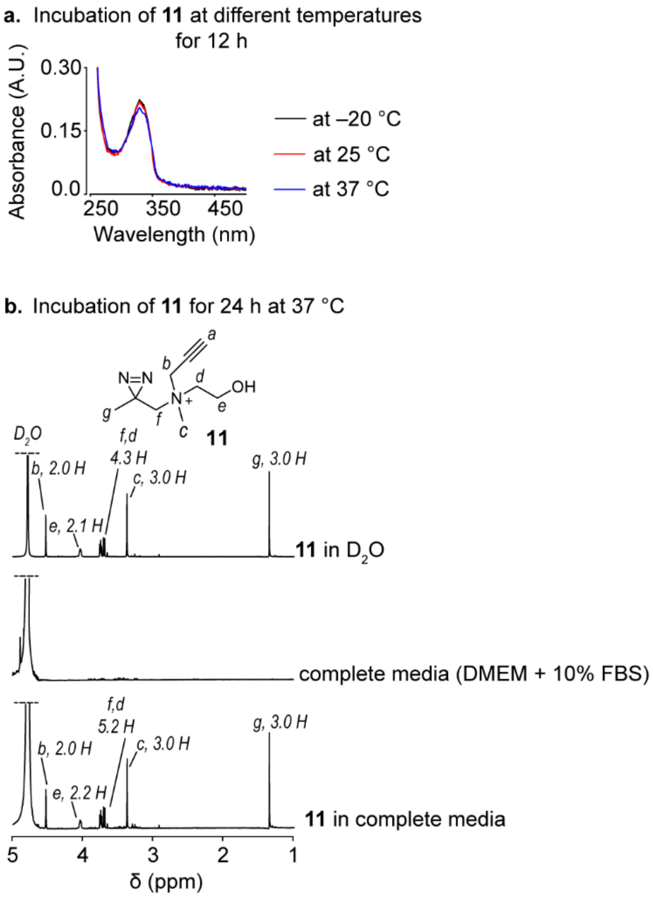


**Figure S15**. Stability of **11**. a) The thermal stability of **11** was monitored by incubating it at the depicted temperatures for 12 h under dark followed by recording the UV absorption spectra of each sample dissolved in water to a concentration of 4.1 mM. b) The $^1$H NMR spectra (400 MHz) of a solution of **11** in $D_2O$ (top) and in DMEM + $D_2O$ (bottom) after incubation at 37 °C for 12 h under dark with tetramethylsilane (TMS) as the reference. The peaks in the spectra are assigned to the protons in the structure of **11** by annotating them by using the symbols *a-g* and the integral values in terms of the number of protons for each peak is denoted next to it. The $^1$H NMR spectrum (400 MHz) of a solution of complete media in $D_2O$ (100 µL of complete media added to 600 µL $D_2O$) is provided in the middle panel.

### 8. Metabolic labeling of HEK293, HeLa, and MDA-MB-231 cells with 11

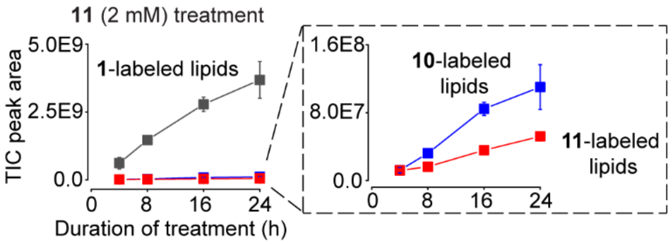


**Figure S16**. A plot of the TIC peak areas versus time wherein the areas under the TIC plots were obtained from the PIS for **1**, **10**, and **11**-labeled lipids on the lipids extracted from HEK293 cells cultured in presence of **11** (2 mM) for 4, 8, 16, and 24 h. The representative TIC and XIC plots for the aforementioned treatments are depicted in Figure S17. Each data point is an average of three biological replicates and the error bars represent the standard error values.

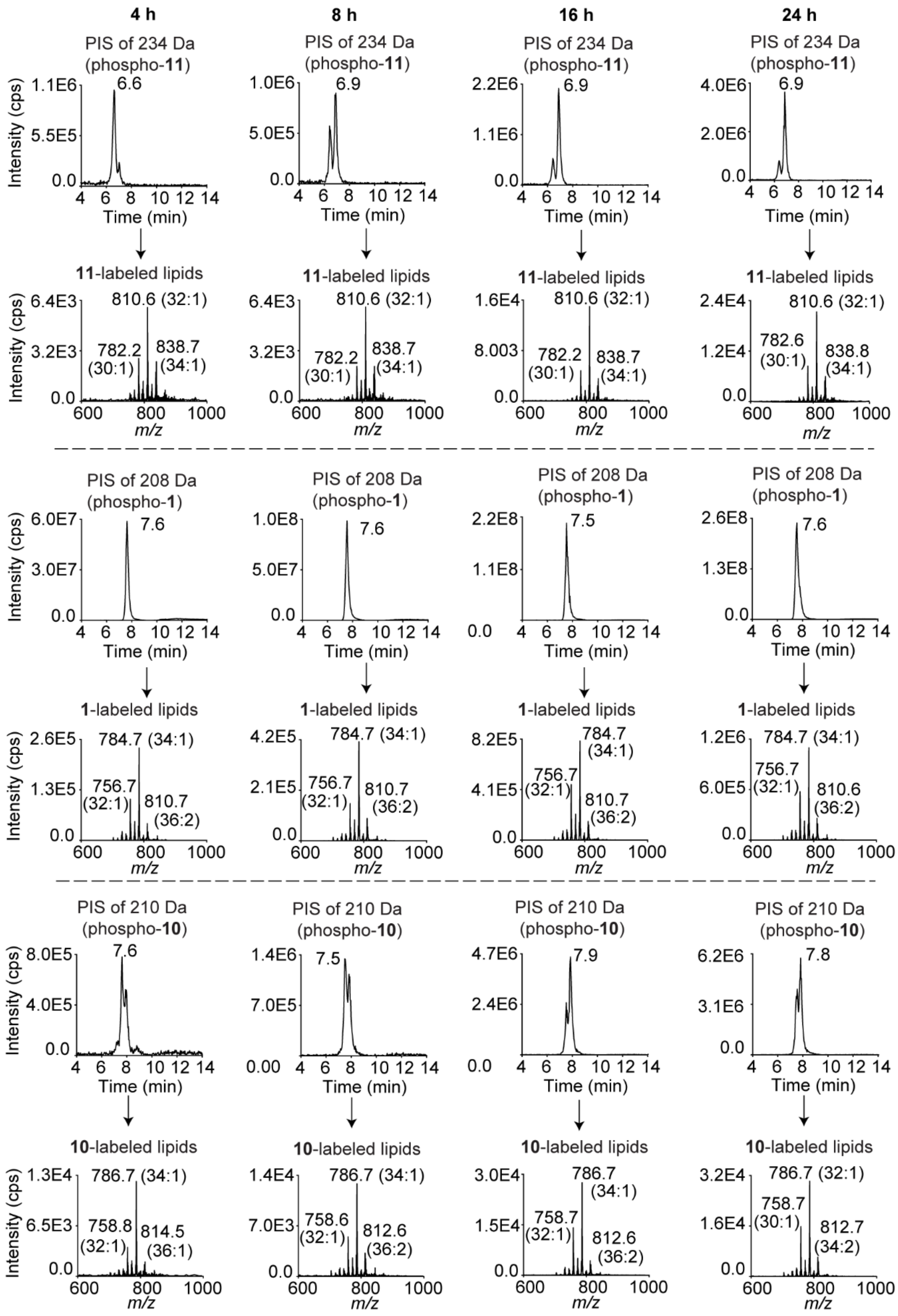


**Figure S17**. *Legend on the next page.*

*Figure S17 legend continued from the last page.* Kinetics for the formation of the metabolically labeled choline lipids produced in HEK293 cells cultured in presence of **11**. Lipids extracted from HEK293 cells that were administered with **11** (2 mM) for 4 h, 8 h, 16 h, and 24 h were subjected to LC-MS/MS and analyzed for the formation of **11**, **1**, and **10**-labeled lipids by employing a PIS of 234 Da, 208 Da, and 210 Da respectively. Each chromatogram is representative of at least three biological replicates. The area under the TIC peaks were manually calculated using the Analyst software to generate the plots in Figure S16 and S18 (left).

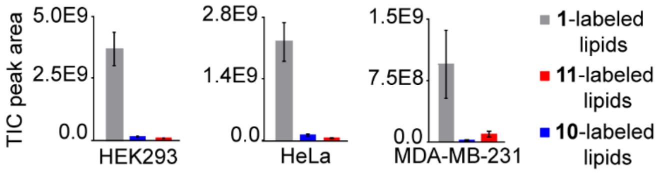


**Figure S18**. A bar graph representing the TIC peak areas for **1**, **10**, and **11**-labeled lipids extracted from HEK293, HeLa and MDA-MB-231 mammalian cells cultured for 24 h in the presence of **11** (2 mM). Each peak area value is an average of three biological replicates and the error bars represent the standard error values. The representative TIC and XIC plots for these experiments are depicted in Figures S17, S19 and S20.

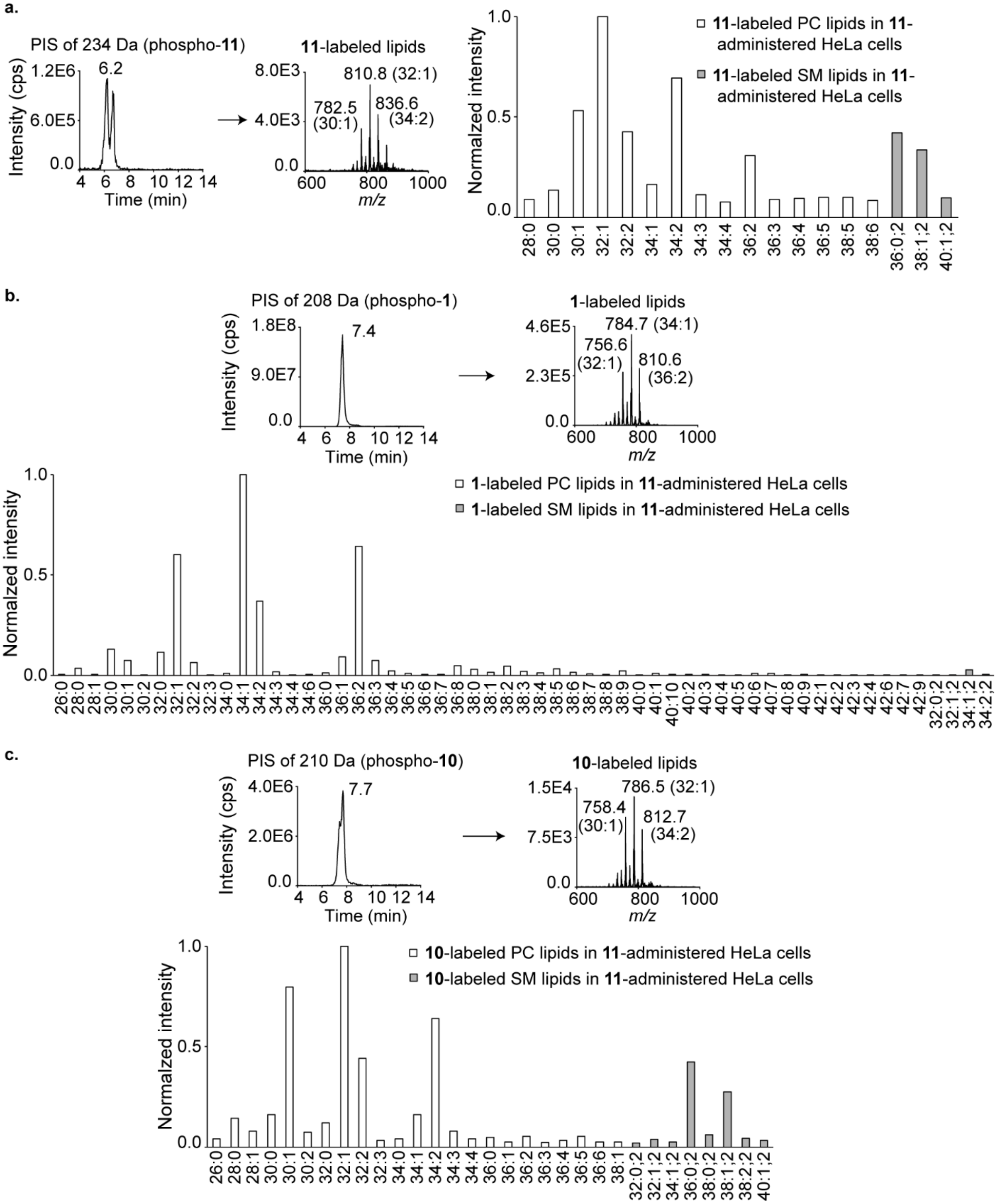


**Figure S19.** Characterization of the metabolically labeled choline lipids produced in HeLa cells cultured in presence of **11**. a) Left: Lipids extracted from HeLa cells that were cultured in presence of **11** (2 mM) for 24 h were subjected to LC-MS/MS and analyzed for the formation of **11**-labeled lipids by employing a PIS of 234 Da. The area under the peak on the TIC was manually calculated using the Analyst software to generate the plot on Figure S18 (middle). Right: A bar graph obtained from the LipidView software denoting all the **11**-labeled PC and SM lipids identified from the XIC on the left panel. The *y*-axis has been normalized to the most intense peak *i.e.*, lipid 32:1 of the XIC. *Legend continued to the next page.*

*Figure S19 legend continued from the previous page*. b) Top: Lipids extracted from HeLa cells that were cultured in presence of **11** (2 mM) for 24 h were subjected to LC-MS/MS and analyzed for the formation of **1**-labeled lipids by employing a PIS of 208 Da. The area under the peak on the TIC was manually calculated using the Analyst software to generate the plot on Figure S18 (middle). Bottom: A bar graph obtained from the LipidView software denoting all **1**-labeled PC and SM lipids identified from the XIC on the top panel. The *y*-axis has been normalized to the most intense peak *i.e.*, lipid 34:1 of the XIC. c) Top: Lipids extracted from HeLa cells that were cultured in presence of **11** (2 mM) for 24 h were subjected to LC-MS/MS and analyzed for the formation of **10**-labeled lipids by employing a PIS of 210 Da. The area under the TIC peaks were manually calculated using the Analyst software to generate the plot on Figure S18 (middle). Bottom: A bar graph obtained from the LipidView software denoting all **10**-labeled PC and SM lipids identified from the XIC on the top panel. The *y*-axis has been normalized to the most intense peak *i.e.*, lipid 32:1 of the XIC.

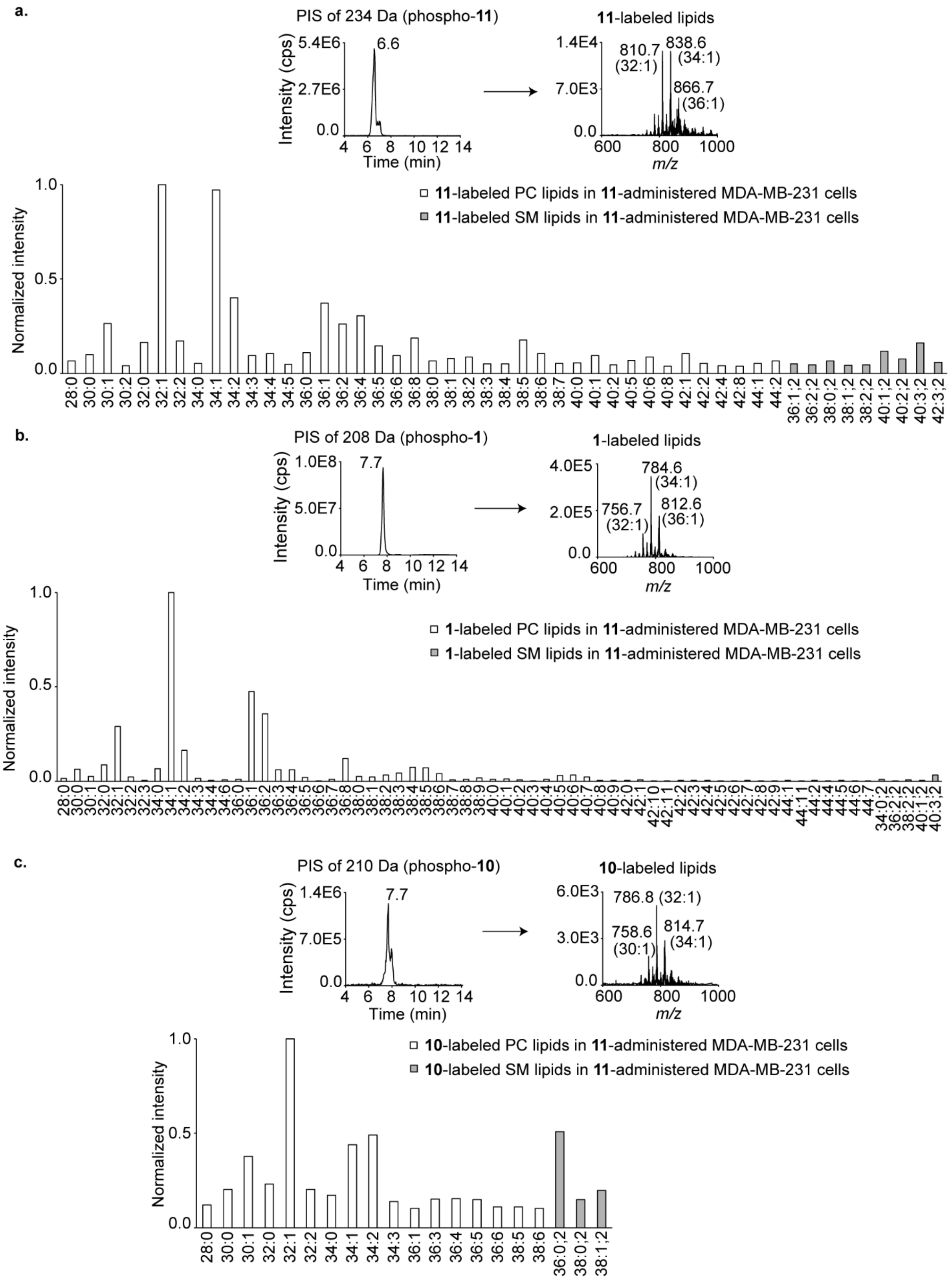


**Figure S20**. *Legend on the next page.*

*Figure S20 legend continued from the previous page.* Characterization of the metabolically labeled choline lipids produced in MDA-MB-231 cells cultured in presence of **11**. a) Top: Lipids extracted from MDA-MB-231 cells cultured in presence of **11** (2 mM) for 24 h were subjected to LC-MS/MS and analyzed for the formation of **11**-labeled lipids by employing a PIS of 234 Da. The area under the peak on the TIC was manually calculated using the Analyst software to generate the plot on Figure S18 (right). Bottom: A bar graph obtained from the LipidView software denoting all **11**-labeled PC and SM lipids identified from the XIC in the top panel. The *y*-axis has been normalized to the most intense peak *i.e.*, lipid 32:1 of the XIC. b) Top: Lipids extracted from MDA-MB-231 cells cultured in presence of **11** (2 mM) for 24 h were subjected to LC-MS/MS and analyzed for the formation of **1**-labeled lipids by employing a PIS of 208 Da. The area under the peak on the TIC was manually calculated using the Analyst software to generate the plot on Figure S18 (right). Bottom: A bar graph obtained from the LipidView software denoting all **1**-labeled PC and SM lipids identified from the XIC in the top panel. The *y*-axis has been normalized to the most intense peak *i.e.*, lipid 34:1 of the XIC. c) Top: Lipids extracted from MDA-MB-231 cells cultured in presence of with **11** (2 mM) for 24 h were subjected to LC-MS/MS and analyzed for the formation of **10**-labeled lipids by employing a PIS of 210 Da. The area under TIC peaks were manually calculated using the Analyst software to generate the plot on Figure S18 (right). Bottom: A bar graph obtained from the LipidView software denoting all **10**-labeled PC and SM lipids identified from the XIC in the top panel. The *y*-axis has been normalized to the most intense peak *i.e.*, lipid 32:1 of the XIC.

### 9. Metabolic labeling studies on the β-diazirine compounds 12, S26 and 13

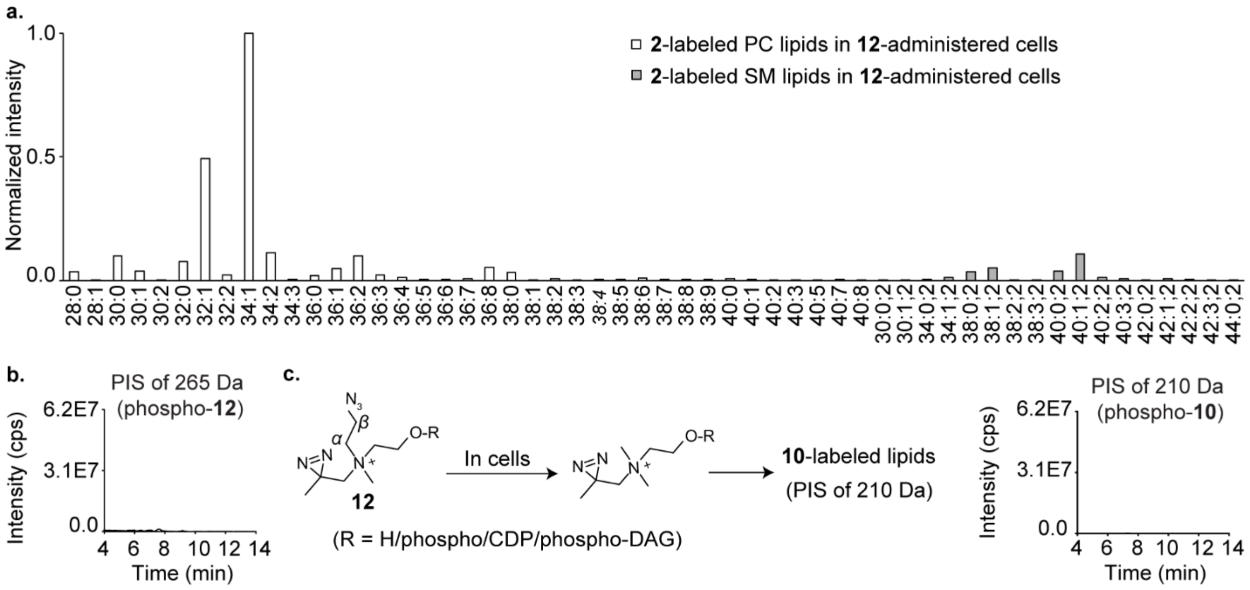


**Figure S21**. a) LipidView analysis of **2**-labeled PC and SM lipids obtained from the HEK293 cultured for 24 h in the presence of **12** (2 mM). These lipids were identified from the XIC depicted in Figure 3e (bottom panel) of the main text. The *y*-axis has been normalized to the most intense peak *i.e.*, lipid 34:1 of the XIC. b) Lipids extracted from HEK293 cells cultured for 24 h in the presence of **12** (2 mM) were subjected to LC-MS/MS and analyzed for the formation of **12**-labeled lipids by employing a PIS of 265 Da. The resultant TIC shows no peak for **12**-labeled lipids. c) Lipids extracted from HEK293 cells cultured for 24 h in the presence of **12** (2 mM) were subjected to LC-MS/MS and analyzed for the formation of **10**-labeled lipids by employing a PIS of 210 Da. The resultant TIC shows no peak for **10**-labeled lipids.

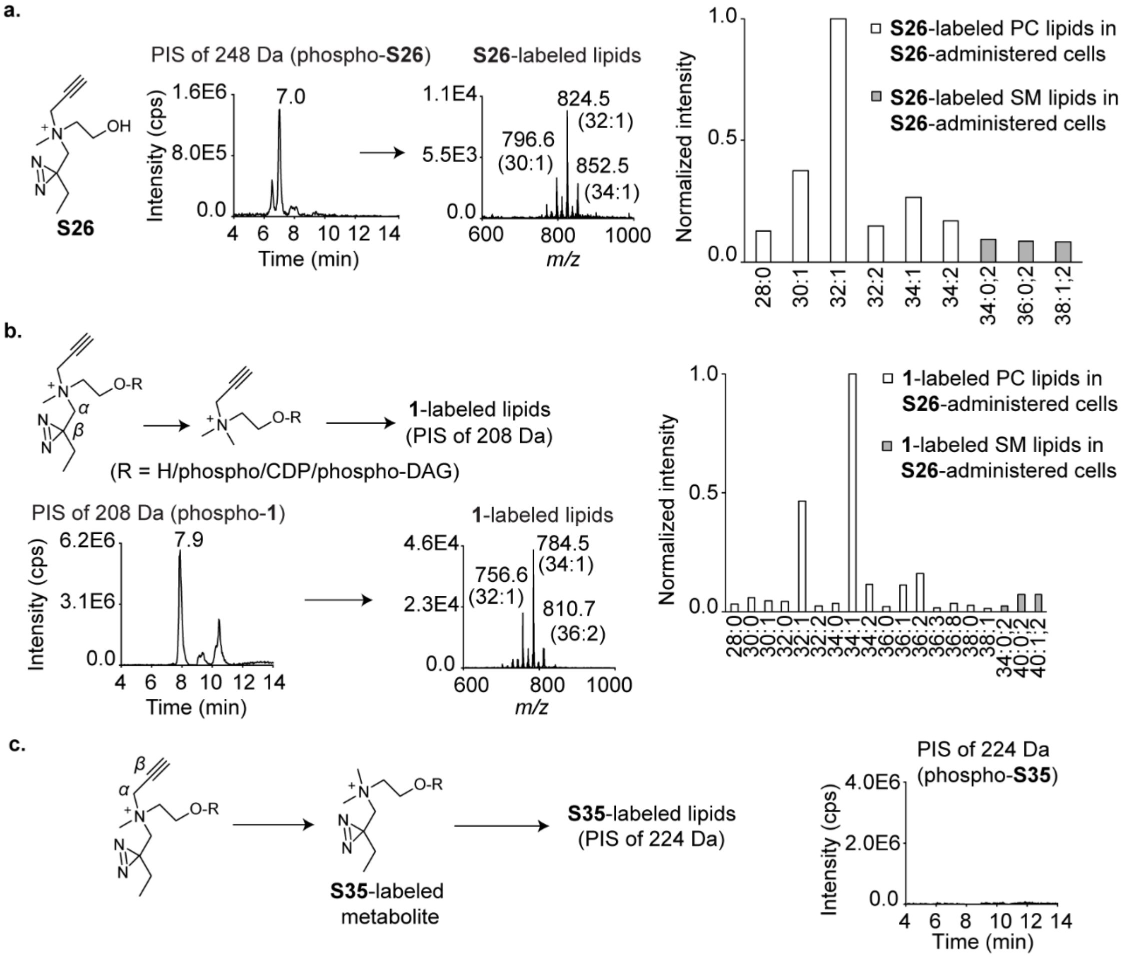


**Figure S22.** a) Characterization of the metabolically labeled choline lipids produced in HEK293 cells cultured in presence of **S26**. a) Left: Lipids extracted from HEK293 cells cultured in the presence of **S26** (2 mM) for 24 h were subjected to LC-MS/MS and analyzed for the formation of **S26**-labeled lipids by employing a PIS of 248 Da. Right: A bar graph obtained from the LipidView software denoting all **S26**-labeled PC and SM lipids identified from the XIC on the left panel. The *y*-axis has been normalized to the most intense peak *i.e.*, lipid 32:1 of the XIC. b) Left: Lipids extracted from HEK293 cells that were cultured in presence of **S26** (2 mM) for 24 h were subjected to LC-MS/MS and analyzed for the formation of **1**-labeled lipids by employing a PIS of 208 Da. Right: A bar graph obtained from the LipidView software denoting all **1**-labeled PC and SM lipids identified from the XIC on the left panel. The *y*-axis has been normalized to the most intense peak *i.e.*, lipid 34:1 of the XIC. c) Lipids extracted from HEK293 cells cultured for 24 h in the presence of **S26** (2 mM) were subjected to LC-MS/MS and analyzed for the formation of **S35**-labeled lipids by employing a PIS of 224 Da. The resultant TIC shows no peak for **S35**-labeled lipids.

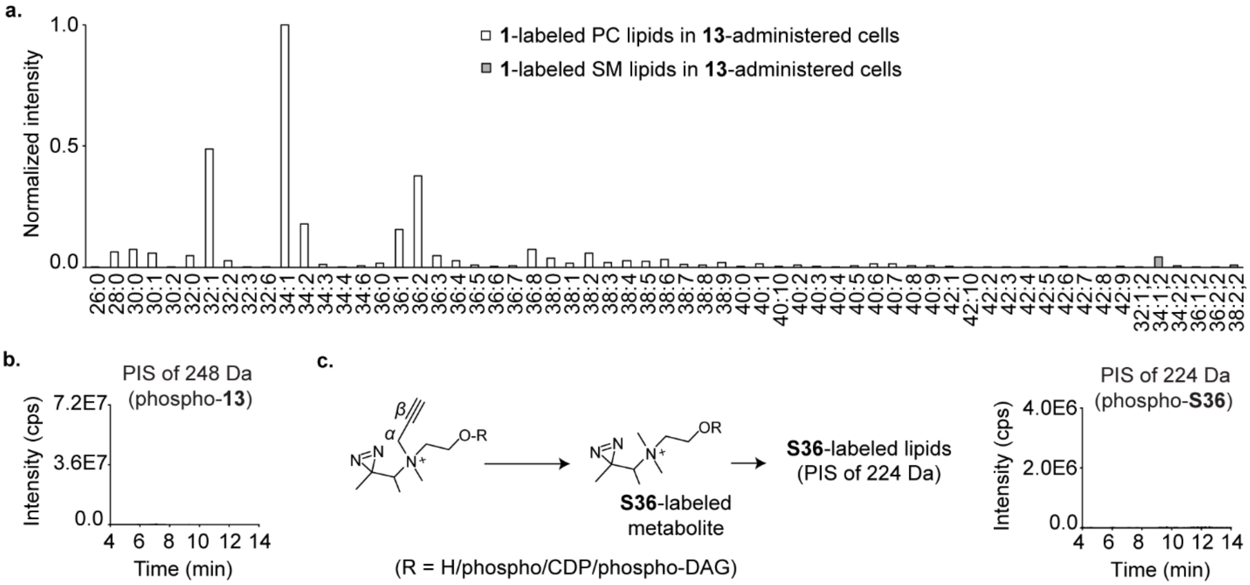


**Figure S23**. a) LipidView analysis of **1**-labeled PC and SM lipids obtained from HEK293 cells cultured for 24 h in the presence of **13** (2 mM). These lipids were identified from the XIC depicted in Figure 3f (third panel from the top) of the main text. The *y*-axis has been normalized to the most intense peak *i.e.*, lipid 34:1 of the XIC. b) Lipids extracted from HEK293 cells cultured for 24 h in the presence of **13** (2 mM) were subjected to LC-MS/MS and analyzed for the formation of **13**-labeled lipids by employing a PIS of 248 Da. The resultant TIC shows no peak for **13**-labeled lipids. c) Lipids extracted from HEK293 cells that were cultured in presence of **13** (2 mM) for 24 h were subjected to LC-MS/MS and analyzed for the formation of **S36**-labeled lipids by employing a PIS of 224 Da. The resultant TIC shows no peak for **S36**-labeled lipids.

### 10. Metabolic labeling studies on D9 choline 15

For each treatment, 3 million HEK293 cells in 10 mL of complete medium were seeded in two 10 cm culture dishes and incubated in a $CO_2$ incubator for 24 h. Subsequently, each flask was administered 200 µL of a solution of **15** in PBS (0.1 M, 0.5 M, and 1 M stocks yielding final concentrations of 2 mM, 10 mM, and 20 mM respectively) and after a 24 h-long incubation in a $CO_2$ incubator, the adhered cells were washed with ice-cold PBS (2 × 5 mL), resuspended in ice-cold PBS (5 mL) and subjected to lipid extraction as per the protocol described in sub-section 5A above. Cells not treated with **15** served as control. Subsequently, the extracted lipid samples were subjected to LC-MS/MS for the identification of **16**-labeled lipids using a PIS of 190 Da as described in sub-section 5 and Table S1 above. Figure 3g (bottom panel) shows the dose-dependent increase in the formation of **16**-labeled lipids.

### 11. Metabolic labeling studies on the γ-diazirine analogs 17 and 18

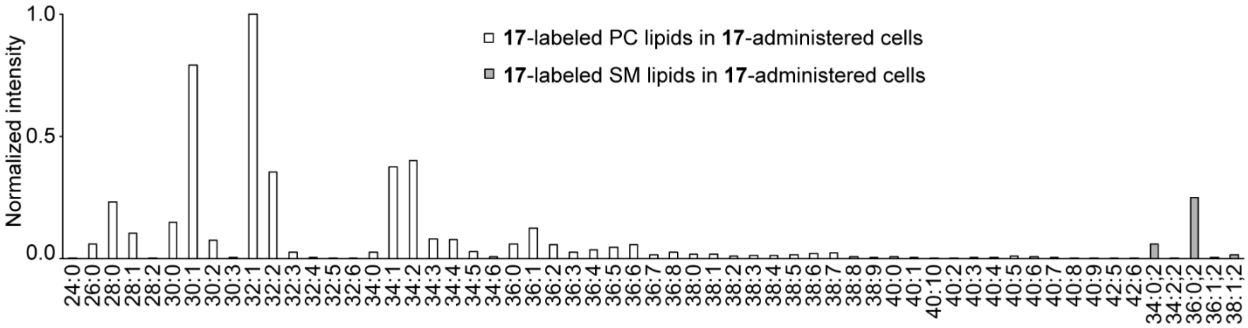


**Figure S24**. LipidView analysis of **17**-labeled PC and SM lipids identified from the XIC in Figure 4a of the main text. The *y*-axis has been normalized to the most intense peak *i.e.*, lipid 32:1 of the XIC.

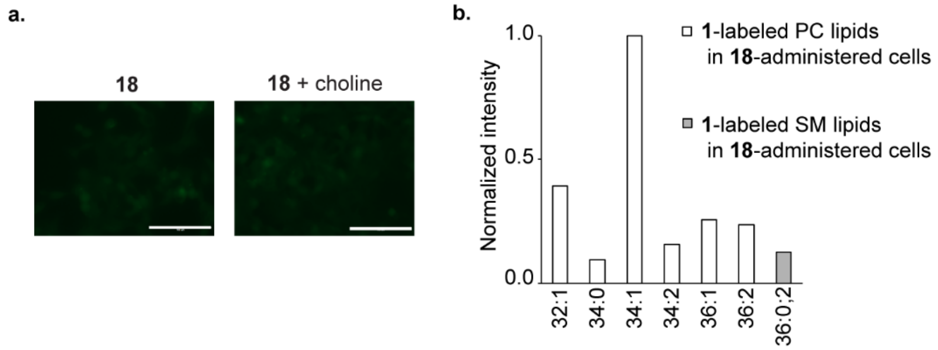


**Figure S25**. a) HEK293 cells cultured for 24 h in the presence of **18** (2 mM; image on the left), and **18** and choline (2 mM each; image on the right) were fixed, treated with 5-azido fluorescein under standard click chemistry conditions, and visualized using the EVOS imaging system under the GFP channel. The scale bars represent 100 μm. b) LipidView analysis of **1**-labeled PC and SM lipids obtained from HEK293 cells cultured for 24 h in the presence of **18** (2 mM). These lipids were identified from the XIC depicted in Figure 4c (bottom panel) of the main text. The *y*-axis has been normalized to the most intense peak *i.e.*, lipid 34:1 of the XIC.

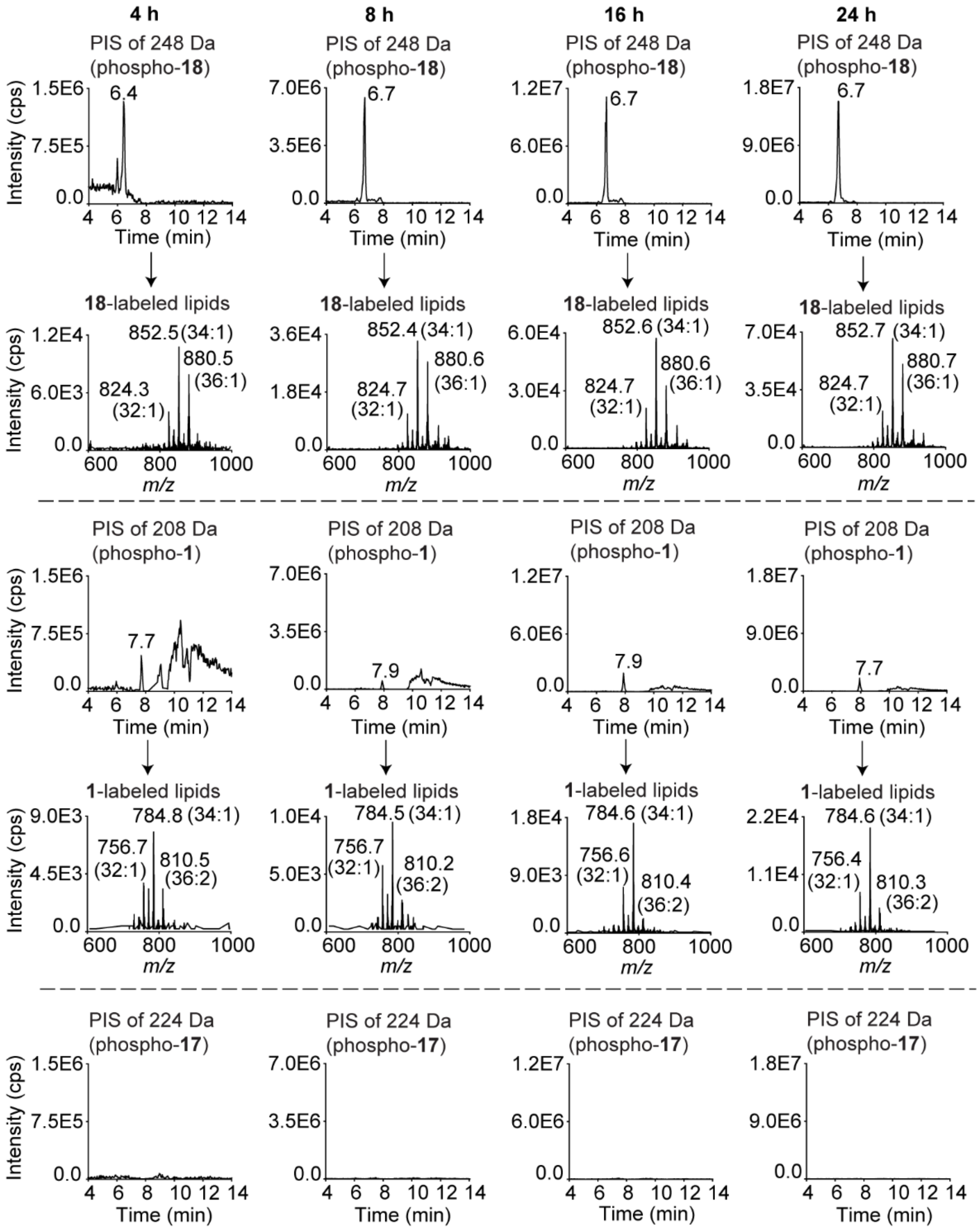


**Figure S26.** Kinetics for the formation of the metabolically labeled choline lipids in HEK293 cells cultured in presence of **18**. Lipids extracted from HEK293 cells that were administered with **18** (2 mM) for 4 h, 8 h, 16 h, and 24 h were subjected to LC-MS/MS and analyzed for the formation of **18**, **1**, and **17**-labeled lipids by employing a PIS of 248 Da, 208 Da, and 224 Da respectively. Each chromatogram is representative of three biological replicates. The area under the TIC peaks were manually calculated using the Analyst software to generate the plot on Figure 4e and 4f (left) of the main text.

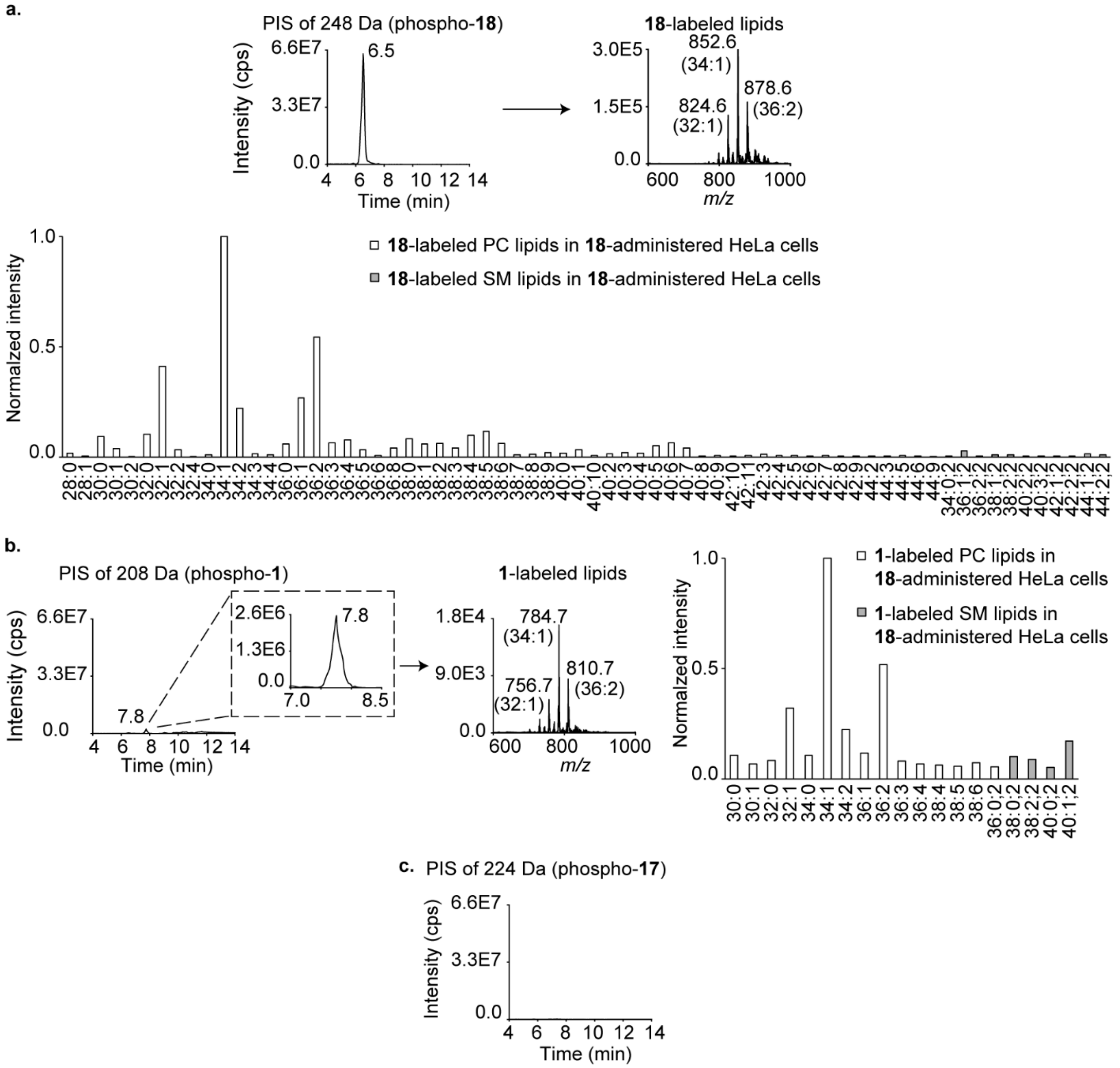


**Figure S27.** Characterization of the metabolically labeled choline lipids produced in HeLa cells cultured in presence of **18**. a) Top: Lipids extracted from HeLa cells cultured in presence of **18** (2 mM) for 24 h were subjected to LC-MS/MS and analyzed for the formation of **18**-labeled lipids by employing a PIS of 248 Da. The area under the TIC peaks were manually calculated using the Analyst software to generate the plot on Figure 4f (middle) of the main text. Bottom: A bar graph obtained from the LipidView software denoting all **18**-labeled PC and SM lipids identified from the XIC in the top panel. The *y*-axis has been normalized to the most intense peak *i.e.*, lipid 34:1 of the XIC. b) Left: Lipids extracted from HeLa cells cultured for 24 h in presence of **18** (2 mM) were subjected to LC-MS/MS and analyzed for the formation of **1**-labeled lipids by employing a PIS of 208 Da. The area under the peak on the TIC was manually calculated using the Analyst software to generate the plot on Figure 4f (middle) of the main text. Right: A bar graph obtained from the LipidView software denoting all **1**-labeled PC and SM lipids identified from the XIC in the left panel. The *y*-axis has been normalized to the most intense peak *i.e.*, lipid 34:1 of the XIC. c) Lipids extracted from HeLa cells cultured for 24 h in presence of **18** (2 mM) were subjected to LC-MS/MS and analyzed for the formation of **17**-labeled lipids by employing a PIS of 224 Da. The resultant TIC shows no peak for **17**-labeled lipids.

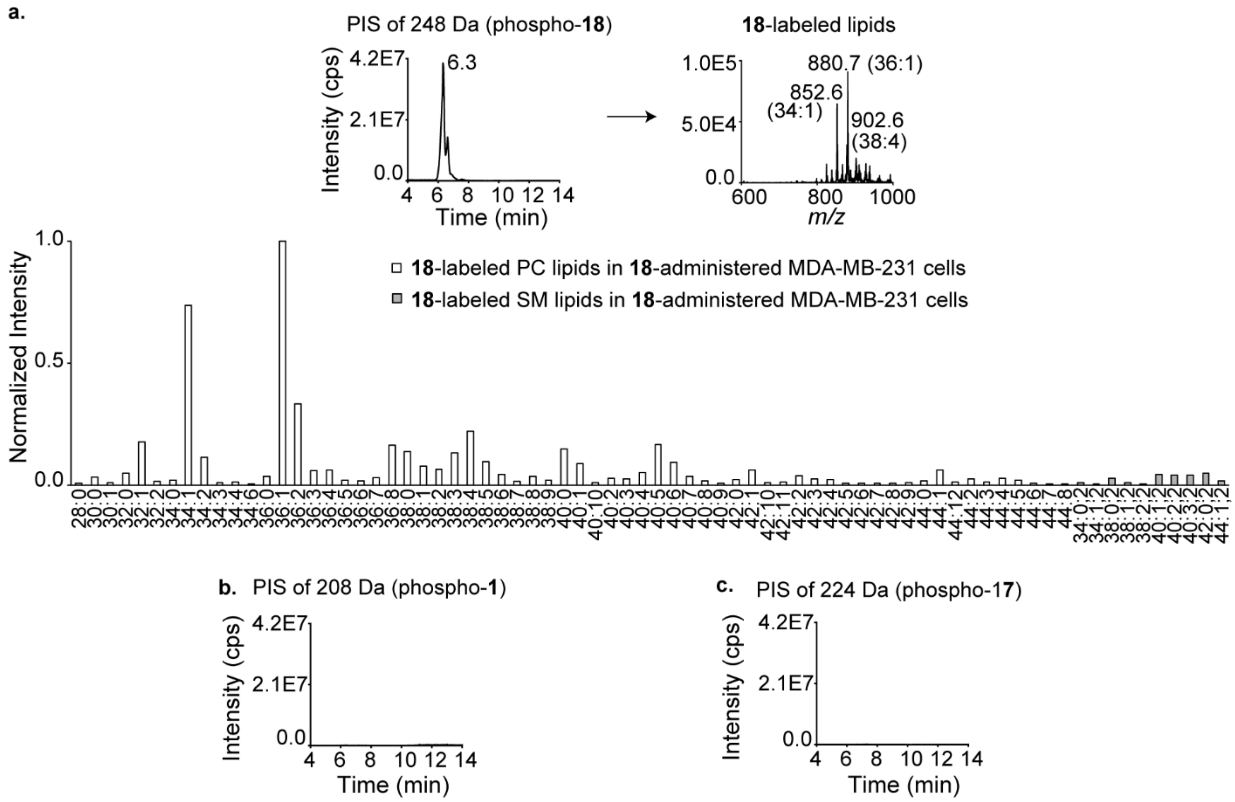


**Figure S28.** Characterization of the metabolically labeled choline lipids produced in MDA-MB-231 cells cultured in presence of **18**. a) Top: Lipids extracted from MDA-MB-231 cells cultured for 24 h in presence of **18** (2 mM) were subjected to LC-MS/MS and analyzed for the formation of **18**-labeled lipids by employing a PIS of 248 Da. The area under the TIC peaks were manually calculated using the Analyst software to generate the plot on Figure 4f (right) of the main text. Bottom: LipidView analysis of **18**-labeled PC and SM lipids identified from the XIC in the top panel. The *y*-axis has been normalized to the most intense peak *i.e.*, lipid 36:1 of the XIC. b) Lipids extracted from MDA-MB-231 cells cultured for 24 h in presence of **18** (2 mM) were subjected to LC-MS/MS and analyzed for the formation of **1**-labeled lipids by employing a PIS of 208 Da. The resultant TIC shows no peak for **1**-labeled lipids. c) Lipids extracted from MDA-MB-231 cells cultured for 24 h in presence of **18** (2 mM) were subjected to LC-MS/MS and analyzed for the formation of **17**-labeled lipids by employing a PIS of 224 Da. The resultant TIC shows no peak for **17**-labeled lipids.

### 12. In-gel fluorescence experiments

A. <u>Gel-based analysis of crosslinked proteins in HEK293 cells</u>: Six 10 cm culture dishes each containing HEK293 cells (10 mL of a 0.2 million cells per mL culture in complete media) were placed in a $CO_2$ incubator for 12 h. Subsequently, 317 µL of a 65 mM solution of either **1**, **11**, **12**, **13**, **S26**, or **18** in PBS was administered into each flask (final concentration of each of these choline analogs in the flask was 2 mM) and the cultures were allowed to grow in the $CO_2$ incubator for another 24 h. Cells not treated with any choline analog served as control. After the incubation, cells were washed with ice-cold DMEM containing 10% FBS (1 × 5 mL) and then with ice-cold PBS (2 × 5 mL).

Subsequently, cells were overlaid with ice-cold PBS (10 mL) and exposed to 356 nm UV as described in sub-section 1 above for 30 min on ice. After the UV exposure, cells were scraped into ice-cold PBS (5 mL) and pelleted by centrifugation (1,000 *g*, 5 min, 4 °C). The resultant cell pellet was resuspended by gentle pipetting in PBS containing EDTA-free protease inhibitor cocktail from Roche (467 µL) and incubated on ice for 30 min. Cells were lysed by probe sonication on ice (8 pulses of 5 sec each with a gap of 5 sec between each pulse, 30% amplitude on a Branson 450 Digital Sonifier), and to the cell lysate (50 µL containing 1 mg/mL protein concentration estimated by employing the Bradford assay), click chemistry reagents were added. Specifically, to this 50 µL cell lysate, 1 µL of a solution of 50 mM $CuSO_4.5H_2O$ in water (final concentration, 0.9 mM), 1 µL of a solution of 50 mM TCEP in PBS (final concentration, 0.9 mM), 3 µL of a solution of 1.7 mM TBTA in 1:4 *t*-butanol:DMSO (final concentration, 0.09 mM), and 1 µL of a 1.25 mM solution of Alexa Fluor 594 azide in DMSO (final concentration, 22 µM) were added and the mixture was incubated for 1 h at room temperature under dark after mixing by pipetting. For the azido analog **12**, Alexa Fluor 594 azide was replaced with 1 µL of a 1.25 mM solution of Alexa Fluor 594 alkyne in DMSO (final concentration, 22 µM). The click reactions were quenched by heating the samples with 2X SDS loading dye (56 µL) for 5 min at 98 °C and the samples were then subjected to SDS-PAGE analysis (10% gel) followed by fluorescence imaging of the gel by GE Typhoon Trio system with 532 nm excitation, and 610 nm (BP30) emission filter. Finally, the gel was stained with coomassie blue and the gel images thus obtained are depicted in Figure 5c of the main text.

B. UV-irradiation time optimization for crosslinking of **18**-labeled metabolites in HEK293 cells: Five 10 cm culture dishes each containing HEK293 cells (10 mL of a 0.2 million cells per mL culture in complete media) were placed in a $CO_2$ incubator for 12 h. Subsequently, 317 µL of a 65 mM solution of **18** in PBS was administered into each flask (final concentration of **18** in each flask was 2 mM) and the cultures were allowed to grow in the $CO_2$ incubator for another 24 h. After the incubation, cells in each plate were washed with ice-cold DMEM containing 10% FBS (1 × 5 mL) and then with ice-cold PBS (2 × 5 mL). Subsequently, cells were overlaid with ice-cold PBS (10 mL) and each plate was exposed to 356 nm UV as described in sub-section 1 above for either 0, 15, 30, 45, or 60 min on ice. After the UV light exposure, cells from each plate were lysed and subjected to the in-gel fluorescence protocol described in sub-section 12A above. The in-gel fluorescence and the coomassie stained gel images thus obtained are depicted in Figure 5b of the main text.

C. Dose dependent crosslinking of **18** in HEK293 cells: Ten 10 cm culture dishes each containing HEK293 cells (10 mL of a 0.2 million cells per mL culture in complete media) were placed in a $CO_2$ incubator for 12 h. Subsequently, 15.4 µL, 77 µL, 154 µL, and 317 µL of a 65 mM solution of **18** in PBS was administered into two plates each (final concentration of **18** were 0.1 mM, 0.5 mM, 1 mM, and 2 mM respectively) and the cultures were allowed to grow in the $CO_2$ incubator for another 24 h. Two flasks containing cells not administered with **18** served as control. After the incubation, the cells were subjected to the in-gel fluorescence protocol described in sub-section 12A above. The in-gel fluorescence and the coomassie stained gel images thus obtained are depicted in Figure 5a of the main text.

D. Membrane protein fractionation of crosslinked proteins in HEK293 cells: Six 10 cm culture dishes each containing HEK293 cells (10 mL of a 0.4 million cells per mL culture in complete media) were placed in a $CO_2$ incubator for 12 h. Subsequently, 317 µL of a 65 mM solution of **18** in PBS was added to each flask (final concentration of **18** in each flask was 2 mM) and the cultures were incubated in the $CO_2$ incubator for another 24 h. After the incubation, the cells were washed with ice-cold DMEM containing 10% FBS (5 mL) and then with ice-cold PBS (2 × 5 mL), and then overlaid with ice-cold PBS (10 mL) before irradiating them with 356 nm UV radiation as described in sub-section 1 above for 30 min on ice. Subsequently, the cells were scraped into ice-cold PBS (5 mL) and pelleted by centrifugation (1,000 *g*, 15 min, 4 °C). The supernatant was decanted and the cell pellet was resuspended in an ice-cold solution of PBS containing EDTA-free protease inhibitor cocktail (3 mL) by gentle pipetting and the cells were lysed using probe sonication as per the protocols described above in sub-section 12A. This cell lysate was centrifuged (180 *g*, 3 min, 4 °C) to remove any unlysed cells and the resultant supernatant was ultracentrifuged (100,000 *g*, 45 min, 4 °C) to separate the soluble proteins from the membrane proteins. The supernatant thus obtained contained the soluble cellular proteins. The membrane fraction pellet was washed with ice-cold PBS (5 × 5 mL), treated with a solution of 1% *w/v* SDS in EDTA-free protease inhibitor cocktail containing PBS (1 mL) and probe sonicated (12 pulses of 5 sec each with a gap of 10 sec between each pulse, 30% amplitude on a Branson 450 Digital Sonifier) to give a clear solution. Proteins in both soluble and membrane fractions were diluted to 1 mg/mL in PBS by employing the Bradford assay followed by subjecting them to click chemistry and in-gel fluorescence SDS-PAGE protocols described in sub-section 12A above. The in-gel fluorescence and the coomassie-stained gel images thus obtained are depicted in Figure S29 below.

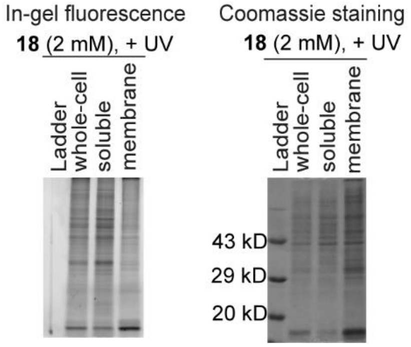


**Figure S29.** Membrane protein fractionation of crosslinked proteins in HEK293 cells cultured for 24 h in the presence of **18**. HEK293 cells administered with **18** (2 mM) were UV-irradiated and the cell lysates were fractionated into soluble and membrane proteins by ultracentrifugation. Each fraction was treated with Alexa Fluor 594 azide under standard click chemistry conditions and subjected to SDS-PAGE. The gel was visualized under a fluorescence imager (left) and subsequently stained with coomassie (right).

### 13. Biotinylation and affinity purification of crosslinked proteins

Three 10 cm culture dishes each containing HEK293 cells (10 mL of a 0.3 million cells per mL culture in complete media) were placed in a $CO_2$ incubator for 12 h. Subsequently, each dish was administered 317 µL of either a 16.3 mM, 32.5 mM or 65 mM solution of **18** in PBS (final concentrations of **18** were 0.5 mM, 1 mM and 2 mM respectively) and placed in a $CO_2$ incubator for another 24 h. The cells were then washed with ice-cold DMEM media containing 10% FBS (5 mL) and then with ice-cold PBS (2 × 5 mL), overlaid with ice-cold PBS (10 mL) and exposed to 356 nm UV as described in sub-section 1 above for 30 min on ice. Cells subjected to the same treatment as described above but not exposed to UV light served as the -UV control. After the +UV or -UV treatments, cells were scraped into ice-cold PBS (5 mL) and pelleted by centrifugation (1,000 *g*, 5 min, 4 °C). The resultant cell pellets were resuspended by gentle pipetting in PBS containing EDTA-free protease inhibitor cocktail from Roche (467 µL) and incubated on ice for 30 min. The cells were lysed by probe sonication on ice (8 pulses of 5 sec each with a gap of 5 sec between each pulse, 30% amplitude on a Branson 450 Digital Sonifier), and the protein concentration in the cell lysate was estimated by employing the Bradford assay. The cell lysate was then subjected to click chemistry reagents as reported previously.[10] Specifically, to a volume of the cell lysate containing 1 mg of protein, was added 20 µL of a solution of 25 mM biotin azide in DMSO (final concentration, 0.5 mM), 20 µL of a solution of 50 mM TCEP in PBS (final concentration, 1 mM), 59 µL of a solution of 1.7 mM TBTA in 1:4 *t*-butanol:DMSO (final concentration, 0.1 mM ), and 20 µL of a solution of 50 mM $CuSO_4.5H_2O$ in water (final concentration, 1 mM). The total volume of this reaction mixture was made up to 1 mL by adding PBS. The mixture was then incubated at room temperature for 90 min. Subsequently, the protein was extracted from this reaction mixture by performing methanol-chloroform extraction. Briefly, ice-cold methanol (480 µL) and ice-cold chloroform (160 µL) were added to the reaction mixture and the resultant heterogeneous mixture was vortexed for 1 min. Water (400 µL at room temperature) was then added to this mixture followed by vortexing for 1 min and the mixture was centrifuged (14,000 *g*, 5 min, room temperature) to achieve phase separation between an aqueous layer on the top and an organic layer at the bottom with a cloudy layer of protein at the interface of the two layers. The aqueous layer at the top was removed by aspiration and to the remaining cloudy layer of protein and the lower organic layer, ice-cold methanol (300 µL) was added followed by vortexing for 1 min. Subsequently, the proteins were pelleted by centrifugation (14,000 *g*, 5 min at room temperature) and the supernatant was aspirated off. To the resulting pellet was added ice-cold methanol (480 µL) and ice-cold chloroform (160 µL) and the mixture was subjected to methanol-chloroform extraction exactly as mentioned above and the pellet thus obtained was resuspended in 1% *w/v* SDS in PBS (800 µL) by vortexing for 2 min, probe sonicated at room temperature (6 pulses of 5 sec each with a gap of 5 sec between each pulse, 60% amplitude on a Branson 450 Digital Sonifier) and incubated at 37 °C for 15 min (with shaking at 1,400 rpm on an Eppendorf Thermomixer). The resulting solution was centrifuged (2,000 *g*, 5 min at room temperature) to separate any insoluble material and the supernatant was transferred to a centrifuge tube and diluted with PBS to a final volume of 4 mL so that the final SDS concentration was reduced to 0.2% *w/v*. This solution was set aside and avidin-agarose resin (200 µL of a 50% *v/v* suspension in glycerol) was washed with PBS at room temperature. Specifically, the resin was resuspended in PBS (500 µL) by gentle pipetting and the resulting suspension was centrifuged (400 *g*, 5 min at room temperature) and the supernatant was aspirated off. The pellet containing avidin-agarose resin was washed again by resuspension in PBS (500 µL) by gentle pipetting and centrifugation (400 *g*, 5 min at room temperature) followed by removal of the supernatant. The pellet thus obtained contained washed avidin-agarose resin that was further resuspended in 100 µL PBS and this suspension was transferred to the aforementioned 4 mL solution of methanol-chloroform extracted proteins dissolved in 0.2% *w/v* SDS in PBS. This mixture was incubated for 2 h at room temperature with constant rotation (10 rpm) on a 360° vertical rotating mixer (Biobase). Subsequently, the resin was spun down from the suspension (400 *g*, 5 min at room temperature) and the supernatant was removed. The resin was then washed with 0.2% *w/v* SDS solution in PBS (10 × 1 mL) and then with PBS (3 × 1 mL) as per the procedure described above that was used to wash the avidin-agarose resin with PBS. To elute the proteins bound to the resin, it was treated with 2X SDS loading dye (100 µL) and incubated at room temperature for 15 min, then heated at 98 °C for 5 min. This mixture was briefly centrifuged (5,000 *g*, 1 min at room temperature) and the supernatant was collected as the desired eluate. This eluate was separated by SDS-PAGE (10% gel) followed by electrophoretically transferring the proteins to a PVDF membrane. The membrane was then treated with 5% *w/v* BSA solution in PBS for 1 h at room temperature, incubated with anti-biotin antibody (details in Table S2) for 3 h at room temperature. Subsequently, the membrane was washed three times with PBST and incubated with shaking in PBST for 10 min after each wash and then it was incubated in anti-mouse IgG conjugated to HRP for 2 h at room temperature (details in Table S2). Finally, the membrane was washed with PBST three times with intermittent 10 min incubations in PBST with shaking. To visualize the proteins present in the eluate, the blot was developed using the ECL kit (Invitrogen) as per the instructions in manufacturer's protocol and imaged for chemiluminescence using ChemiDoc imaging system (Biorad). The blot thus obtained is depicted in Figure 5d of the main text.

### 14. Sample preparation for quantitative proteomics (TMT) and target validation by Western blotting

For protein identification, the same procedure for the isolation of crosslinked proteins as described above in sub-section 13 was followed except that instead of three 10 cm culture dishes, six culture dishes were administered with a final concentration of 2 mM of **18** (three +UV and three -UV experiments), and the avidin column-bound proteins were not eluted. Instead, the protein-bound avidin-agarose resin thus obtained was outsourced to the Thermo Fisher Scientific Center for Multiplexed Proteomics at the Harvard Medical School (https://tcmp.hms.harvard.edu/) where it was subjected to the sixplex tandem mass tag (TMT) analysis on the three +UV and the three -UV samples (experimental details below in sub-section **15**) that yielded a list of 902 proteins that were identified in all the six samples (Sheet 1 of Table S3). For each sample, the raw intensities of all the peaks for peptides obtained from individual proteins were added to yield a quantitative estimate of the amount of that protein. The resulting values obtained for the three +UV samples were averaged and divided by the averaged value obtained from the three -UV samples, to yield the enrichment ratio for each protein (the raw data for each of the six samples is provided in Sheet 1 of Table S3). Unpaired two-tailed *t*-test was done on the raw intensity values obtained for each protein from the set of three +UV and the set of three -UV samples. The 674 proteins that gave the enrichment ratio (+UV/-UV) of $\geq 2$ and p-value $\leq 0.05$ were deemed "high-confidence" protein hits (Sheet 1 of Table S3). The volcano plot for -log(p-value) *vs* $\log_2$(enrichment ratio) for all the 902 proteins is depicted in Figure 5e of the main text. Seven proteins from the list of high confidence protein hits (p32, nucleolin, COX 4, ezrin, PGRMC1, VDAC1, and annexin A2) were selected for the validation by Western blot. For validation of the aforementioned proteins, the same procedure for the avidin-enrichment of the crosslinked proteins as described above in sub-section 13 was followed except that the experiments were performed on one 10 cm culture dish administered with a final concentration of 2 mM of **18**. Instead of blotting against anti-biotin as was done in the procedure described in sub-section 13, the BSA blocked membrane was incubated with either of the following primary antibodies: anti-p32, anti-nucleolin, anti-COX 4, anti-ezrin, anti-PGRMC1, anti-VDAC1, or anti-annexin A2 followed by incubation with the secondary antibody conjugated to HRP. The details of the primary antibodies and the secondary antibodies conjugated to HRP are described in Table S2 below. The blot was developed using the ECL kit (Invitrogen) as per the instructions in manufacturer's protocol and imaged for chemiluminescence using LAS-3000 imager (Fujifilm). The blot thus obtained is depicted in Figure 5f of the main text.

**Table S2**. Details of antibodies used for Western blot and ELISA-based binding assays

| S. No. | Protein name | Manufacturer | Catalog No. | Dilution used | Data Figure |
|---|---|---|---|---|---|
| 1 | Anti-p32 | Abcam | ab24733 | 1:1000 | Figure 5g |
| 2 | Anti-nucleolin | Abcam | ab129200 | 1:1000 | Figure 5g |
| 3 | Anti-COX 4 | Cell Signaling Technology | 4850 | 1:1000 | Figure 5g |
| 4 | Anti-ezrin | Abcam | ab4069 | 1:1000 | Figure 5g |
| 5 | Anti-PGRMC1 | Cell Signaling Technology | 13856 | 1:1000 | Figure 5g |
| 6 | Anti-VDAC1 | Abcam | ab154856 | 1:1000 | Figure 5g |
| 7 | Anti-annexin A2 | Abcam | ab54771 | 1:1000 | Figure 5g |
| 8 | Anti-biotin | Santa Cruz | sc-53179 | 1:4000 | Figures 5e and S30c |
| 9 | Streptavidin-HRP | Invitrogen | 434323 | 1:5000 | Figure 7a, 7c-e |
| 10 | Goat-anti rabbit IgG conjugated to HRP | Abcam | ab205718 | 1:10000 | Figure 5g |
| 11 | Goat anti-mouse IgG conjugated to HRP | Abcam | ab205719 | 1:10000 | Figures 5e, 5g, and S30c |

**15. Quantitative proteomics (TMT) experiments**

A. <u>On-bead digestion:</u> The avidin-agarose resin containing the enriched proteins (obtained as described in sub-section 14 above) was washed three times with 50 mM Tris-HCl buffer at pH 8.0 followed by resuspension in a solution of 1 M urea in 50 mM Tris-HCl at pH 8.0. The initial protein digestion was performed with trypsin (5 ng/µL) at 37 °C for 1 h. After the initial trypsin digestion, samples were centrifuged briefly and the supernatant was collected. The beads were further washed three times with a solution of 1 M urea in 50 mM Tris-HCl at pH 8.0. All the bead washes were pooled in the same tube and digested overnight at room temperature by incubating in trypsin (5 ng/µL). Digested peptides were reduced with 5 mM DTT followed by alkylation with 15 mM iodoacetamide, quenching alkylation with 10 mM DTT and finally quenching the digestion process with TFA. Acidified digested peptides were desalted over $C_{18}$STAGE tips. Desalted peptides were reconstituted with 200 mM EPPS buffer at pH 8.0 and labeled with sixplex tandem mass tag (TMT) reagents. The sixplex labeling reactions were performed for 2 h at room temperature. Modification of tyrosine residue with TMT was reversed by the addition of 0.5% hydroxylamine for 15 min and the reaction was quenched with 0.5% TFA. Samples were combined at a 1:1:1:1:1:1 ratio and the resultant sample was further desalted using a STAGE tip and reconstituted with 5% acetonitrile + 5% formic acid for mass spectrometric analysis.

B. Liquid chromatography-MS3 spectrometry (LC-MS3): The labeled peptide sample from the previous sub-section 15B was analyzed with an LC-MS3 data collection strategy on an Orbitrap Fusion mass spectrometer (Thermo Fisher Scientific) equipped with a Thermo Easy-nLC 1200 for online sample handling and peptide separations.[11] The resuspended peptide from the previous step was loaded onto a 100 µm inner diameter fused-silica micro capillary with a needle tip pulled to an internal diameter less than 5 µm. The column was packed in-house to a length of 35 cm with a $C_{18}$ reverse phase resin (GP118 resin 1.8 µm, 120 Å, Sepax Technologies). The peptides were separated using a 180 min linear gradient from 3% to 25% buffer B (100% acetonitrile + 0.125% formic acid) equilibrated with buffer A (3% acetonitrile + 0.125% formic acid) at a flow rate of 600 nL/min across the column. The scan sequence for the Fusion Orbitrap began with an MS1 spectrum (Orbitrap analysis, resolution 120,000, 400−1400 *m/z* scan range, AGC target $2 \times 10^5$, maximum injection time 100 ms, dynamic exclusion of 90 seconds). The "Top10" precursors were selected for MS2 analysis, which consisted of CID (quadrupole isolation set at 0.5 Da and ion trap analysis, AGC $8 \times 10^3$, Collision Energy 35%, maximum injection time 150 ms). The top ten precursors from each MS2 scan were selected for MS3 analysis (synchronous precursor selection), in which precursors were fragmented by HCD prior to Orbitrap analysis (Collision Energy 55%, max AGC $1 \times 10^5$, maximum injection time 150 msec, isolation window 2.5 Da, resolution 60,000).

C. LC-MS3 data analysis: A suite of in-house software tools were used to for .RAW file processing and controlling peptide and protein level false discovery rates, assembling proteins from peptides, and protein quantification from peptides as previously described. MS/MS spectra were searched against a UniProt Human database with both the forward and reverse sequences. Database search criteria are as follows: tryptic with two missed cleavages, a precursor mass tolerance of 50 ppm, fragment ion mass tolerance of 1.0 Da, static alkylation of cysteine (57.02146 Da), static TMT labeling of lysine residues and *N*-termini of peptides (229.162932 Da), and variable oxidation of methionine (15.99491 Da). TMT reporter ion intensities were measured using a 0.003 Da window around the theoretical *m*/*z* for each reporter ion in the MS3 scan. Peptide spectral matches with poor quality MS3 spectra were excluded from quantitation (<100 summed signal-to-noise across 10 channels and <0.5 precursor isolation specificity).

### 16. Analysis of the high-confidence proteins obtained from proteomics studies

To identify the subcellular localization of all the 674 high-confidence protein hits, their corresponding UniProt IDs were submitted to the retrieve/ID mapping feature of the UniProt database. The proteins were characterized into seven categories: lysosome, golgi, ER, cell membrane, mitochondria, nucleus, and cytoplasm and the total number of proteins in each location were counted manually. The list thus obtained is provided in Sheet 2 of Table S3 and a graph summarizing this data is depicted in the Figure 5g of the main text. The total number of proteins among the seven categories was higher than 674 because of multiple annotations for several proteins. The same UniProt database was used to characterize these 674 high-confidence proteins as membrane or soluble. Proteins that possessed known or predicted transmembrane domains were labeled as membrane proteins and the remaining ones were considered soluble. The resulting data is depicted in Sheet 2 of Table S3. Using the same retrieve/ID mapping feature, the 674 UniProt IDs were also sorted for subclasses of enzymes namely, oxidoreductases, transferases, hydrolases, lyases, isomerases, and ligases. The number of proteins belonging to each enzyme subclass were counted manually and a plot summarizing this data is depicted in Figure 5i of the main text and the protein-wise list thus obtained is provided in Sheet 7 of Table S3. The retrieve/ID mapping tool on UniProt database was also used to categorize proteins as DrugBank targets (data depicted in Figure 5k and Sheet 10 of Table S3).

In addition to this analysis, the UniProt IDs of the 674 protein hits were also submitted to the PANTHER classification system resulting in the classification of 449 of these proteins into sixteen functional classes—translational proteins, cytoskeletal proteins, receptors, nucleic acid metabolism proteins, transfer/carrier proteins, chaperones, adaptors, transporters, metabolite interconversion enzymes, protein modifying enzymes, membrane traffic proteins, protein-binding activity modulators, calcium-binding proteins, intercellular signal molecules, gene-specific transcriptional regulators, and chromatin binding/regulatory proteins. The list thus obtained is depicted in Sheet 6 of Table S3 and a plot summarizing the data on the ten most abundant protein classes is depicted in Figure 5h of the main text.

The gene IDs of the 674 high-confidence protein hits were analyzed by the DAVID Bioinformatics Database to categorize proteins that are genetically liked to the following disease classes—aging, cancer, cardiovascular, developmental, hematological, immune, infection, metabolic, neurological, pharmacogenomic, psychiatric, renal, and reproduction (Sheet 8 of Table S3 and Figure 5j).

To determine whether the mRNAs of our 674 high-confidence protein hits are known to be overexpressed in any type of cancer, the corresponding gene IDs of each of the high-confidence protein hits were manually submitted to the Oncomine database and the results of this analysis are summarized in Sheet 9 of Table S3.

**17. Characterizing the binding of p32 to choline metabolites and lipids**

A. Recombinant production of His tagged p32: A plasmid containing the gene encoding for the *N*-terminus 6X His tagged p32 protein (henceforth referred to as His-p32) in the pET15b vector was used to transform *E. coli* BL21 (DE3) cells. LB broth cultures (1 L) containing ampicillin (100 µg/mL) were inoculated with the transformed *E. coli* BL21 (DE3) cells and induced with IPTG (0.5 mM) on reaching an O.D. (600 nm) of 0.4-0.6 at 37 °C. The induced culture was pelleted (6,000 rpm, 10 min at room temperature) after 6 h of growth and the supernatant was discarded. The efficiency of the induction was evaluated by performing SDS-PAGE analysis (Figure S30a). Consistent with the previous reports,[12] although the molecular weight of His-p32 is 26.1 kDa, we observed that it gave a band at ~34 kDa. To purify the protein, the cell pellet was resuspended by vigorous vortexing in 20 mL of HEPES buffer (20 mM at pH 7.6) containing 0.5 M NaCl, 1.5 mM $MgCl_2$, 1 mM DTT, 0.1% (*v/v*) TritonX-100, and 20% (*v/v*) glycerol (henceforth known as lysis buffer). The resuspended cells were lysed via probe sonication on ice (150 pulses of 2 sec each with a gap of 3 sec between each pulse, 30% amplitude on a Branson 450 Digital Sonifier) followed by centrifugation at 15,500 *g* at 4 °C for 30 min and the resulting supernatant was retained. A column packed with 1.5 mL of Ni-NTA resin (Qiagen) was washed with 20 mL of lysis buffer at 4 °C and the aforementioned cell lysate containing overexpressed His-p32 was applied to the resin at 4 °C. After collecting the flow-through, the column was washed twice with 5 mL of sodium phosphate buffer (50 mM at pH 6.0) containing 0.3 M NaCl, 10 mM imidazole, and 0.05% Tween-20 at 4 °C. The bound protein was eluted by applying 25 mL of sodium phosphate buffer (50 mM at pH 6.0) containing 0.3 M NaCl, 300 mM imidazole and 0.05% Tween-20 at 4 °C and the eluted fractions were collected as 5 mL aliquots. The collected fractions were characterized by subjecting 10 µL of each fraction to SDS-PAGE (Figure S30b). The fractions that contained the required His-p32 (fraction numbers 1-5) were combined and used for further processing as described in sub-section 17B and 17C below.

B. Testing the binding of choline metabolites with His-p32 using DARTS: The fractions containing His-p32 obtained as described in the sub-section 17A above were combined and dialyzed against Tris HCl buffer (50 mM at pH 8.0 containing 150 mM NaCl). Subsequently, the dialyzed protein was concentrated by using 5 kDa molecular weight cutoff Vivaspin spin concentrators (GE life sciences), and quantified by measuring the absorbance of the concentrated protein solution at 280 nm. A solution of 3.13 mg/mL His-p32 in Tris HCl buffer (50 mM at pH 8.0 containing 150 mM NaCl) was supplemented with EDTA-free protease inhibitor cocktail (from a 10X stock solution in water), 50 mM NaF (from a stock solution of 500 mM in water), 1 mM PMSF (from a stock solution of 100 mM in ethanol), and 1% *v/v* NP-40 (stock solution of 10% *v/v* in water). This protein solution was used to test the binding of His-p32 with choline metabolites and lipids by employing drug affinity responsive target stability (DARTS) assays.[13] To perform these assays, 40 µL binding reaction mixture samples (the "+" samples) were constituted that contained 2.5 mg/mL His-p32 (32 µL of the aforementioned 3.13 mg/mL stock solution was added), 10 mM $CaCl_2$ (2 µL of a 200 mM stock solution in water was added), and 1-50 mM choline/ CDP-choline/GPC/phosphocholine (a 50 mM metabolite stock solution was used for achieving the final concentrations of 1 mM and 5 mM in the samples, whereas a 500 mM stock solution in water was used for achieving concentrations of 10 mM, 25 mM, and 50 mM). These samples were incubated at 25 °C for 4 h with shaking at 600 rpm in an Eppendorf Thermomixer. In the control "-" samples for choline, CDP choline and GPC, these metabolites were replaced with NaCl at concentrations identical to those for the metabolites in the corresponding "+" samples. In the control "-" samples for phosphocholine, the metabolite was replaced with $CaCl_2$. To test the binding of His-p32 with 10-100 µg/mL PC and SM, these lipids were added from their 1 mg/mL stocks in DMSO to yield the final concentrations of 10 µg/mL, 25 µg/mL, 50 µg/mL, 75 µg/mL or 100 µg/mL. In the corresponding vehicle controls, instead of the lipid stock solutions, equal volume of DMSO was added. To each of these 40 µL reaction mixtures, 4 µL of an 83.3 ng/µL stock of pronase (stock solution made in 50 mM Tris-HCl buffer at pH 8.0 containing 50 mM NaCl and 10 mM $CaCl_2$) was added, and the mixture was incubated at 25 °C for 30 min without shaking (final pronase: protein *w/w* ratio in each sample was 1:300). To the control samples, instead of this pronase stock solution, 4 µL of 50 mM Tris-HCl buffer at pH 8.0 containing 50 mM NaCl and 10 mM $CaCl_2$ was added. The proteolysis reaction was quenched by adding 2X SDS-loading buffer (44 µL) in each sample and heating the resultant mixture at 75 °C for 10 min. The samples were then centrifuged (8,000 *g*, 3 min at room temperature) and the supernatants (15 µL) were subjected to SDS-PAGE (12%

gel) and stained with coomassie. To test the binding of 1-50 mM phosphocholine with another protein, the experiment described above was repeated with the 3.13 mg/mL stock solution of myoglobin instead of His-p32. The gel images thus obtained are depicted in Figure 6a-f of the main text.

C. Biotinylation of His-p32: The fractions containing His-p32 obtained as described in sub-section 17A above were combined and dialyzed against PBS. Subsequently, the dialyzed protein was concentrated by using 5 kDa molecular weight cutoff Vivaspin spin concentrators (GE life sciences) and quantified by measuring the absorbance of the concentrated protein solution at 280 nm. Biotinylation was performed by adding a solution of biotin *N*-hydroxysuccinimide ester in DMSO (950 nmol, 47 µL, 20 eq.) to a solution of His-p32 in PBS (47.5 nmol, 470 µL, 1 eq.). After incubating for 2 h at room temperature, the reaction mixture was desalted by using Zeba spin 7 kDa molecular weight cutoff desalting columns (Thermo Fisher Scientific) and the protein solution was then subjected to Western blotting against the anti-biotin antibody to characterize biotinylation. Specifically, the desalted protein solution was subjected to SDS-PAGE (12% gel) followed by electrophoretically transferring the protein to a PVDF membrane. The membrane was incubated in 5% *w/v* BSA solution in PBS for 1 h at room temperature and then in anti-biotin antibody (details in Table S2) for 3 h at room temperature. Subsequently, the membrane was washed three times with PBST and incubated with shaking in PBST for 10 min after each wash and then it was incubated in anti-mouse IgG conjugated to HRP for 2 h at room temperature (details in Table S2). Finally, the membrane was washed with PBST three times with intermittent 10 min incubations in PBST with shaking. To visualize the biotinylated His-p32, the blot was developed using the ECL kit (Invitrogen) as per the manufacturer's protocol and imaged for chemiluminescence using the ChemiDoc imaging system (Biorad) as depicted in Figure S30c.

D. Probing the binding between p32 and its ligands, Lyp-1, HA, Ac-Tat peptide and MCP1 using ELISA: The stock solutions of the ligands were made in PBS at the following concentrations: Ac-Tat peptide (5 mg/mL), MCP1 (11.5 mM), Lyp-1 (2 mg/mL) and HA (10 mg/mL). These solutions were diluted into the sodium carbonate-bicarbonate buffer (50 mM, pH 9.6) to achieve their respective working concentrations (25 µg/mL for Ac-Tat, 100 nM for MCP1 and 50 µg/mL for both Lyp-1 and HA) and 100 µL of each of these solutions were added to 96 well-plates and incubated for 12 h at 4 °C to immobilize the ligands. Subsequently, each well was washed with PBS (300 µL/well × 3) and blocked with 5% (*w/v*) skimmed milk solution in PBS (300 µL/well) for 2 h at room temperature. The skimmed milk solution was then aspirated off and the wells were washed three times with 0.05% (*v/v*) Tween-20 containing PBS (PBST, 300 µL/well) before introducing solutions (100 µL/well) containing different concentrations of biotinylated His-p32 in PBS (as indicated by the data points of the ELISA plots depicted in Figure 7 of the main text) and choline metabolites (at concentrations mentioned in the legend of Figure 7). Wells administered with 100 µL of PBS instead of biotinylated His-p32 served as blank. Subsequently, the plates were incubated at room temperature for 2 h following which, the solutions were aspirated off and the wells were washed three times with PBST (300 µL/well). The bound biotinylated p32 was detected by incubation in a solution containing HRP conjugated to streptavidin in 5% *w/v* BSA in PBS (100 µL/well, details in Table S2) for 2 h at room temperature. The unbound protein was washed off with PBST (300 µL/well × 3) and the wells were incubated in 100 µL of the substrate solution containing 1.82 mM ABTS (2,2'-azino-bis(3-ethylbenzothiazoline-6-sulfonic acid)) and 0.03% *w/w* hydrogen peroxide solution (diluted from a 30% *w/w* stock) in citrate-phosphate buffer (100 mM, pH 5.0) for 15 min at room temperature under dark. The reaction was stopped by adding sulfuric acid (625 mM, 100 µL/well) and the intensity of the resulting blue-green color was immediately quantified by measuring the absorbance at 405 nm by using an iMark microplate reader (Biorad). The readings obtained from the blank wells containing immobilized Lyp-1/HA/MCP1/Ac-Tat peptide, but not administered with biotinylated p32 were subtracted from those obtained from the experimental samples and these blank-subtracted values were used to generate a saturation binding curve (Figure 7a-d in the main text). Each value is an average of at least three technical replicates.

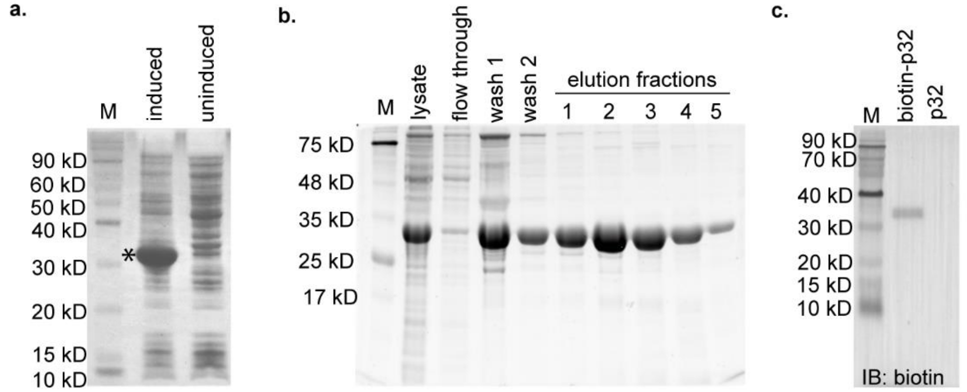


**Figure S30.** Recombinant production and biotinylation of His-p32. a) SDS-PAGE analysis for His-p32 overexpression in IPTG induced BL21 (DE3) cells as described in sub-section 17A. The overexpressed His-p32 band is indicated by the asterisk mark. b) An SDS-PAGE analysis of the Ni-NTA purification of His-p32 as described in sub-section 17A. c) Detection of biotinylated-p32 (loaded in the second lane) using Western blot against biotin as described in sub-section 17C. The third lane was loaded with purified p32 not subjected to biotinylation.

### 18. Investigations on the effects of choline metabolites on the function of enzymes

1. Lactate dehydrogenase (LDH): Enzymatic assays on rabbit muscle LDH (Sigma) were performed at 25 °C in a quartz cuvette by using a UV-Vis spectrophotometer (Perkin Elmer Lambda 25) as per a procedure described previously.[14] The stock solutions of choline, CDP choline, GPC and sodium chloride (20 mM each), phosphocholine and calcium chloride (100 mM each), sodium pyruvate and NADH were made in 0.2 M Tris-HCl buffer at pH 7.3 and the stock solutions of PC and SM lipids (2.5 mg/mL each) were made in DMSO. All enzyme assays were performed in 0.2 M Tris-HCl buffer at pH 7.3. To generate kinetic plots with varying NADH and constant pyruvate concentrations, the final concentrations of reaction components in the kinetic runs were as follows: LDH: 0.06 U/mL (from a stock of 1.2 units/mL in 0.2 M Tris-HCl buffer at pH 7.3), sodium pyruvate: 1 mM (from a stock of 20 mM in 0.2 M Tris-HCl buffer at pH 7.3), and choline metabolite/lipid or control: as depicted in Figure S31b. The enzymatic reactions were initiated by adding NADH (37.5 µM-150 µM from a stock of 3 mM in 0.2 M Tris-HCl buffer at pH 7.3) to the cuvette. To generate kinetic plots with varying pyruvate and constant NADH concentrations, the final concentrations of reaction components in the kinetic runs were as follows: LDH: 0.06 U/mL, choline metabolite/lipid or control: as depicted in Figure S31b, and sodium pyruvate: 300 µM-1 mM (from a stock solution of 20 mM in 0.2 M Tris-HCl buffer at pH 7.3). The enzymatic reactions were initiated by adding NADH (150 µM from a stock solution of 3 mM in 0.2 M Tris-HCl buffer at pH 7.3). The velocity of the enzymatic reaction was monitored by recording the rate of decrease in the absorbance of NADH at 340 nm and the $K_m$ and $V_{max}$ values were obtained from the double reciprocal plots that were generated by using the initial velocity of the reactions. The kinetic runs corresponding to each reaction were repeated at least three times.

2. Citrate Synthase (CS): Enzymatic assays on porcine heart CS (Sigma) were performed at 25 °C in a quartz cuvette and by using a UV-Vis spectrophotometer (Perkin Elmer Lambda 25) as per a procedure described previously.[15] The stock solutions of choline, CDP choline, GPC and sodium chloride (20 mM each), acetyl-CoA (1 mM), and phosphocholine and calcium chloride (100 mM each) were made in water. The 0.5 mM stock of 5,5'-dithiobis-(2-nitrobenzoic acid) or DTNB was made in 0.5 M Tris-HCl buffer at pH 8.1 and the 2 mM stock of oxaloacetic acid was made in 0.1 M triethanolamine-HCl buffer at pH 8.0 containing 1 mM EDTA (henceforth known as buffer A), and that of the CS enzyme was made in 0.1 M Tris-HCl buffer at pH 7.0. The stock solutions of PC and SM lipids were made in DMSO at 2.5 mg/mL. All enzyme assays were performed in 0.1 M Tris-HCl

buffer at pH 8.1. To generate kinetic plots with varying acetyl-CoA and constant oxaloacetic acid concentrations, the final concentrations of reaction components in the kinetic runs were as follows: DTNB: 100 µM (from a stock solution of 0.5 mM in 0.5 M Tris-HCl buffer at pH 8.1), oxaloacetic acid: 100 µM (from a stock solution of 2 mM in buffer A), choline metabolite/lipid or control: as depicted in Figure S32b, and acetyl-CoA: 5 µM–75 µM (from a stock solution of 1 mM in water). The enzymatic reactions were initiated by adding CS enzyme (194 ng/mL from a stock of 3.87 µg/mL in 0.1 M Tris -HCl buffer at pH 7.0) to the cuvette. To generate kinetic plots with varying oxaloacetic acid and constant acetyl-CoA concentrations, the final concentrations of reaction components in the kinetic runs were as follows: acetyl-CoA: 50 µM (from a stock solution of 1 mM in water), DTNB: 100 µM (from a stock solution of 0.5 mM in 0.5 M Tris-HCl buffer at pH 8.1), choline metabolite/lipid or control: as depicted in Figure S32b, and oxaloacetic acid: 5 µM–75 µM (from a stock solution of 1 mM in buffer A). The enzymatic reactions were initiated by adding CS enzyme (194 ng/mL from a stock of 3.87 µg/mL in 0.1 M Tris-HCl buffer at pH 7.0) to the cuvette. The velocity of the enzymatic reaction was monitored by recording the rate of increase in the absorbance at 412 nm which was a result of reduction of DTNB by enzymatically liberated CoA-SH.
The $K_m$ and $V_{max}$ values were obtained from the double reciprocal plots that were generated by using the initial velocity of the reactions. The kinetic runs corresponding to each reaction condition were repeated at least three times.

3. Glyceraldehyde-3-phosphate dehydrogenase (GAPDH): Enzymatic assays on rabbit muscle GAPDH (Sigma) were performed at 25 °C in a quartz cuvette by using a UV-Vis spectrophotometer (Perkin Elmer Lambda 25) by making slight modifications to the procedures described previously.[16, 17] The stock solutions of PC and SM lipids (2.5 mg/mL) were made in DMSO, and those of choline, CDP choline, GPC, sodium chloride , phosphocholine, calcium chloride were made in 15 mM sodium pyrophosphate buffer at pH 8.5 containing 30 mM sodium arsenate (henceforth known as buffer B) at 30 mM. Stock solution of DL-glyceraldehyde-3-phosphate (G-3-P; 7.6 mM), and $NAD^+$ (4.5 mM) were also made in buffer B. All enzyme assays were performed in buffer B containing 3.3 mM DTT (added from a stock of 100 mM in buffer B). To generate kinetic plots with varying G-3-P and constant $NAD^+$ concentrations, the final concentrations of reaction components in the kinetic runs were as follows: GAPDH: 0.38 U/mL (added from a stock of 11.4 units/mL in buffer B), $NAD^+$: 150 µM (added from a stock of 4.5 mM in buffer B), and choline metabolite/lipid or control: as depicted in Figure S33b. The enzymatic reactions were initiated by adding G-3-P (63.3 µM-500 µM from a stock of 7.6 mM in buffer B) to the cuvette. To generate kinetic plots with varying $NAD^+$ and

constant G-3-P concentrations, the final concentrations of reaction components in the kinetic runs were as follows: GAPDH: 0.38 U/mL (added from a stock of 11.4 units/mL in buffer B), choline metabolite/lipid or control: as depicted in Figure S33b, and $NAD^+$: 25 µM-150 µM (from a stock solution of 4.5 mM in buffer B). The enzymatic reactions were initiated by adding G-3-P (126.6 µM from a stock solution of 7.6 mM in buffer B). The velocity of the enzymatic reaction was monitored by recording the rate of increase in the absorbance of NADH at 340 nm and the $K_m$ and $V_{max}$ values were obtained from the double reciprocal plots that were generated by using the initial velocity of the reactions. The kinetic runs corresponding to each reaction were repeated at least three times.

4. Malic enzyme (ME): Enzymatic assays on chicken liver ME (Sigma) were performed at 25 °C in a 96-well plate by using a microtiter plate reader (BioTek) as per a procedure described previously.[18] The stock solutions of choline, CDP choline, GPC, and sodium chloride were made in water at 20 mM. The stocks solution for phosphocholine and calcium chloride were made in water at 150 mM, and those of PC and SM lipids were made in DMSO at 2.5 mg/mL. All enzyme assays were performed in 66.7 mM triethanolamine-HCl buffer at pH 7.4. To generate kinetic plots with varying malic acid and constant $NADP^+$ concentrations, the final concentrations of reaction components in the kinetic runs were as follows: manganese (II) chloride: 5 mM (from a stock solution of 0.1 M in water), $NADP^+$: 300 µM (from a stock solution of 6 mM in water), choline metabolite/lipid or control: as depicted in Figure S34b, and malic acid: 100 µM-1 mM (from a stock of 20 mM in water). The enzymatic reactions were initiated by adding ME (0.033 U/mL from a stock solution of 1 U/mL in 50 mM Tris-HCl buffer at pH 7.5) to the 96-well plate. To generate kinetic plots with varying $NADP^+$ and constant malic acid concentrations, the final concentrations of reaction components in the kinetic runs were as follows: manganese (II) chloride: 5 mM (from a stock solution of 0.1 M in water), malic acid: 800 µM (from a stock of 20 mM in water), choline metabolite/lipid or control: as depicted in Figure S34b, and $NADP^+$: 5 µM-500 µM (from a stock solution of 6 mM in water). The enzymatic reactions were initiated by adding ME (0.033 U/mL from a stock solution of 1 U/mL in 50 mM Tris-HCl buffer at pH 7.5) to the 96-well plate. The velocity of the enzymatic reaction was monitored by recording the rate of increase in the absorbance of NADPH at 340 nm and the $K_m$ and $V_{max}$ values were obtained from the double reciprocal plots that were generated by using the initial velocity of the reactions. The kinetic runs corresponding to each reaction were repeated at least three times.

5. Glutamate dehydrogenase (GLUD1): Enzymatic assays on bovine liver GLUD1 (Sigma) were performed at 25 °C in a quartz cuvette by using a UV-Vis spectrophotometer (Perkin Elmer Lambda 25) as per the procedure described previously.[19] The stock solutions of choline, CDP choline, phosphocholine, GPC, sodium chloride and calcium chloride were made in water at 30 mM, and those of PC and SM lipids were made in DMSO at 2.5 mg/mL. All enzyme assays were performed in 100 mM potassium phosphate buffer at pH 7.0. To generate kinetic plots with varying α-ketoglutarate (α-KG) and constant NADH concentrations, the final concentrations of reaction components in the kinetic runs were as follows: ammonium chloride: 50 mM (from a stock solution of 1.5 M in water), NADH: 150 µM (from a stock solution of 5 mM in water), choline metabolite/lipid or control: as depicted in Figure S35b, and α-KG: 100 µM-3 mM (from a stock of 90 mM in water). The enzymatic reactions were initiated by adding GLUD1 (0.16 U/mL from a stock solution of 24 U/mL in 100 mM potassium phosphate buffer at pH 7.0) to the cuvette. To generate kinetic plots with varying NADH and constant α-KG concentrations, the final concentrations of reaction components in the kinetic runs were as follows: ammonium chloride: 50 mM (from a stock solution of 1.5 M in water), α-KG: 250 µM (from a stock solution of 7.5 mM in water), choline metabolite/lipid or control: as depicted in Figure S35b, and NADH: 50 µM-100 µM (from a stock of 5 mM in water). The enzymatic reactions were initiated by adding GLUD1 (0.16 U/mL from a stock solution of 24 U/mL in 100 mM potassium phosphate buffer at pH 7.0) to the cuvette. The velocity of the enzymatic reaction was monitored by recording the rate of decrease in the absorbance of NADH at 340 nm and the $K_m$ and $V_{max}$ values were obtained from the double reciprocal plots that were generated by using the initial velocity of the reactions. The kinetic runs corresponding to each reaction were repeated at least three times.

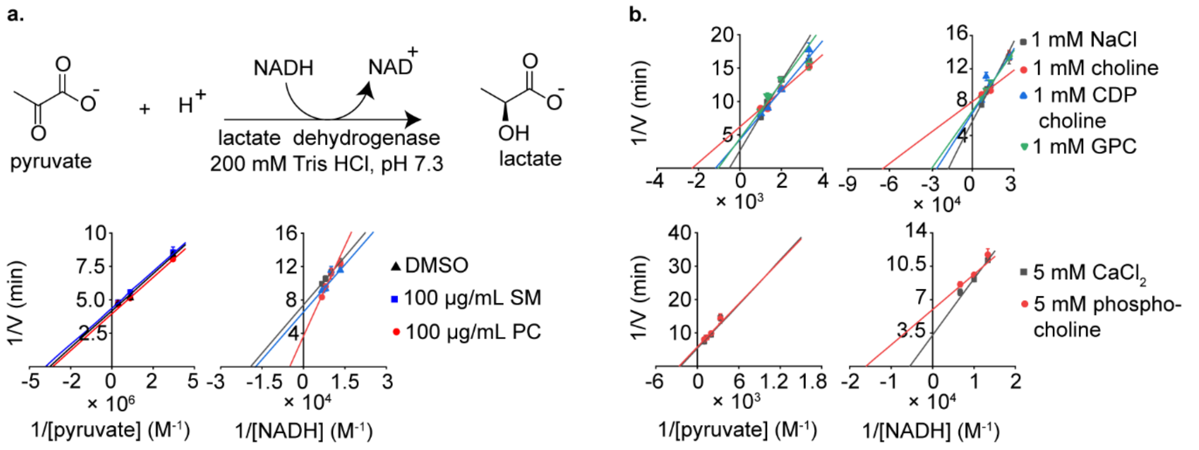


c. Variable pyruvate, constant NADH

| Metabolite/Control | $K_m$ app (M) | $K_m$ metabolite / $K_m$ control | $V_{max}$ (min$^{-1}$) | $V_{max}$ metabolite / $V_{max}$ control |
|---|---|---|---|---|
| DMSO (control) | $(2.95 \pm 0.4) \times 10^{-7}$ | 1 | 0.24 ± 0.01 | 1 |
| PC | $(3.09 \pm 0.3) \times 10^{-7}$ | 1.05 ± 0.2 | 0.23 ± 0.01 | 0.95 ± 0.1 |
| SM | $(2.76 \pm 0.3) \times 10^{-7}$ | 0.94 ± 0.2 | 0.25 ± 0.01 | 1.06 ± 0.1 |
| 1 mM NaCl (control) | $(1.93 \pm 0.5) \times 10^{-3}$ | 1 | 0.38 ± 0.1 | 1 |
| 1 mM choline | $(4.34 \pm 0.6) \times 10^{-4}$ | 0.22 ± 0.1 | 0.16 ± 0.01 | 0.43 ± 0.2 |
| 1 mM CDP choline | $(8.51 \pm 1.1) \times 10^{-4}$ | 0.44 ± 0.1 | 0.23 ± 0.03 | 0.61 ± 0.2 |
| 1 mM GPC | $(9.60 \pm 1.8) \times 10^{-4}$ | 0.50 ± 0.2 | 0.23 ± 0.05 | 0.60 ± 0.3 |
| 5 mM $CaCl_2$ (control) | $(4.09 \pm 1.2) \times 10^{-4}$ | 1 | 0.19 ± 0.03 | 1 |
| 5 mM phosphocholine | $(3.77 \pm 0.6) \times 10^{-4}$ | 0.92 ± 0.3 | 0.17 ± 0.01 | 0.94 ± 0.2 |

d. Variable NADH, constant pyruvate

| Metabolite/Control | $K_m$ app (M) | $K_m$ metabolite / $K_m$ control | $V_{max}$ (min$^{-1}$) | $V_{max}$ metabolite / $V_{max}$ control |
|---|---|---|---|---|
| DMSO (control) | $(5.22 \pm 0.4) \times 10^{-5}$ | 1 | 0.14 ± 0.00 | 1 |
| PC | $(1.95 \pm 0.5) \times 10^{-4}$ | 3.73 ± 0.9 | 0.27 ± 0.07 | 2.01 ± 0.5 |
| SM | $(5.79 \pm 1.1) \times 10^{-5}$ | 1.11 ± 0.2 | 0.15 ± 0.02 | 1.13 ± 0.1 |
| 1 mM NaCl (control) | $(5.75 \pm 0.7) \times 10^{-5}$ | 1 | 0.18 ± 0.01 | 1 |
| 1 mM choline | $(1.54 \pm 0.7) \times 10^{-5}$ | 0.27 ± 0.1 | 0.12 ± 0.01 | 0.69 ± 0.1 |
| 1 mM CDP choline | $(3.85 \pm 0.7) \times 10^{-5}$ | 0.67 ± 0.1 | 0.15 ± 0.01 | 0.84 ± 0.1 |
| 1 mM GPC | $(3.37 \pm 0.1) \times 10^{-5}$ | 0.59 ± 0.1 | 0.14 ± 0.00 | 0.80 ± 0.1 |
| 5 mM $CaCl_2$ (control) | $(1.82 \pm 0.2) \times 10^{-4}$ | 1 | 0.31 ± 0.05 | 1 |
| 5 mM phosphocholine | $(6.22 \pm 2.3) \times 10^{-5}$ | 0.34 ± 0.1 | 0.17 ± 0.03 | 0.54 ± 0.1 |

**Figure S31.** Characterizing the effect of choline metabolites on the activity of lactate dehydrogenase. a) The enzymatic reaction of lactate dehydrogenase. b) Double reciprocal plots generated by calculating the initial velocity of each enzyme-catalyzed reaction in the presence of the designated choline metabolites or in the presence of salt/vehicle solvent instead of the metabolites. Specifically, whereas assays performed with choline, CDP choline and GPC contained 1 mM concentrations of these metabolites (used in their chloride salt forms), in the corresponding metabolite-devoid experiments, these metabolites were replaced with 1 mM NaCl. Assays with phosphocholine were performed in the presence of 5 mM phosphocholine (used as phosphocholine chloride calcium salt), whereas the corresponding metabolite-devoid experiments contained 5 mM $CaCl_2$. PC and SM were introduced into the assays in their DMSO-solubilized forms and the corresponding negative controls contained an identical volume of the DMSO vehicle. c) and d) Table depicting the value of $K_m$ and $V_{max}$ of the enzymatic reactions.

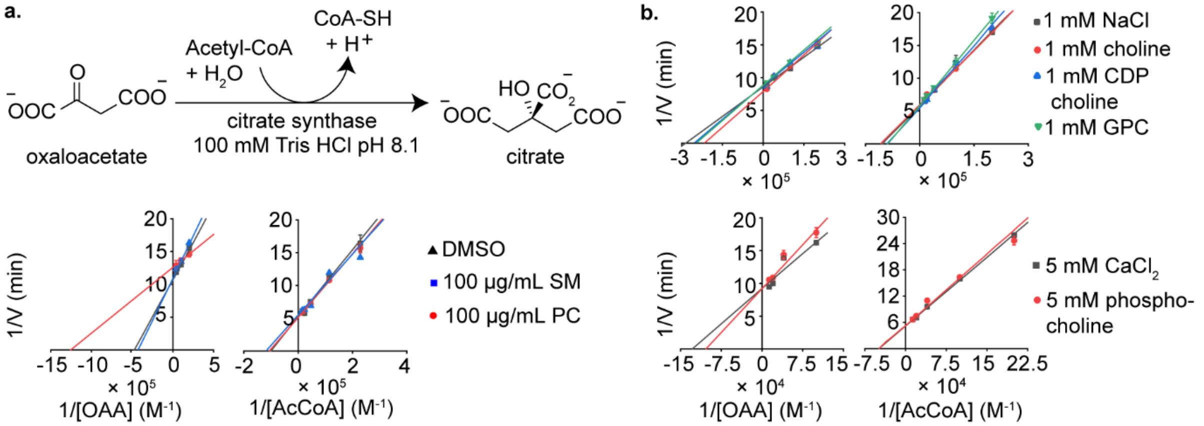


c.

Variable AcCoA, constant oxaloacetate

| Metabolite/Control | $K_m$ app (M) | $K_m$ metabolite / $K_m$ control | $V_{max}$ (min$^{-1}$) | $V_{max}$ metabolite / $V_{max}$ control |
|---|---|---|---|---|
| DMSO (control) | $(1.14 \pm 0.1) \times 10^{-5}$ | 1 | 0.19 ± 0.01 | 1 |
| PC | $(1.07 \pm 0.03) \times 10^{-5}$ | 0.94 ± 0.1 | 0.19 ± 0.002 | 1.00 ± 0.05 |
| SM | $(9.60 \pm 2.1) \times 10^{-6}$ | 0.84 ± 0.2 | 0.18 ± 0.03 | 0.94 ± 0.16 |
| 1 mM NaCl (control) | $(9.84 \pm 0.1) \times 10^{-6}$ | 1 | 0.17 ± 0.001 | 1 |
| 1 mM choline | $(9.25 \pm 0.3) \times 10^{-6}$ | 0.94 ± 0.04 | 0.17 ± 0.003 | 0.95 ± 0.02 |
| 1 mM CDP choline | $(1.09 \pm 0.04) \times 10^{-5}$ | 1.11 ± 0.1 | 0.18 ± 0.003 | 1.01 ± 0.02 |
| 1 mM GPC | $(1.12 \pm 0.04) \times 10^{-5}$ | 1.14 ± 0.04 | 0.17 ± 0.001 | 0.97 ± 0.01 |
| 5 mM $CaCl_2$ (control) | $(2.06 \pm 0.005) \times 10^{-5}$ | 1 | 0.20 ± 0.004 | 1 |
| 5 mM phosphocholine | $(2.05 \pm 0.09) \times 10^{-5}$ | 1.00 ± 0.1 | 0.19 ± 0.01 | 0.96 ± 0.06 |

d.

Variable oxaloacetate, constant AcCoA

| Metabolite/Control | $K_m$ app (M) | $K_m$ metabolite / $K_m$ control | $V_{max}$ (min$^{-1}$) | $V_{max}$ metabolite / $V_{max}$ control |
|---|---|---|---|---|
| DMSO (control) | $(2.11 \pm 0.2) \times 10^{-6}$ | 1 | 0.093 ± 0.002 | 1 |
| PC | $(7.97 \pm 0.7) \times 10^{-7}$ | 0.38 ± 0.1 | 0.079 ± 0.001 | 0.86 ± 0.02 |
| SM | $(2.34 \pm 0.4) \times 10^{-6}$ | 1.11 ± 0.2 | 0.092 ± 0.004 | 0.99 ± 0.1 |
| 1 mM NaCl (control) | $(3.20 \pm 0.2) \times 10^{-6}$ | 1 | 0.116 ± 0.003 | 1 |
| 1 mM choline | $(4.62 \pm 0.4) \times 10^{-6}$ | 1.32 ± 0.2 | 0.124 ± 0.01 | 1.07 ± 0.1 |
| 1 mM CDP choline | $(3.95 \pm 0.6) \times 10^{-6}$ | 1.13 ± 0.2 | 0.115 ± 0.01 | 0.99 ± 0.1 |
| 1 mM GPC | $(4.05 \pm 0.2) \times 10^{-6}$ | 1.16 ± 0.1 | 0.114 ± 0.002 | 0.98 ± 0.03 |
| 5 mM $CaCl_2$ (control) | $(7.79 \pm 1.4) \times 10^{-6}$ | 1 | 0.109 ± 0.01 | 1 |
| 5 mM phosphocholine | $(9.68 \pm 1.9) \times 10^{-6}$ | 1.24 ± 0.3 | 0.109 ± 0.01 | 0.99 ± 0.1 |

**Figure S32.** Characterizing the activity of citrate synthase in presence of choline metabolites and lipids. a) The enzymatic reaction of citrate synthase. b) Double reciprocal plots generated by calculating the initial velocity of each enzyme-catalyzed reaction in the presence of the designated choline metabolites or in the presence of salt/vehicle solvent instead of the metabolites. Specifically, whereas assays performed with choline, CDP choline and GPC contained 1 mM concentrations of these metabolites, in the corresponding metabolite-devoid experiments, these metabolites were replaced with 1 mM NaCl. Assays with phosphocholine were performed in the presence of 5 mM phosphocholine (used as phosphocholine chloride calcium salt), whereas the corresponding metabolite-devoid experiments contained 5 mM $CaCl_2$. PC and SM were introduced into the assays in their DMSO-solubilized forms and the corresponding negative controls contained an identical volume of the DMSO vehicle. c) and d) Table depicting the value of $K_m$ and $V_{max}$ of the enzymatic reactions.

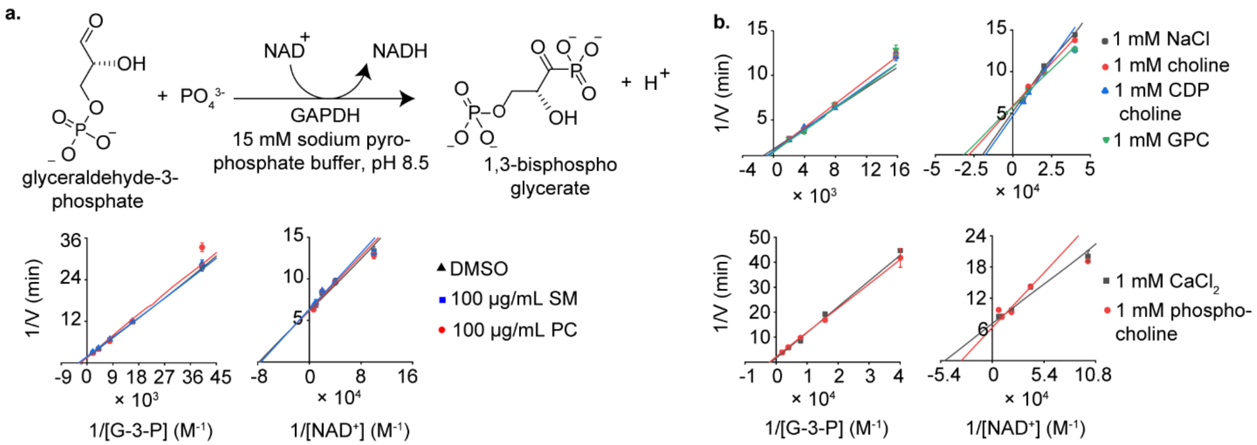


c.

Variable glyceraldehyde-3-phosphate, constant NAD+ (control)

| Metabolite/Control | $K_m$ app (M) | $K_m$ metabolite / $K_m$ control | $V_{max}$ ($min^{-1}$) | $V_{max}$ metabolite / $V_{max}$ control |
|---|---|---|---|---|
| DMSO (control) | $(4.19 \pm 0.2) \times 10^{-4}$ | 1 | 0.64 ± 0.01 | 1 |
| PC | $(4.69 \pm 0.5) \times 10^{-4}$ | 1.12 ± 0.11 | 0.70 ± 0.09 | 1.08 ± 0.15 |
| SM | $(3.41 \pm 0.1) \times 10^{-4}$ | 0.81 ± 0.02 | 0.54 ± 0.02 | 0.84 ± 0.03 |
| 1 mM NaCl (control) | $(3.29 \pm 0.4) \times 10^{-4}$ | 1 | 0.58 ± 0.07 | 1 |
| 1 mM choline | $(4.35 \pm 0.1) \times 10^{-4}$ | 1.32 ± 0.17 | 0.65 ± 0.05 | 1.14 ± 0.15 |
| 1 mM CDP choline | $(4.05 \pm 0.2) \times 10^{-4}$ | 1.23 ± 0.16 | 0.66 ± 0.05 | 1.15 ± 0.16 |
| 1 mM GPC | $(4.75 \pm 1.2) \times 10^{-4}$ | 1.44 ± 0.41 | 0.76 ± 0.23 | 1.32 ± 0.43 |
| 1 mM $CaCl_2$(control) | $(6.02 \pm 0.5) \times 10^{-4}$ | 1 | 0.58 ± 0.08 | 1 |
| 1 mM phosphocholine | $(5.17 \pm 0.2) \times 10^{-4}$ | 0.86 ± 0.07 | 0.53 ± 0.02 | 0.90 ± 0.13 |

d.

Variable NAD+, constant glyceraldehyde-3-phosphate

| Metabolite/Control | $K_m$ app (M) | $K_m$ metabolite / $K_m$ control | $V_{max}$ ($min^{-1}$) | $V_{max}$ metabolite / $V_{max}$ control |
|---|---|---|---|---|
| DMSO (control) | $(1.83 \pm 0.3) \times 10^{-5}$ | 1 | 0.17 ± 0.01 | 1 |
| PC | $(1.60 \pm 0.2) \times 10^{-5}$ | 0.87 ± 0.15 | 0.17 ± 0.01 | 1.00 ± 0.06 |
| SM | $(1.86 \pm 0.3) \times 10^{-5}$ | 1.02 ± 0.22 | 0.16 ± 0.01 | 0.97 ± 0.05 |
| 1 mM NaCl (control) | $(4.99 \pm 0.9) \times 10^{-5}$ | 1 | 0.19 ± 0.01 | 1 |
| 1 mM choline | $(3.53 \pm 0.6) \times 10^{-5}$ | 0.71 ± 0.18 | 0.17 ± 0.01 | 0.91 ± 0.16 |
| 1 mM CDP choline | $(5.63 \pm 0.8) \times 10^{-5}$ | 1.13 ± 0.27 | 0.21 ± 0.02 | 1.12 ± 0.20 |
| 1 mM GPC | $(3.14 \pm 0.4) \times 10^{-5}$ | 0.63 ± 0.14 | 0.17 ± 0.01 | 0.91 ± 0.16 |
| 1 mM $CaCl_2$ (control) | $(2.02 \pm 0.3) \times 10^{-5}$ | 1 | 0.14 ± 0.01 | 1 |
| 1 mM phosphocholine | $(3.13 \pm 0.4) \times 10^{-5}$ | 1.55 ± 0.32 | 0.16 ± 0.02 | 1.12 ± 0.16 |

**Figure S33.** Characterizing the activity of GAPDH in presence of choline metabolites and lipids. a) The enzymatic reaction of GAPDH. b) Double reciprocal plots generated by calculating the initial velocity of each enzyme-catalyzed reaction in the presence of the designated choline metabolites or in the presence of salt/vehicle solvent instead of the metabolites. Specifically, whereas assays performed with choline, CDP choline and GPC contained 1 mM concentrations of these metabolites, in the corresponding metabolite-devoid experiments, these metabolites were replaced with 1 mM NaCl. Assays with phosphocholine were also performed in the presence of 1 mM phosphocholine (used as phosphocholine chloride calcium salt), whereas the corresponding metabolite-devoid experiments contained 1 mM $CaCl_2$. PC and SM were introduced into the assays in their DMSO-solubilized forms and the corresponding negative controls contained an identical volume of the DMSO vehicle. c) and d) Table depicting the value of $K_m$ and $V_{max}$ of the enzymatic reactions.

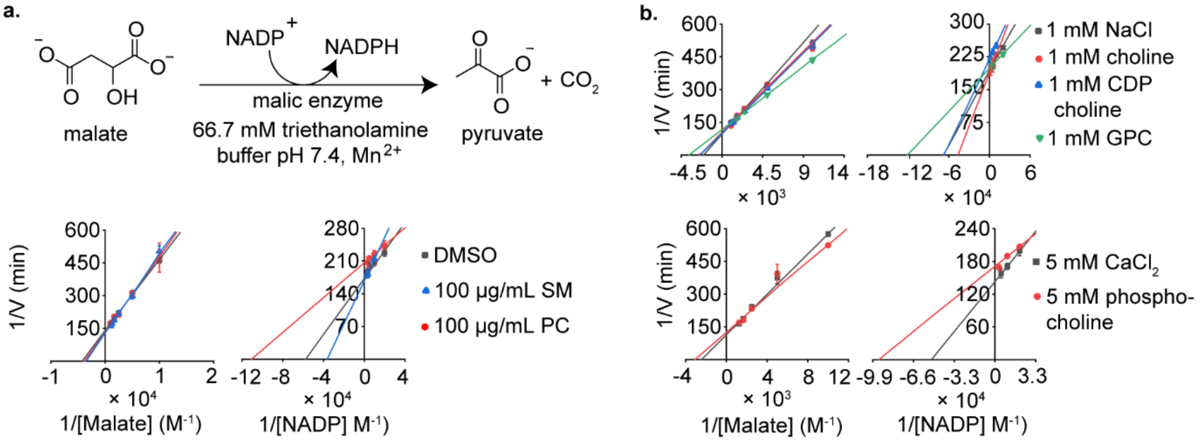


c. Variable malate, constant NADP+

| Metabolite/Control | $K_m$ app (M) | $K_m$ metabolite / $K_m$control | $V_{max}$ (min$^{-1}$) | $V_{max}$ metabolite / $V_{max}$ control |
|---|---|---|---|---|
| DMSO (control) | (2.38 ± 0.05) × $10^{-4}$ | 1 | (7.25 ± 0.09) × $10^{-3}$ | 1 |
| PC | (2.62 ± 0.2) × $10^{-4}$ | 1.10 ± 0.08 | (7.53 ± 0.3) × $10^{-3}$ | 1.04 ± 0.04 |
| SM | (2.89 ± 0.09) × $10^{-4}$ | 1.22 ± 0.05 | (7.95 ± 0.2) × $10^{-3}$ | 1.10 ± 0.03 |
| 1 mM NaCl (control) | (4.92 ± 0.2) × $10^{-4}$ | 1 | (1.05 ± 0.02) × $10^{-2}$ | 1 |
| 1 mM choline | (3.94 ± 0.3) × $10^{-4}$ | 0.80 ± 0.06 | (9.54 ± 0.7) × $10^{-3}$ | 0.91 ± 0.07 |
| 1 mM CDP choline | (4.10 ± 0.07) × $10^{-4}$ | 0.83 ± 0.03 | (1.00 ± 0.02) × $10^{-2}$ | 0.95 ± 0.04 |
| 1 mM GPC | (2.74 ± 0.06) × $10^{-4}$ | 0.56 ± 0.02 | (8.59 ± 0.2) × $10^{-3}$ | 0.82 ± 0.03 |
| 5 mM $CaCl_2$(control) | (4.11 ± 0.1) × $10^{-4}$ | 1 | (8.88 ± 0.5) × $10^{-3}$ | 1 |
| 5 mM phosphocholine | (3.24 ± 0.3) × $10^{-4}$ | 0.79 ± 0.07 | (7.94 ± 0.6) × $10^{-3}$ | 0.89 ± 0.08 |

d. Variable NADP+, constant malate

| Metabolite/Control | $K_m$ app (M) | $K_m$ metabolite / $K_m$control | $V_{max}$ (min$^{-1}$) | $V_{max}$ metabolite / $V_{max}$ control |
|---|---|---|---|---|
| DMSO (control) | (1.74 ± 0.3) × $10^{-5}$ | 1 | (5.81 ± 0.08) × $10^{-3}$ | 1 |
| PC | (9.02 ± 1.3) × $10^{-6}$ | 0.52 ± 0.1 | (4.85 ± 0.05) × $10^{-3}$ | 0.83 ± 0.01 |
| SM | (2.76 ± 0.02) × $10^{-5}$ | 1.59 ± 0.23 | (5.95 ± 0.01) × $10^{-3}$ | 1.02 ± 0.01 |
| 1 mM NaCl (control) | (1.48 ± 0.2) × $10^{-5}$ | 1 | (5.22 ± 0.1) × $10^{-3}$ | 1 |
| 1 mM choline | (2.15 ± 0.1) × $10^{-5}$ | 1.46 ± 0.21 | (5.30 ± 0.05) × $10^{-3}$ | 1.02 ± 0.02 |
| 1 mM CDP choline | (1.49 ± 0.3) × $10^{-5}$ | 1.01 ± 0.22 | (4.58 ± 0.09) × $10^{-3}$ | 0.88 ± 0.02 |
| 1 mM GPC | (8.28 ± 0.6) × $10^{-6}$ | 0.56 ± 0.08 | (5.05 ± 0.04) × $10^{-3}$ | 0.97 ± 0.02 |
| 5 mM $CaCl_2$ (control) | (1.95 ± 0.06) × $10^{-5}$ | 1 | (6.98 ± 0.05) × $10^{-3}$ | 1 |
| 5 mM phosphocholine | (1.06 ± 0.2) × $10^{-5}$ | 0.55 ± 0.09 | (5.85 ± 0.2) × $10^{-3}$ | 0.84 ± 0.02 |

**Figure S34.** Characterizing the activity of malic enzyme in presence of choline metabolites and lipids. a) The enzymatic reaction of malic enzyme. b) Double reciprocal plots generated by calculating the initial velocity of each enzyme-catalyzed reaction in the presence of the designated choline metabolites or in the presence of salt/vehicle solvent instead of the metabolites. Specifically, whereas assays performed with choline, CDP choline and GPC contained 1 mM concentrations of these metabolites, in the corresponding metabolite-devoid experiments, these metabolites were replaced with 1 mM NaCl. Assays with phosphocholine were performed in the presence of 5 mM phosphocholine (used as phosphocholine chloride calcium salt), whereas the corresponding metabolite-devoid experiments contained 5 mM $CaCl_2$. PC and SM were introduced into the assays in their DMSO-solubilized forms and the corresponding negative controls contained an identical volume of the DMSO vehicle. c) and d) Table depicting the value of $K_m$ and $V_{max}$ of the enzymatic reactions.

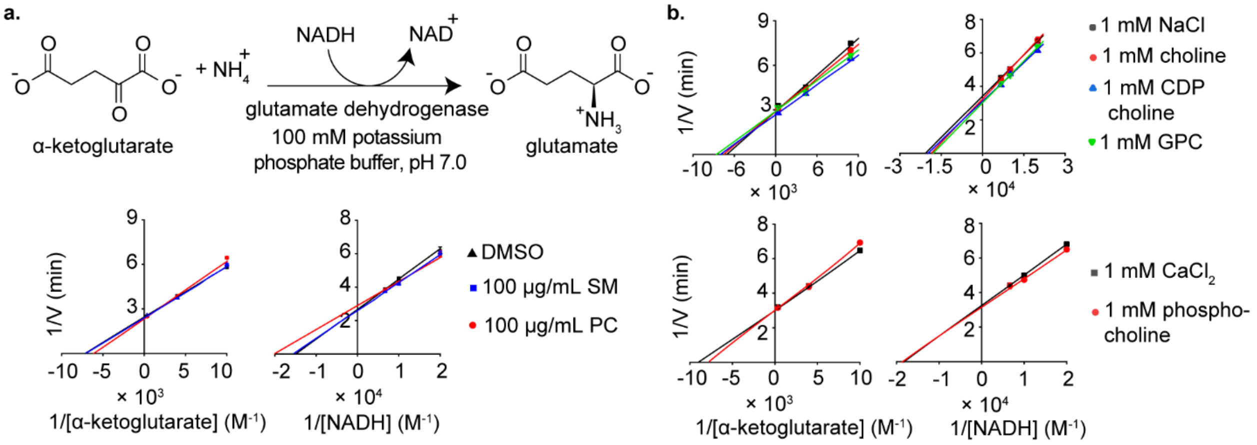


**c.** Variable α-ketoglutarate, constant NADH

| Metabolite/Control | $K_m$ app (M) | $K_m$ metabolite / $K_m$ control | $V_{max}$ (min$^{-1}$) | $V_{max}$ metabolite / $V_{max}$ control |
|---|---|---|---|---|
| DMSO (control) | (1.40 ± 0.1) × $10^{-4}$ | 1 | 0.41 ± 0.01 | 1 |
| PC | (1.63 ± 0.1) × $10^{-4}$ | 1.17 ± 0.1 | 0.42 ± 0.01 | 1.05 ± 0.1 |
| SM | (1.41 ± 0.1) × $10^{-4}$ | 1.01 ± 0.1 | 0.41 ± 0.01 | 1.01 ± 0.1 |
| 1 mM NaCl (control) | (1.54 ± 0.1) × $10^{-4}$ | 1 | 0.35 ± 0.03 | 1 |
| 1 mM choline | (1.45 ± 0.1) × $10^{-4}$ | 0.94 ± 0.1 | 0.35 ± 0.01 | 1.01 ± 0.1 |
| 1 mM CDP choline | (1.36 ± 0.1) × $10^{-4}$ | 0.89 ± 0.1 | 0.37 ± 0.01 | 1.09 ± 0.1 |
| 1 mM GPC | (1.27 ± 0.1) × $10^{-4}$ | 0.82 ± 0.1 | 0.34 ± 0.02 | 0.99 ± 0.1 |
| 1 mM $CaCl_2$ (control) | (1.11 ± 0.01) × $10^{-4}$ | 1 | 0.33 ± 0.01 | 1 |
| 1 mM phosphocholine | (1.28 ± 0.1) × $10^{-4}$ | 1.15 ± 0.04 | 0.33 ± 0.01 | 1.01 ± 0.1 |

**d.** Variable NADH, constant α-ketoglutarate

| Metabolite/Control | $K_m$ app (M) | $K_m$ metabolite / $K_m$ control | $V_{max}$ (min$^{-1}$) | $V_{max}$ metabolite / $V_{max}$ control |
|---|---|---|---|---|
| DMSO (control) | (6.70 ± 0.5 × $10^{-5}$ | 1 | 0.37 ± 0.01 | 1 |
| PC | (4.95 ± 0.7) × $10^{-5}$ | 0.74 ± 0.11 | 0.35 ± 0.02 | 0.93 ± 0.1 |
| SM | (6.43 ± 0.7) × $10^{-5}$ | 0.96 ± 0.13 | 0.39 ± 0.03 | 1.04 ± 0.1 |
| 1 mM NaCl (control) | (4.83 ± 0.1) × $10^{-5}$ | 1 | 0.29 ± 0.01 | 1 |
| 1 mM choline | (5.39 ± 0.1) × $10^{-5}$ | 1.12 ± 0.02 | 0.31 ± 0.01 | 1.05 ± 0.1 |
| 1 mM CDP choline | (5.06 ± 0.5) × $10^{-5}$ | 1.05 ± 0.11 | 0.32 ± 0.02 | 1.11 ± 0.1 |
| 1 mM GPC | (5.56 ± 0.2) × $10^{-5}$ | 1.15 ± 0.05 | 0.33 ± 0.01 | 1.13 ± 0.1 |
| 1 mM $CaCl_2$ (control) | (5.49 ± 0.2) × $10^{-5}$ | 1 | 0.31 ± 0.01 | 1 |
| 1 mM phosphocholine | (5.38 ± 0.5) × $10^{-5}$ | 0.98 ± 0.09 | 0.32 ± 0.02 | 1.04 ± 0.1 |

**Figure S35.** Characterizing the activity of glutamate dehydrogenase in presence of choline metabolites and lipids. a) The enzymatic reaction of glutamate dehydrogenase. b) Double reciprocal plots generated by calculating the initial velocity of each enzyme-catalyzed reaction in the presence of the designated choline metabolites or in the presence of salt/vehicle solvent instead of the metabolites. Specifically, whereas assays performed with choline, CDP choline and GPC contained 1 mM concentrations of these metabolites, in the corresponding metabolite-devoid experiments, these metabolites were replaced with 1 mM NaCl. Assays with phosphocholine were performed in the presence of 1 mM phosphocholine (used as phosphocholine chloride calcium salt), whereas the corresponding metabolite-devoid experiments contained 1 mM $CaCl_2$. PC and SM were introduced into the assays in their DMSO-solubilized forms and the corresponding negative controls contained an identical volume of the DMSO vehicle. c) and d) Table depicting the value of $K_m$ and $V_{max}$ of the enzymatic reactions.

### 20. $^{1}H$ and $^{13}C$ NMR spectra

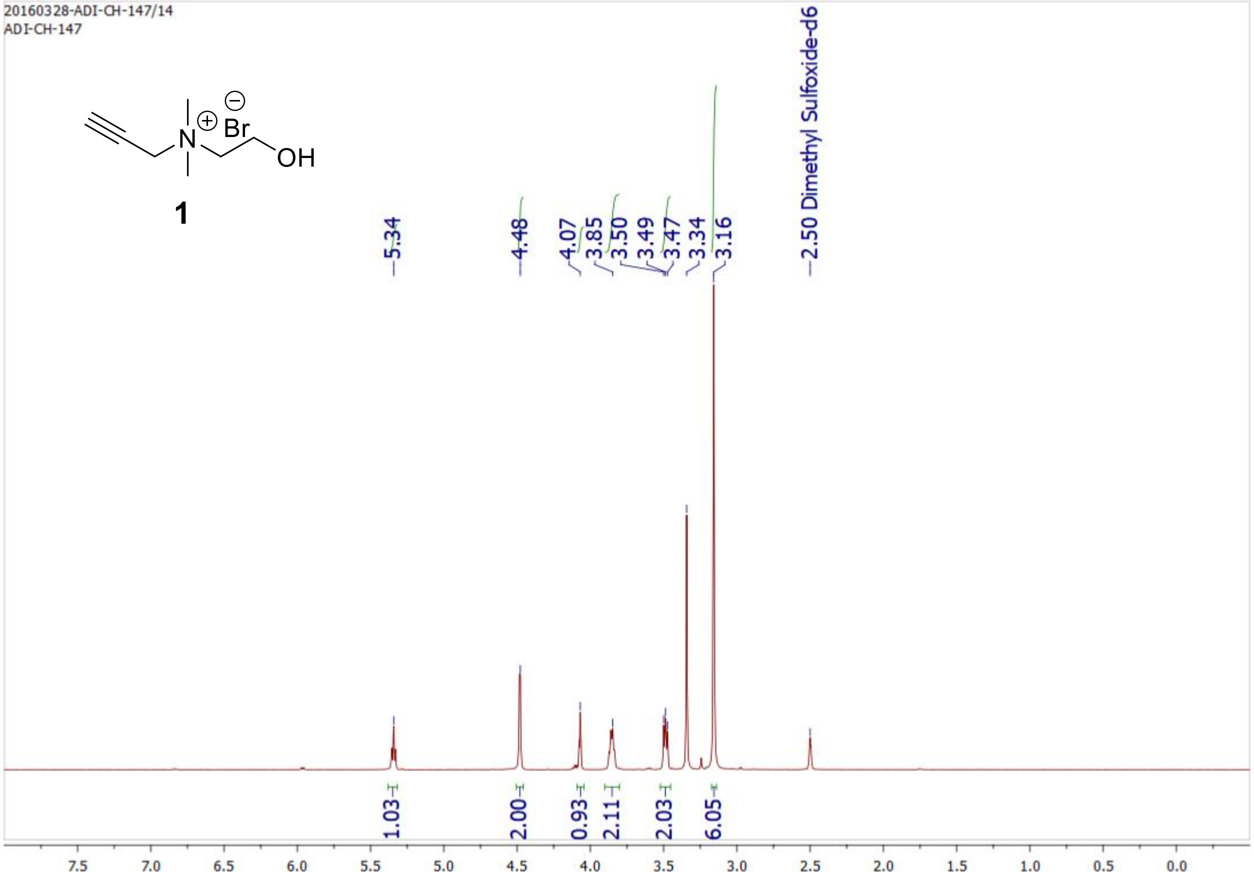


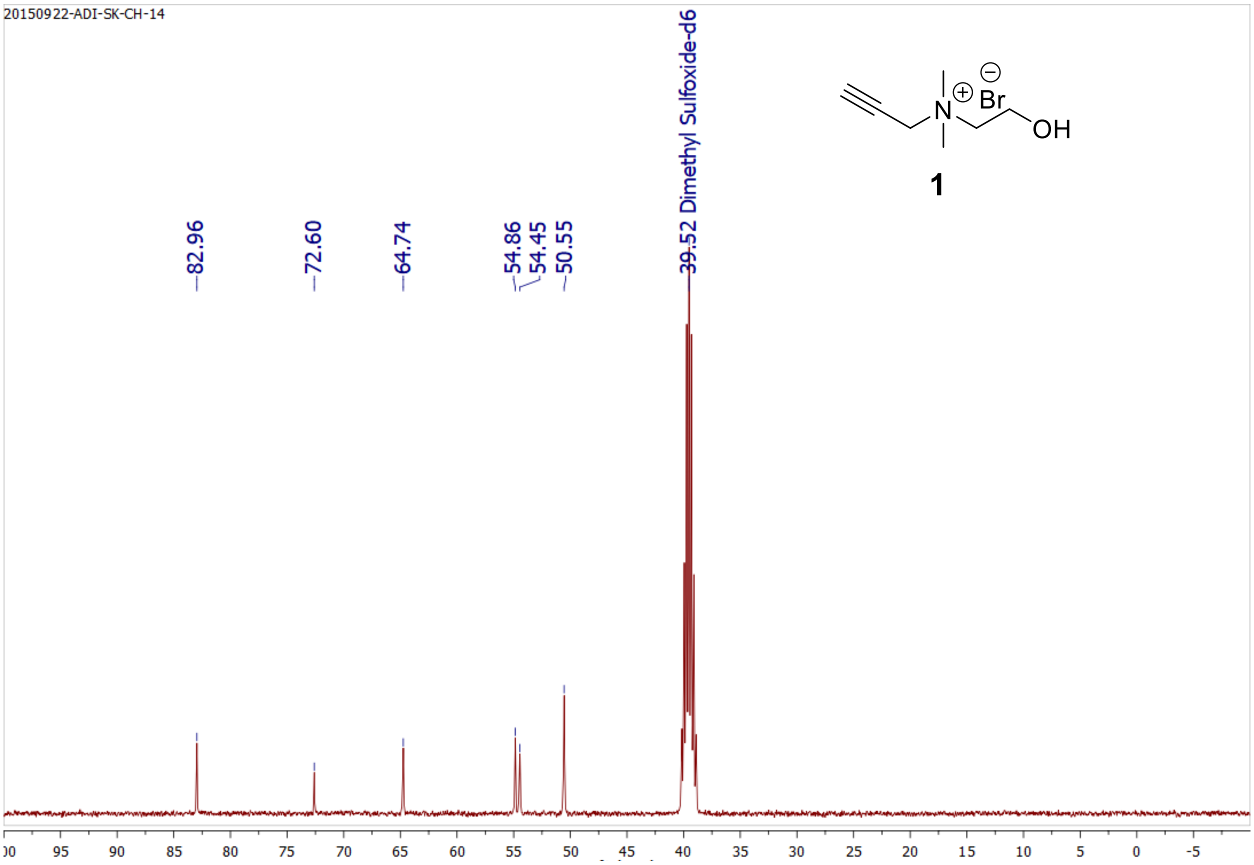

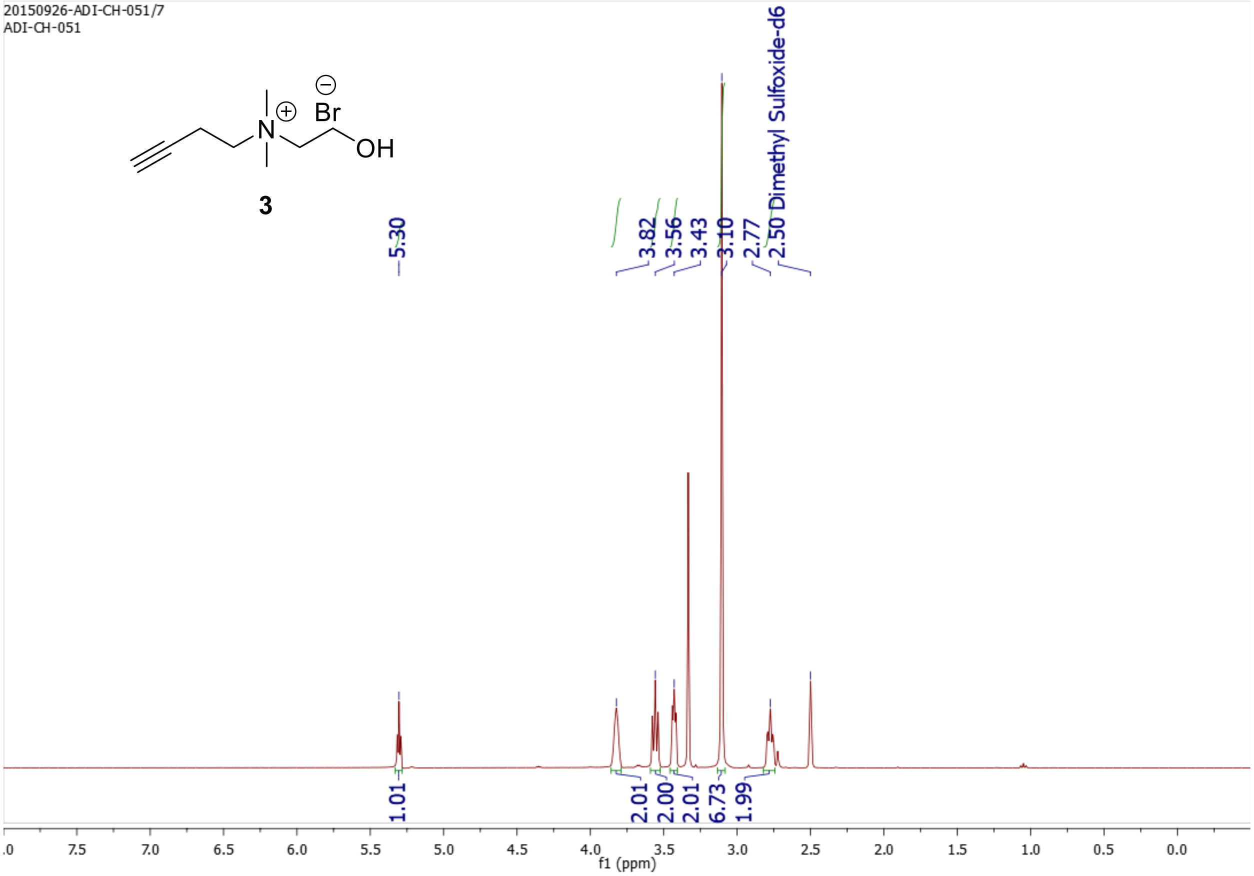
20150926-ADI-CH-051/7
ADI-CH-051
3
5.30
3.82
3.56
3.43
3.10
2.77
2.50 Dimethyl Sulfoxide-d6
1.01
2.01
2.00
2.01
6.73
1.99
f1 (ppm)

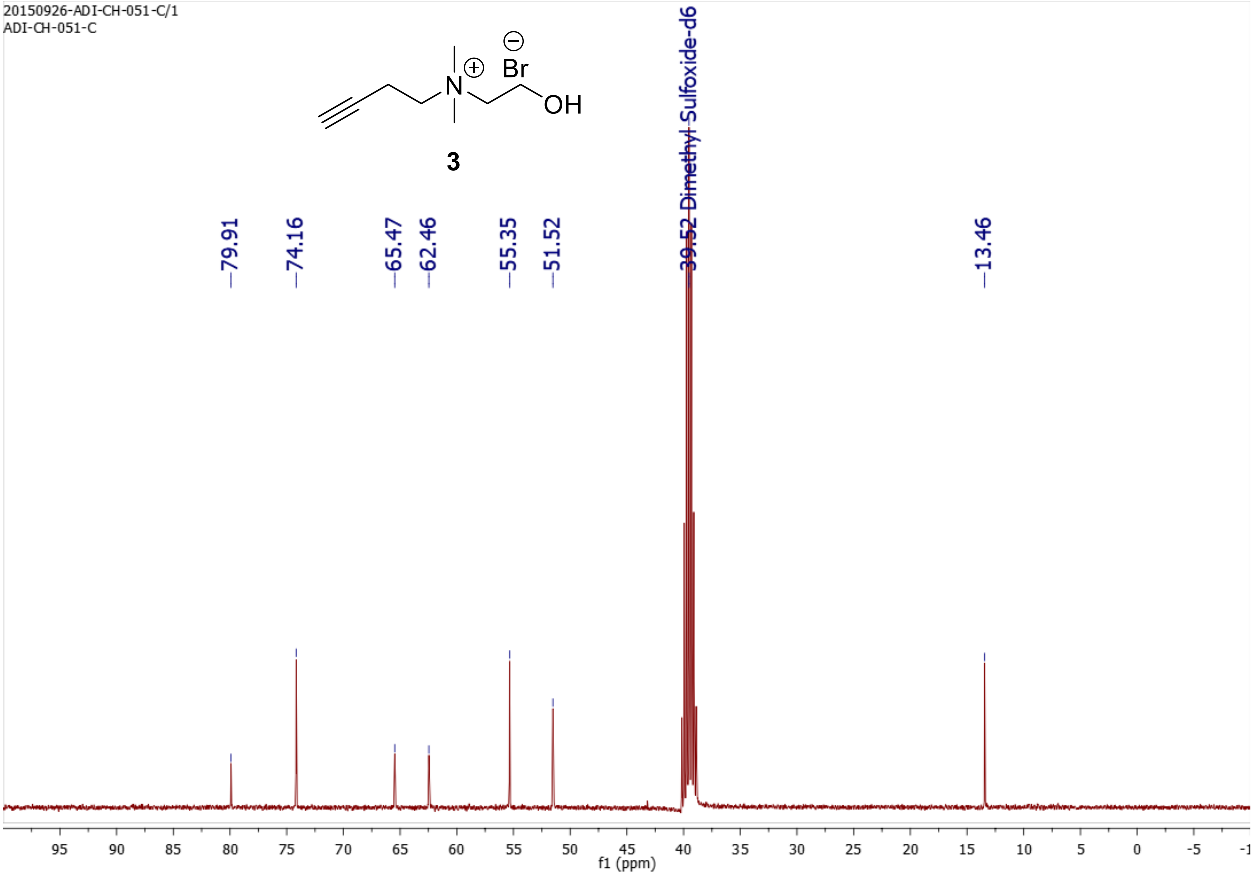
20150926-ADI-CH-051-C/1
ADI-CH-051-C
3
79.91
74.16
65.47
62.46
55.35
51.52
39.52 Dimethyl Sulfoxide-d6
13.46
f1 (ppm)

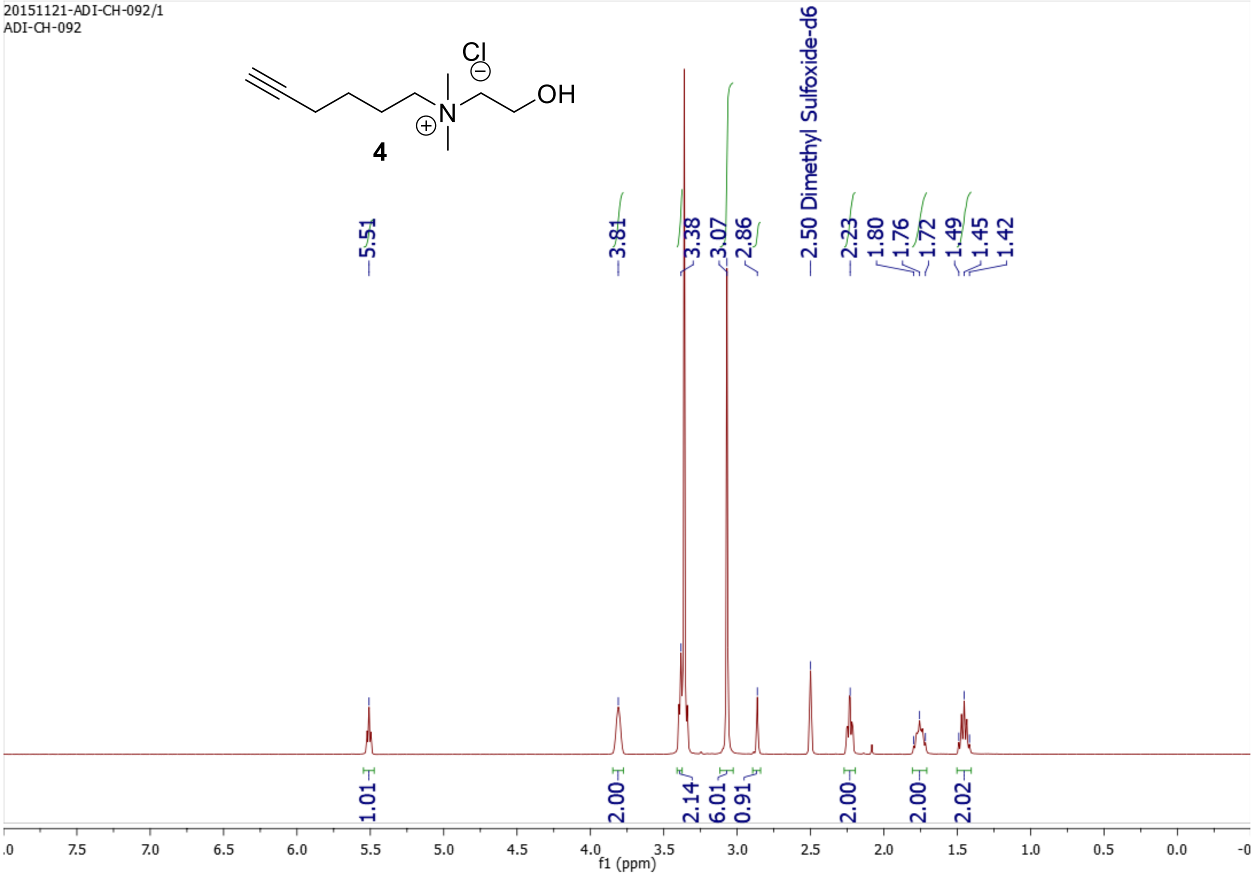
20151121-ADI-CH-092/1
ADI-CH-092
Cl
OH
N
4
5.51
3.81
3.38
3.07
2.86
2.50 Dimethyl Sulfoxide-d6
2.23
1.80
1.76
1.72
1.49
1.45
1.42
1.01
2.00
2.14
6.01
0.91
2.00
2.00
2.02
f1 (ppm)

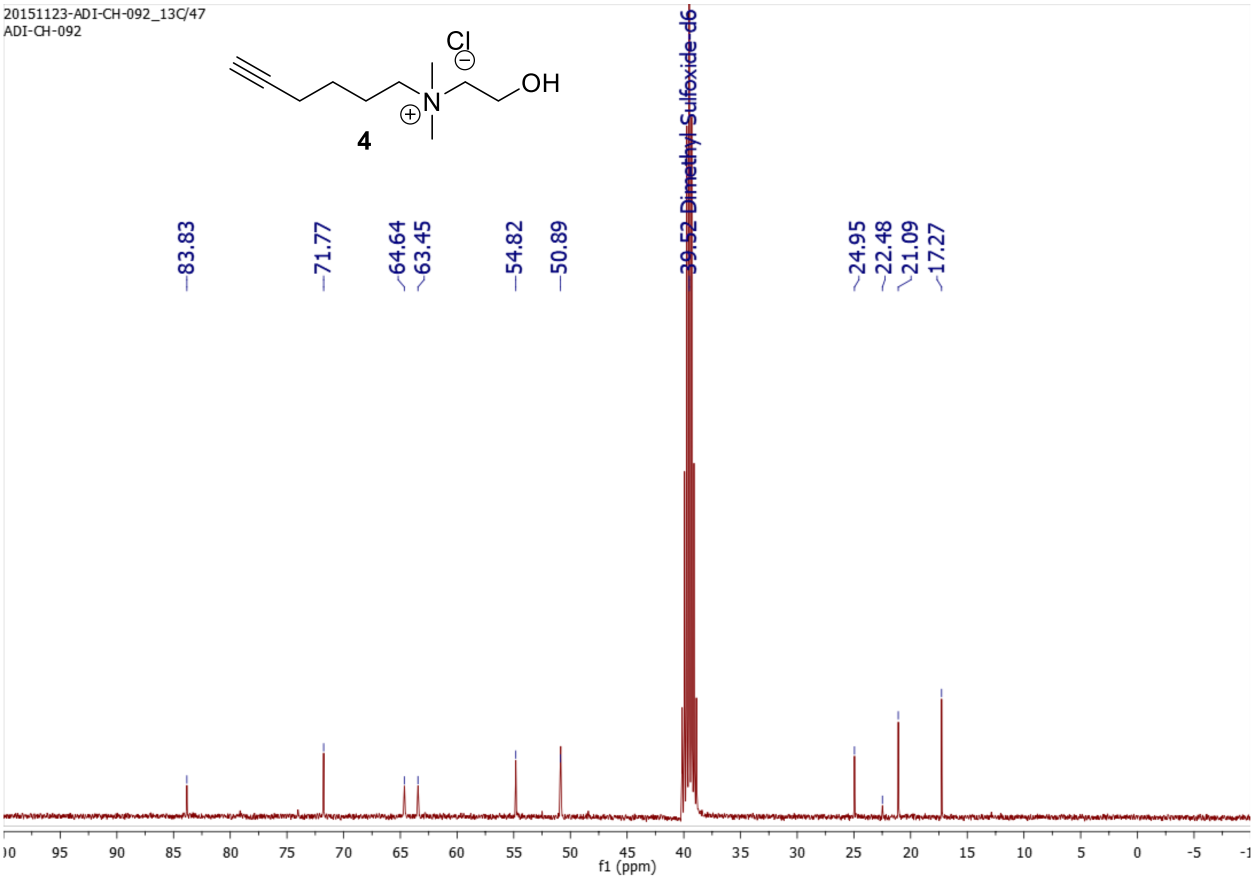
20151123-ADI-CH-092_13C/47
ADI-CH-092
Cl
OH
N
4
83.83
71.77
64.64
63.45
54.82
50.89
39.52 Dimethyl Sulfoxide-d6
24.95
22.48
21.09
17.27
f1 (ppm)

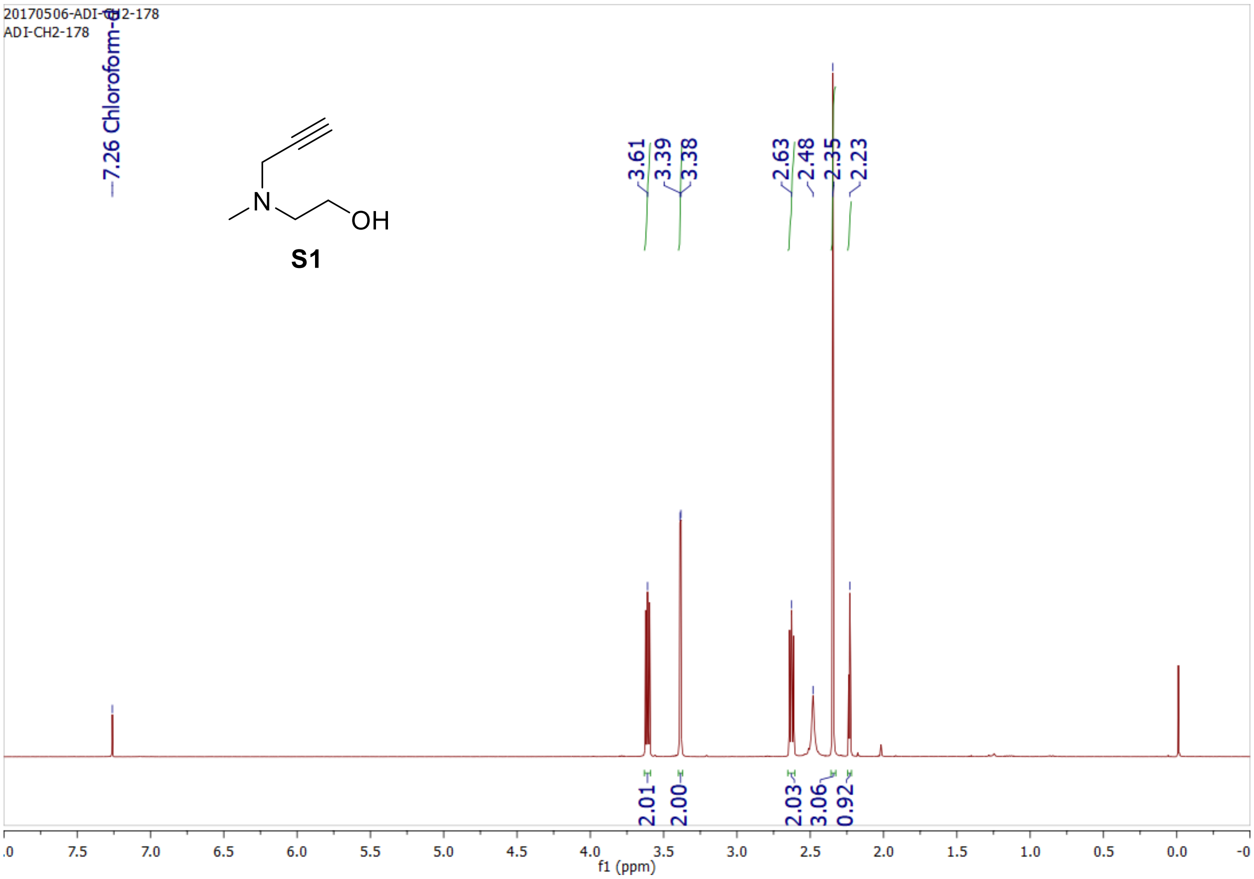
20170506-ADI-CH2-178
ADI-CH2-178
7.26 Chloroform-d
3.61
3.39
3.38
2.63
2.48
2.35
2.23
N
OH
S1
2.01
2.00
2.03
3.06
0.92
f1 (ppm)

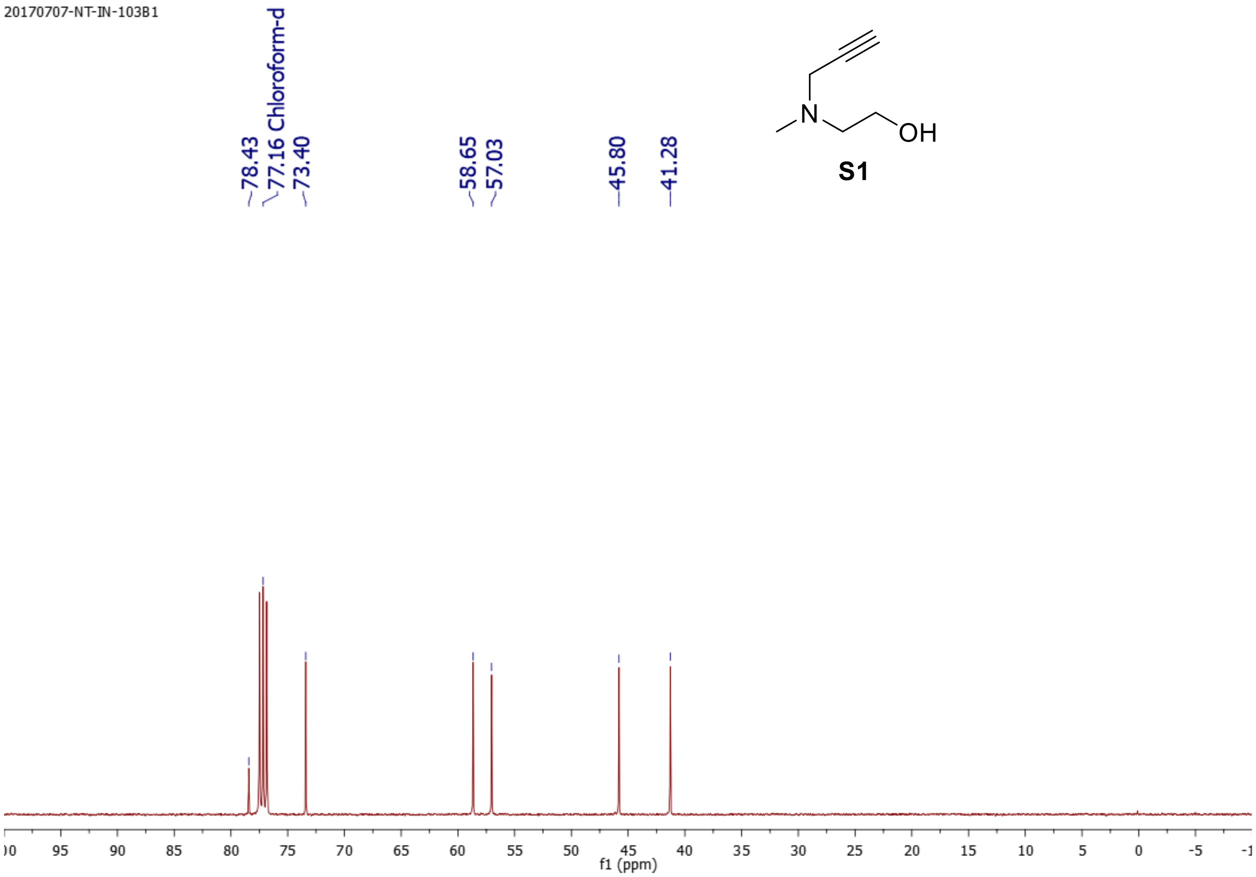
20170707-NT-IN-103B1
78.43
77.16 Chloroform-d
73.40
58.65
57.03
45.80
41.28
N
OH
S1
f1 (ppm)

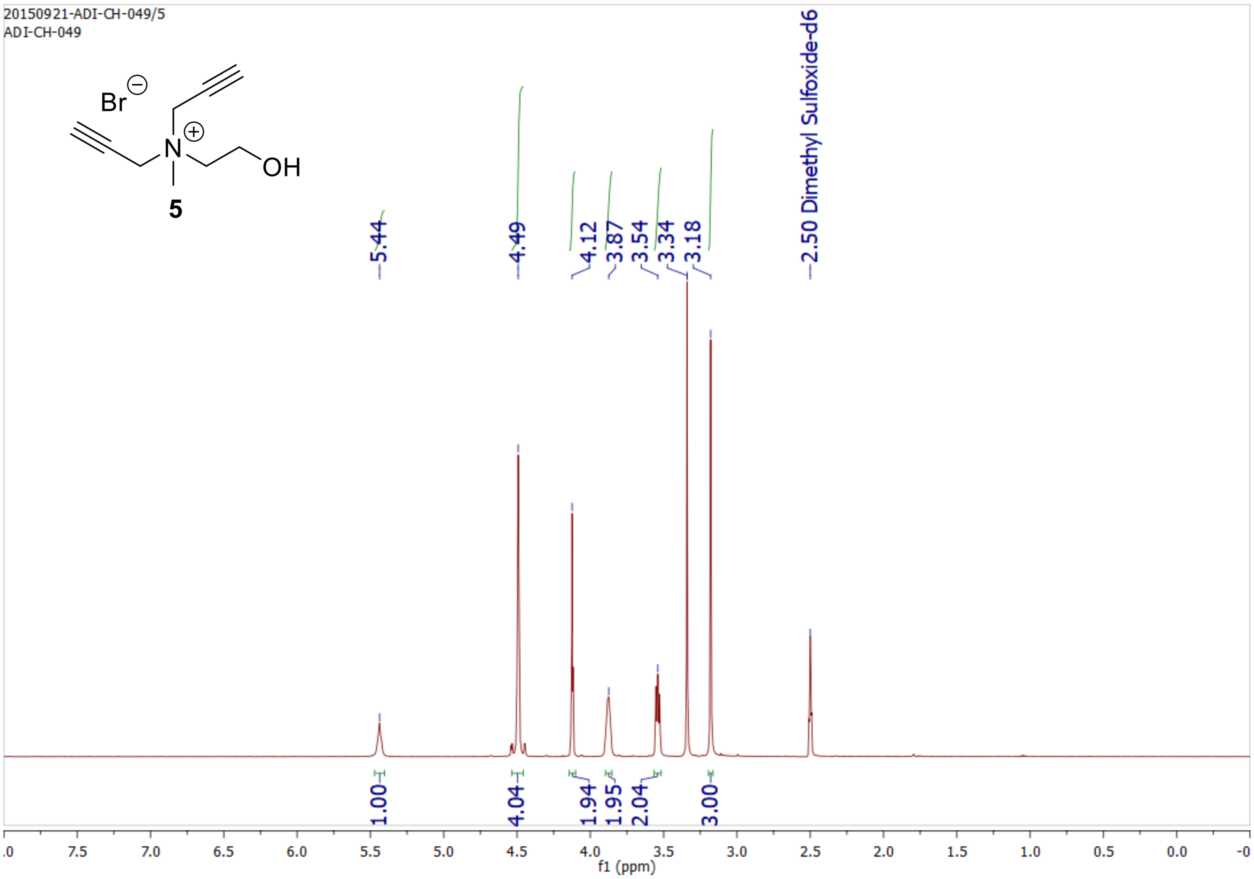
20150921-ADI-CH-049/5
ADI-CH-049
Br
OH
5
5.44
4.49
4.12
3.87
3.54
3.34
3.18
2.50 Dimethyl Sulfoxide-d6
1.00
4.04
1.94
1.95
2.04
3.00
f1 (ppm)

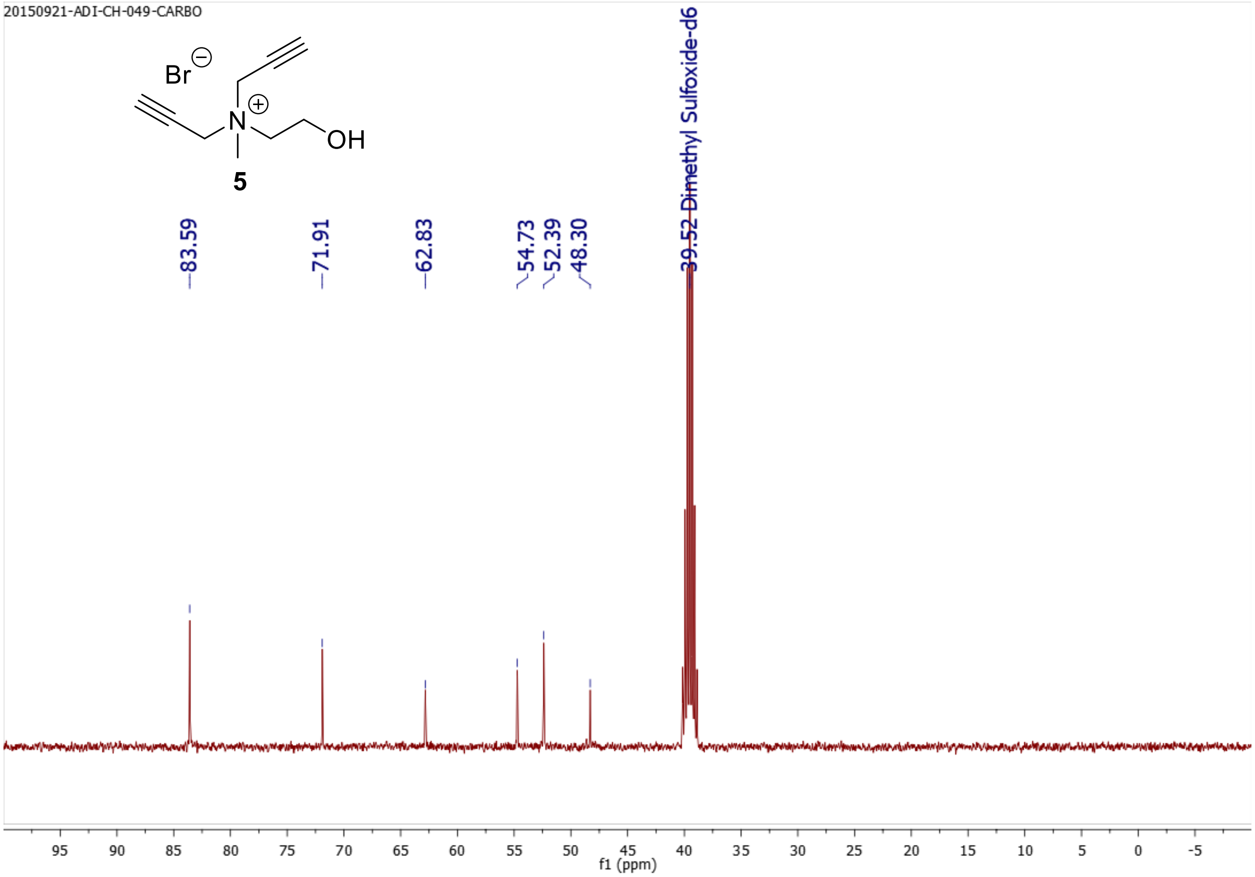
20150921-ADI-CH-049-CARBO
Br
OH
5
83.59
71.91
62.83
54.73
52.39
48.30
39.52 Dimethyl Sulfoxide-d6
f1 (ppm)

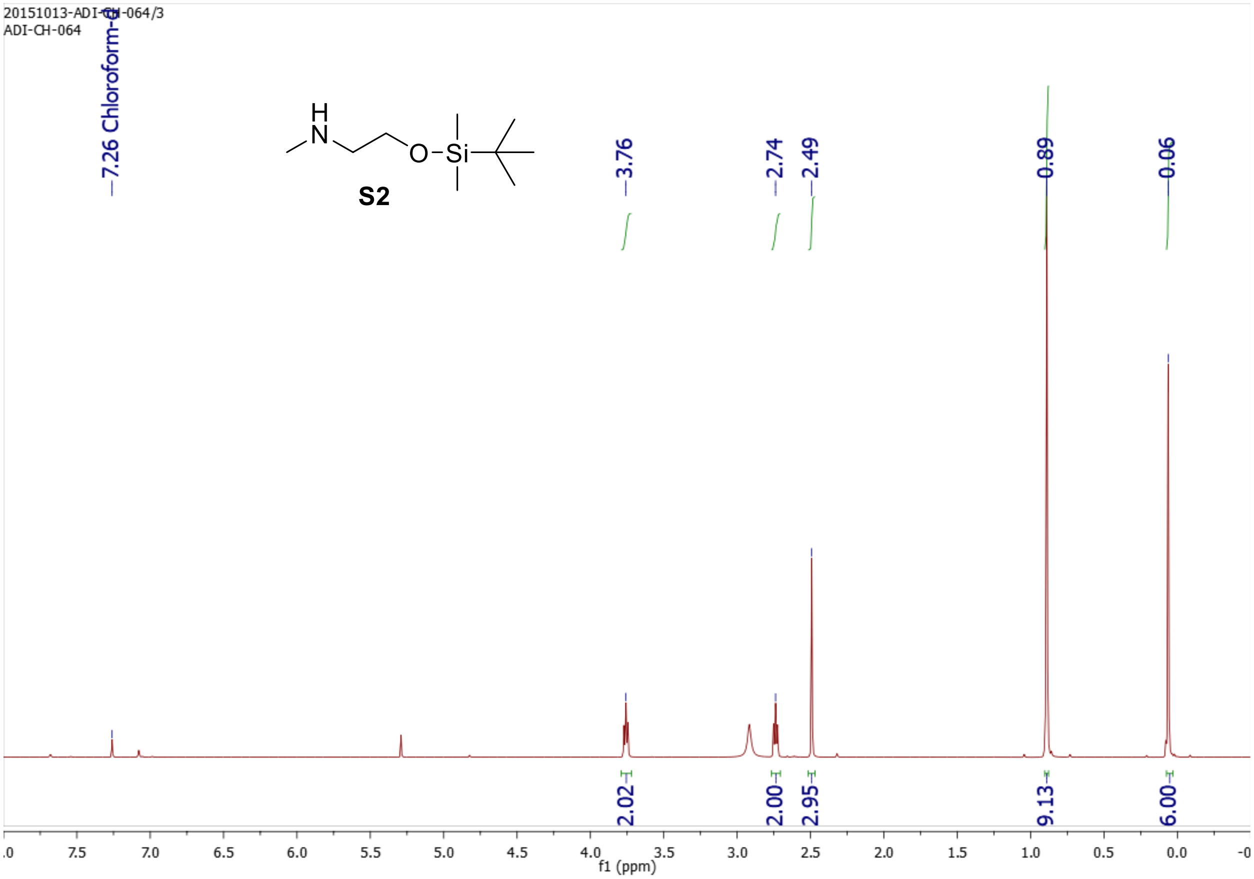
20151013-ADI-CH-064/3
ADI-CH-064
7.26 Chloroform-d
S2
3.76
2.74
2.49
0.89
0.06
2.02
2.00
2.95
9.13
6.00
f1 (ppm)

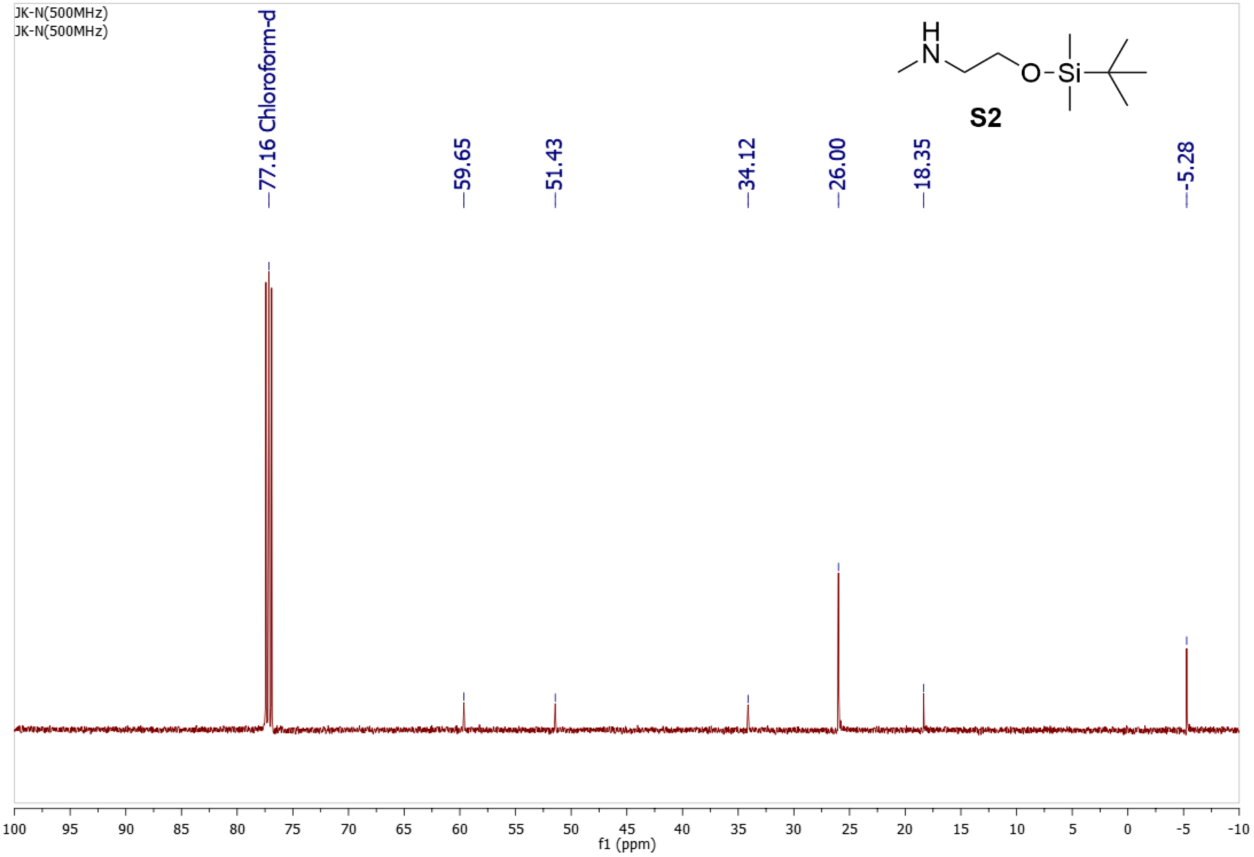
JK-N(500MHz)
JK-N(500MHz)
77.16 Chloroform-d
59.65
51.43
34.12
26.00
18.35
-5.28
S2
f1 (ppm)

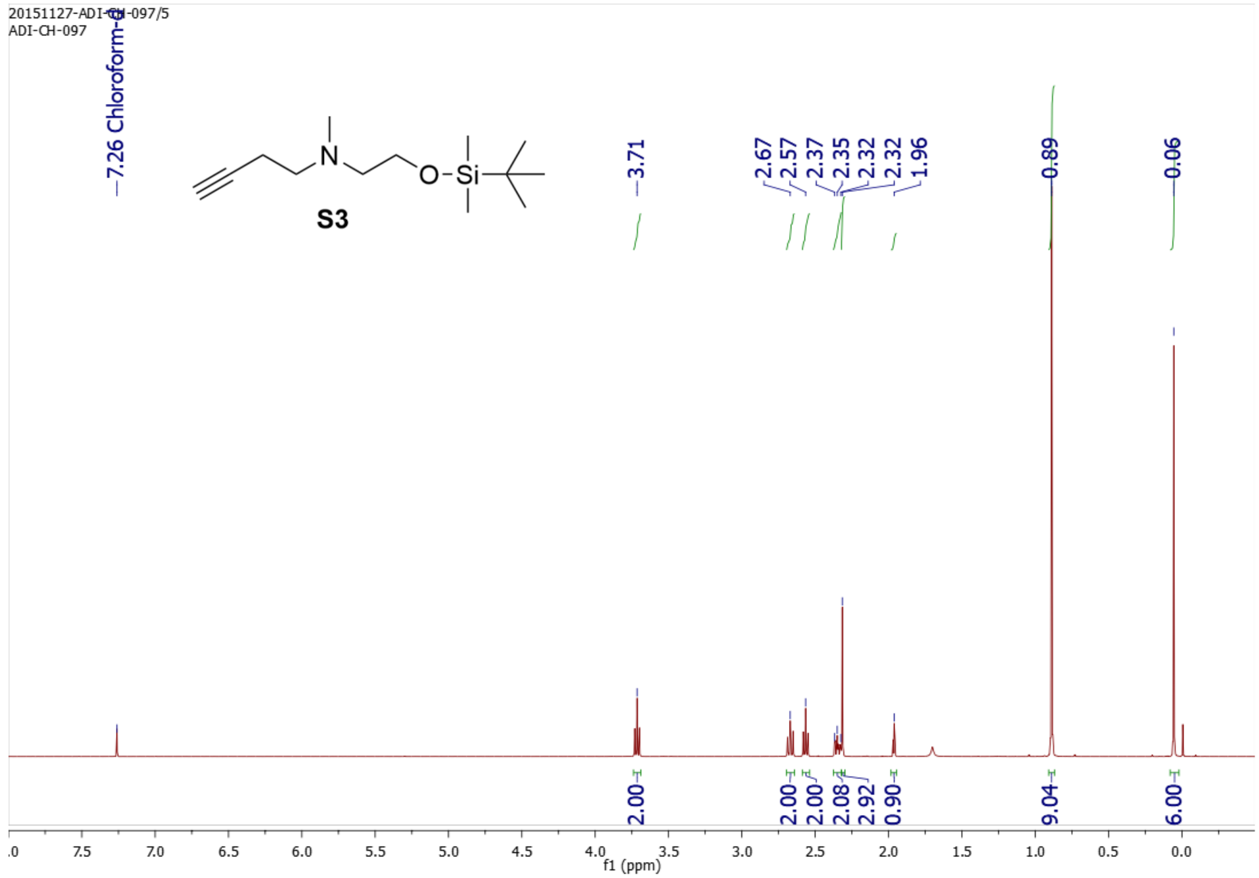

20151127-ADI-CH-097/5
ADI-CH-097
7.26 Chloroform-d
S3
3.71
2.67
2.57
2.37
2.35
2.32
2.32
1.96
0.89
0.06
2.00
2.00
2.00
2.08
2.92
0.90
9.04
6.00
f1 (ppm)


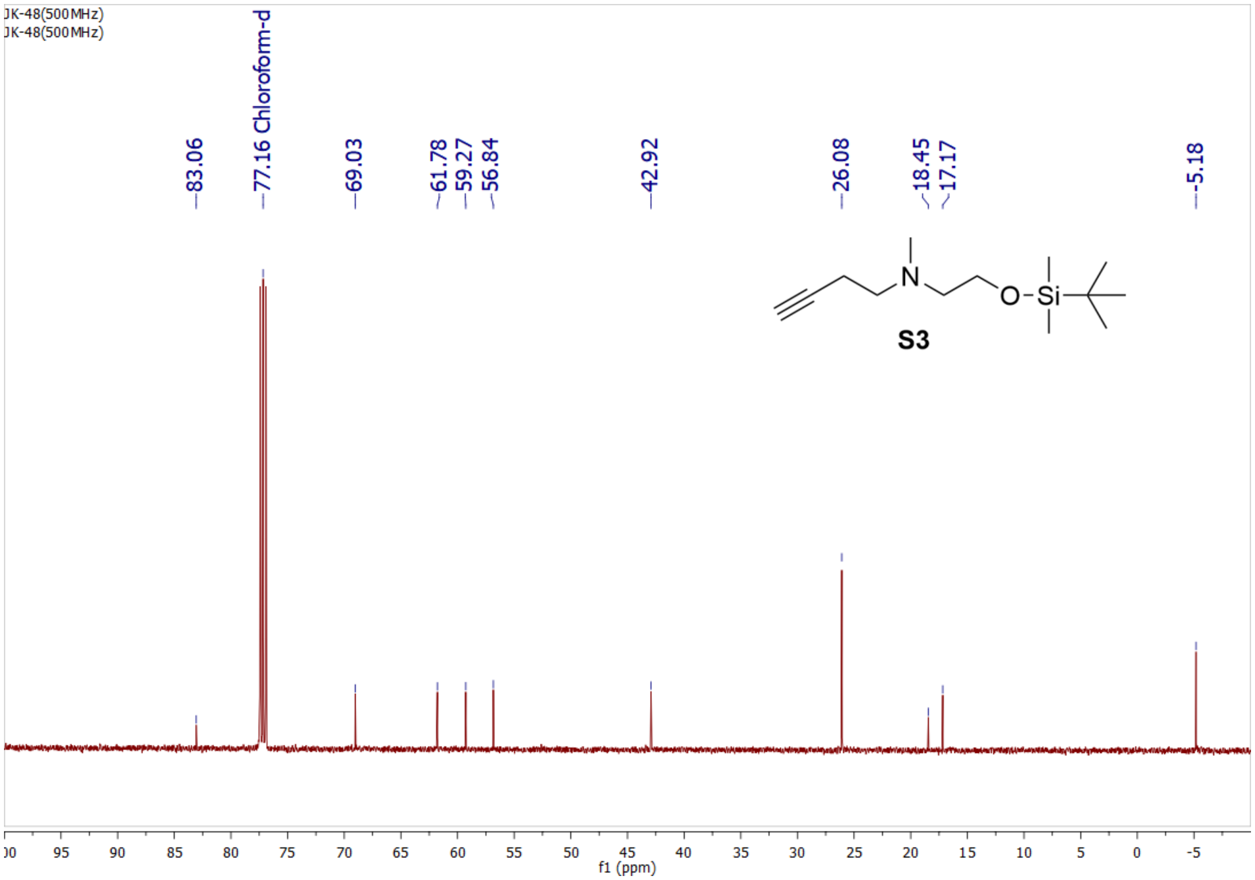

JK-48(500MHz)
JK-48(500MHz)
83.06
77.16 Chloroform-d
69.03
61.78
59.27
56.84
42.92
26.08
18.45
17.17
-5.18
S3
f1 (ppm)

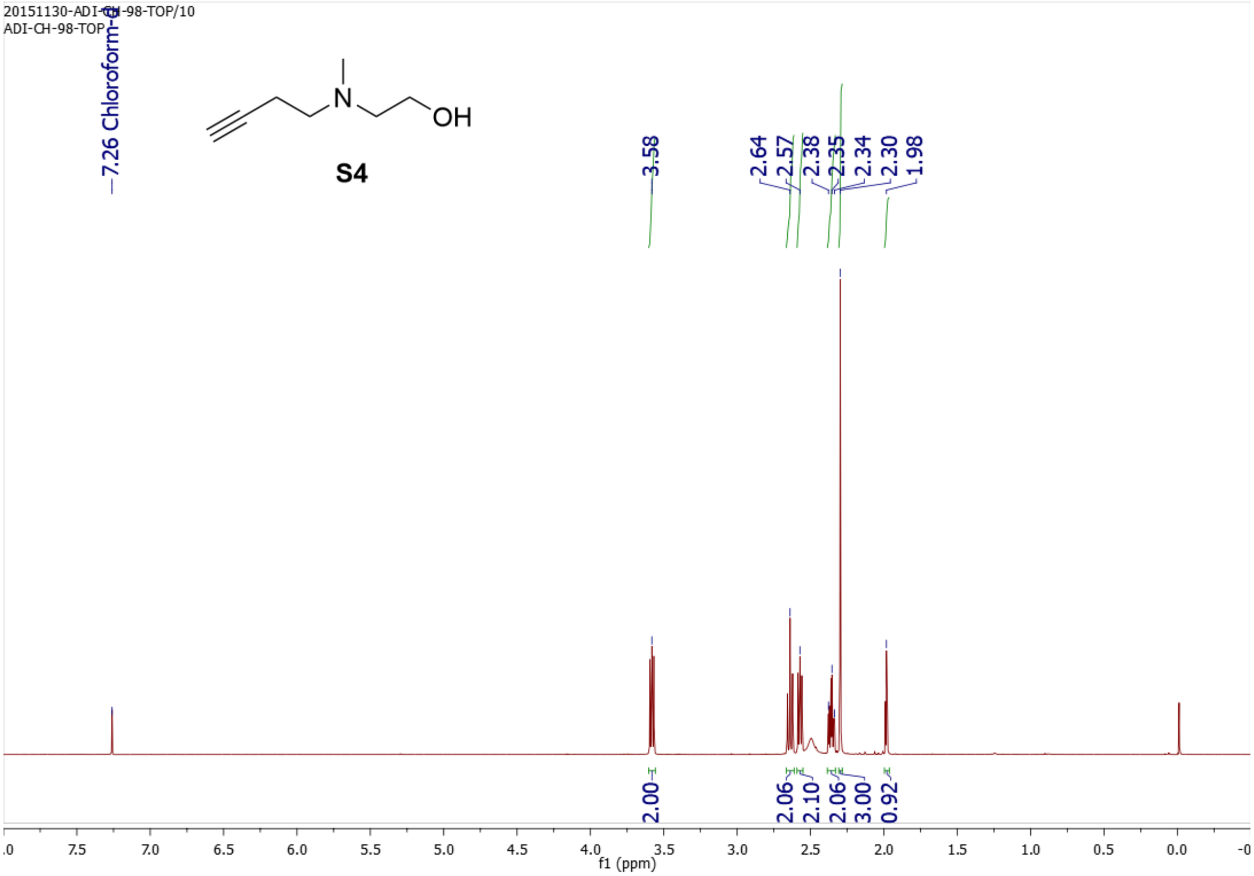
20151130-ADI-CH-98-TOP/10
ADI-CH-98-TOP
7.26 Chloroform-d
N
OH
S4
3.58
2.64
2.57
2.38
2.35
2.34
2.30
1.98
2.00
2.06
2.10
2.06
3.00
0.92
f1 (ppm)

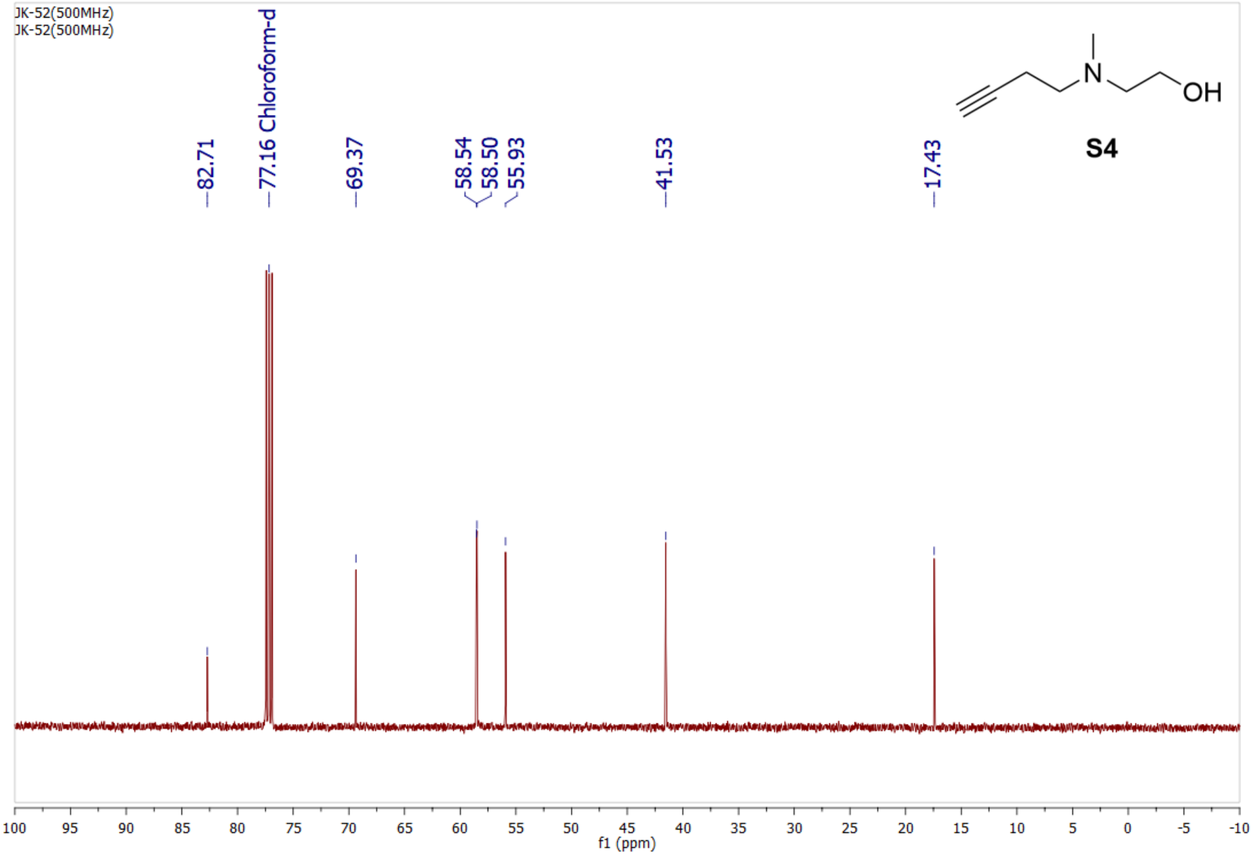
JK-52(500MHz)
JK-52(500MHz)
82.71
77.16 Chloroform-d
69.37
58.54
58.50
55.93
41.53
17.43
N
OH
S4
f1 (ppm)

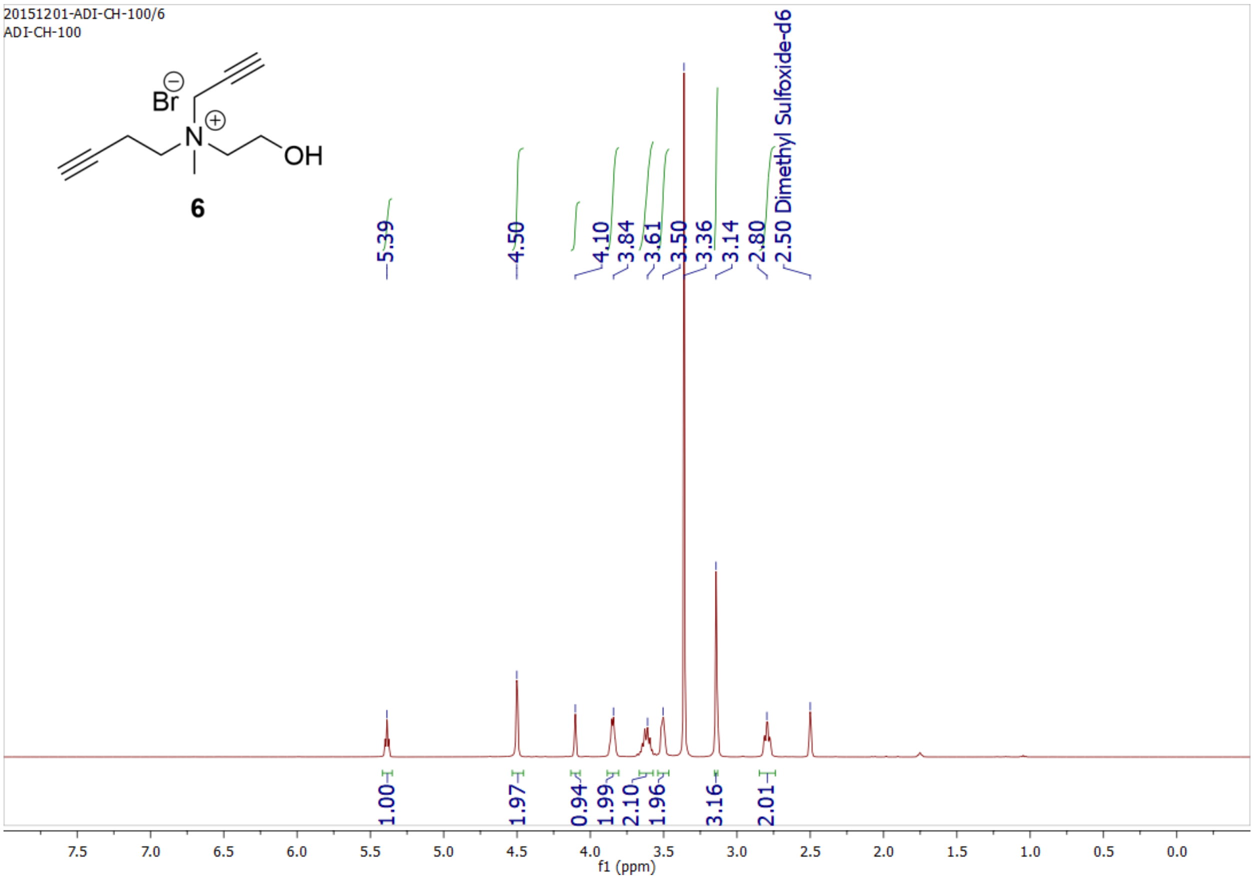

20151201-ADI-CH-100/6
ADI-CH-100
Br
N
OH
6
5.39
4.50
4.10
3.84
3.61
3.50
3.36
3.14
2.80
2.50 Dimethyl Sulfoxide-d6
1.00
1.97
0.94
1.99
2.10
1.96
3.16
2.01
7.5
7.0
6.5
6.0
5.5
5.0
4.5
4.0
3.5
3.0
2.5
2.0
1.5
1.0
0.5
0.0
f1 (ppm)


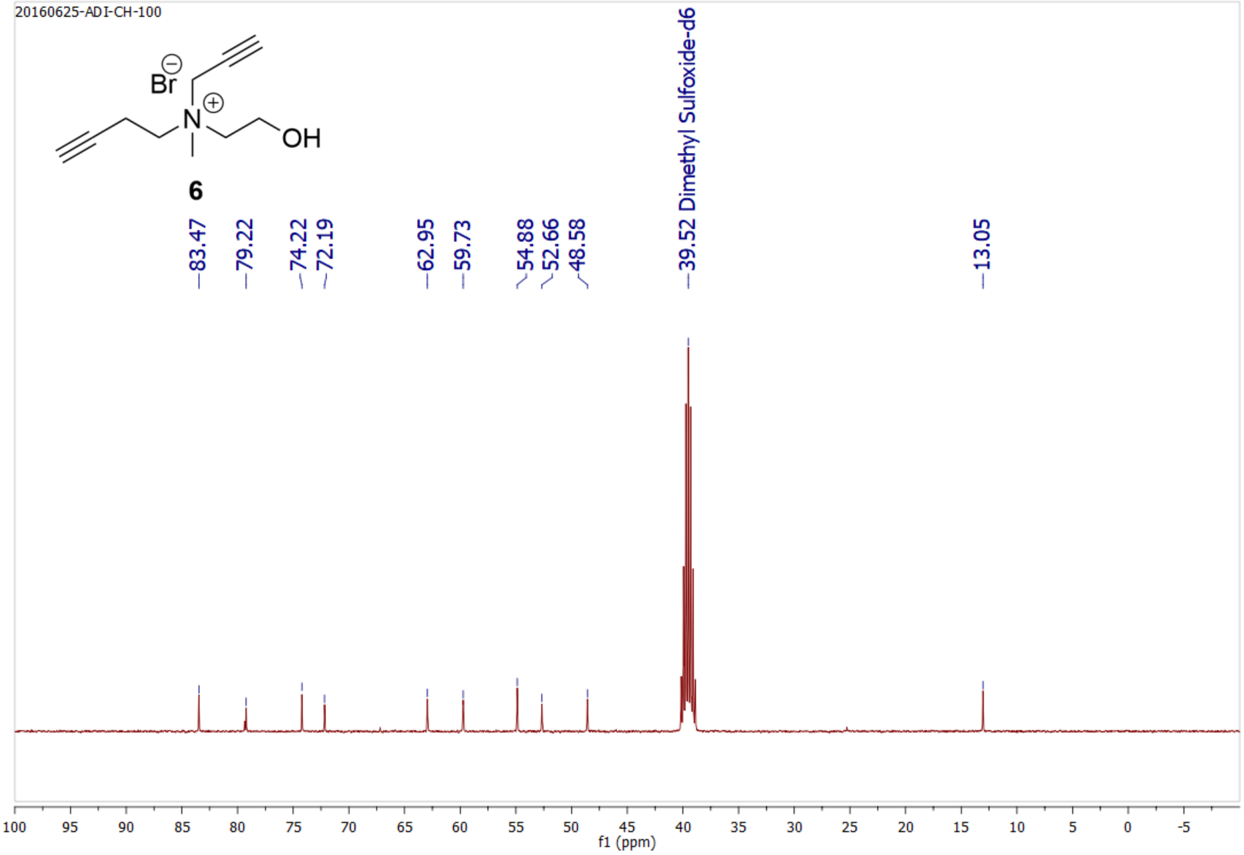

20160625-ADI-CH-100
Br
N
OH
6
83.47
79.22
74.22
72.19
62.95
59.73
54.88
52.66
48.58
39.52 Dimethyl Sulfoxide-d6
13.05
100
95
90
85
80
75
70
65
60
55
50
45
40
35
30
25
20
15
10
5
0
-5
f1 (ppm)

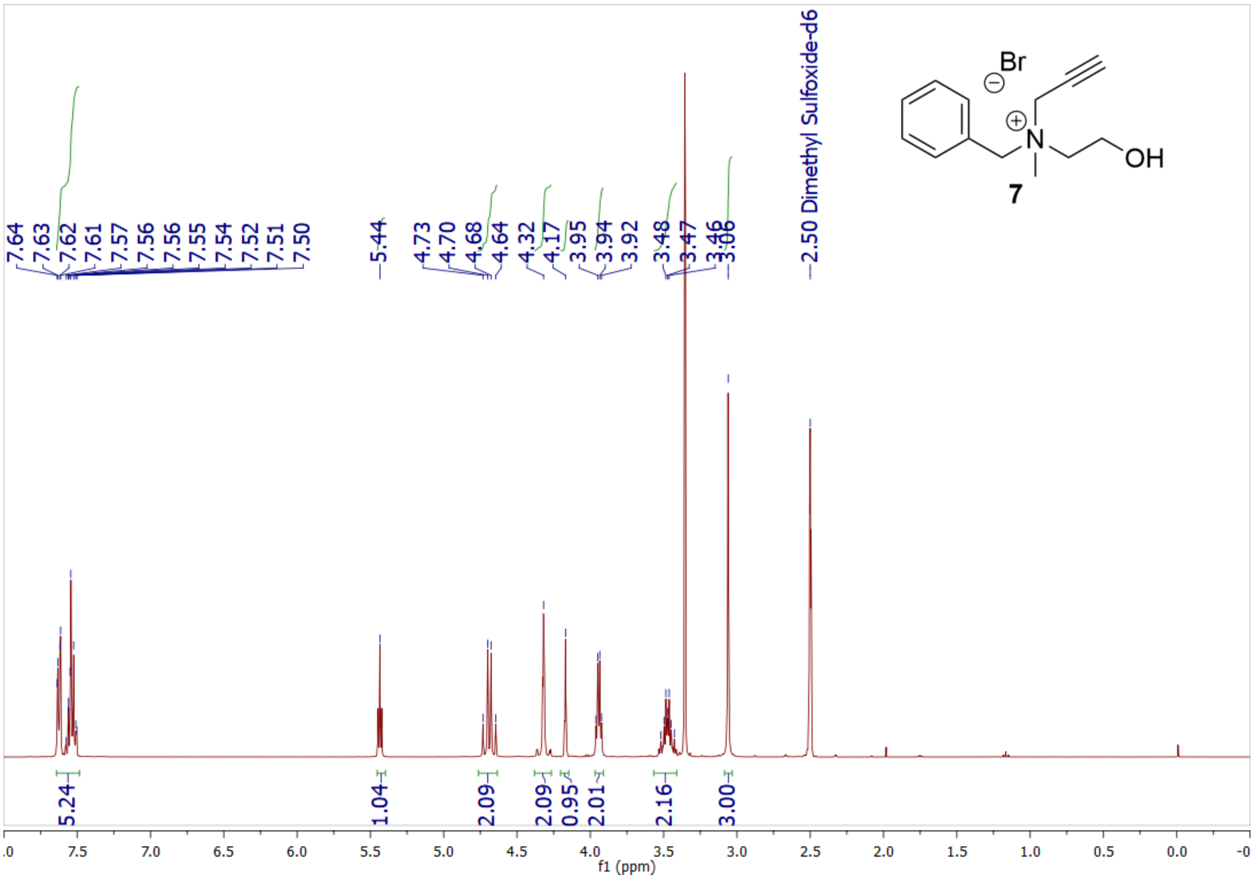

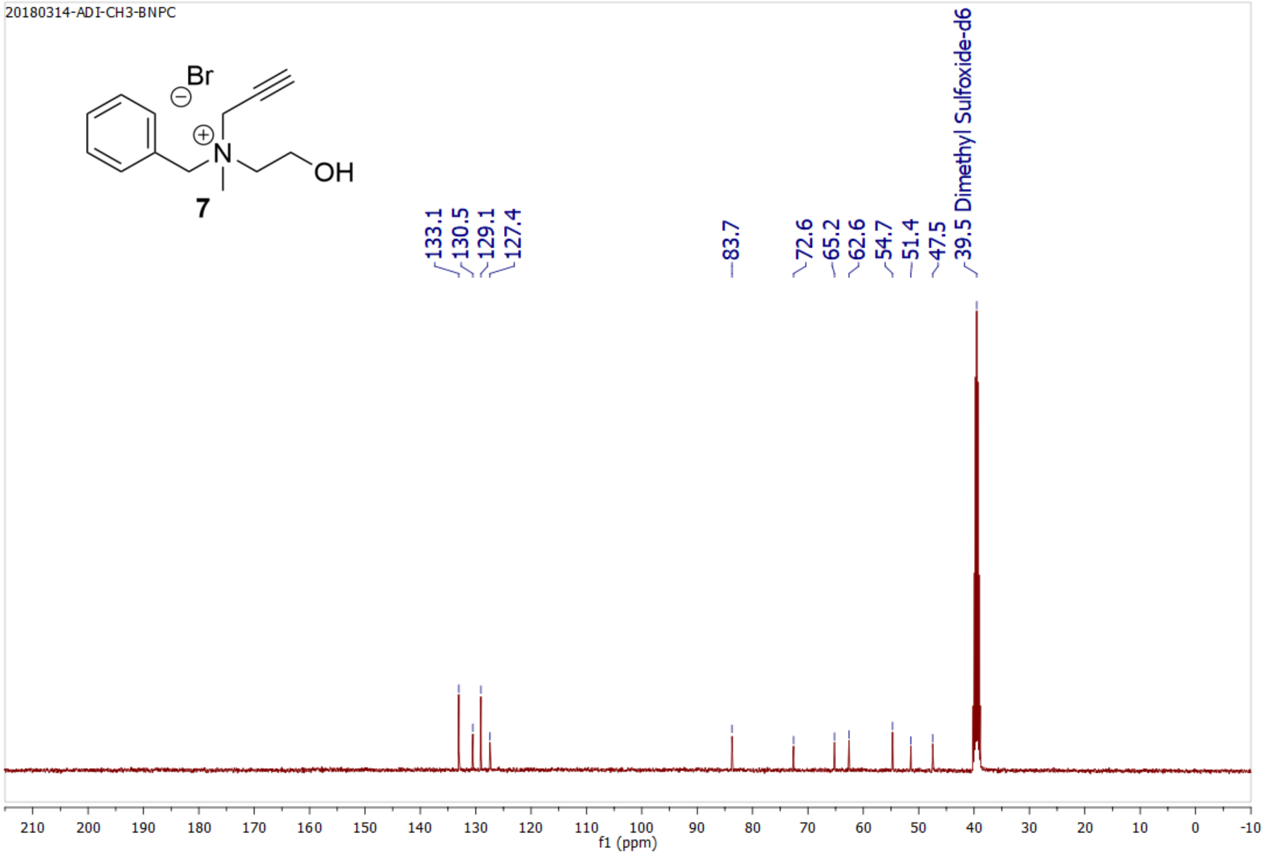

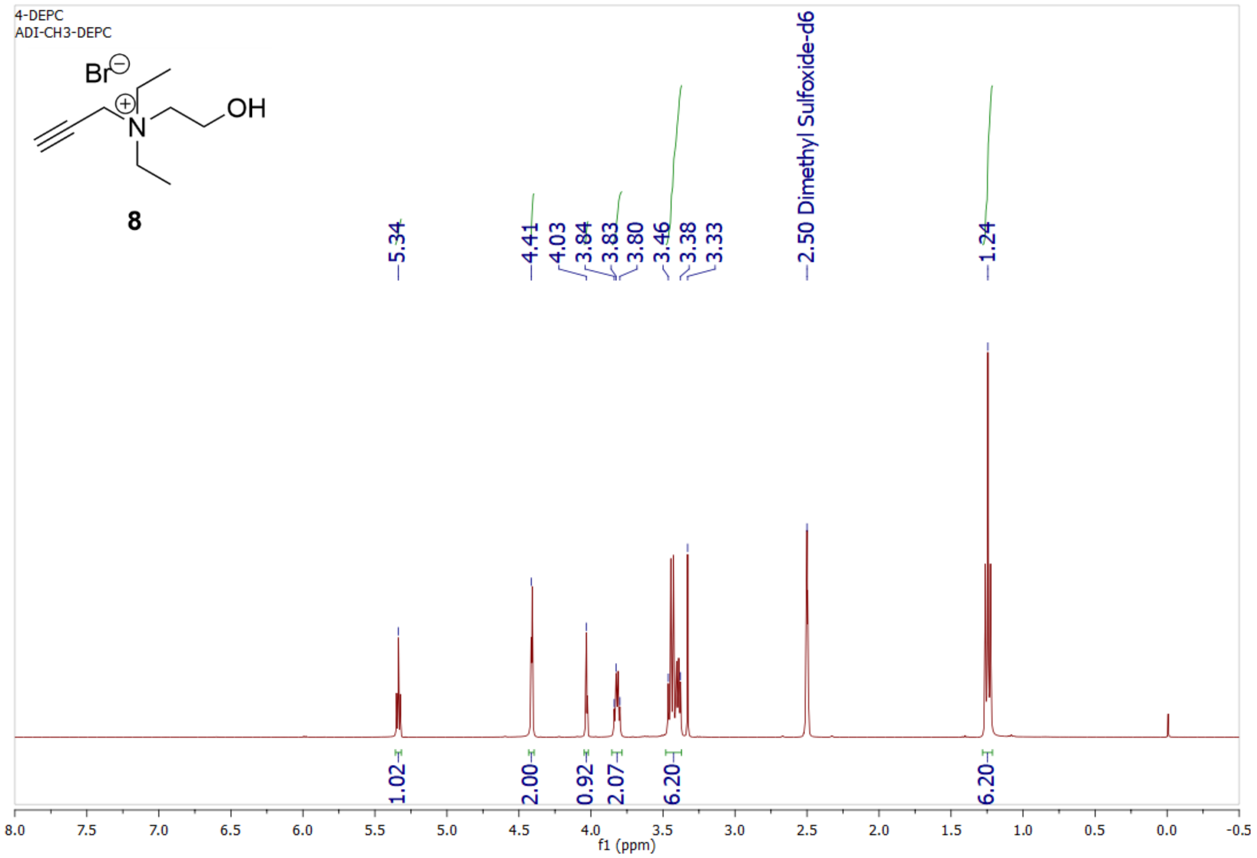
4-DEPC
ADI-CH3-DEPC
Br
OH
N
8
5.34
4.41
4.03
3.84
3.83
3.80
3.46
3.38
3.33
2.50 Dimethyl Sulfoxide-d6
1.24
1.02
2.00
0.92
2.07
6.20
6.20
f1 (ppm)

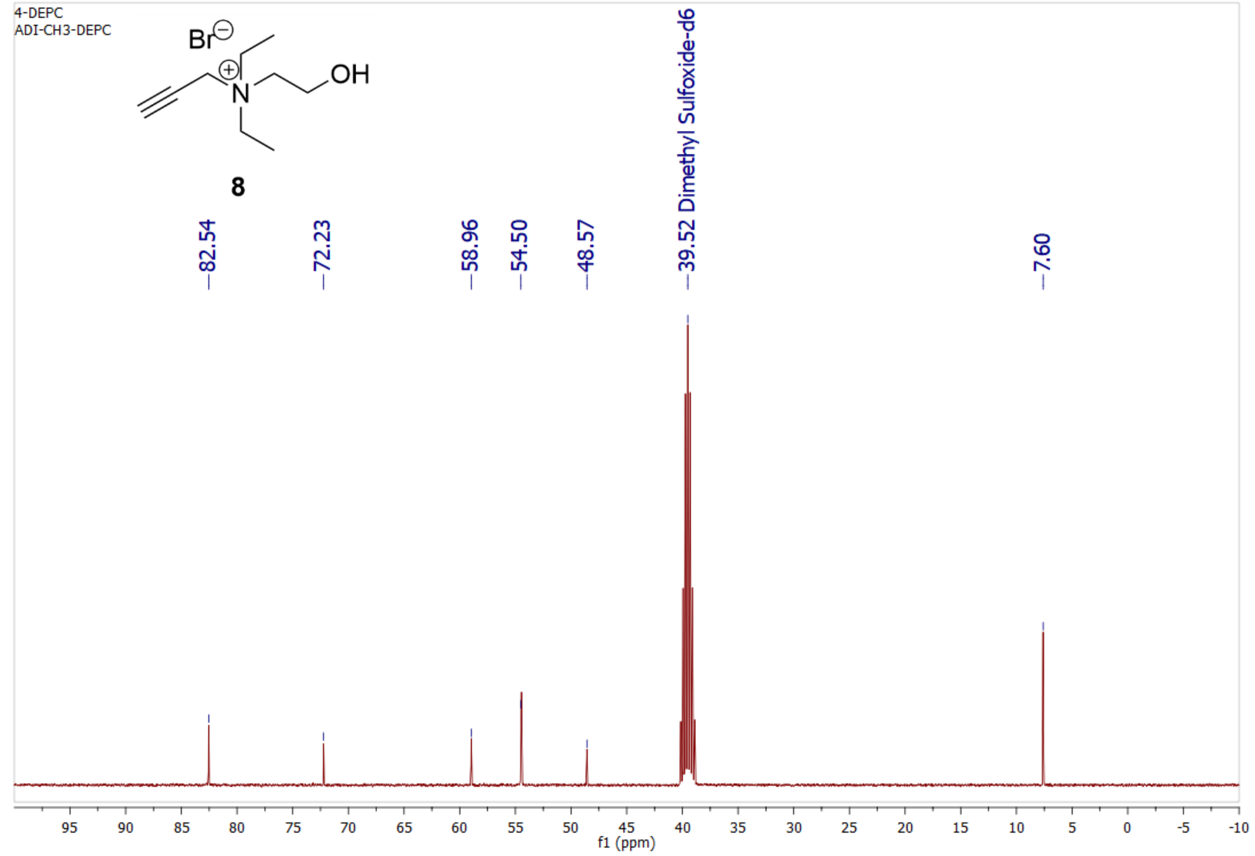
4-DEPC
ADI-CH3-DEPC
Br
OH
N
8
82.54
72.23
58.96
54.50
48.57
39.52 Dimethyl Sulfoxide-d6
7.60
f1 (ppm)

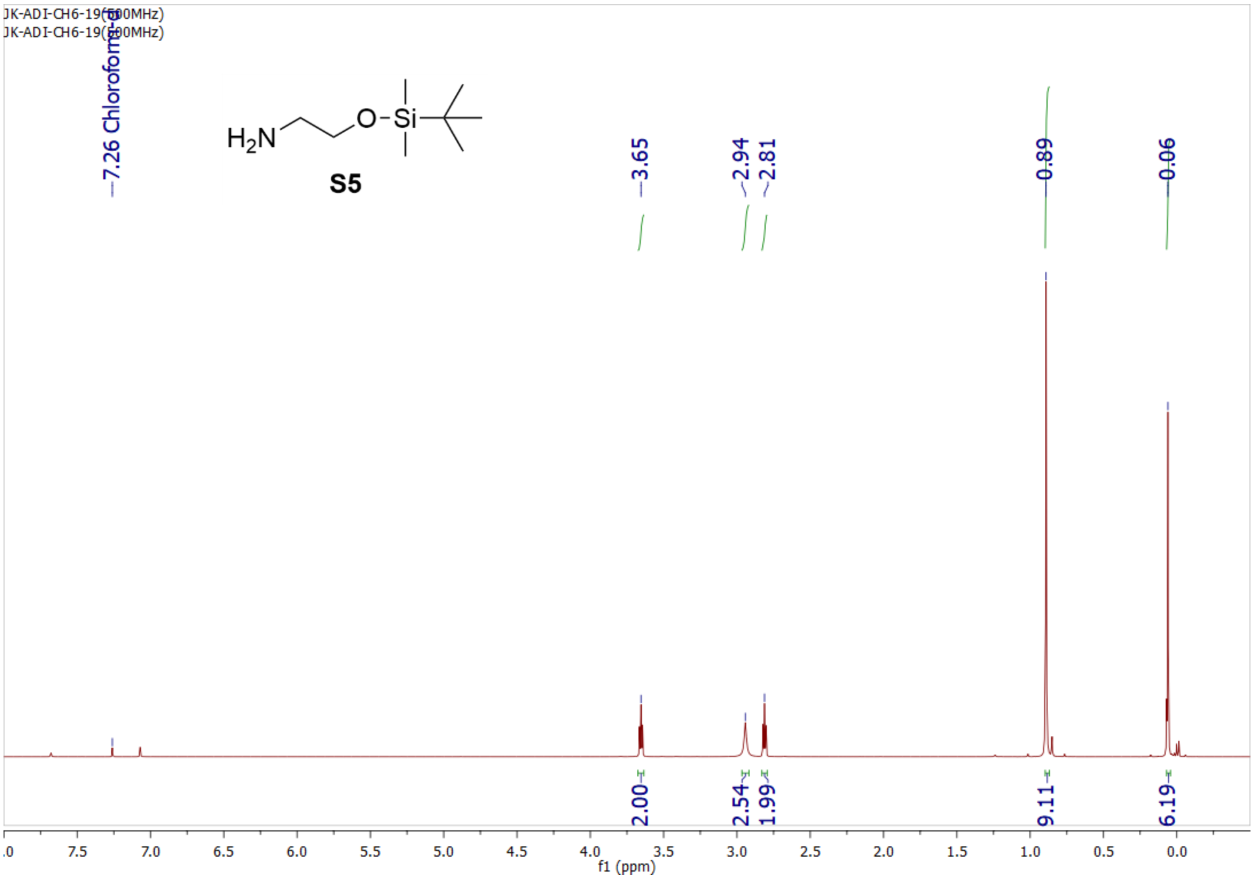
JK-ADI-CH6-19(500MHz)
JK-ADI-CH6-19(500MHz)
7.26 Chloroform-d
$H_2N$
O
Si
S5
3.65
2.94
2.81
0.89
0.06
2.00
2.54
1.99
9.11
6.19
f1 (ppm)

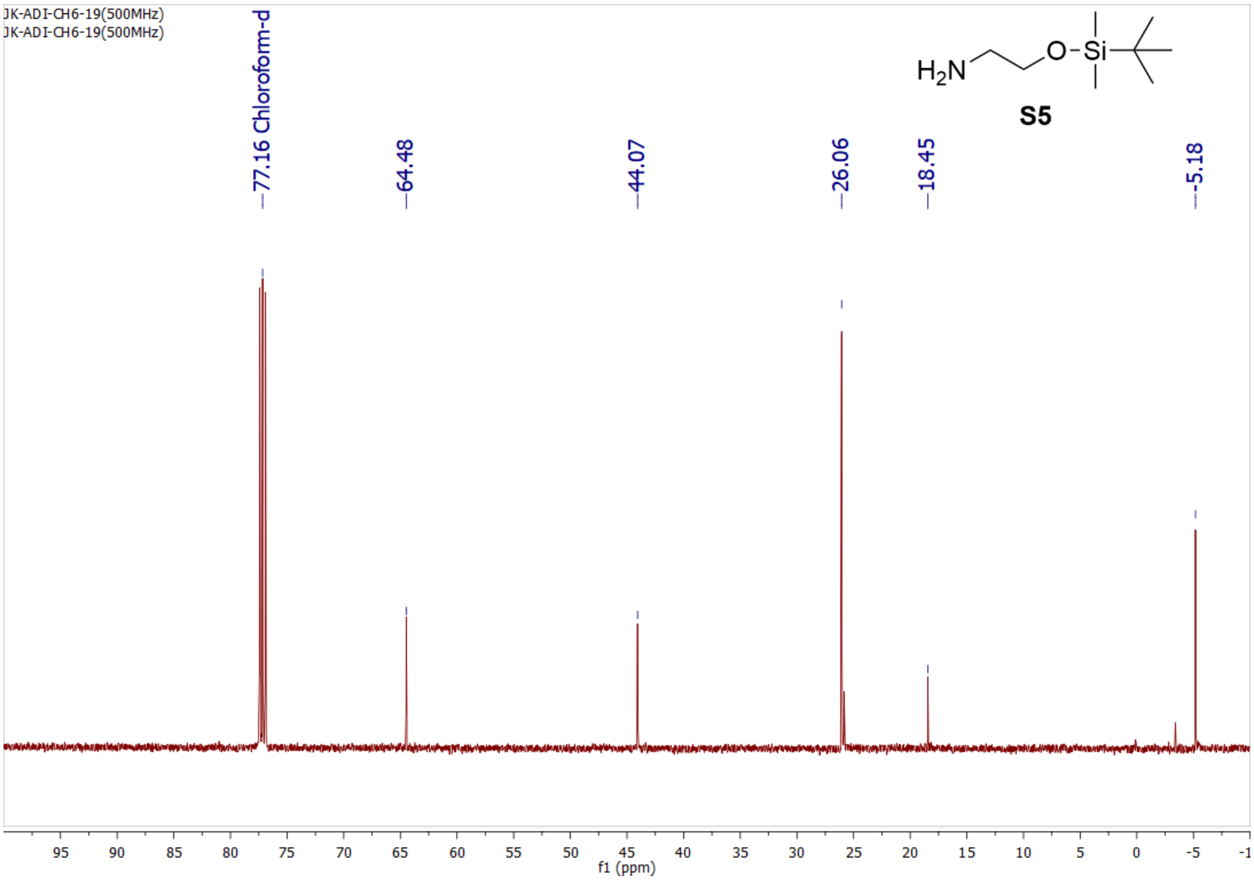
JK-ADI-CH6-19(500MHz)
JK-ADI-CH6-19(500MHz)
77.16 Chloroform-d
64.48
44.07
26.06
18.45
-5.18
$H_2N$
O
Si
S5
f1 (ppm)

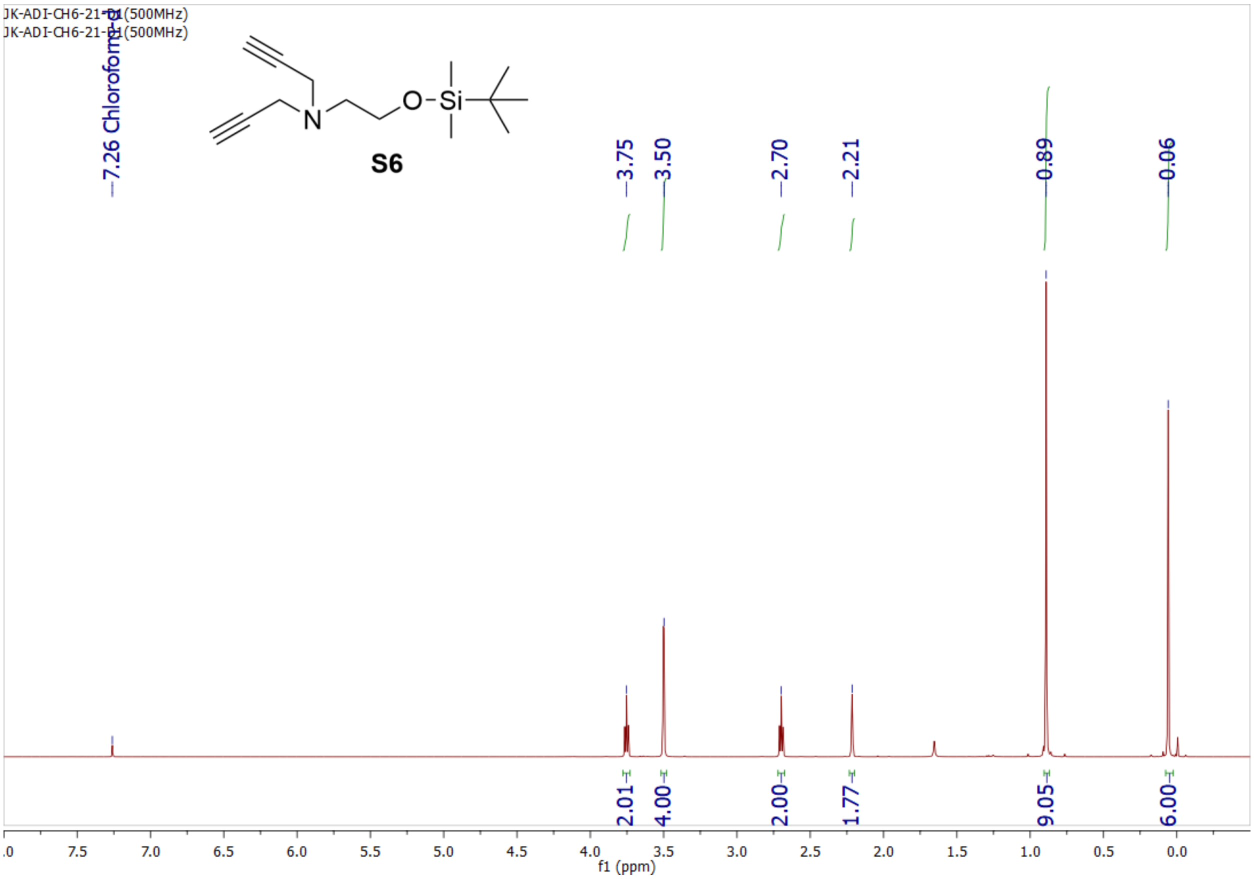
JK-ADI-CH6-21-D1(500MHz)
JK-ADI-CH6-21-D1(500MHz)
7.26 Chloroform-d
S6
3.75
3.50
2.70
2.21
0.89
0.06
2.01
4.00
2.00
1.77
9.05
6.00
f1 (ppm)

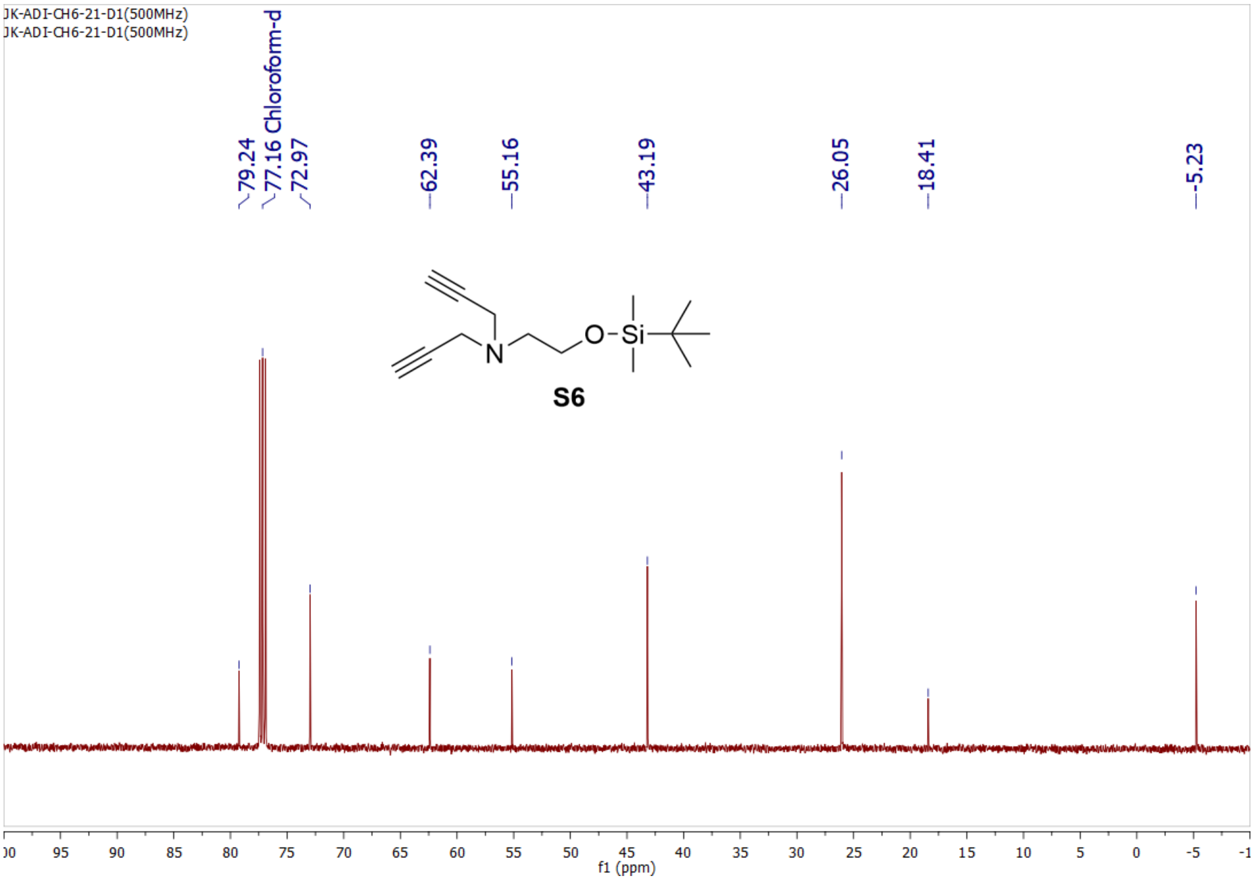
JK-ADI-CH6-21-D1(500MHz)
JK-ADI-CH6-21-D1(500MHz)
79.24
77.16 Chloroform-d
72.97
62.39
55.16
43.19
26.05
18.41
-5.23
S6
f1 (ppm)

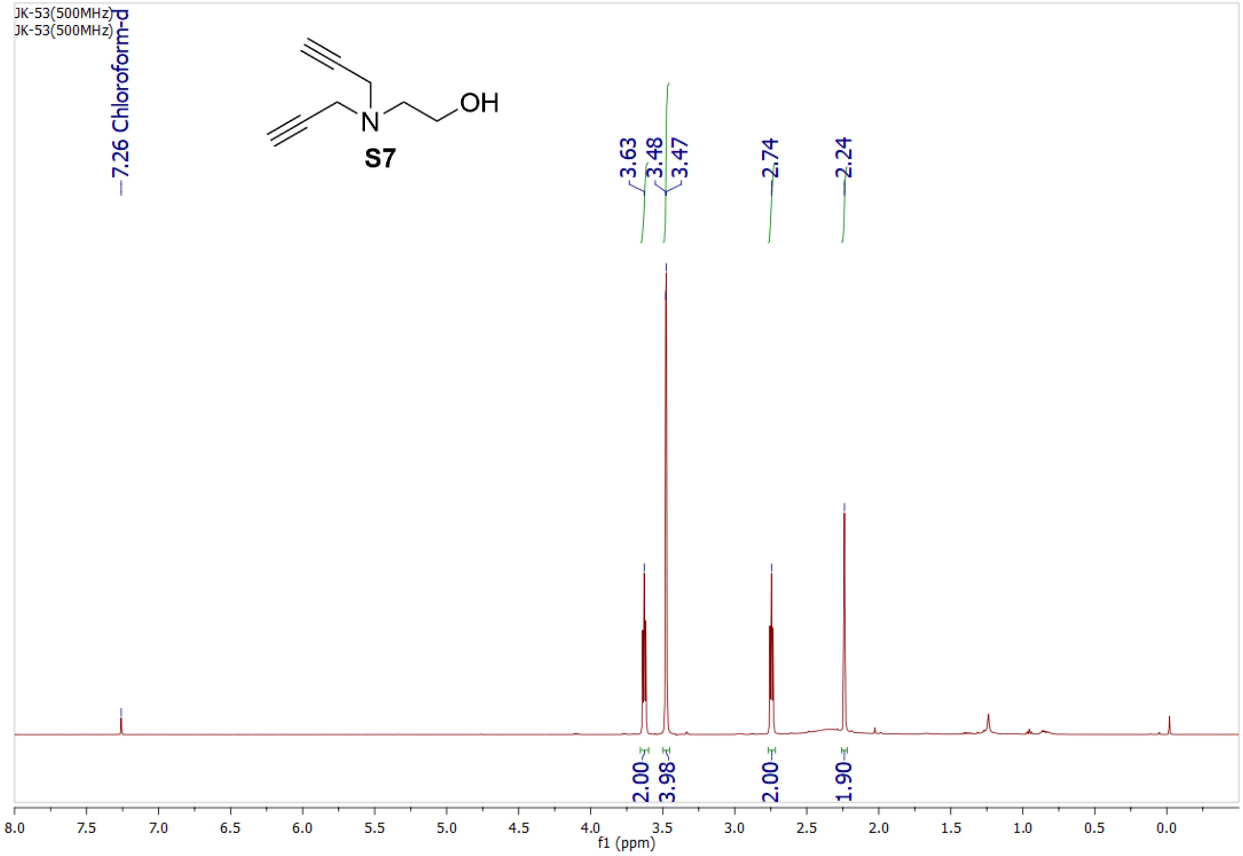
JK-53(500MHz)
JK-53(500MHz)
7.26 Chloroform-d
OH
N
S7
3.63
3.48
3.47
2.74
2.24
2.00
3.98
2.00
1.90
8.0 7.5 7.0 6.5 6.0 5.5 5.0 4.5 4.0 3.5 3.0 2.5 2.0 1.5 1.0 0.5 0.0
f1 (ppm)

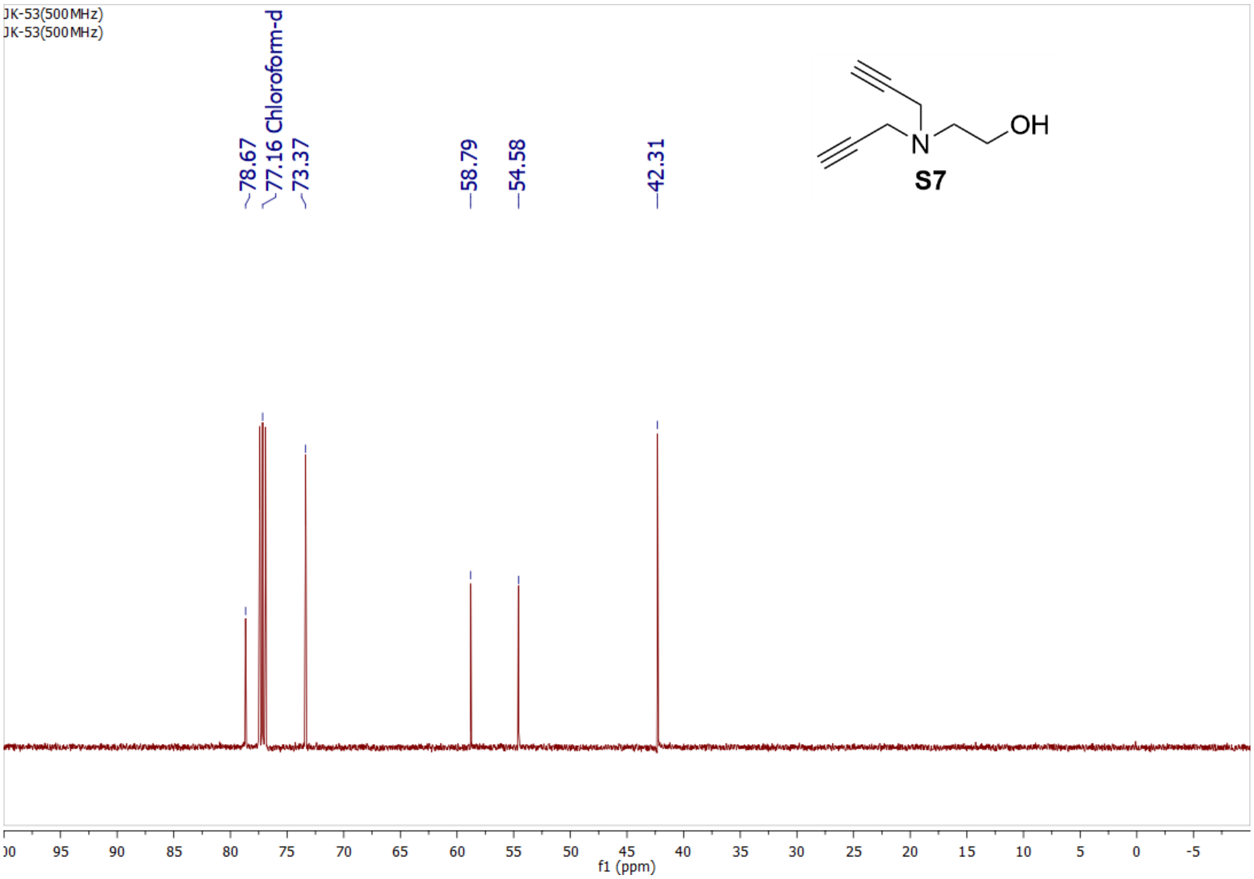
JK-53(500MHz)
JK-53(500MHz)
78.67
77.16 Chloroform-d
73.37
58.79
54.58
42.31
OH
N
S7
95 90 85 80 75 70 65 60 55 50 45 40 35 30 25 20 15 10 5 0 -5
f1 (ppm)

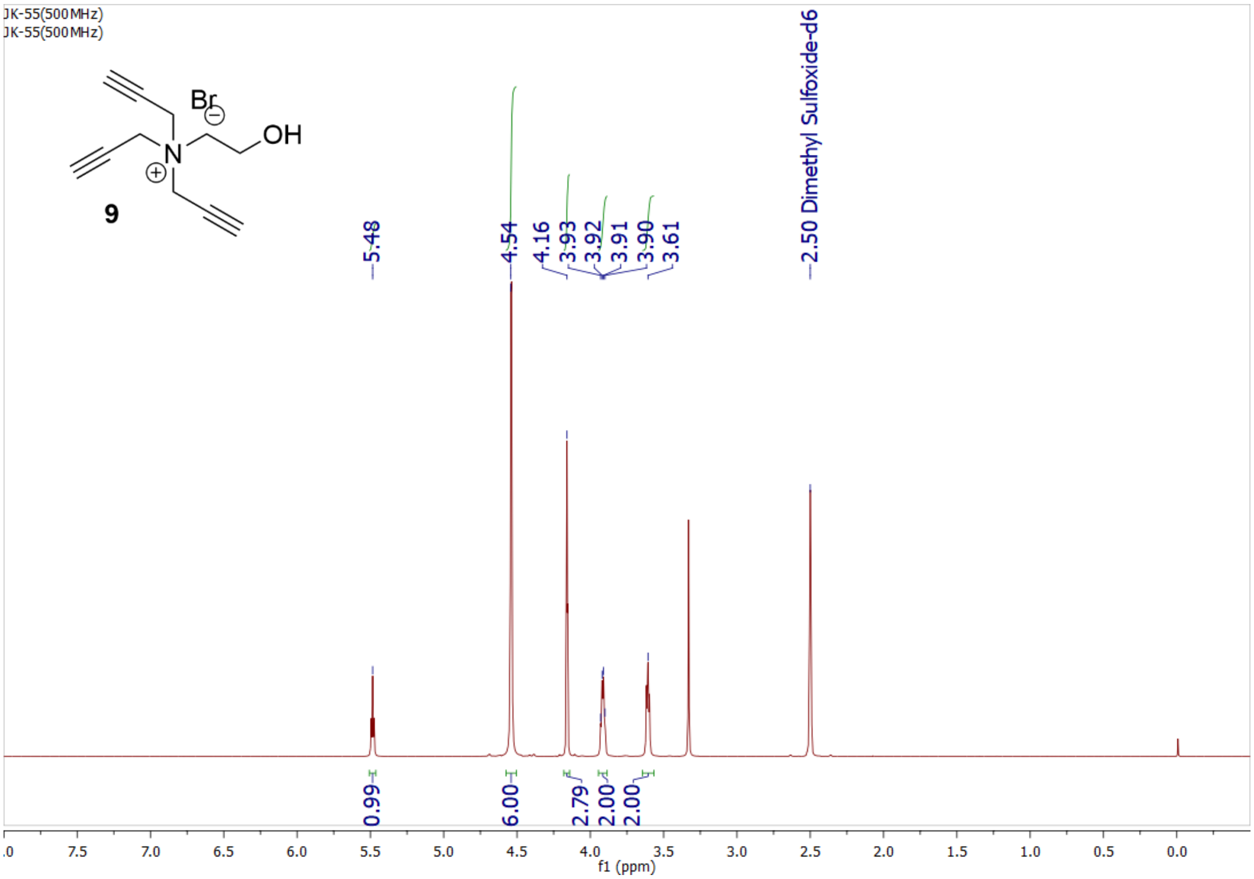
JK-55(500MHz)
JK-55(500MHz)
Br
OH
9
5.48
4.54
4.16
3.93
3.92
3.91
3.90
3.61
2.50 Dimethyl Sulfoxide-d6
0.99
6.00
2.79
2.00
2.00
f1 (ppm)

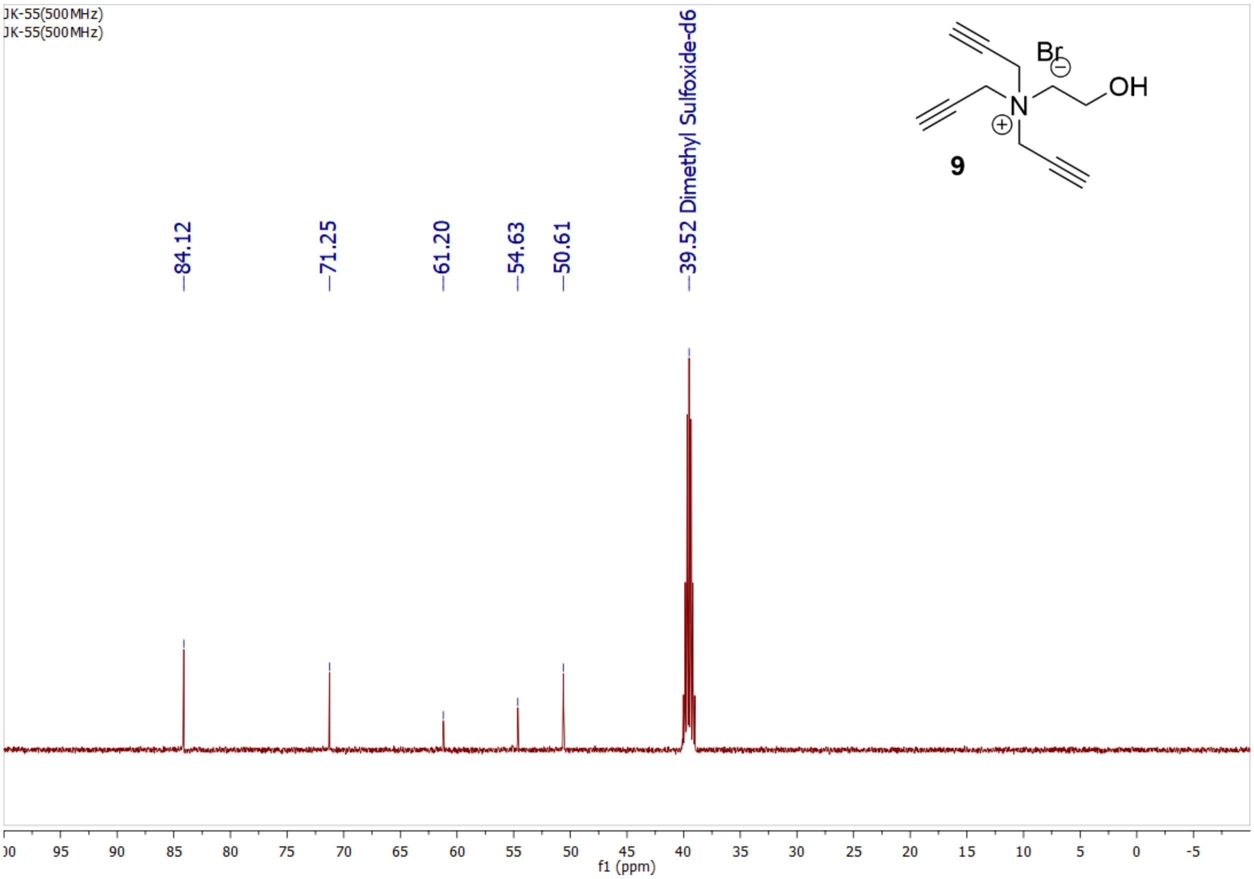
JK-55(500MHz)
JK-55(500MHz)
Br
OH
9
84.12
71.25
61.20
54.63
50.61
39.52 Dimethyl Sulfoxide-d6
f1 (ppm)

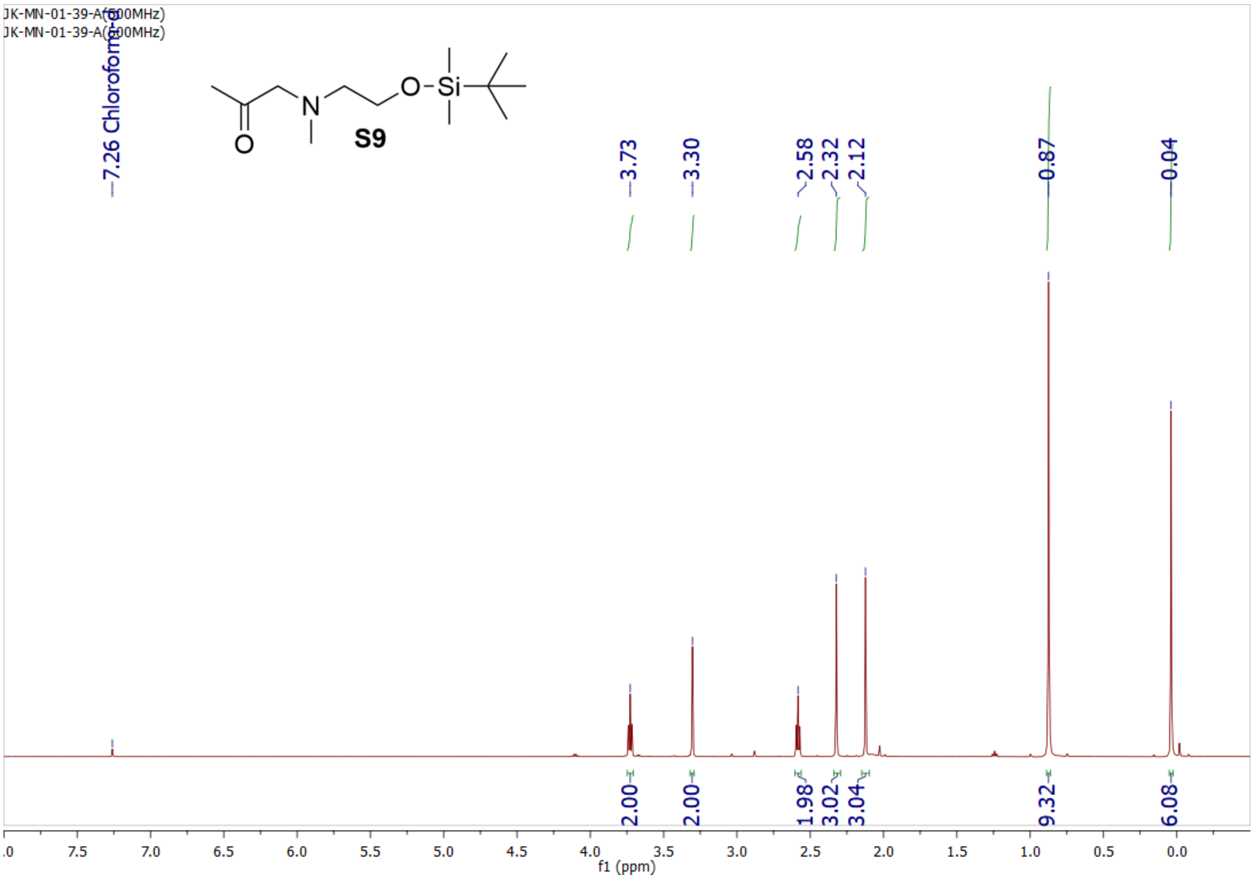
JK-MN-01-39-A(500MHz)
JK-MN-01-39-A(500MHz)
7.26 Chloroform-d
S9
3.73
3.30
2.58
2.32
2.12
0.87
0.04
2.00
2.00
1.98
3.02
3.04
9.32
6.08
f1 (ppm)

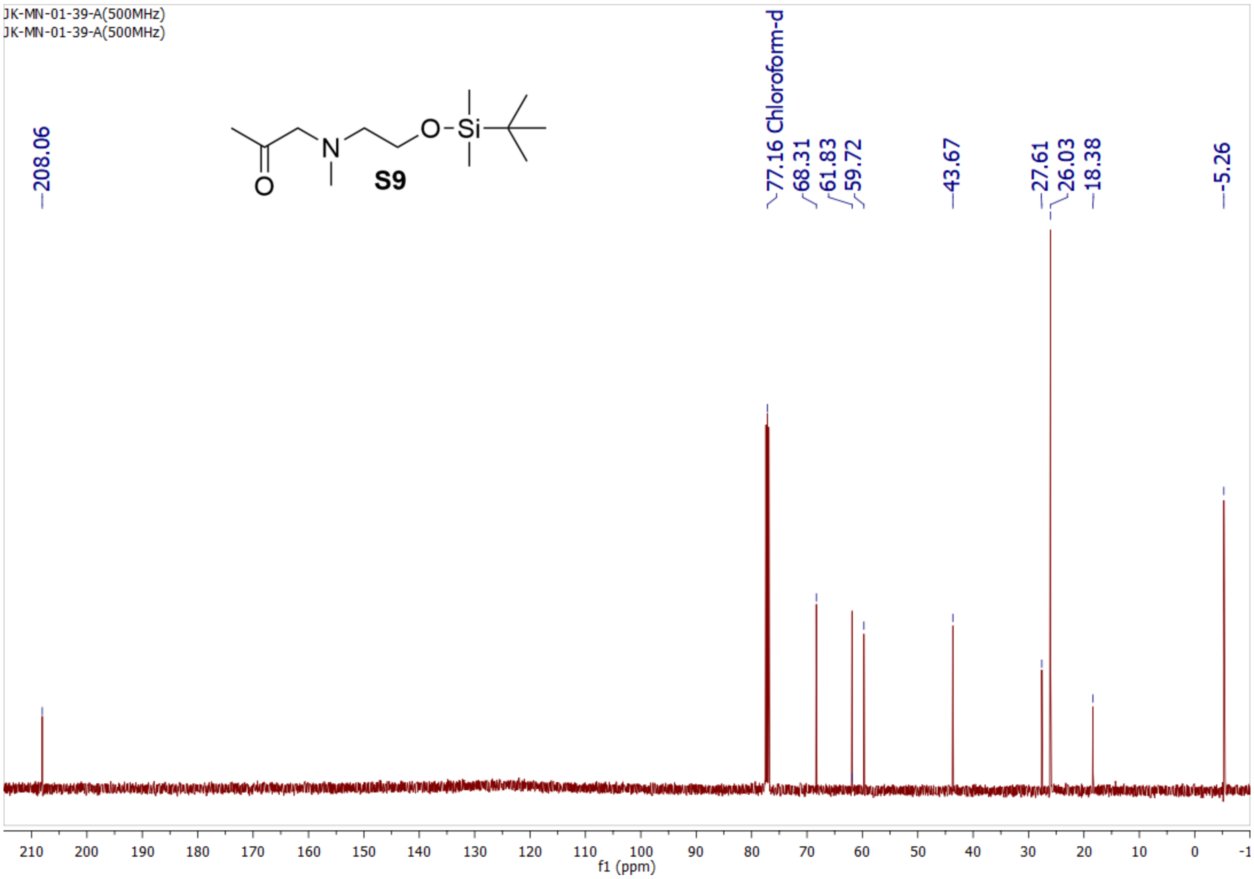
JK-MN-01-39-A(500MHz)
JK-MN-01-39-A(500MHz)
208.06
S9
77.16 Chloroform-d
68.31
61.83
59.72
43.67
27.61
26.03
18.38
-5.26
f1 (ppm)

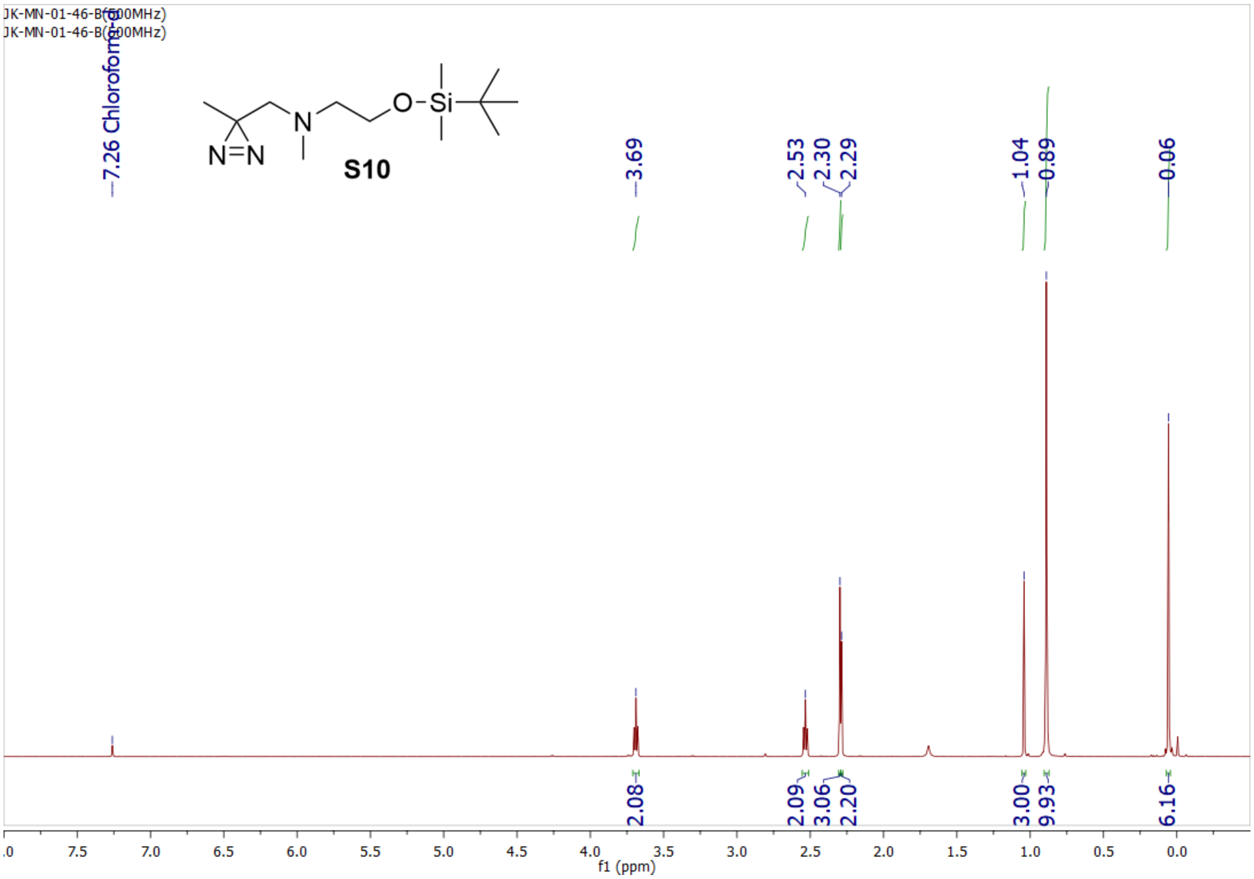
JK-MN-01-46-B(500MHz)
JK-MN-01-46-B(500MHz)
7.26 Chloroform-d
S10
3.69
2.53
2.30
2.29
1.04
0.89
0.06
2.08
2.09
3.06
2.20
3.00
9.93
6.16
f1 (ppm)

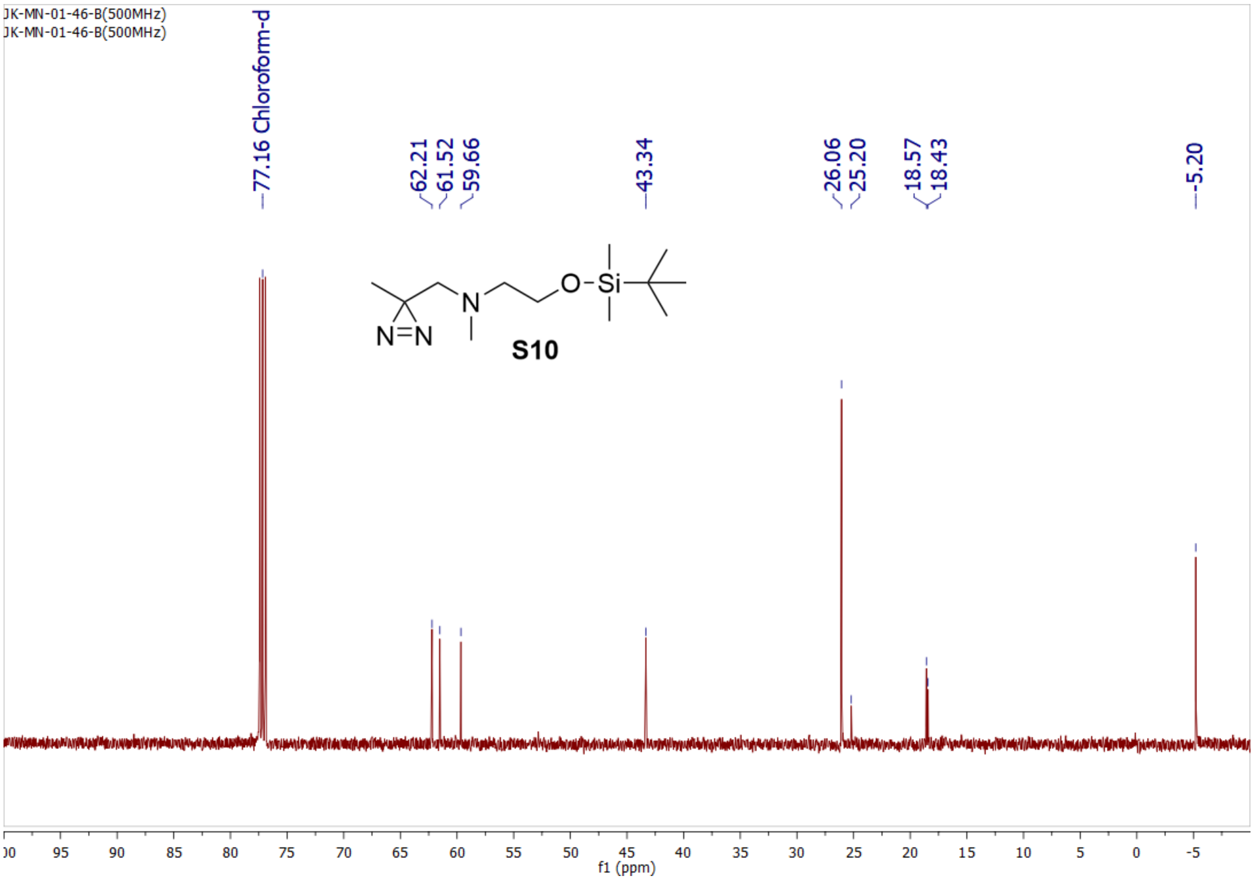
JK-MN-01-46-B(500MHz)
JK-MN-01-46-B(500MHz)
77.16 Chloroform-d
62.21
61.52
59.66
43.34
26.06
25.20
18.57
18.43
-5.20
S10
f1 (ppm)

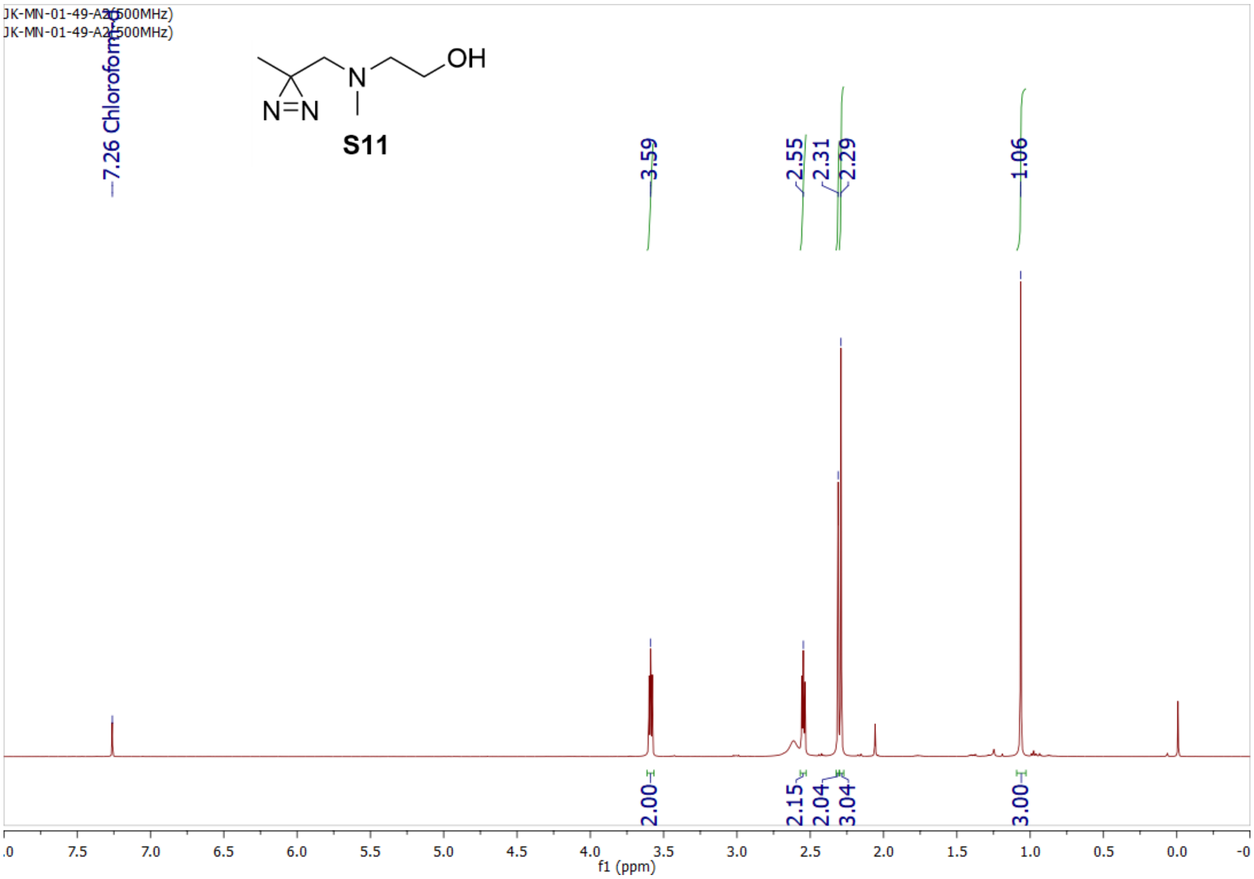
JK-MN-01-49-A2(500MHz)
JK-MN-01-49-A2(500MHz)
7.26 Chloroform-d
OH
N
N=N
S11
3.59
2.55
2.31
2.29
1.06
2.00
2.15
2.04
3.04
3.00
f1 (ppm)

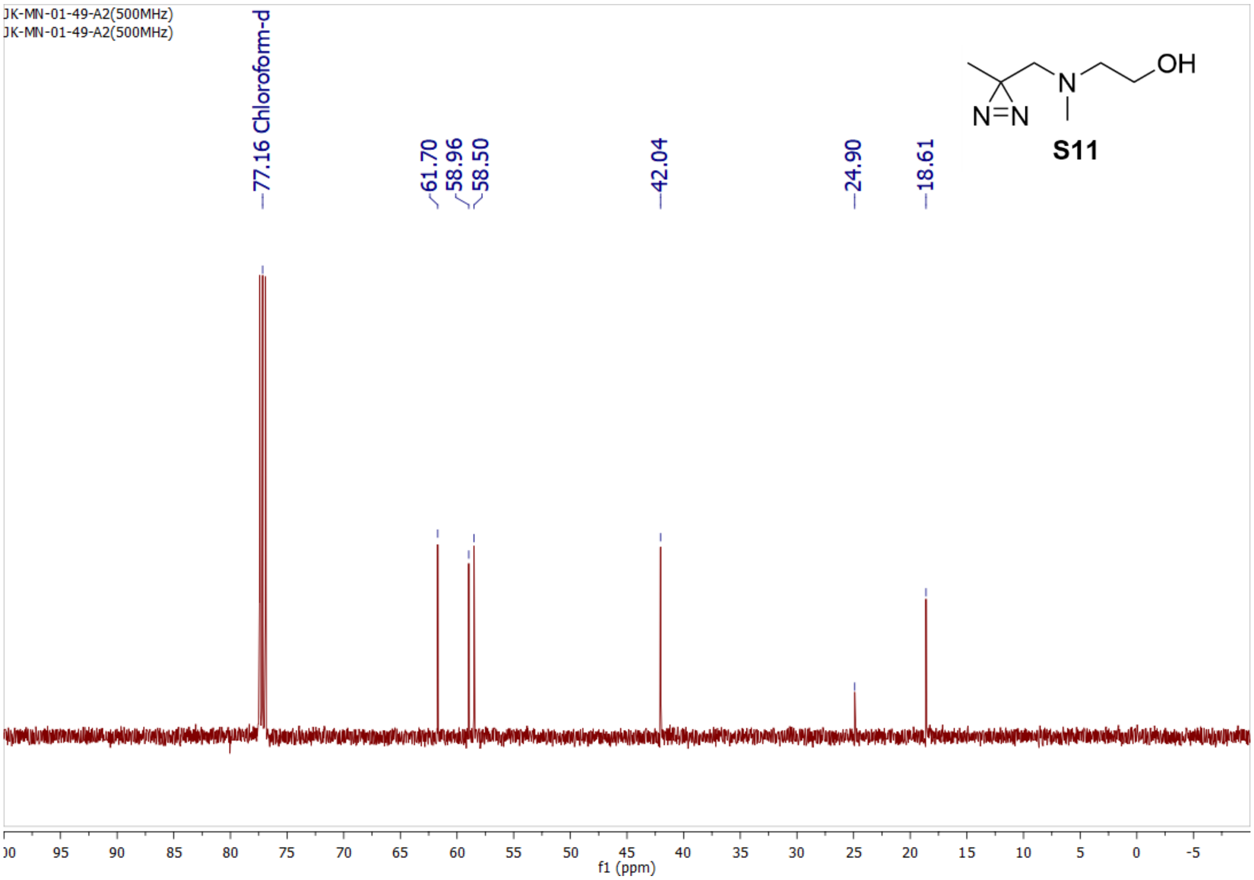
JK-MN-01-49-A2(500MHz)
JK-MN-01-49-A2(500MHz)
77.16 Chloroform-d
61.70
58.96
58.50
42.04
24.90
18.61
OH
N
N=N
S11
f1 (ppm)

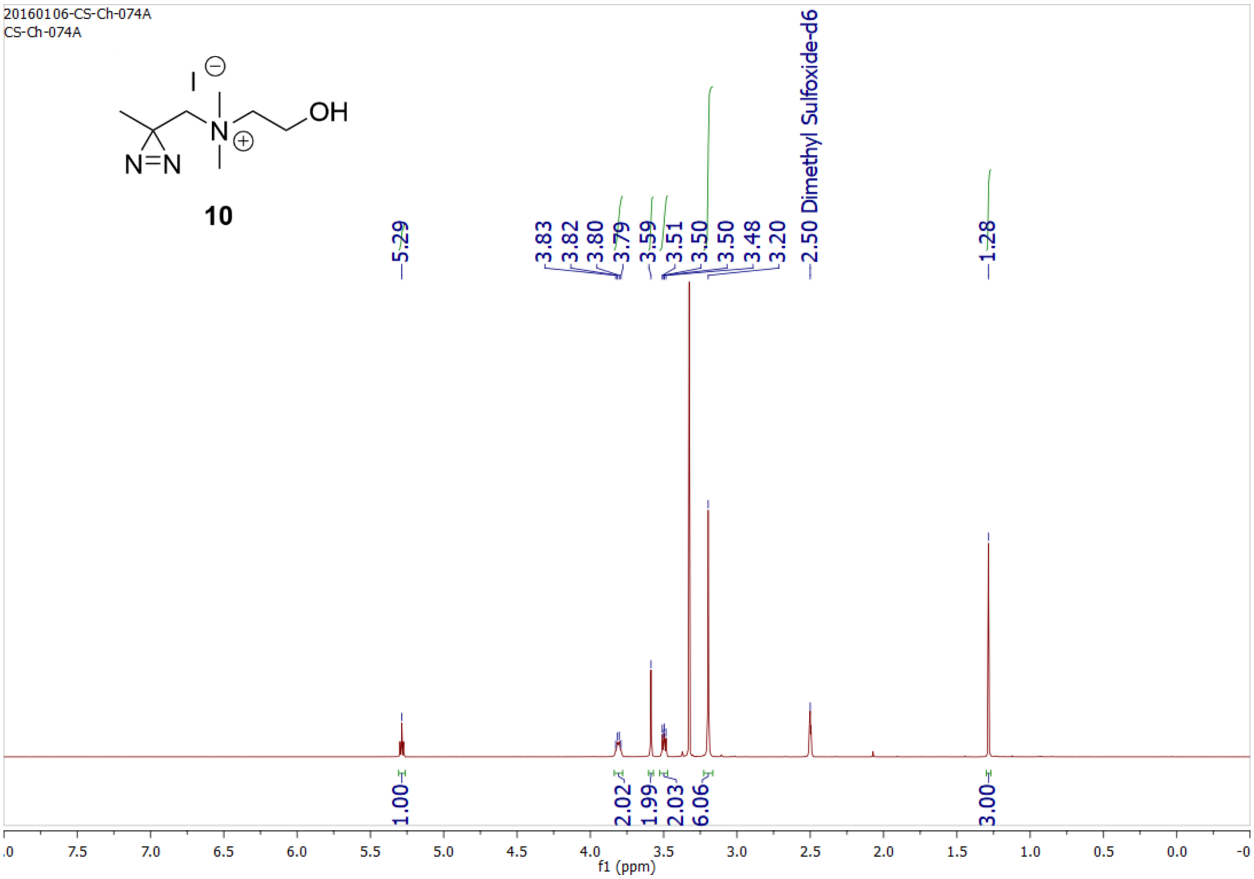
20160106-CS-Ch-074A
CS-Ch-074A
10
5.29
3.83
3.82
3.80
3.79
3.59
3.51
3.50
3.50
3.48
3.20
2.50 Dimethyl Sulfoxide-d6
1.28
1.00
2.02
1.99
2.03
6.06
3.00
f1 (ppm)

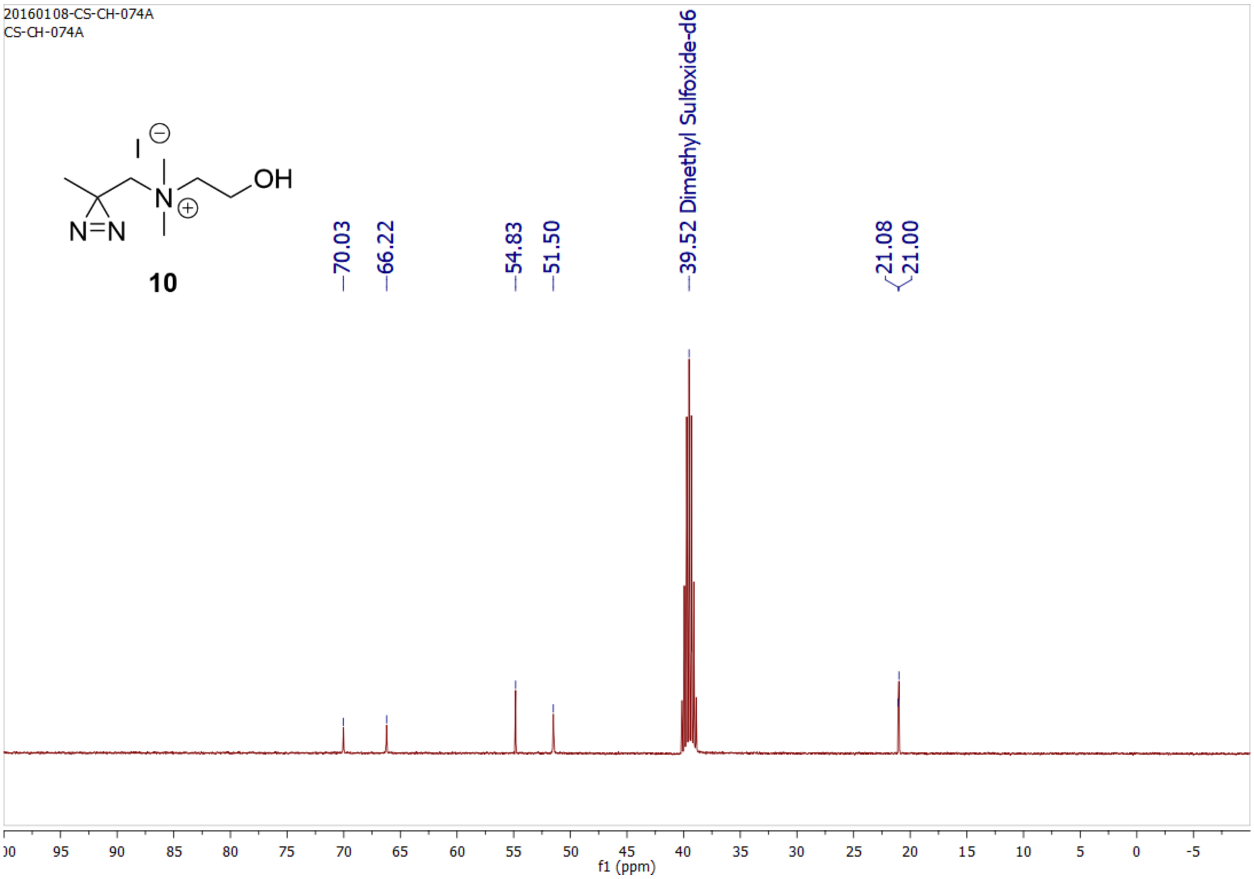
20160108-CS-CH-074A
CS-CH-074A
10
70.03
66.22
54.83
51.50
39.52 Dimethyl Sulfoxide-d6
21.08
21.00
f1 (ppm)

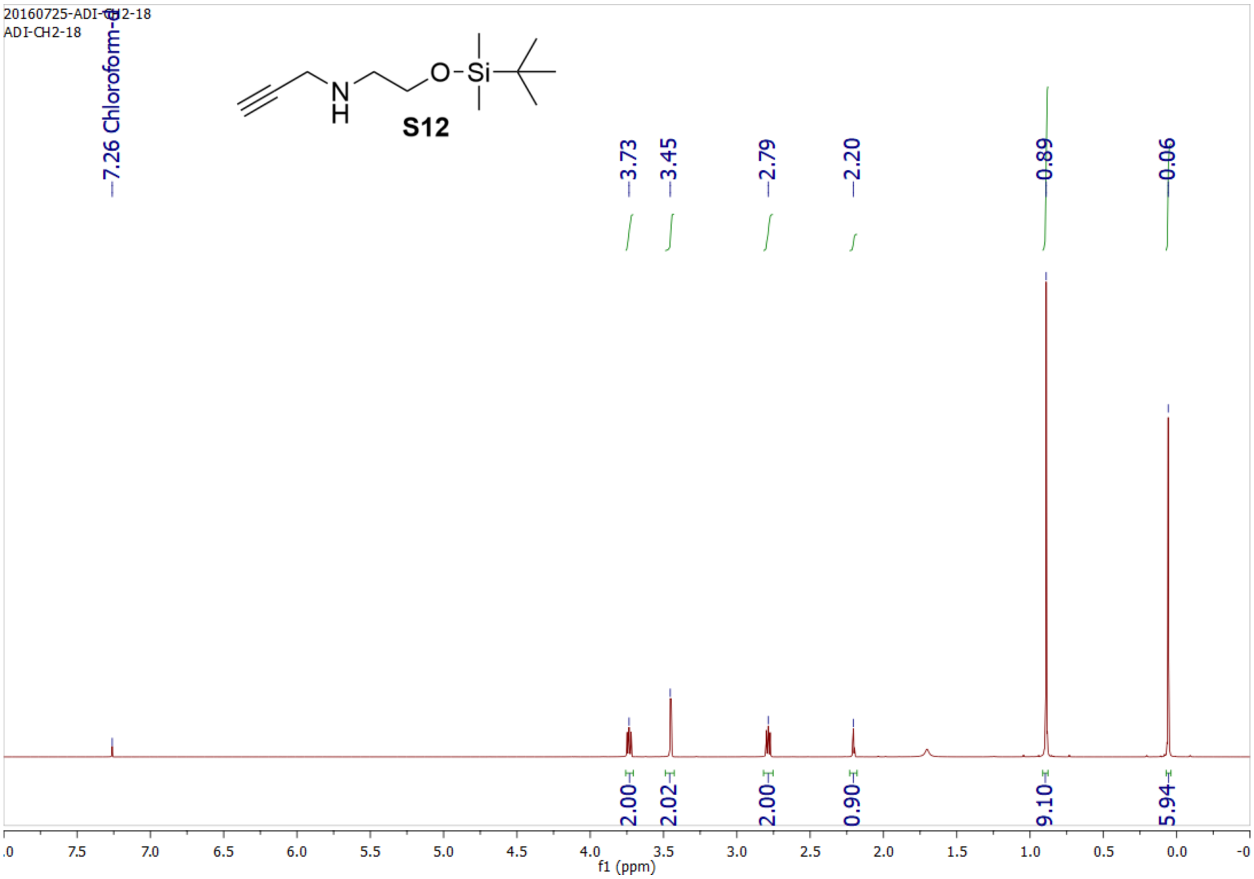
20160725-ADI-CH2-18
ADI-CH2-18
7.26 Chloroform-d
S12
3.73
3.45
2.79
2.20
0.89
0.06
2.00
2.02
2.00
0.90
9.10
5.94
f1 (ppm)

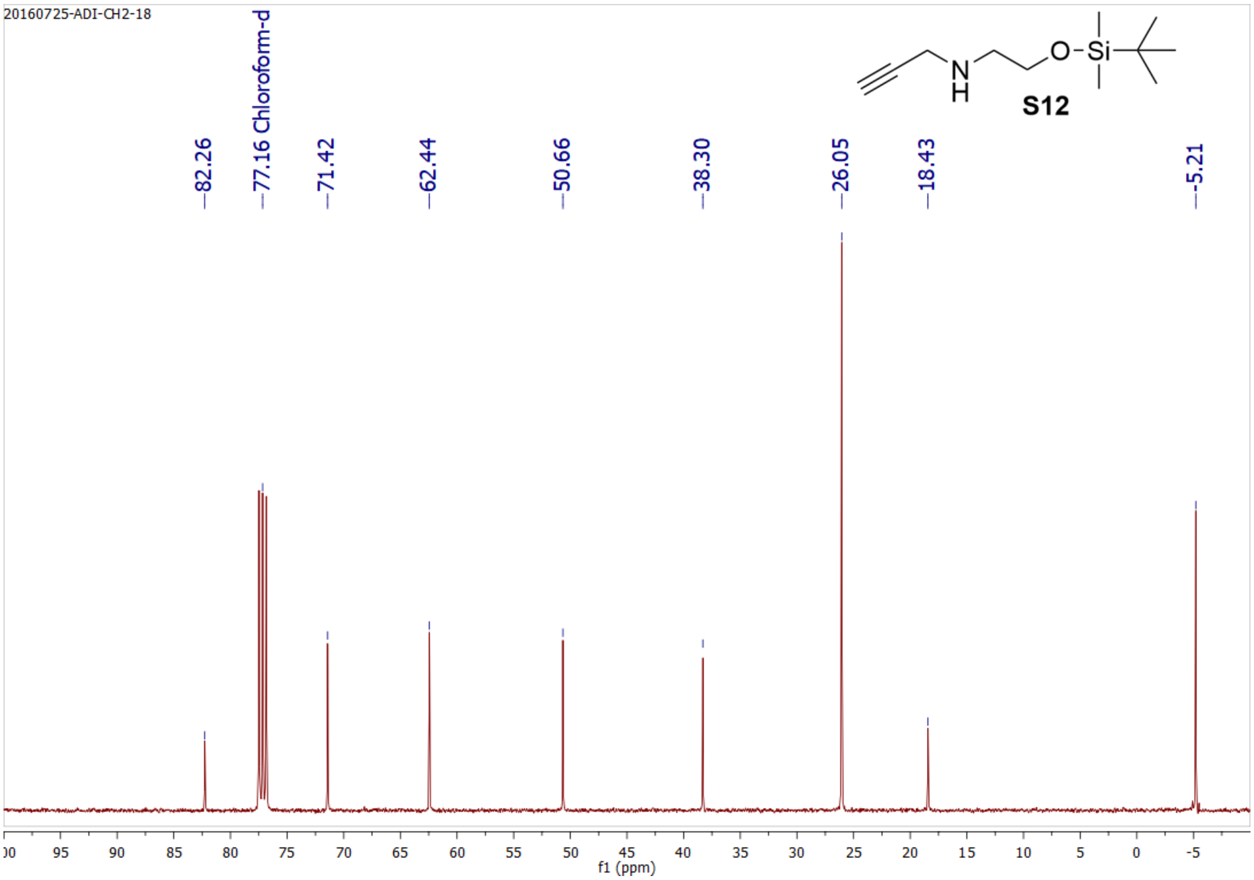
20160725-ADI-CH2-18
S12
82.26
77.16 Chloroform-d
71.42
62.44
50.66
38.30
26.05
18.43
-5.21
f1 (ppm)

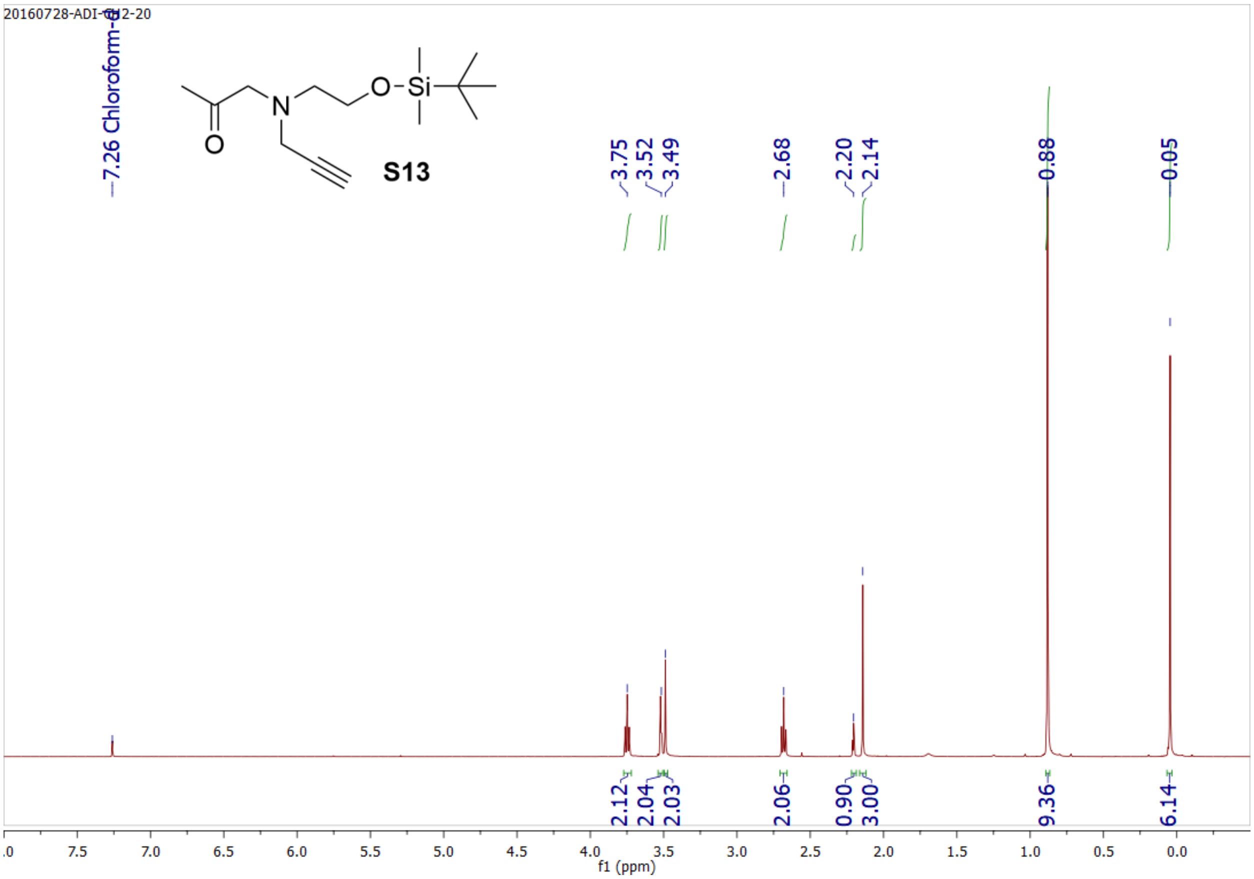

20160728-ADI-CH2-20
7.26 Chloroform-d
S13
3.75
3.52
3.49
2.68
2.20
2.14
0.88
0.05
2.12
2.04
2.03
2.06
0.90
3.00
9.36
6.14
f1 (ppm)


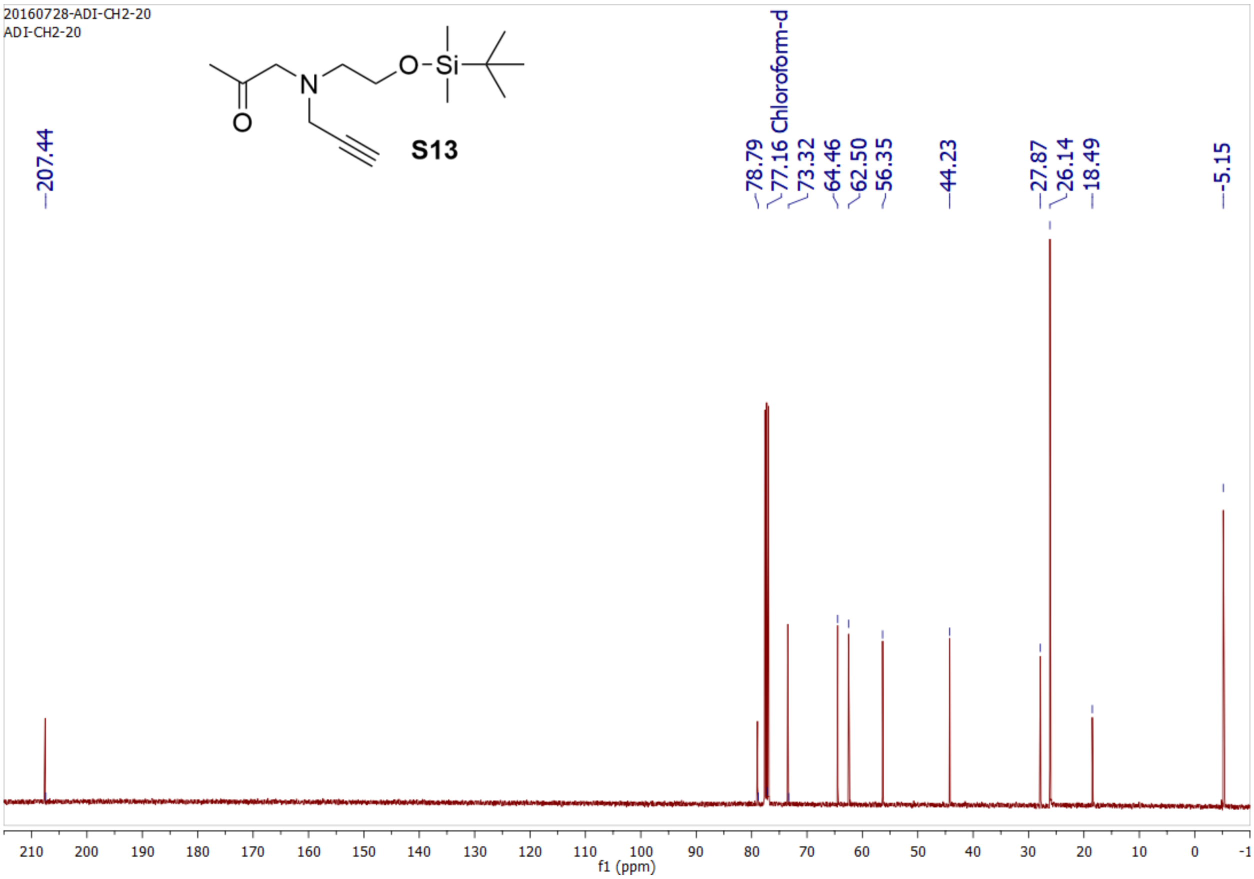

20160728-ADI-CH2-20
ADI-CH2-20
S13
207.44
78.79
77.16 Chloroform-d
73.32
64.46
62.50
56.35
44.23
27.87
26.14
18.49
-5.15
f1 (ppm)

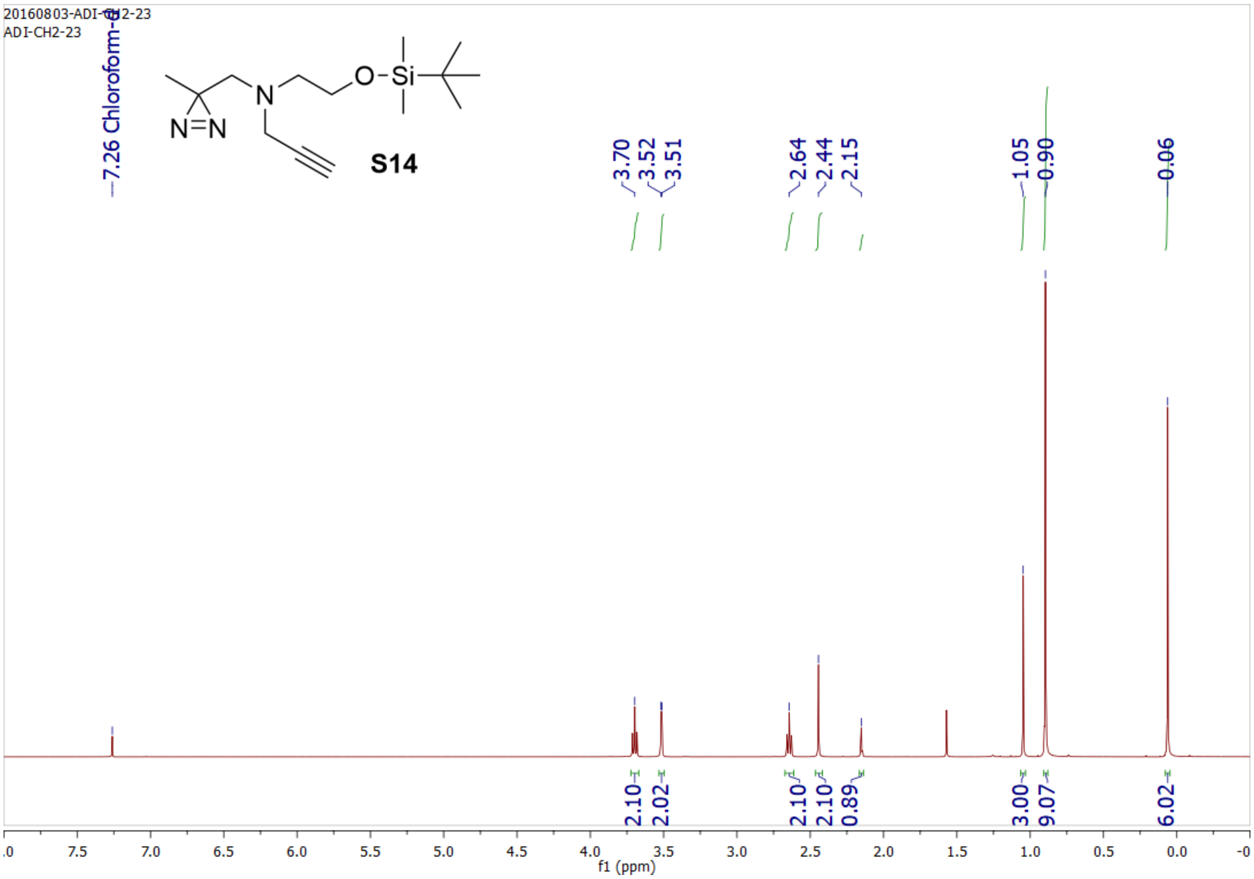
20160803-ADI-CH2-23
ADI-CH2-23
7.26 Chloroform-d
S14
3.70
3.52
3.51
2.64
2.44
2.15
1.05
0.90
0.06
2.10
2.02
2.10
2.10
0.89
3.00
9.07
6.02
f1 (ppm)

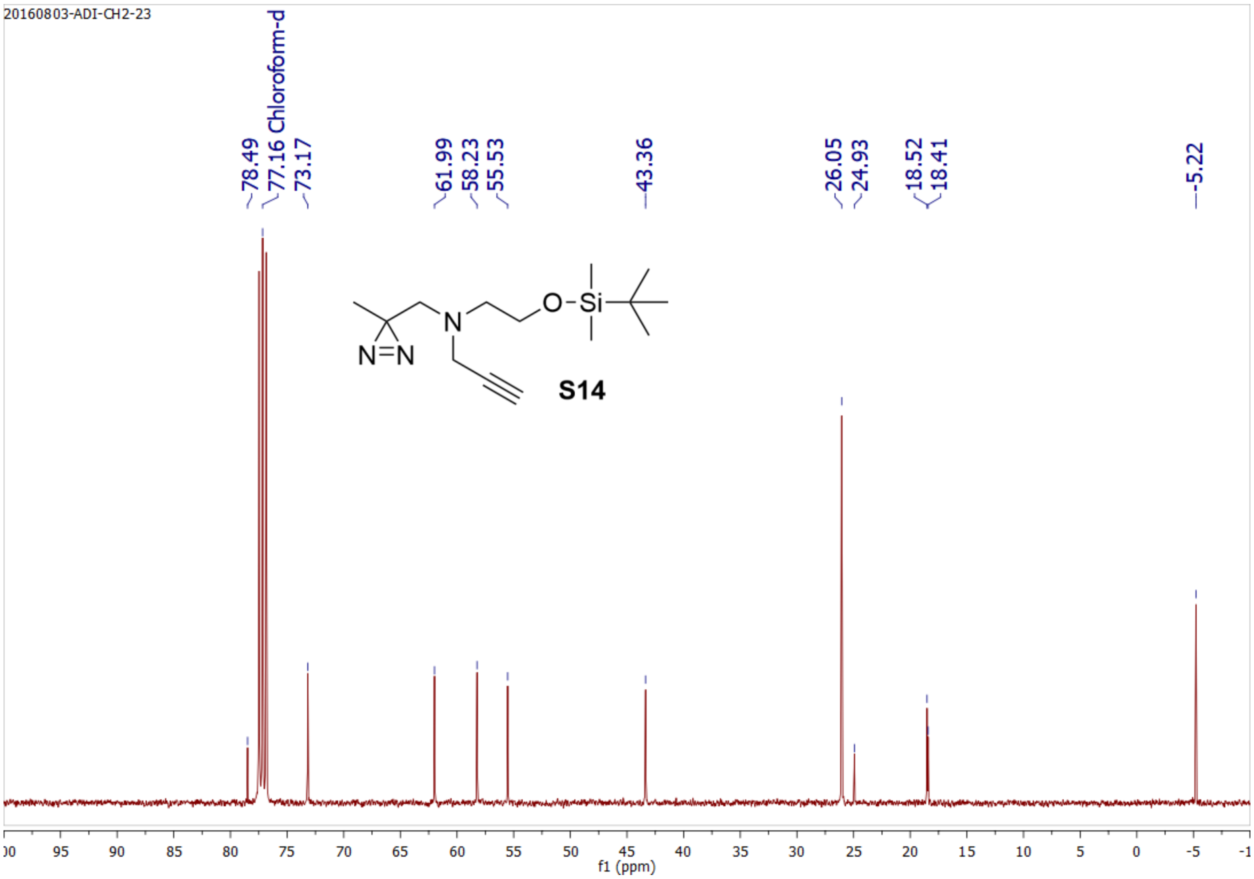
20160803-ADI-CH2-23
78.49
77.16 Chloroform-d
73.17
61.99
58.23
55.53
43.36
26.05
24.93
18.52
18.41
-5.22
S14
f1 (ppm)

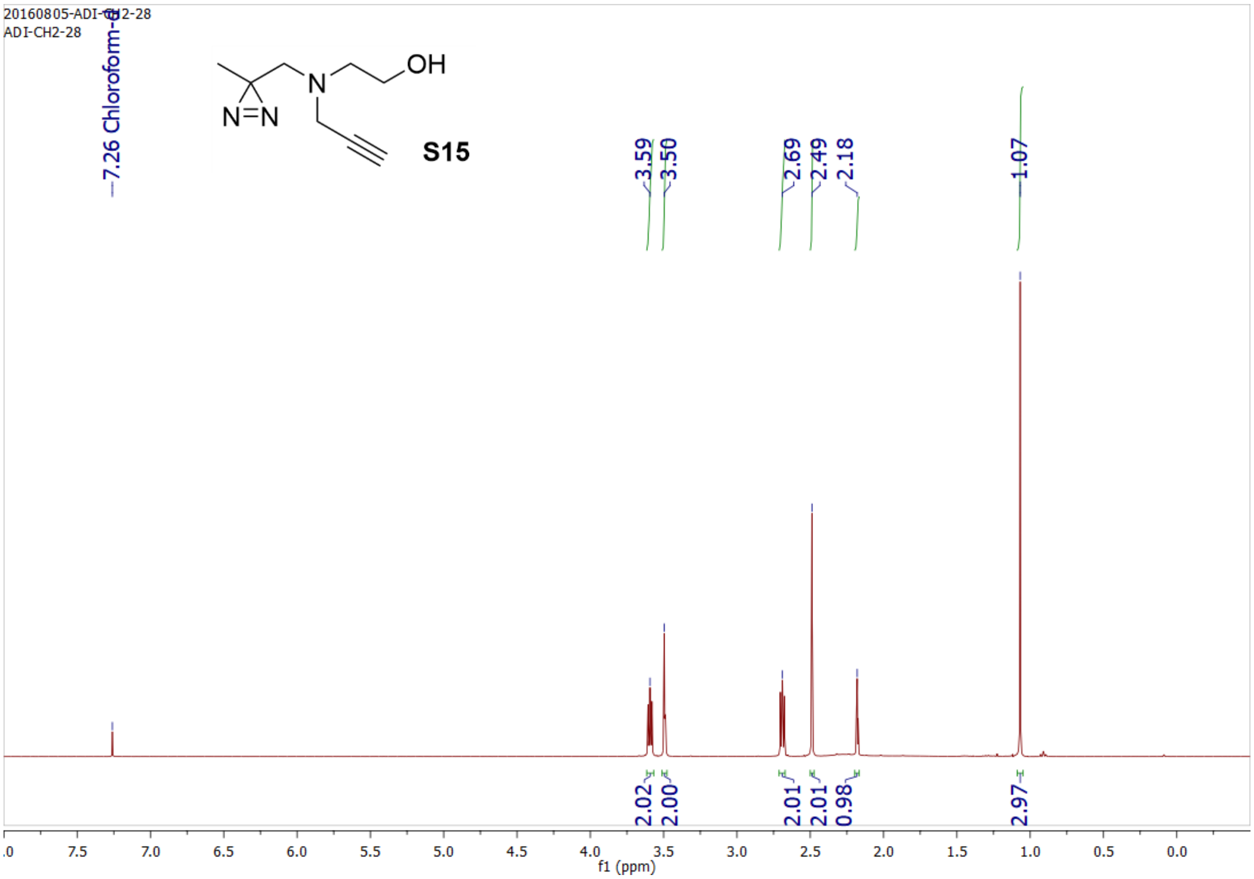
20160805-ADI-CH2-28
ADI-CH2-28
7.26 Chloroform-d
OH
N
N=N
S15
3.59
3.50
2.69
2.49
2.18
1.07
2.02
2.00
2.01
2.01
0.98
2.97
f1 (ppm)

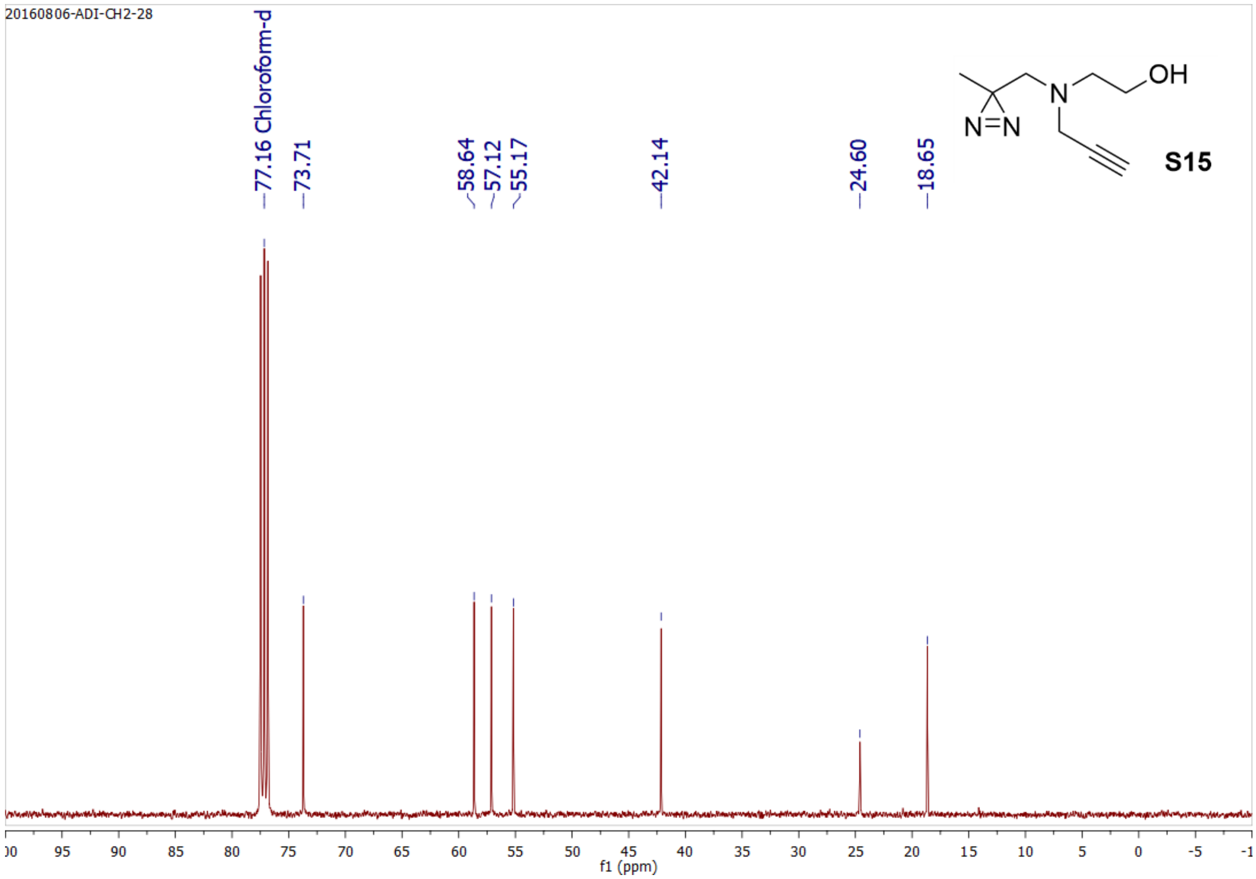
20160806-ADI-CH2-28
77.16 Chloroform-d
73.71
58.64
57.12
55.17
42.14
24.60
18.65
OH
N
N=N
S15
f1 (ppm)

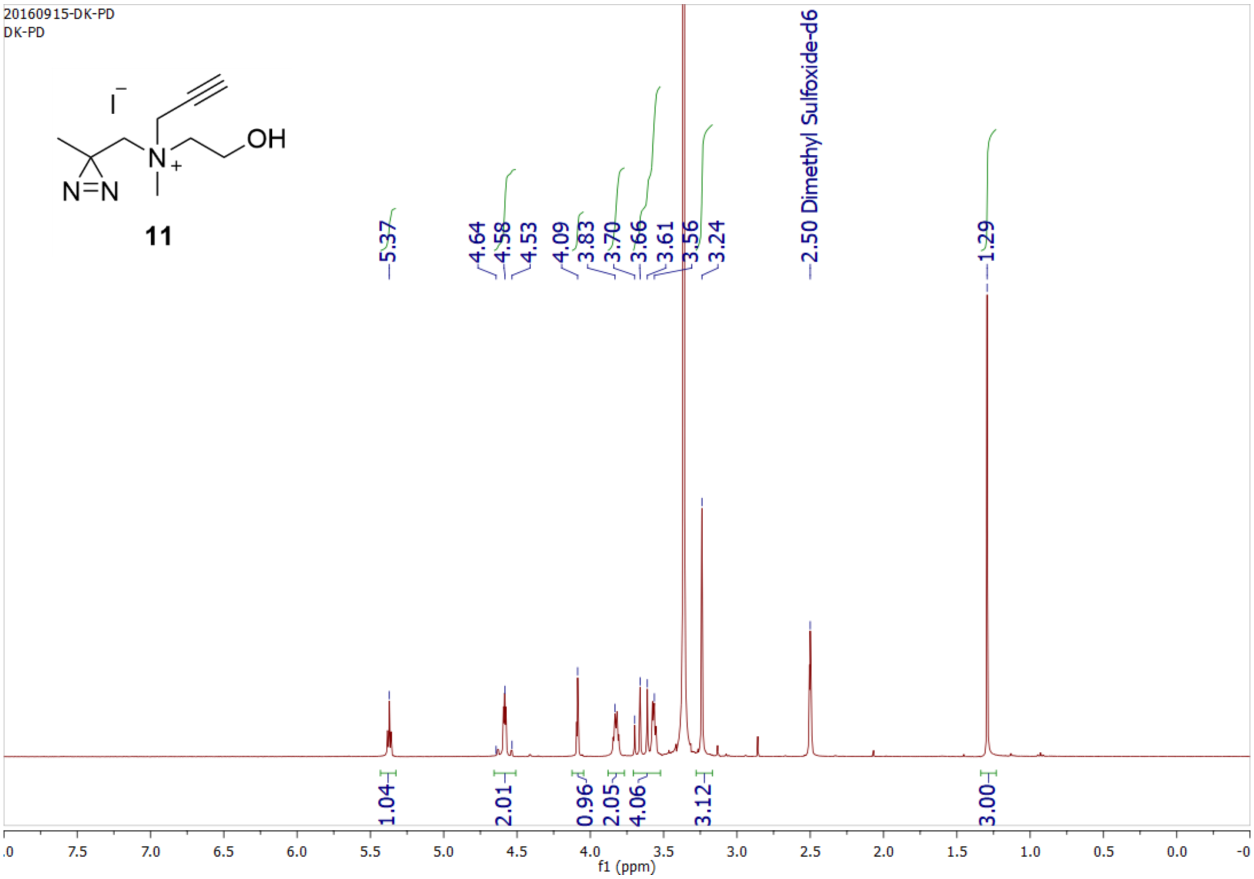

20160915-DK-PD
DK-PD
OH
N
I
N=N
11
5.37
4.64
4.58
4.53
4.09
3.83
3.70
3.66
3.61
3.56
3.24
2.50 Dimethyl Sulfoxide-d6
1.29
1.04
2.01
0.96
2.05
4.06
3.12
3.00
f1 (ppm)


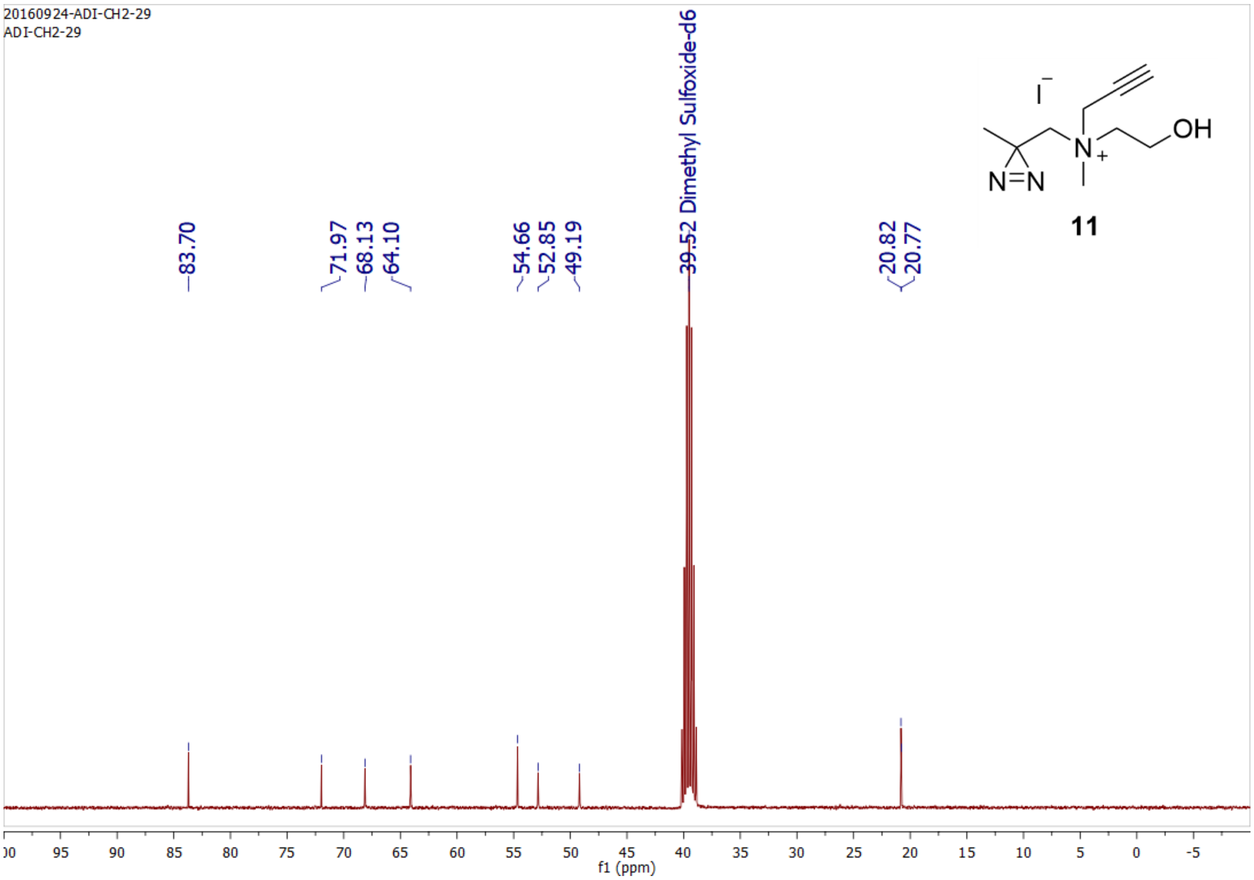

20160924-ADI-CH2-29
ADI-CH2-29
83.70
71.97
68.13
64.10
54.66
52.85
49.19
39.52 Dimethyl Sulfoxide-d6
20.82
20.77
OH
N
I
N=N
11
f1 (ppm)

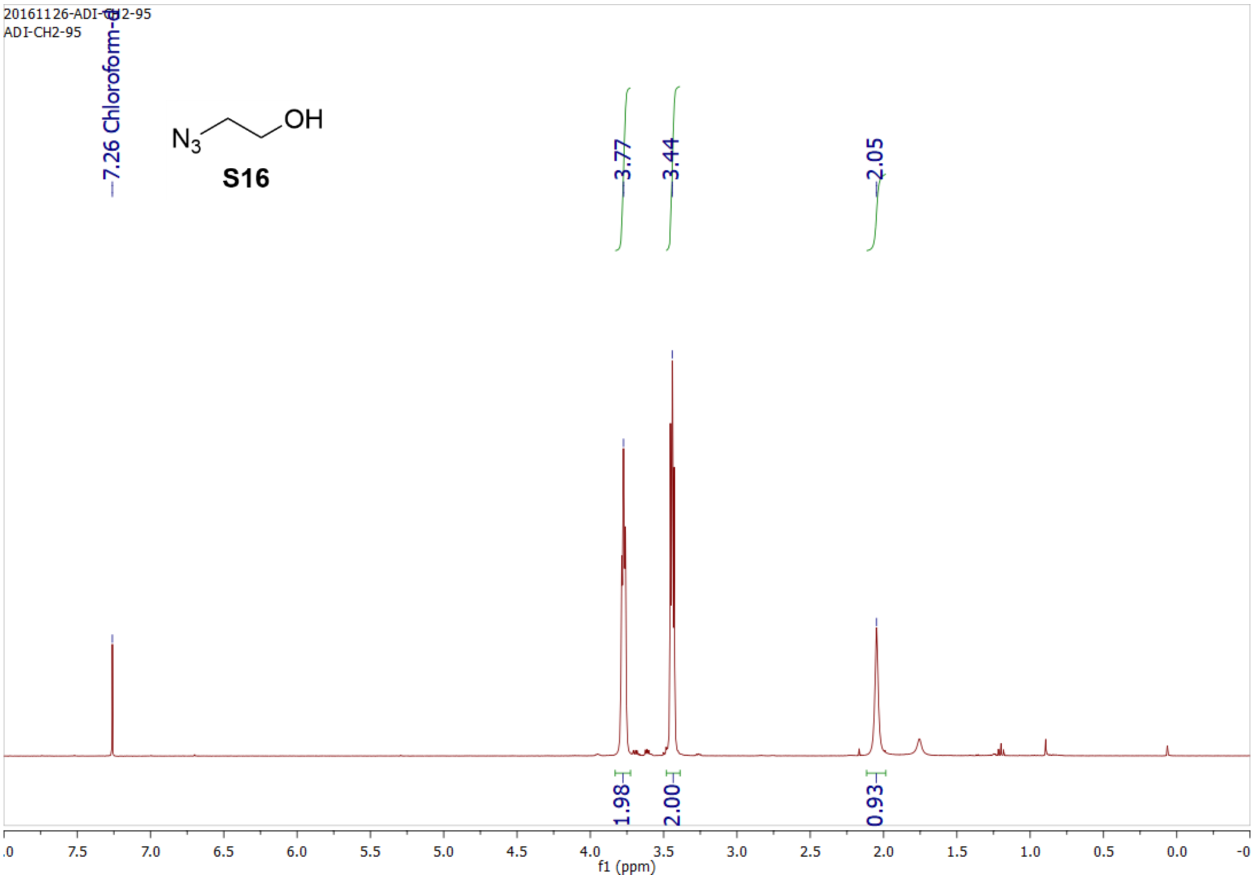
20161126-ADI-CH2-95
ADI-CH2-95
7.26 Chloroform-d
N3
OH
S16
3.77
3.44
2.05
1.98
2.00
0.93
f1 (ppm)

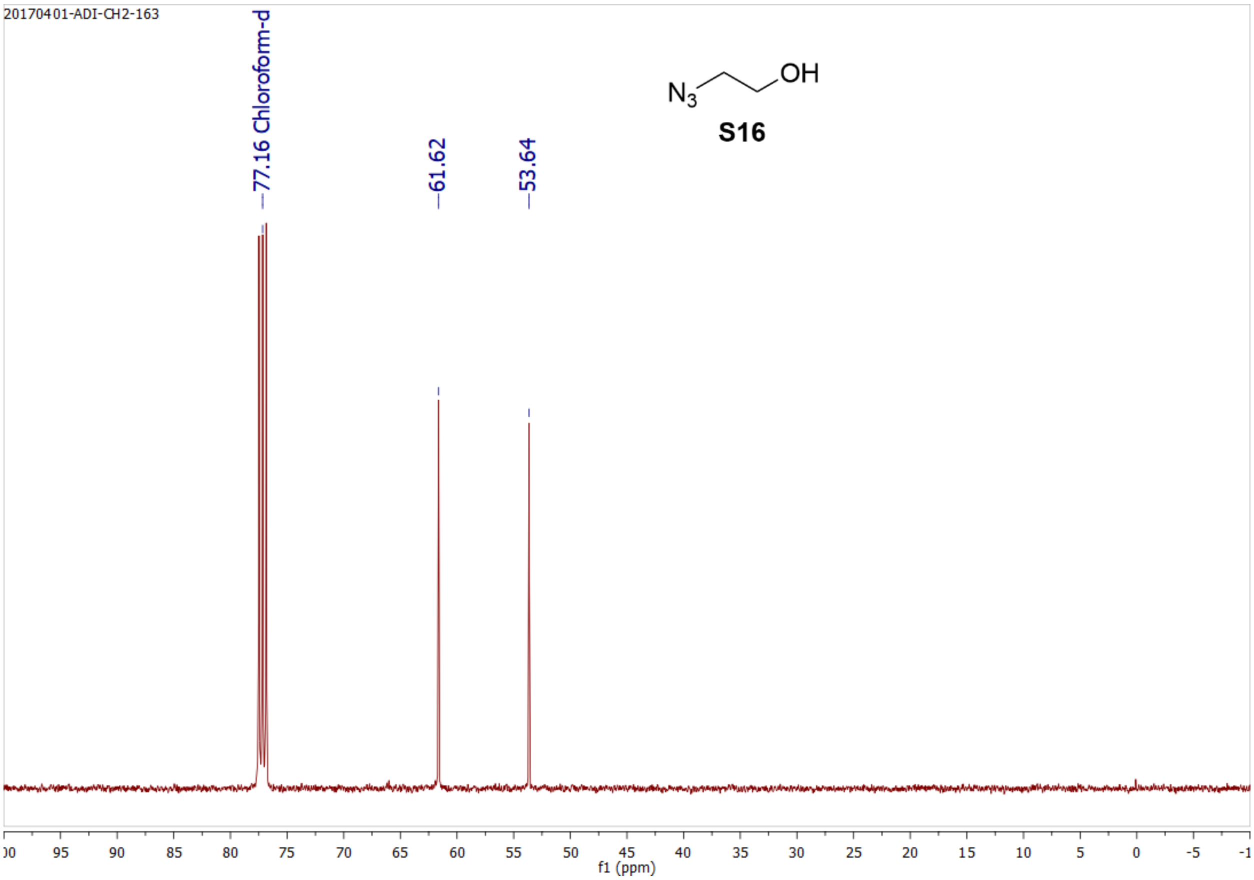
20170401-ADI-CH2-163
77.16 Chloroform-d
61.62
53.64
N3
OH
S16
f1 (ppm)

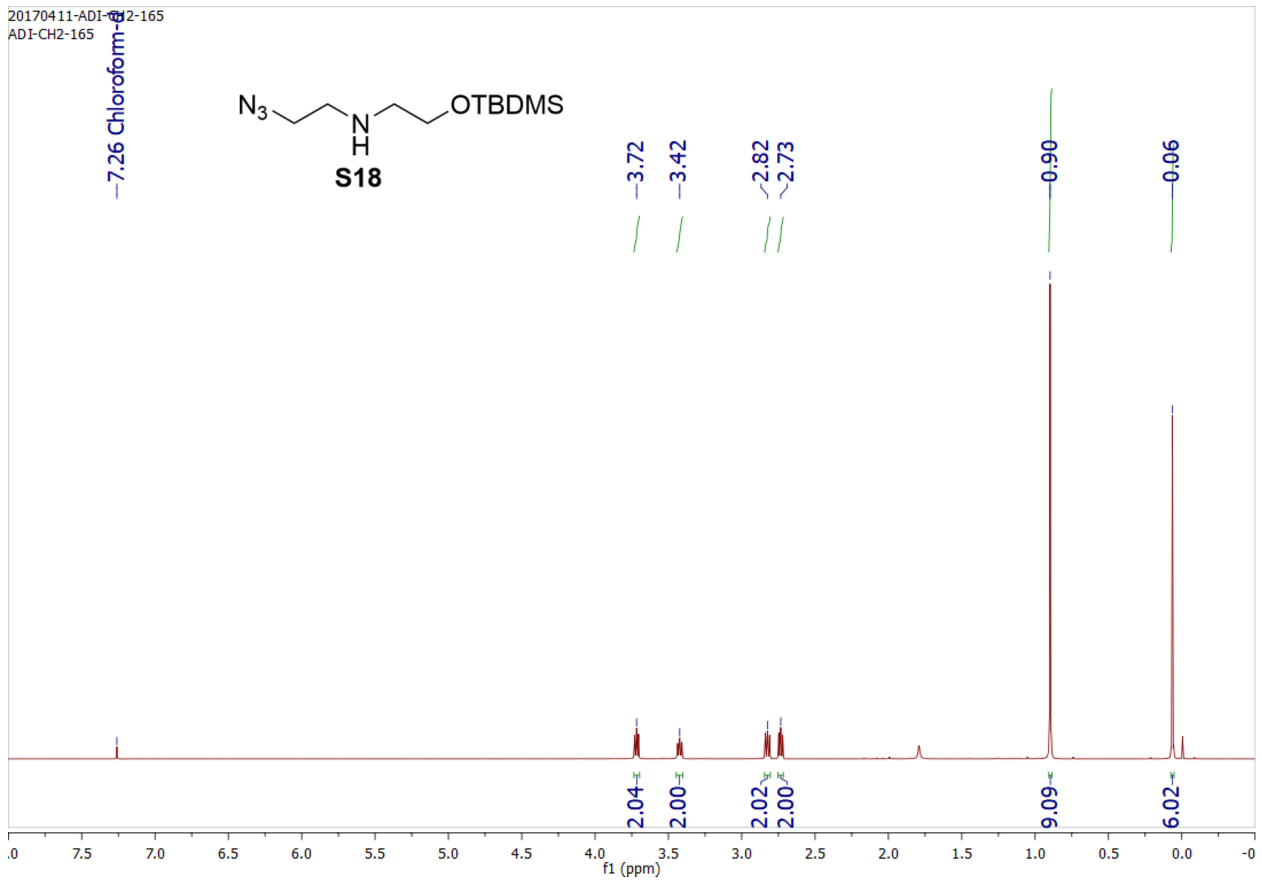
20170411-ADI-CH2-165
ADI-CH2-165
7.26 Chloroform-d
N3 N H OTBDMS
S18
3.72
3.42
2.82
2.73
0.90
0.06
2.04
2.00
2.02
2.00
9.09
6.02
f1 (ppm)

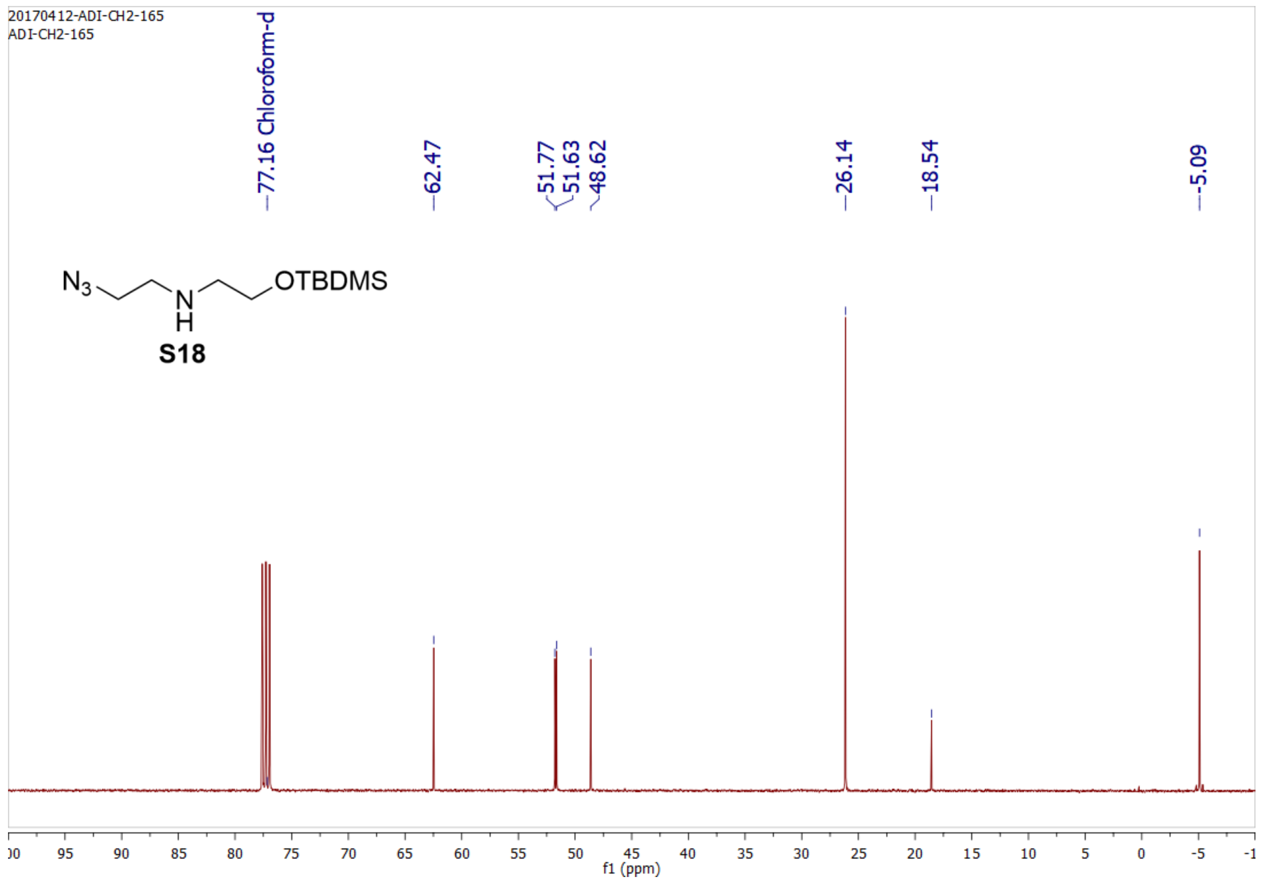
20170412-ADI-CH2-165
ADI-CH2-165
77.16 Chloroform-d
62.47
51.77
51.63
48.62
26.14
18.54
-5.09
N3 N H OTBDMS
S18
f1 (ppm)

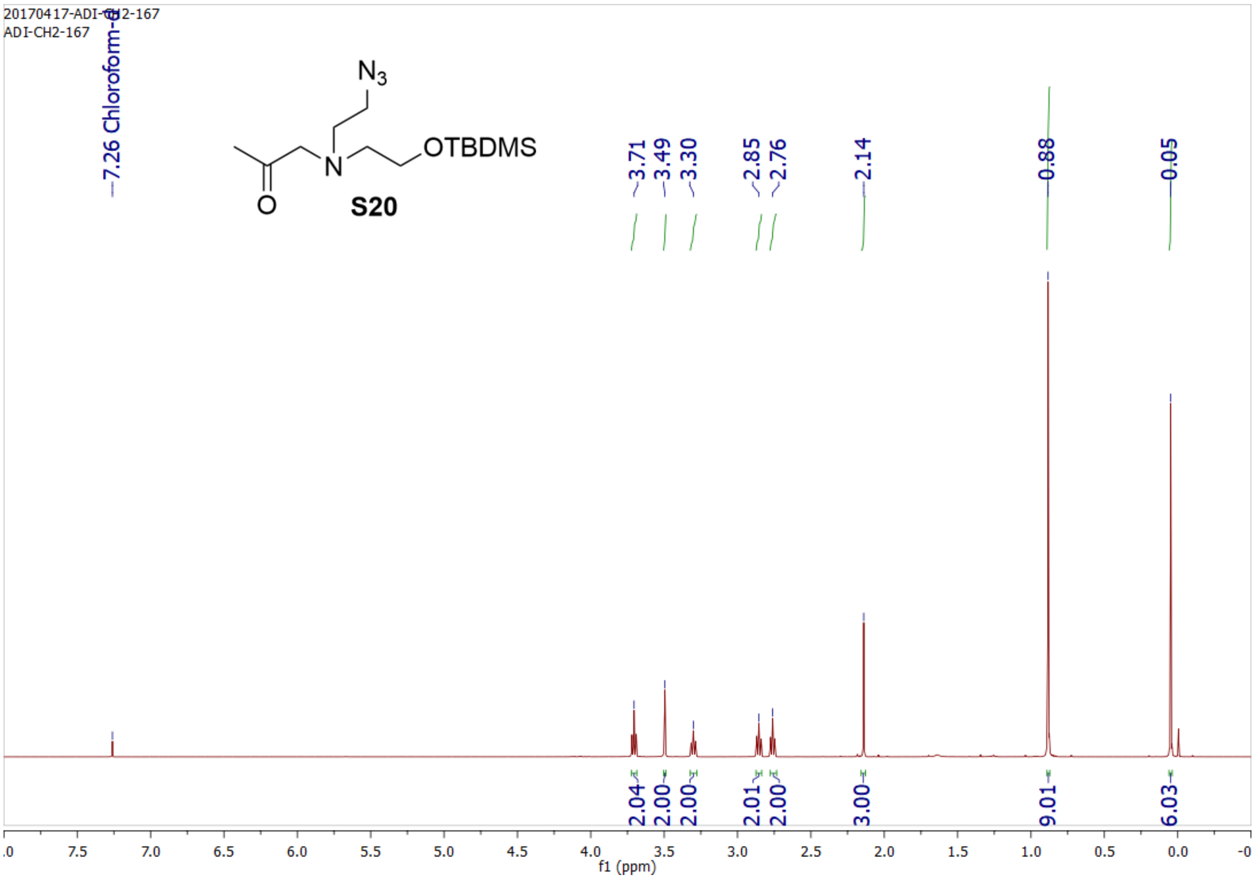
20170417-ADI-CH2-167
ADI-CH2-167
7.26 Chloroform-d
N3
OTBDMS
O
N
S20
3.71
3.49
3.30
2.85
2.76
2.14
0.88
0.05
2.04
2.00
2.00
2.01
2.00
3.00
9.01
6.03
f1 (ppm)

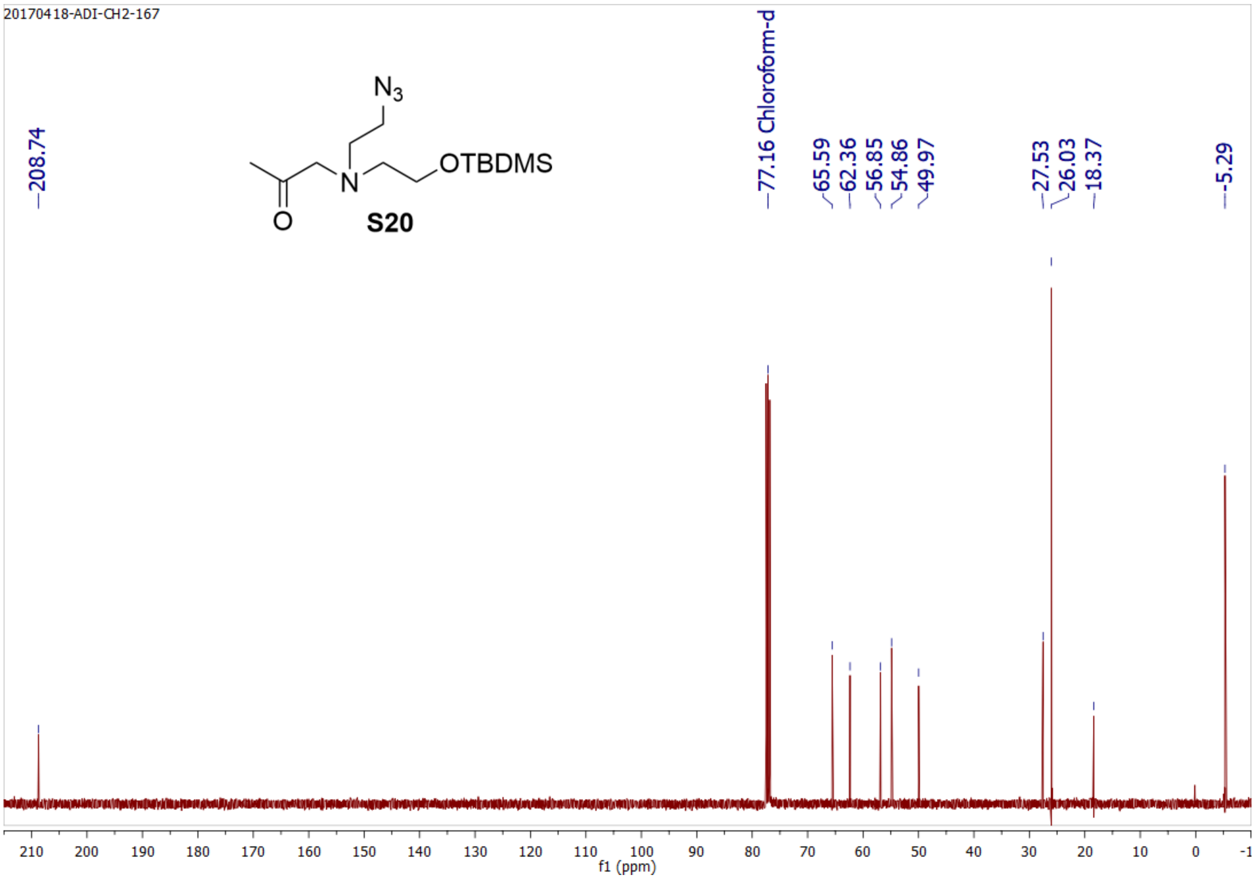
20170418-ADI-CH2-167
208.74
N3
OTBDMS
O
N
S20
77.16 Chloroform-d
65.59
62.36
56.85
54.86
49.97
27.53
26.03
18.37
-5.29
f1 (ppm)

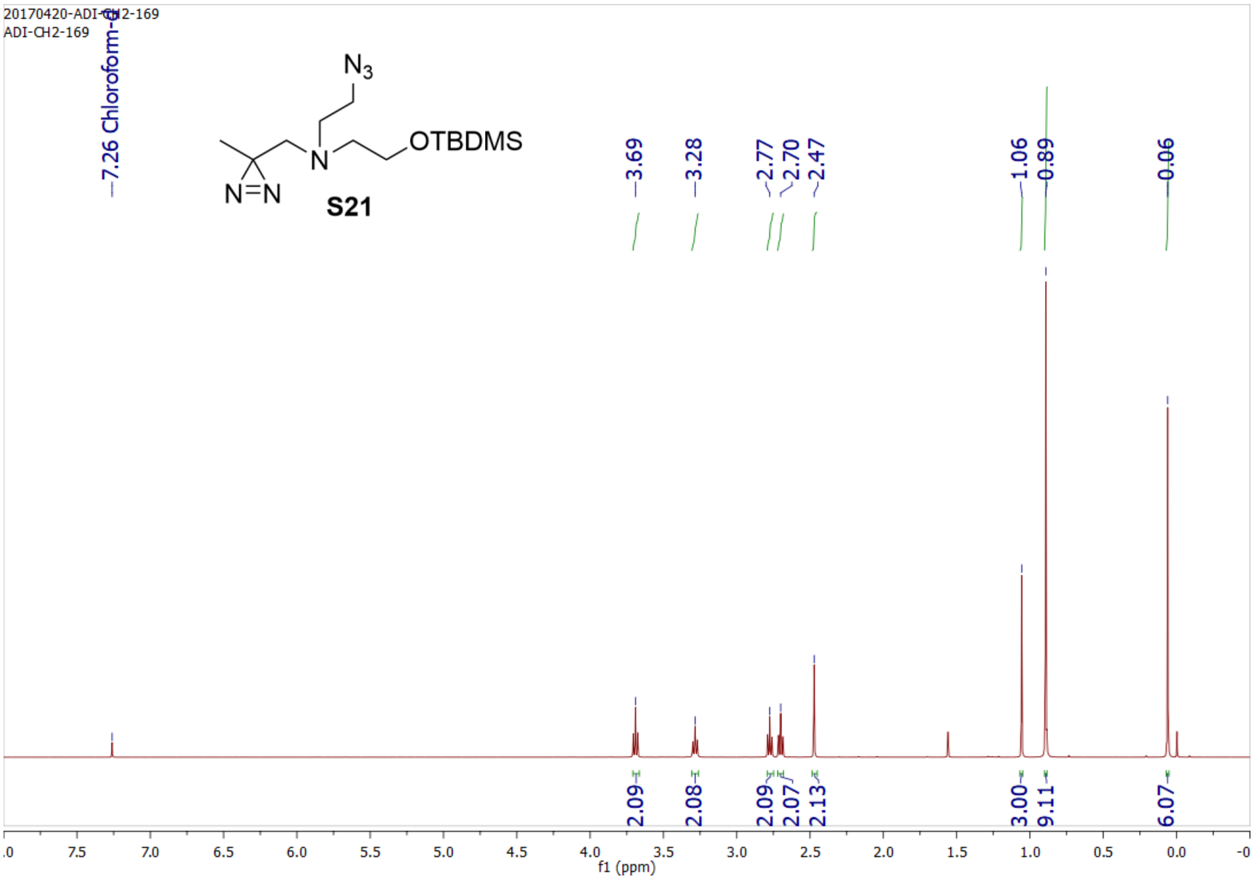
20170420-ADI-CH2-169
ADI-CH2-169
7.26 Chloroform-d
N3
OTBDMS
N
N=N
S21
3.69
3.28
2.77
2.70
2.47
1.06
0.89
0.06
2.09
2.08
2.09
2.07
2.13
3.00
9.11
6.07
f1 (ppm)

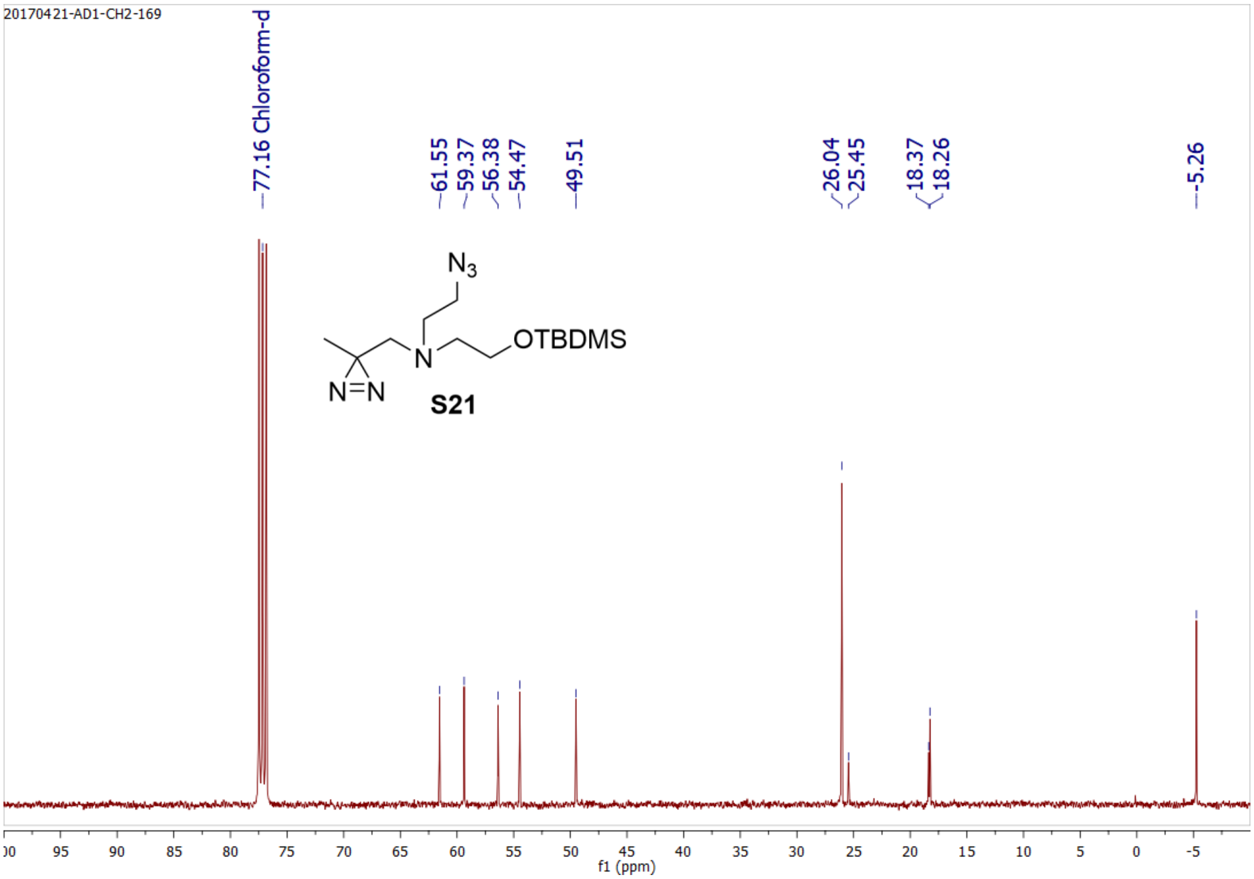
20170421-AD1-CH2-169
77.16 Chloroform-d
61.55
59.37
56.38
54.47
49.51
26.04
25.45
18.37
18.26
-5.26
N3
OTBDMS
N
N=N
S21
f1 (ppm)

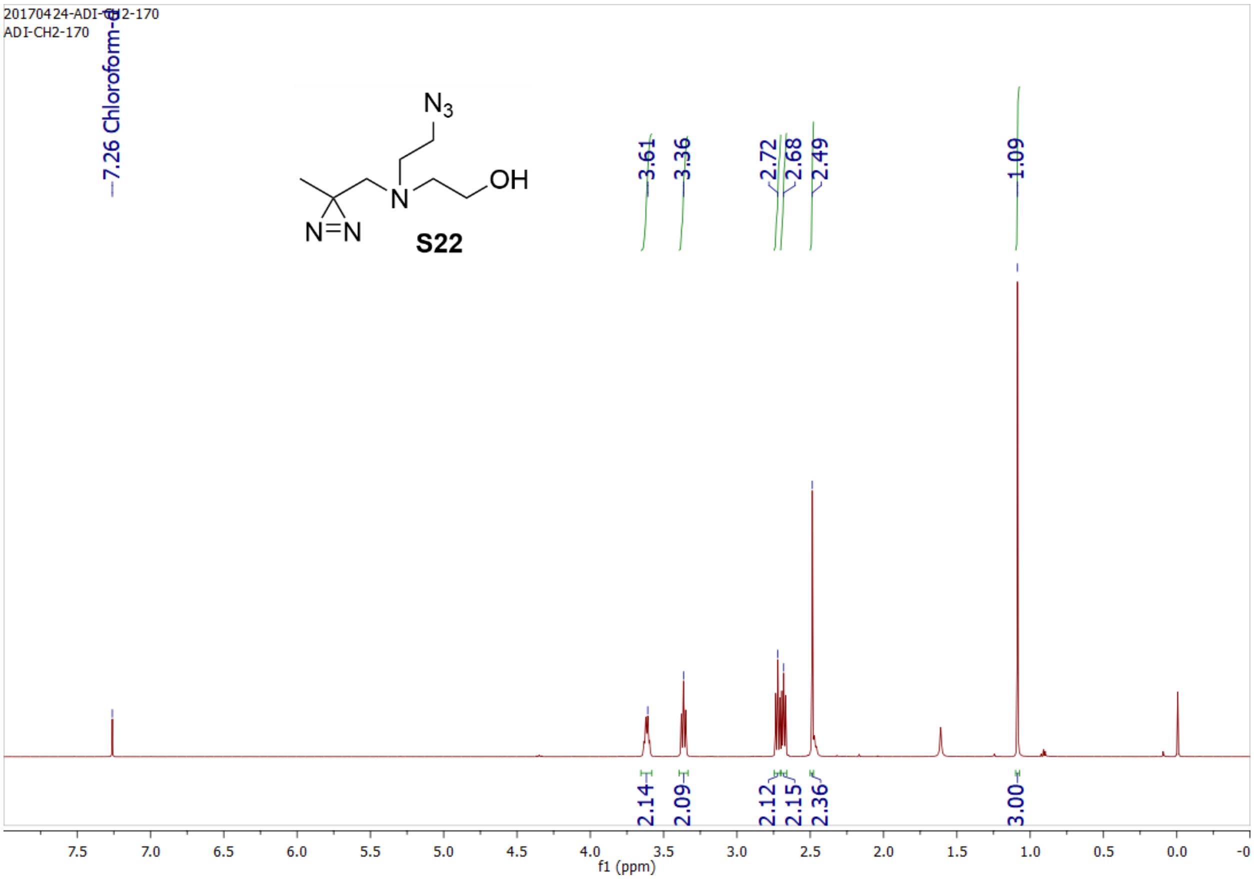
20170424-ADI-CH2-170
ADI-CH2-170
7.26 Chloroform-d
N3
OH
N
N=N
S22
3.61
3.36
2.72
2.68
2.49
1.09
2.14
2.09
2.12
2.15
2.36
3.00
f1 (ppm)

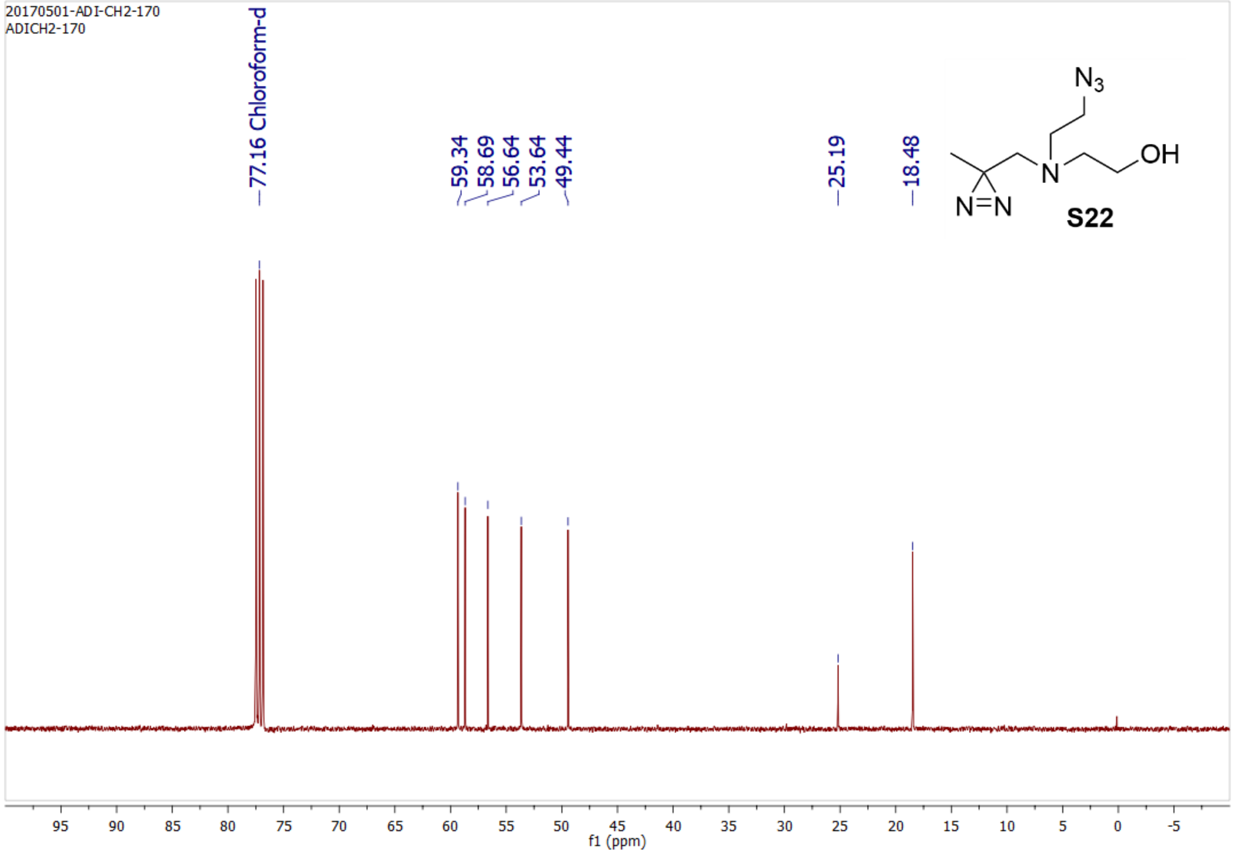
20170501-ADI-CH2-170
ADICH2-170
77.16 Chloroform-d
59.34
58.69
56.64
53.64
49.44
25.19
18.48
N3
OH
N
N=N
S22
f1 (ppm)

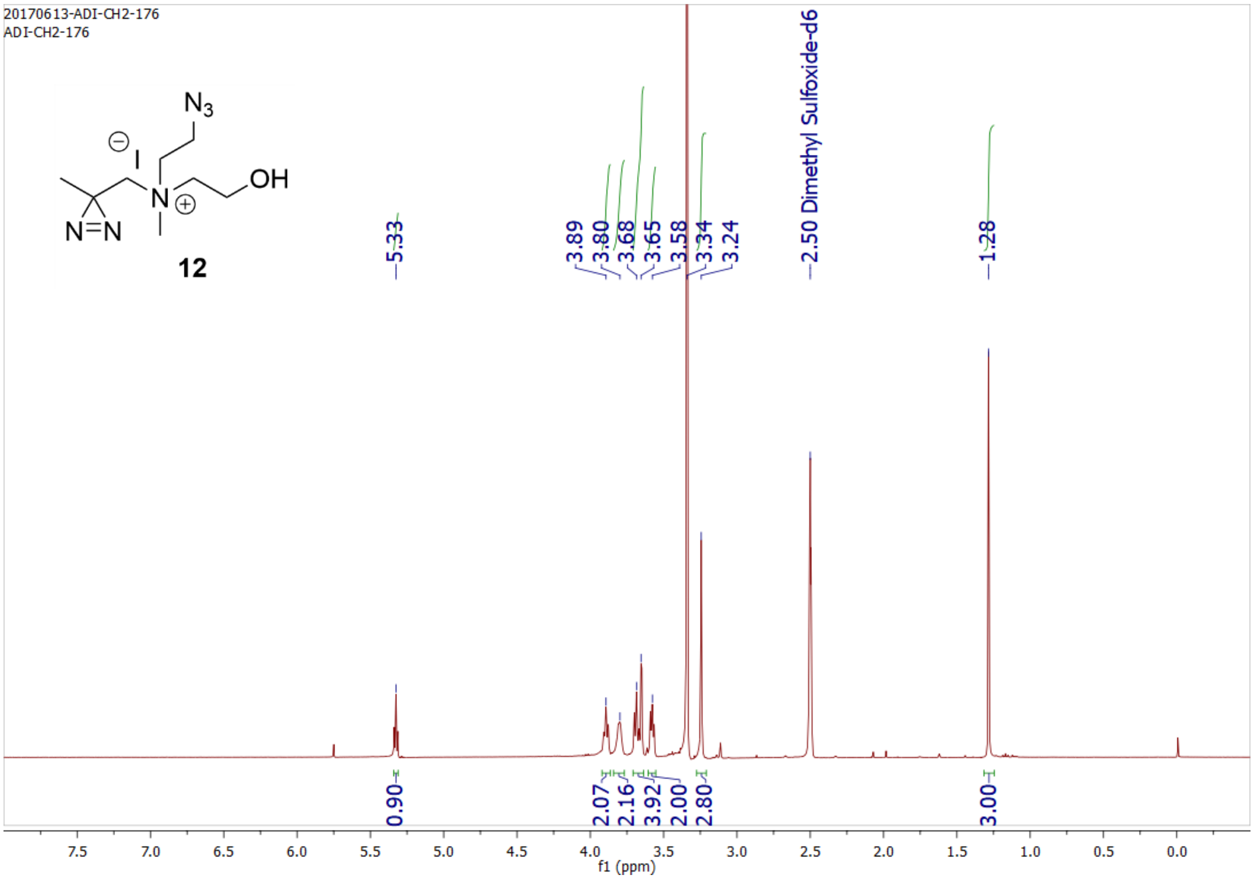

20170613-ADI-CH2-176
ADI-CH2-176
12
5.33
3.89
3.80
3.68
3.65
3.58
3.34
3.24
2.50 Dimethyl Sulfoxide-d6
1.28
0.90
2.07
2.16
3.92
2.00
2.80
3.00
f1 (ppm)


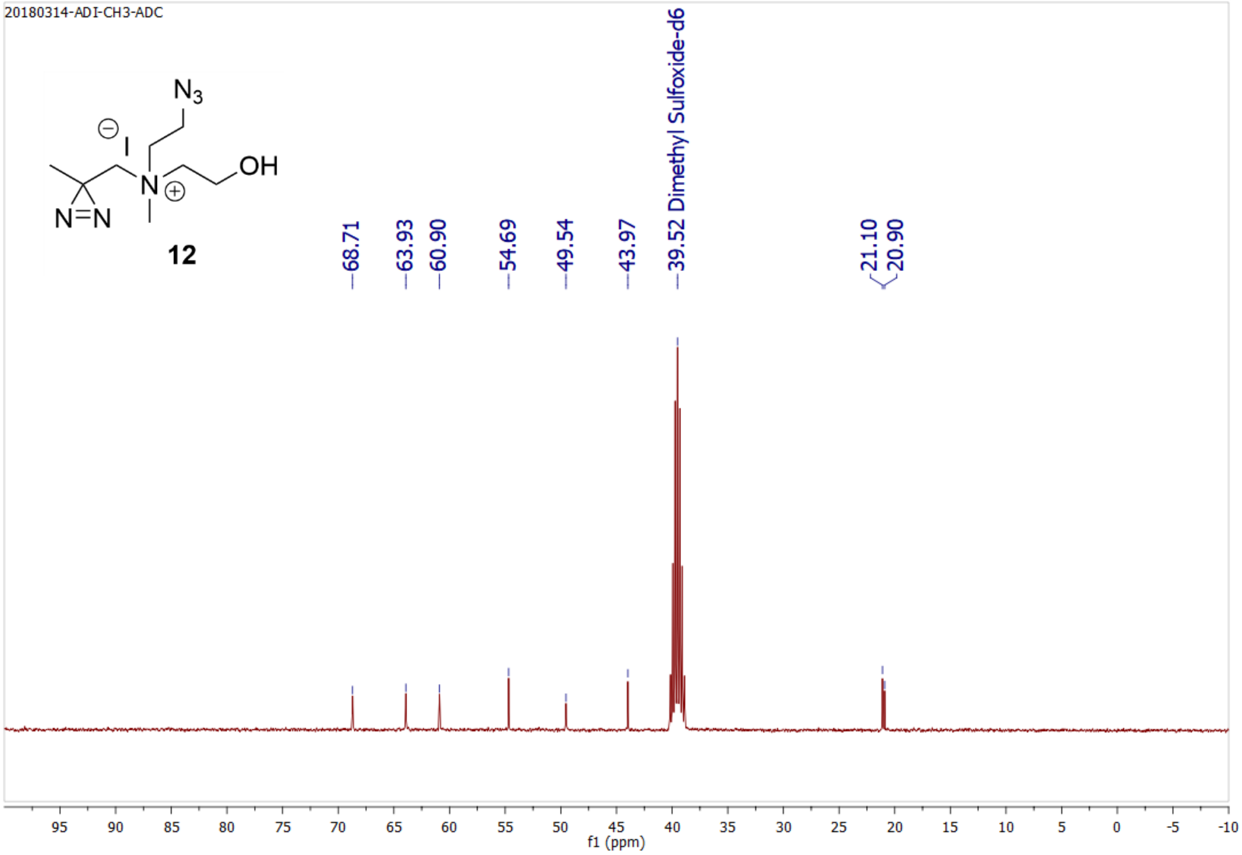

20180314-ADI-CH3-ADC
12
68.71
63.93
60.90
54.69
49.54
43.97
39.52 Dimethyl Sulfoxide-d6
21.10
20.90
f1 (ppm)

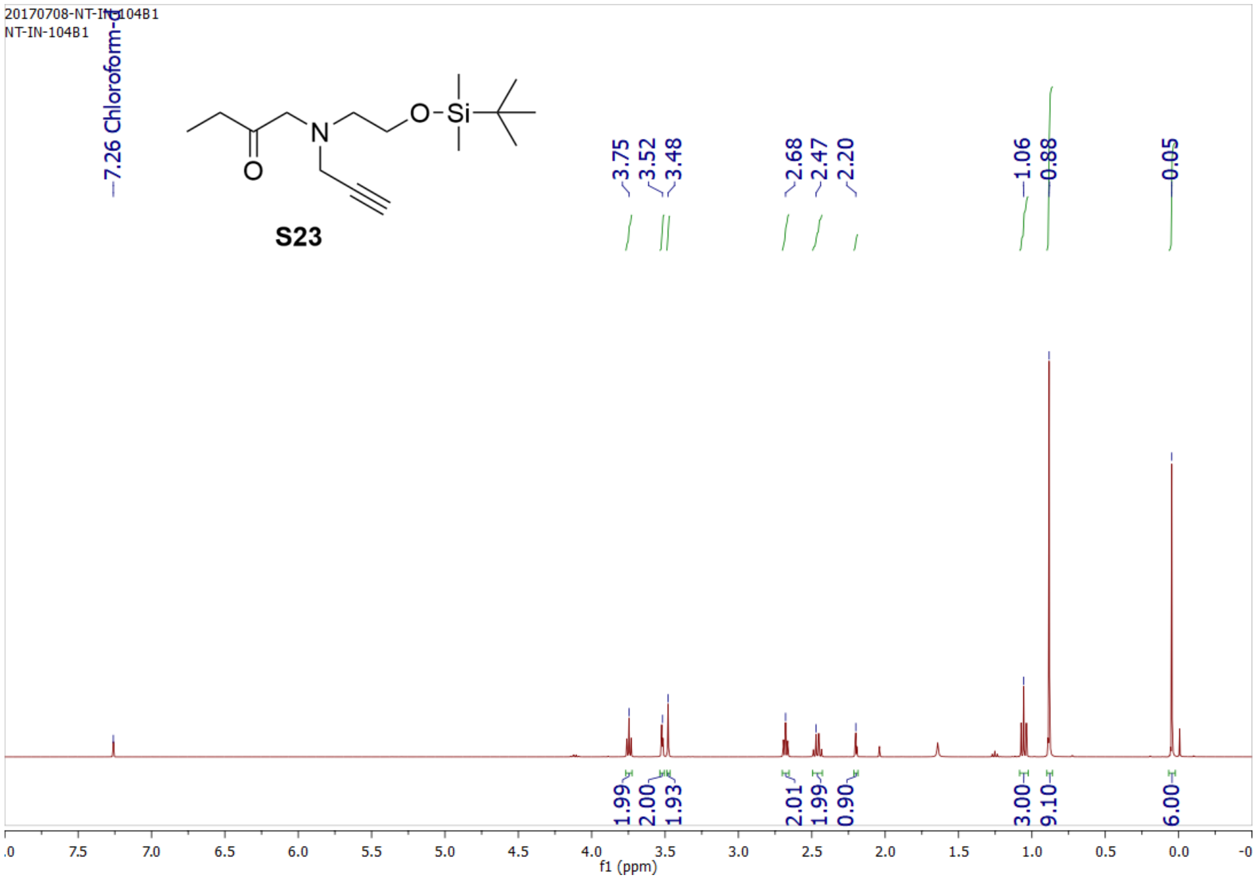
20170708-NT-IN-104B1
NT-IN-104B1
7.26 Chloroform-d
S23
3.75
3.52
3.48
2.68
2.47
2.20
1.06
0.88
0.05
1.99
2.00
1.93
2.01
1.99
0.90
3.00
9.10
6.00
f1 (ppm)

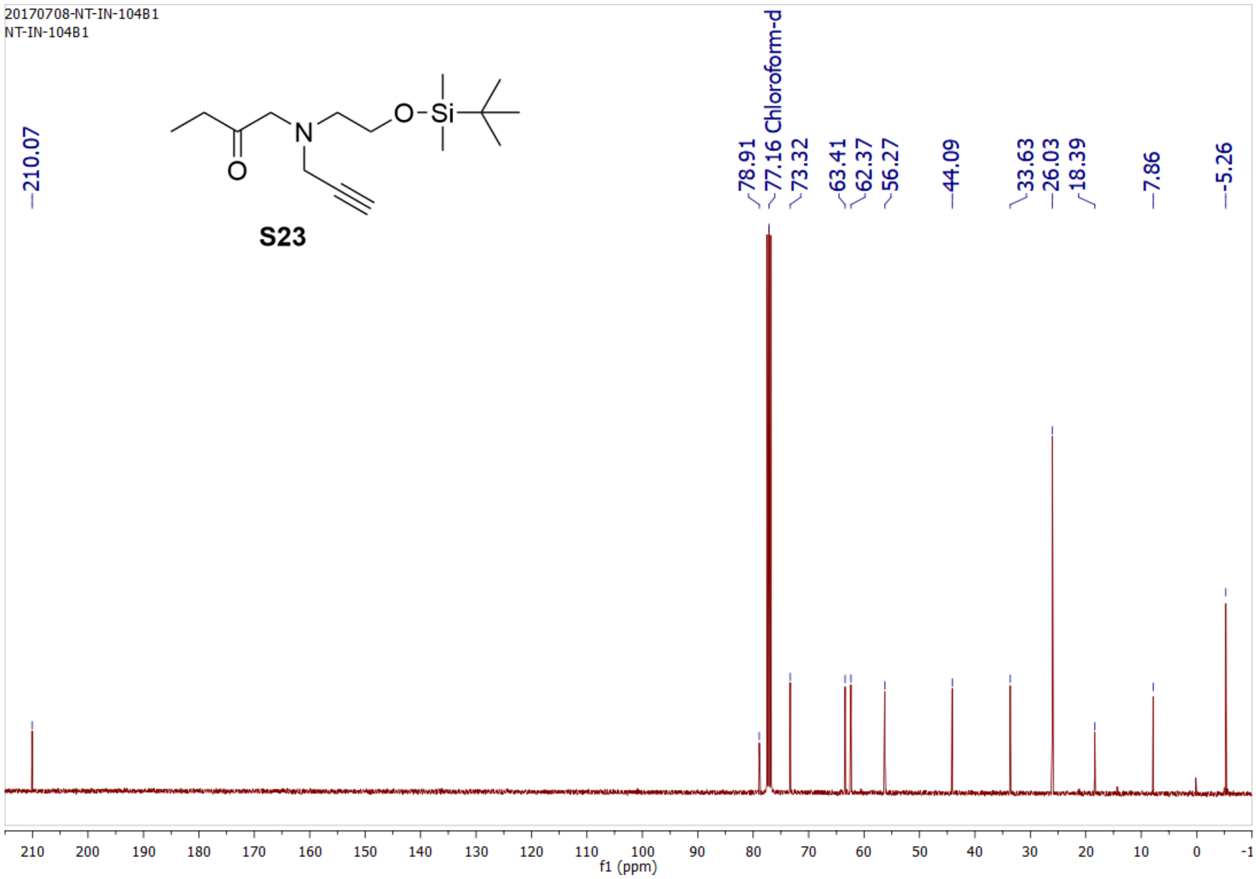
20170708-NT-IN-104B1
NT-IN-104B1
210.07
S23
78.91
77.16 Chloroform-d
73.32
63.41
62.37
56.27
44.09
33.63
26.03
18.39
7.86
-5.26
f1 (ppm)

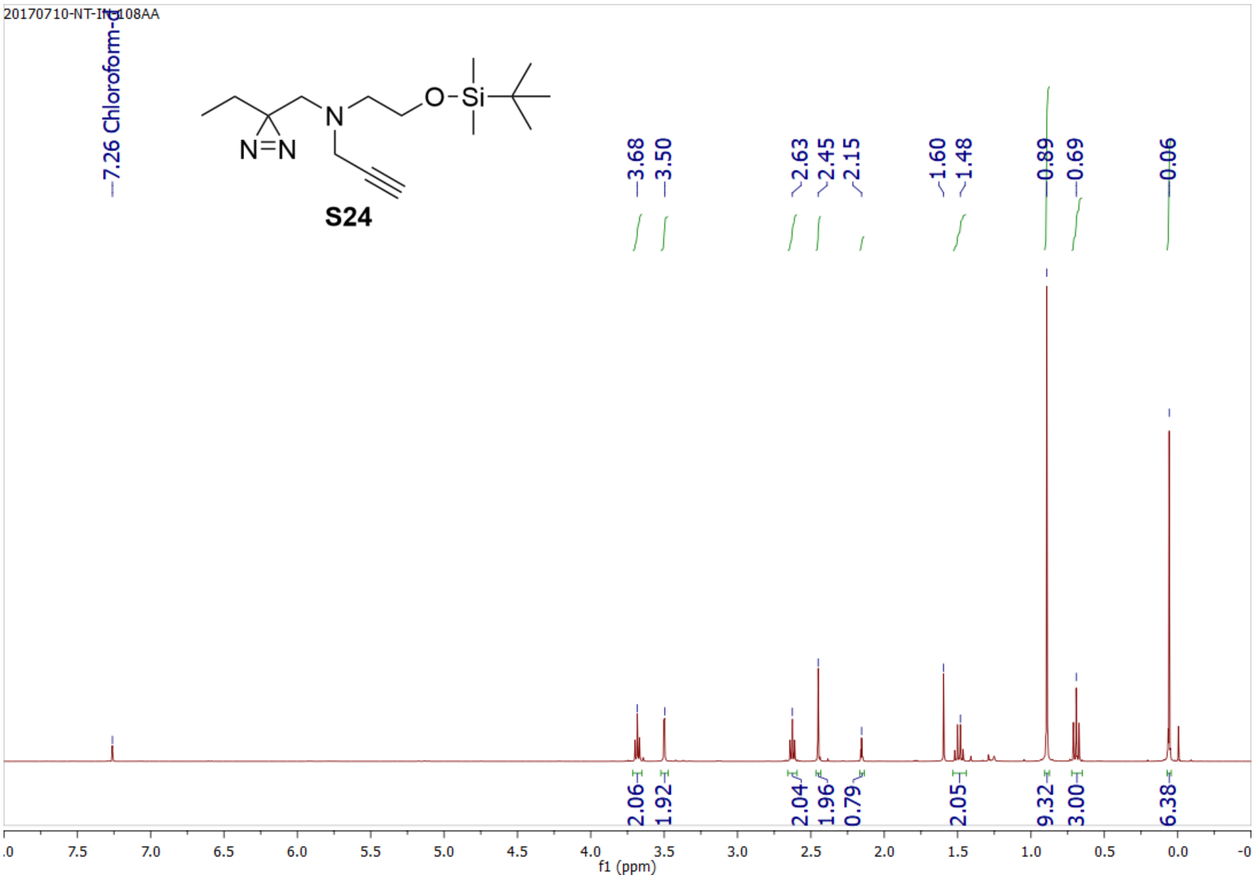
20170710-NT-IN-108AA
7.26 Chloroform-d
S24
3.68
3.50
2.63
2.45
2.15
1.60
1.48
0.89
0.69
0.06
2.06
1.92
2.04
1.96
0.79
2.05
9.32
3.00
6.38
f1 (ppm)

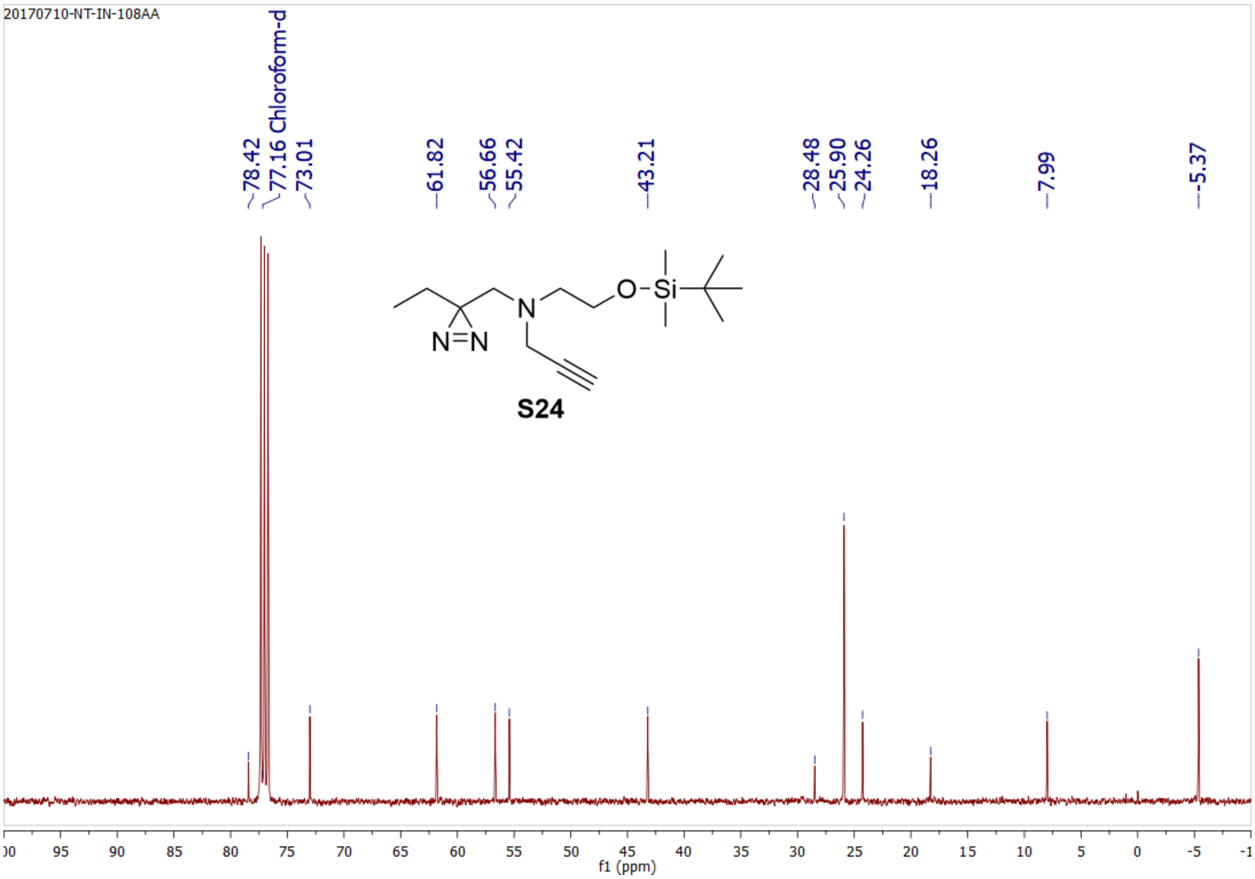
20170710-NT-IN-108AA
78.42
77.16 Chloroform-d
73.01
61.82
56.66
55.42
43.21
28.48
25.90
24.26
18.26
7.99
-5.37
S24
f1 (ppm)

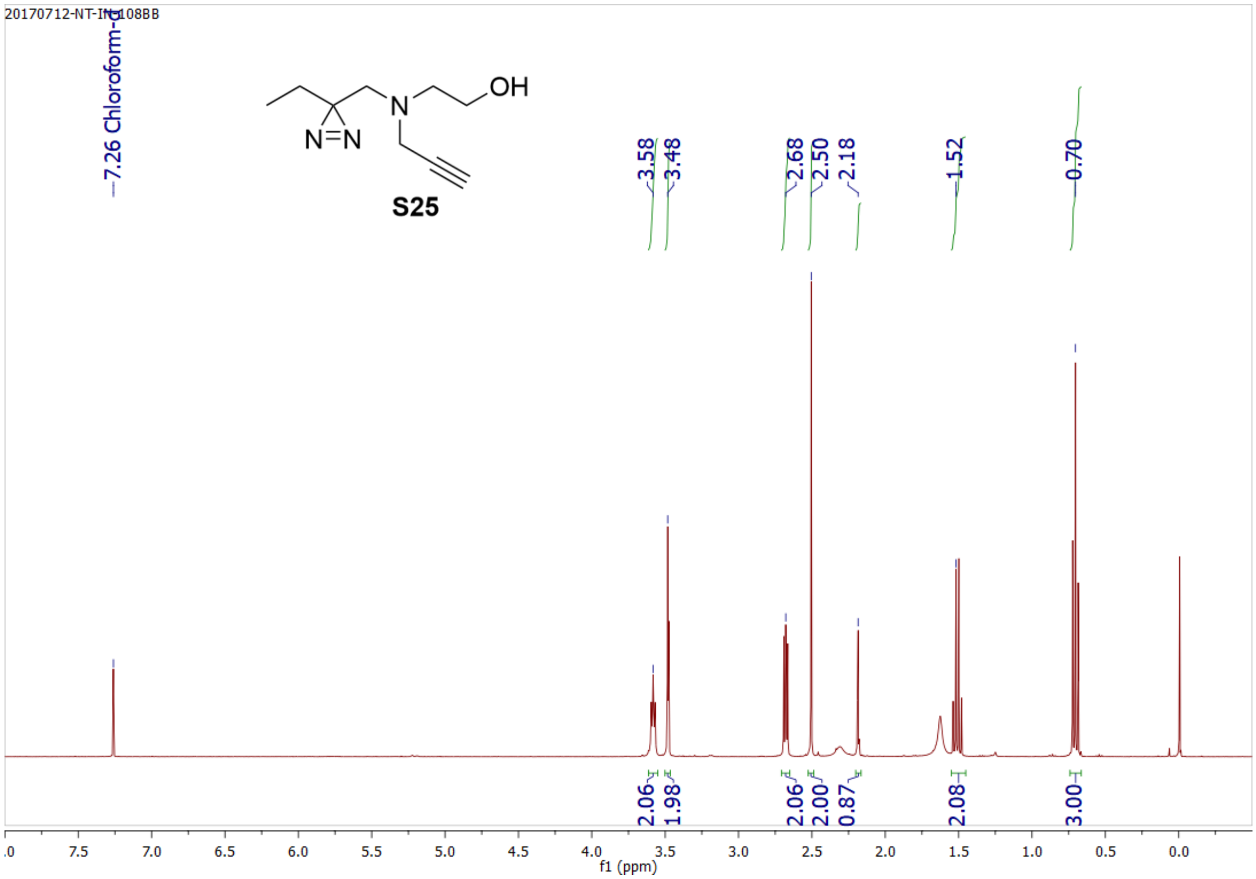
20170712-NT-IN-108BB
7.26 Chloroform-d
3.58
3.48
2.68
2.50
2.18
1.52
0.70
S25
OH
N
N=N
2.06
1.98
2.06
2.00
0.87
2.08
3.00
f1 (ppm)

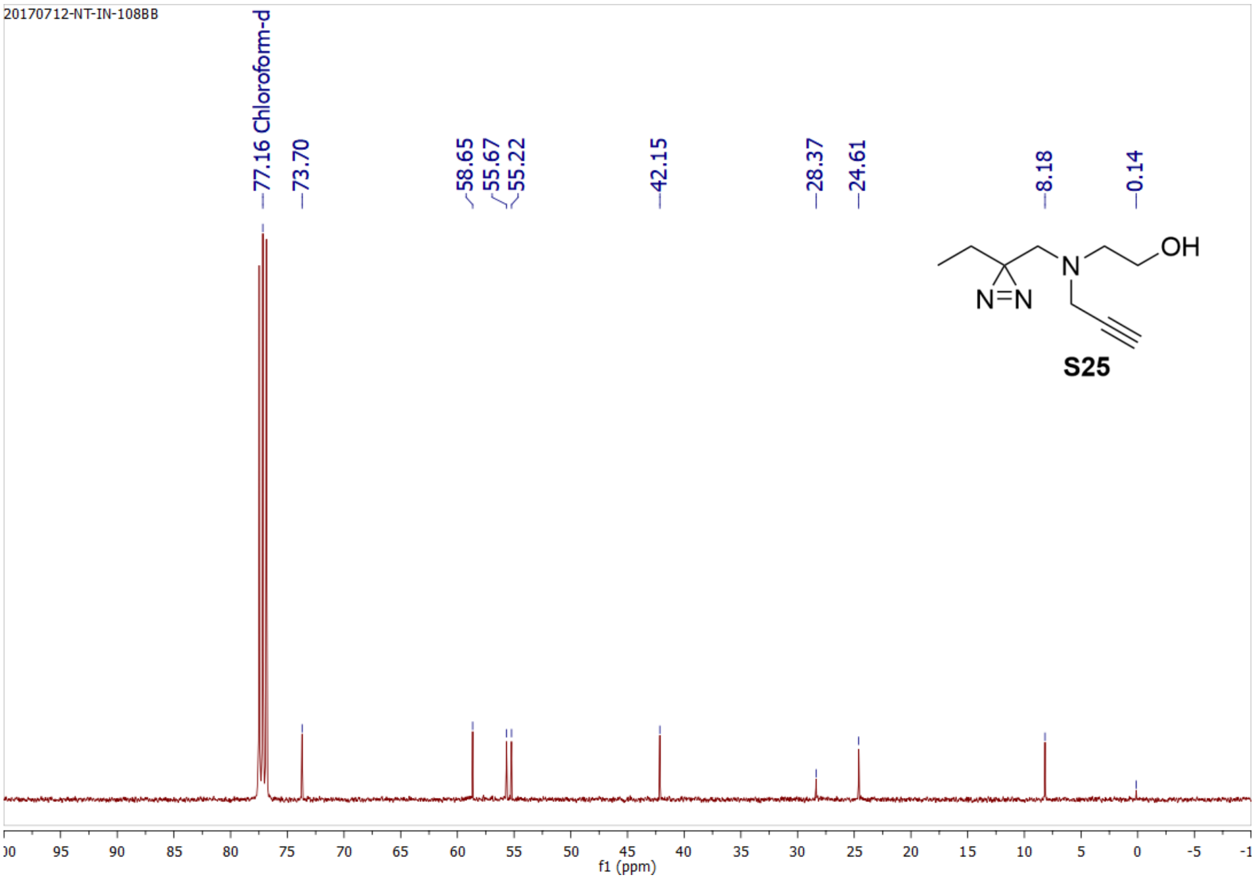
20170712-NT-IN-108BB
77.16 Chloroform-d
73.70
58.65
55.67
55.22
42.15
28.37
24.61
8.18
0.14
OH
N
N=N
S25
f1 (ppm)

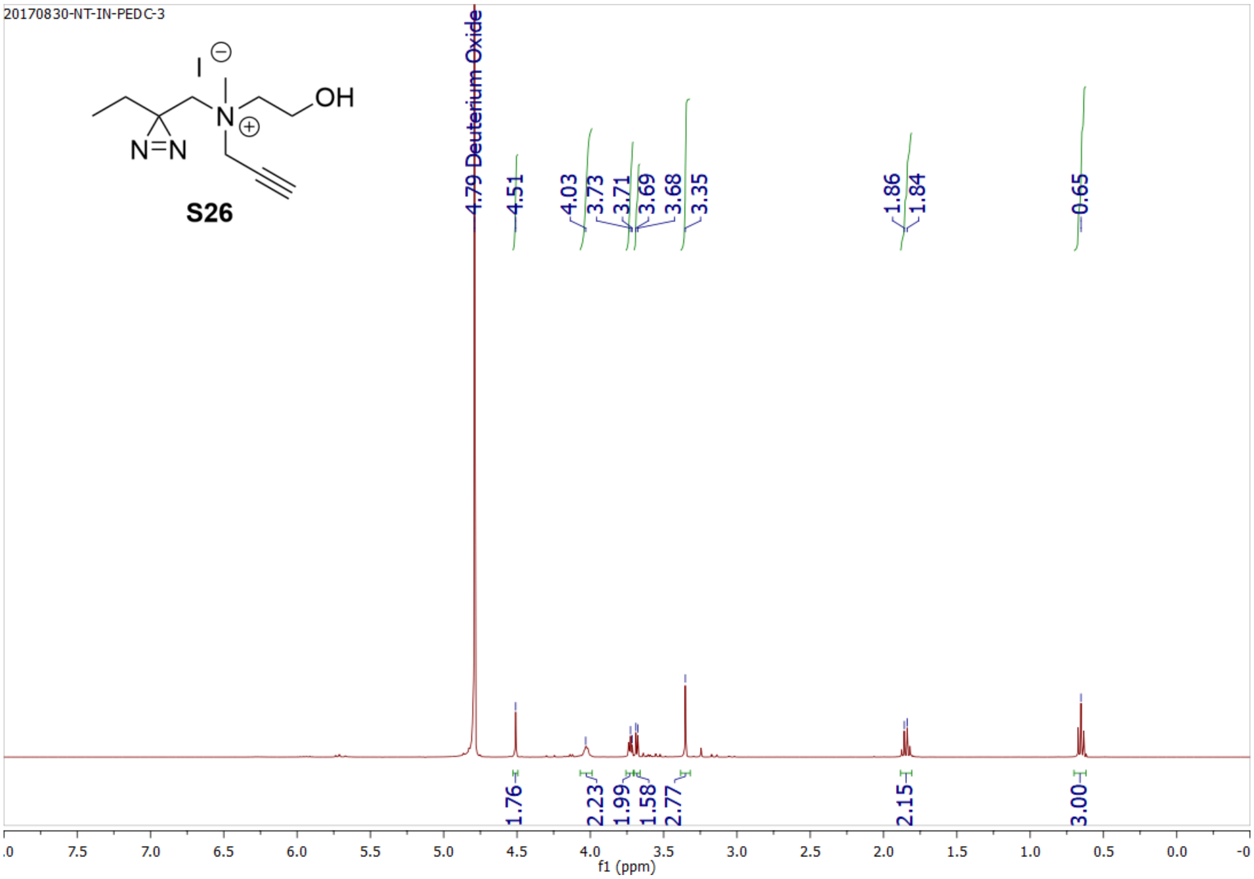
20170830-NT-IN-PEDC-3
S26
4.79 Deuterium Oxide
4.51
4.03
3.73
3.71
3.69
3.68
3.35
1.86
1.84
0.65
1.76
2.23
1.99
1.58
2.77
2.15
3.00
f1 (ppm)

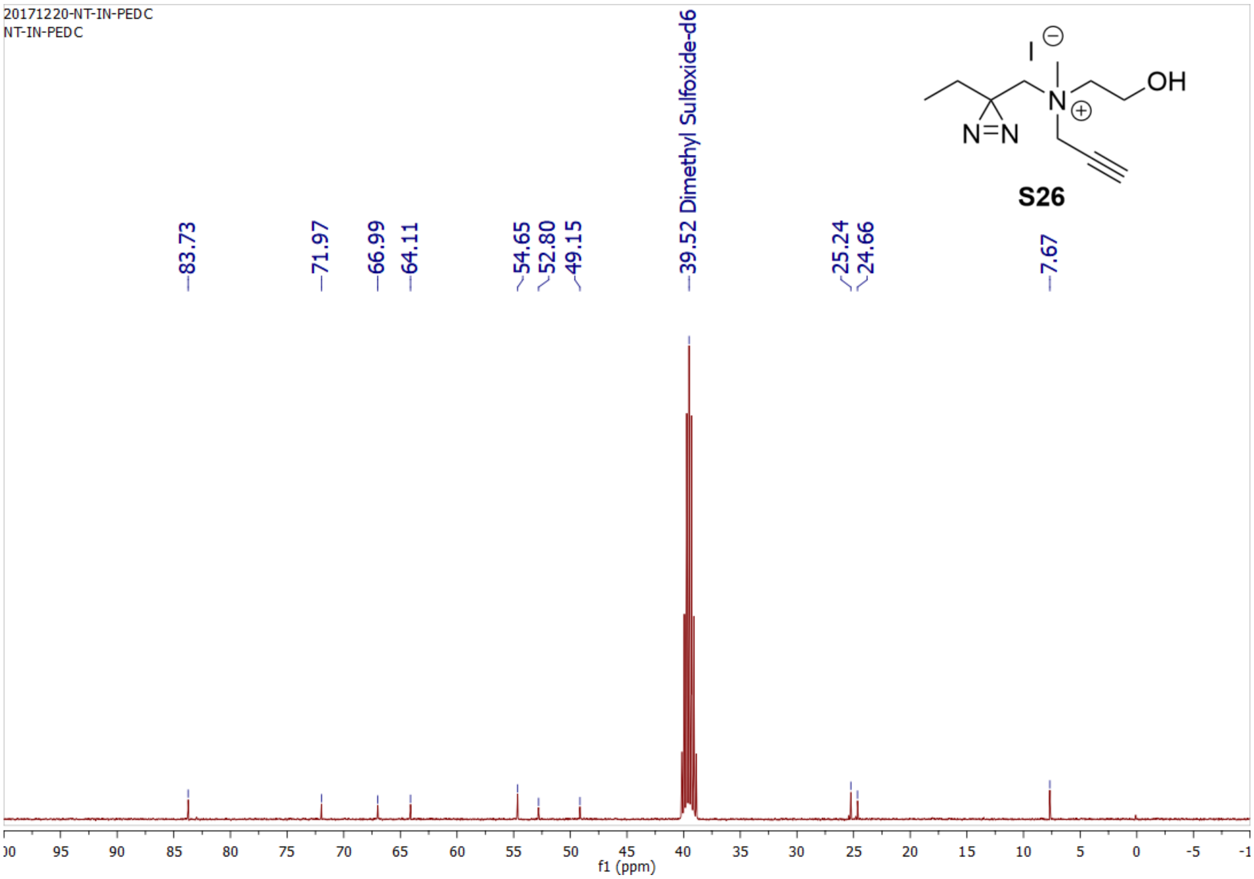
20171220-NT-IN-PEDC
NT-IN-PEDC
S26
83.73
71.97
66.99
64.11
54.65
52.80
49.15
39.52 Dimethyl Sulfoxide-d6
25.24
24.66
7.67
f1 (ppm)

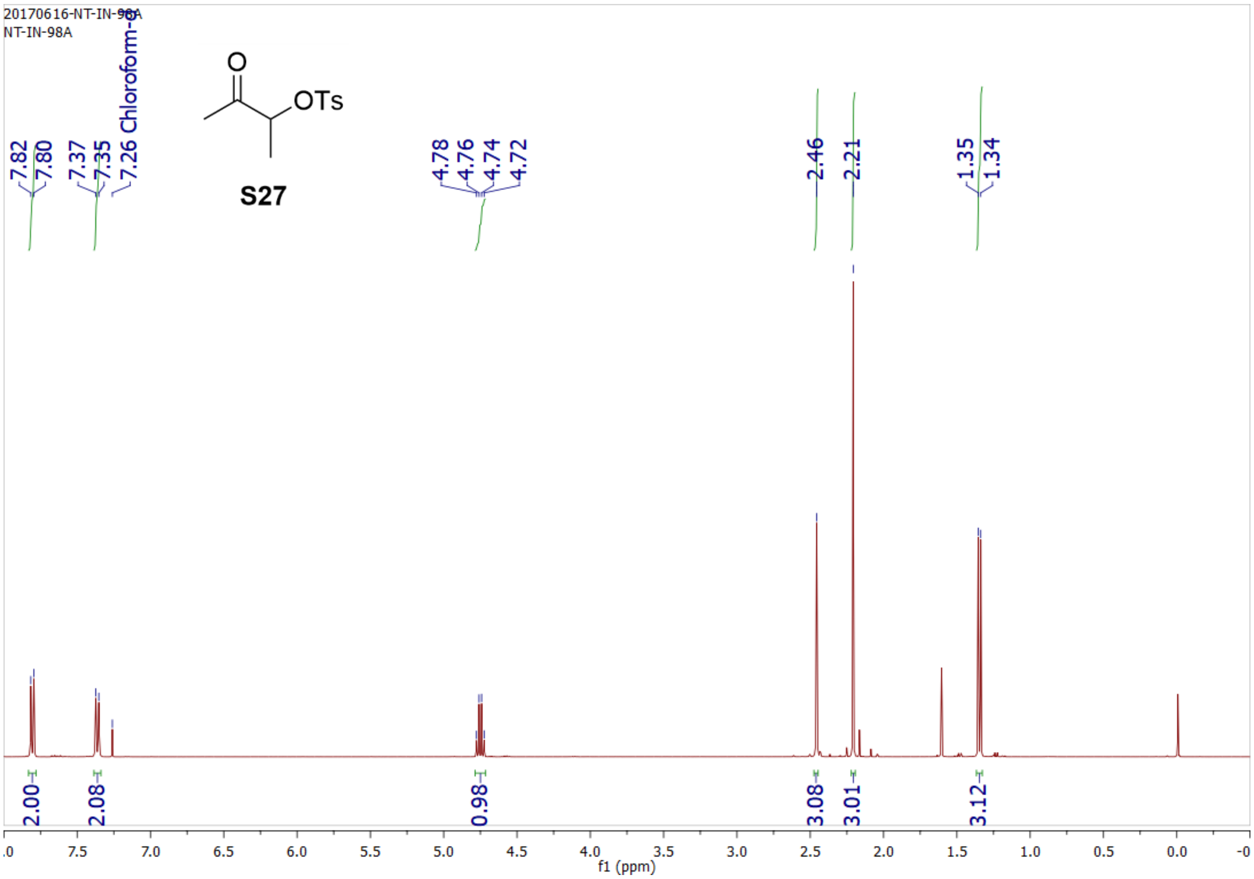
20170616-NT-IN-98A
NT-IN-98A
S27
OTs
O
7.82
7.80
7.37
7.35
7.26 Chloroform-d
4.78
4.76
4.74
4.72
2.46
2.21
1.35
1.34
2.00
2.08
0.98
3.08
3.01
3.12
f1 (ppm)

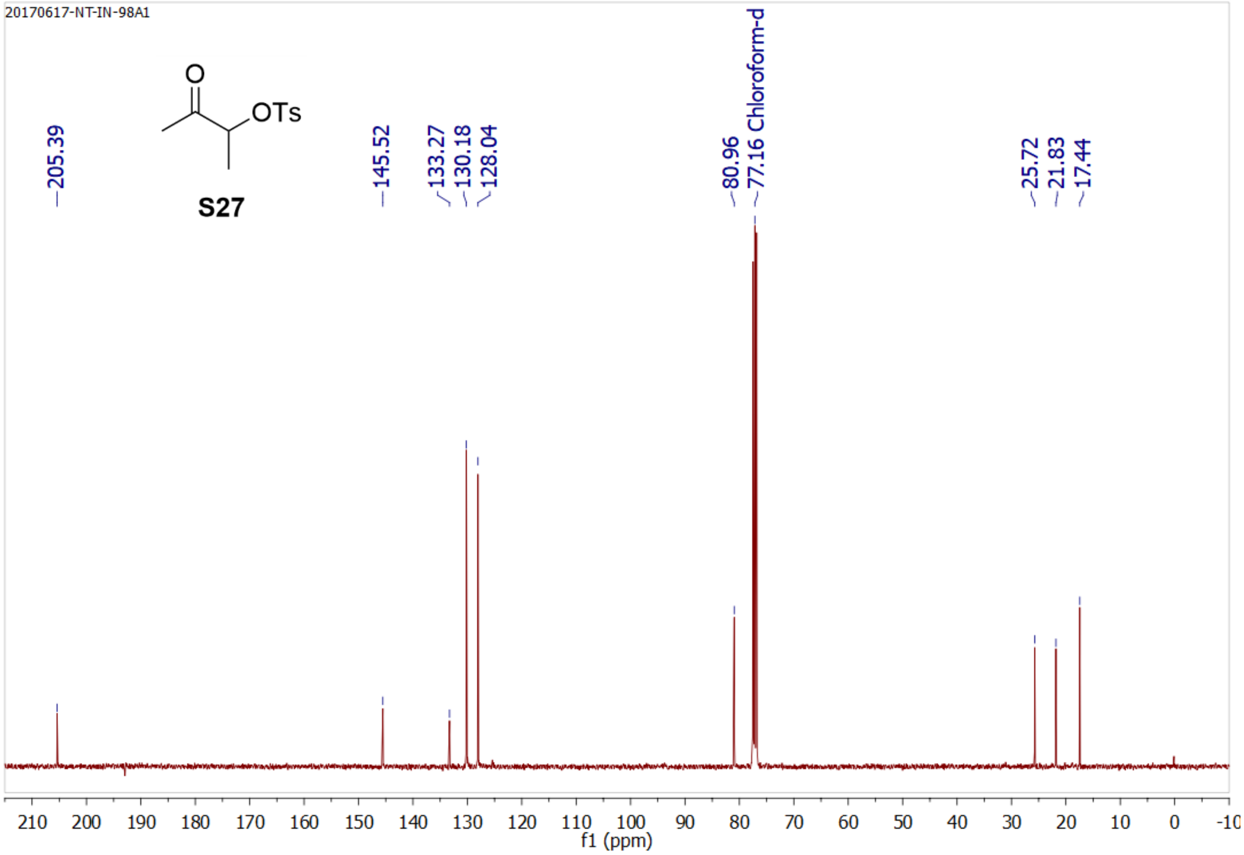
20170617-NT-IN-98A1
S27
OTs
O
205.39
145.52
133.27
130.18
128.04
80.96
77.16 Chloroform-d
25.72
21.83
17.44
f1 (ppm)

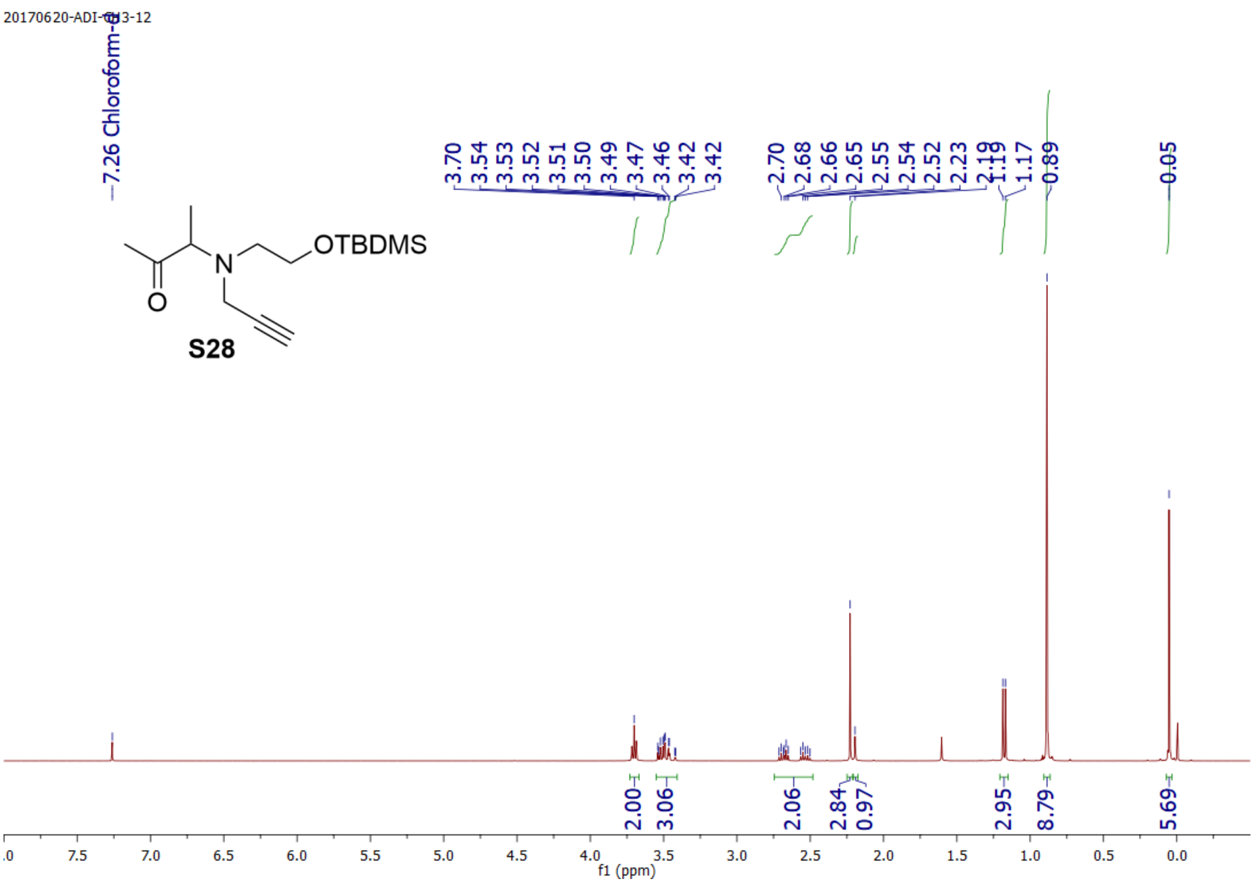
20170620-ADI-CH3-12
7.26 Chloroform-d
3.70 3.54 3.53 3.52 3.51 3.50 3.49 3.47 3.46 3.42 3.42
2.70 2.68 2.66 2.65 2.55 2.54 2.52 2.23 2.19 1.19 1.17 0.89 0.05
OTBDMS
N
O
S28
2.00 3.06 2.06 2.84 0.97 2.95 8.79 5.69
f1 (ppm)

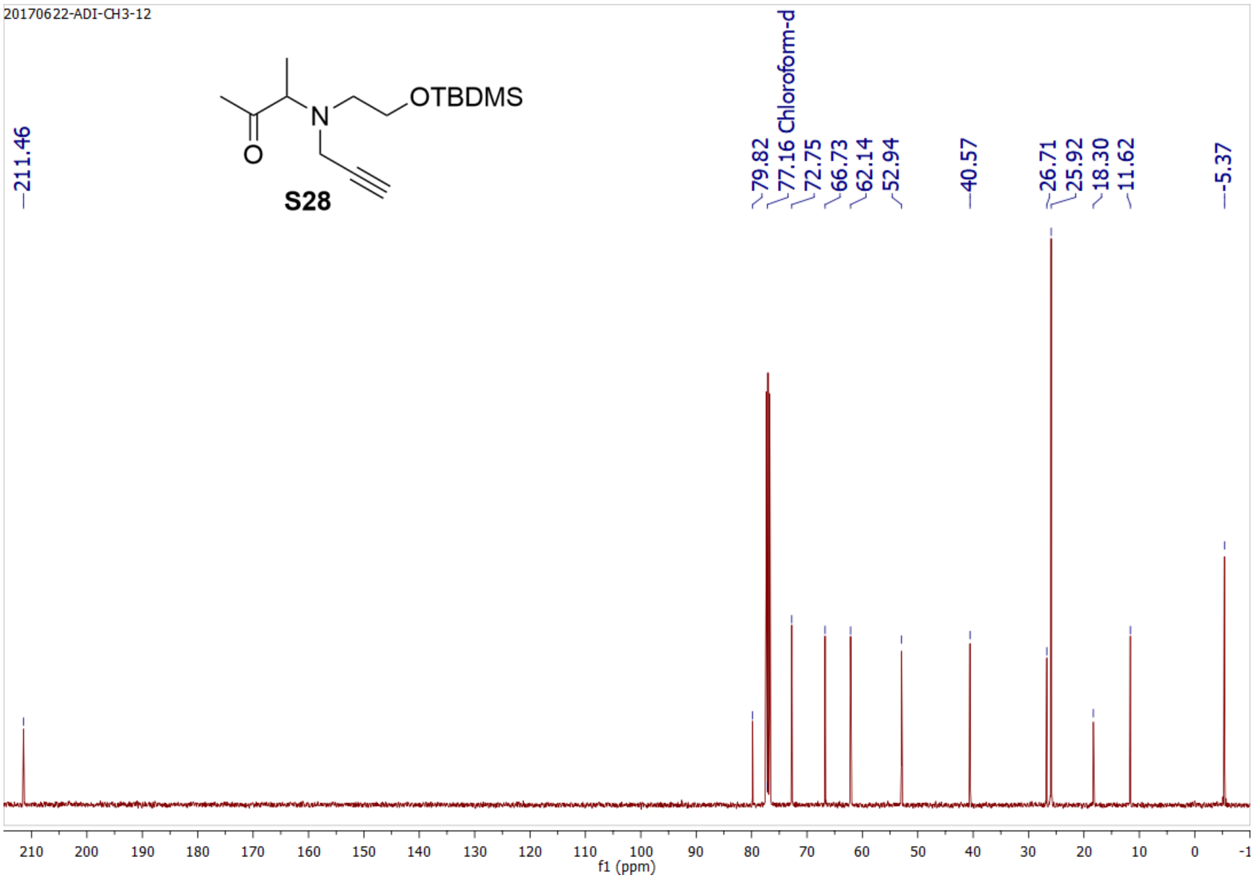
20170622-ADI-CH3-12
211.46
79.82 77.16 Chloroform-d 72.75 66.73 62.14 52.94 40.57 26.71 25.92 18.30 11.62 -5.37
OTBDMS
N
O
S28
f1 (ppm)

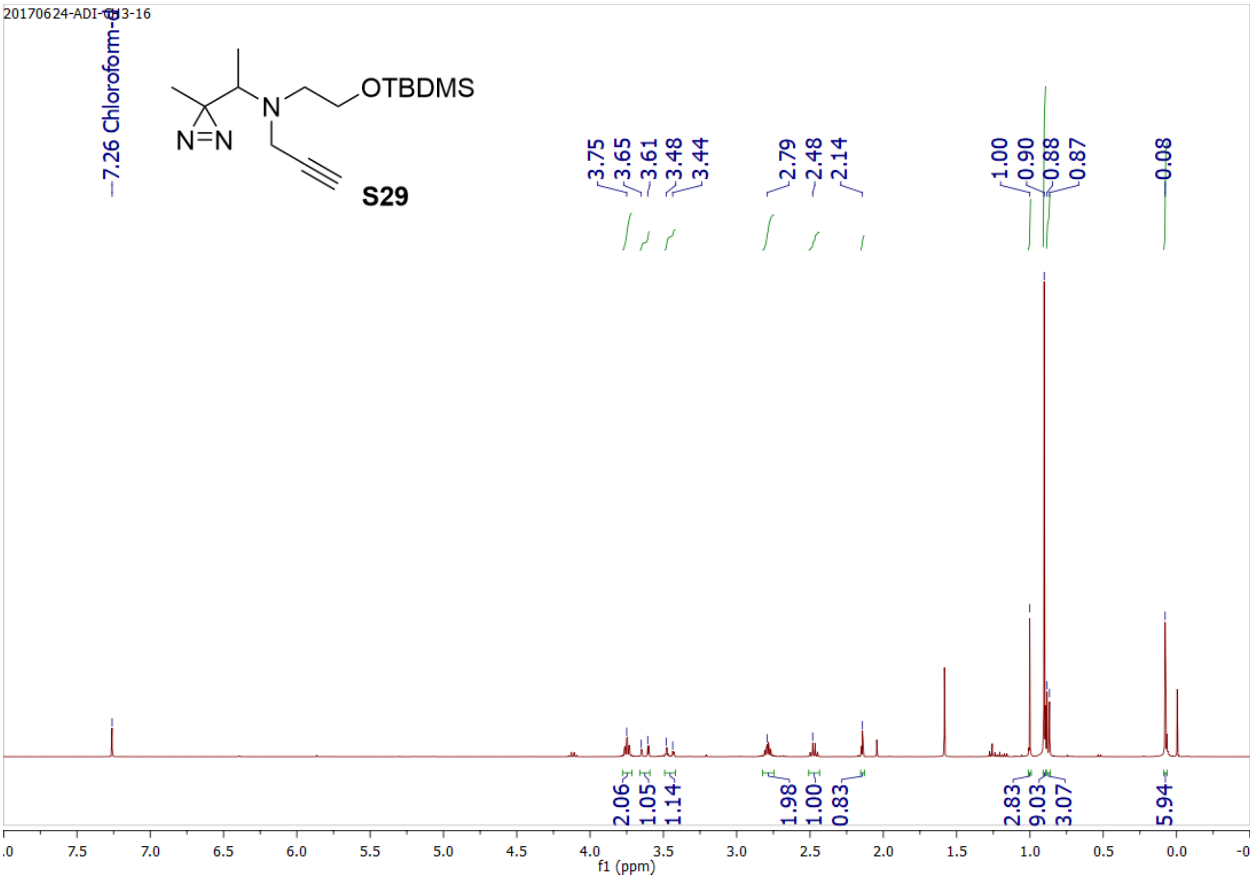

20170624-ADI-CH3-16
7.26 Chloroform-d
OTBDMS
N
N=N
S29
3.75
3.65
3.61
3.48
3.44
2.79
2.48
2.14
1.00
0.90
0.88
0.87
0.08
2.06
1.05
1.14
1.98
1.00
0.83
2.83
9.03
3.07
5.94
f1 (ppm)


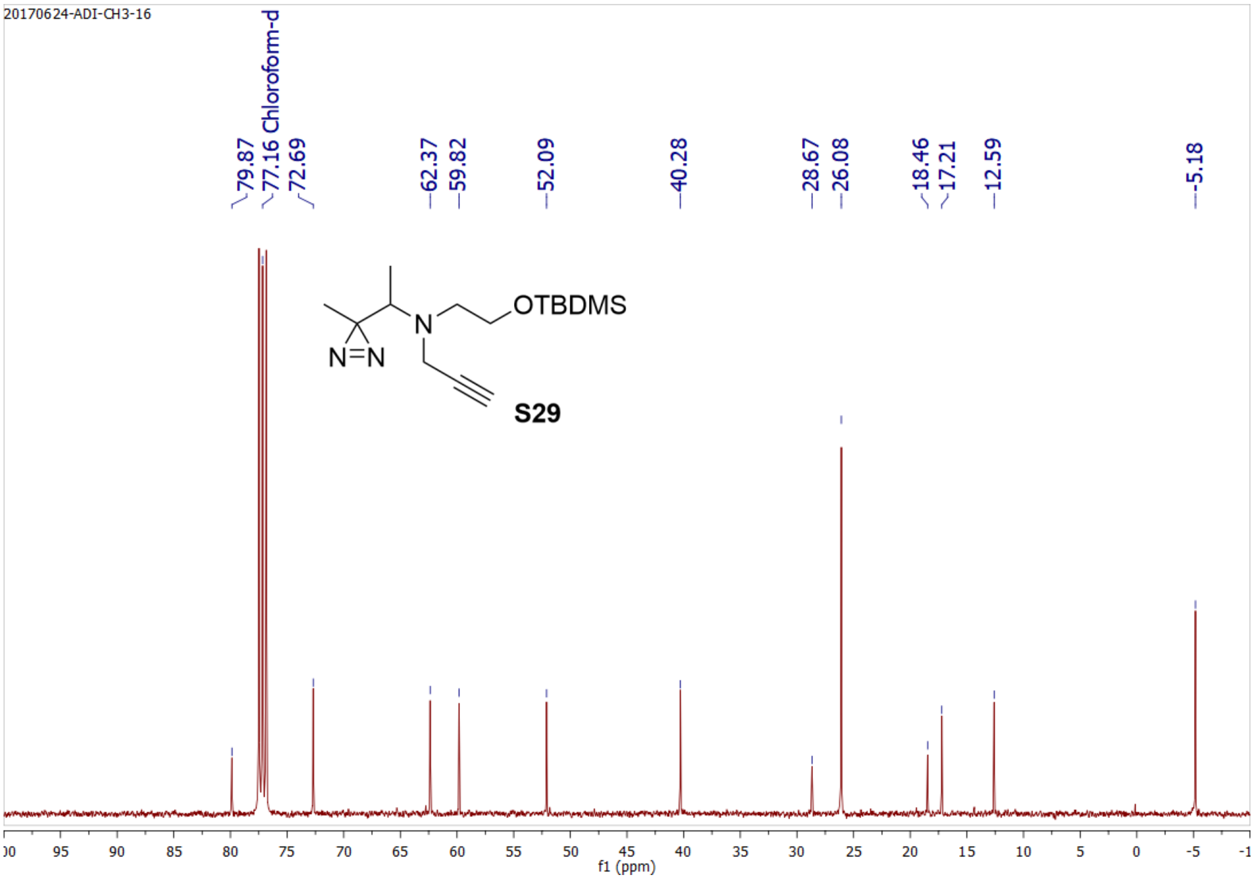

20170624-ADI-CH3-16
79.87
77.16 Chloroform-d
72.69
62.37
59.82
52.09
40.28
28.67
26.08
18.46
17.21
12.59
-5.18
OTBDMS
N
N=N
S29
f1 (ppm)

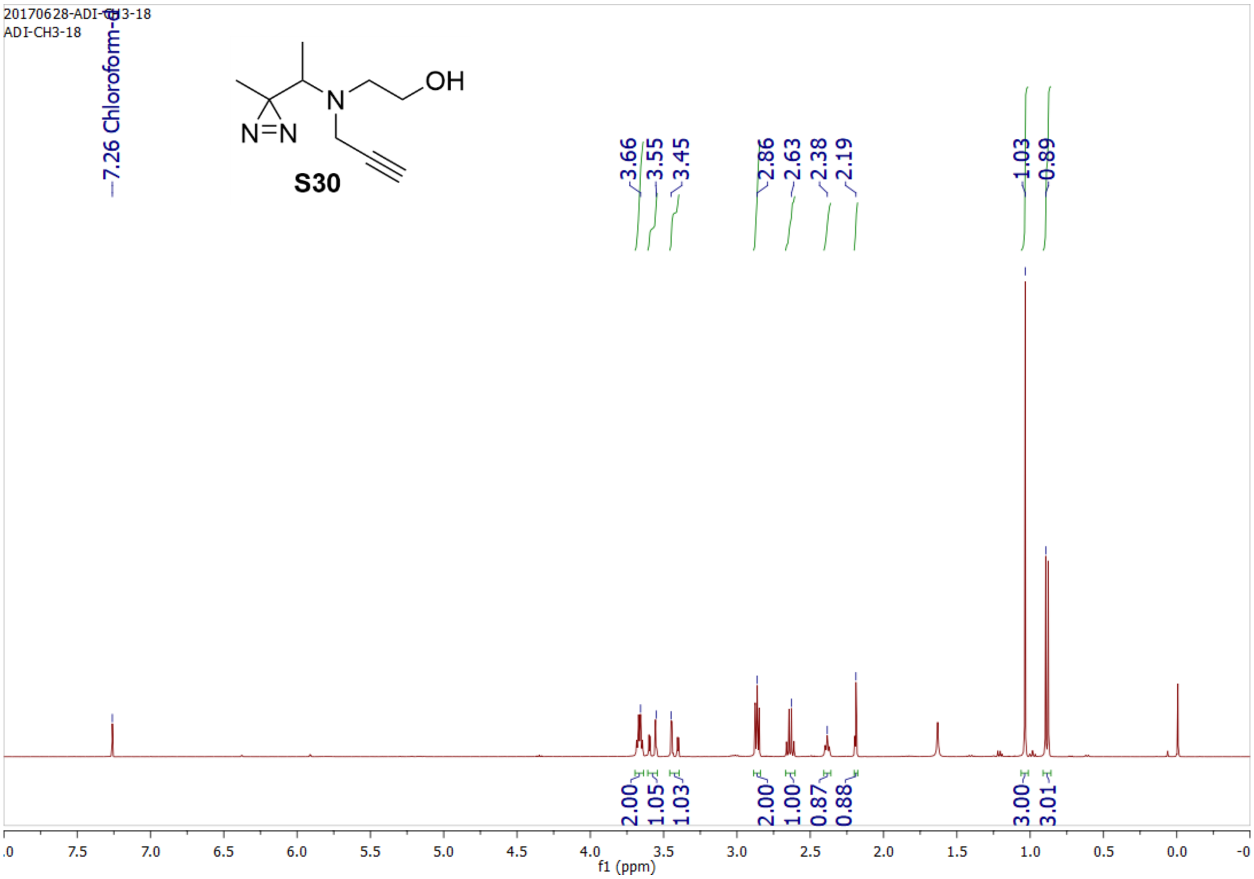
20170628-ADI-CH3-18
ADI-CH3-18
7.26 Chloroform-d
OH
N
N=N
S30
3.66
3.55
3.45
2.86
2.63
2.38
2.19
1.03
0.89
2.00
1.05
1.03
2.00
1.00
0.87
0.88
3.00
3.01
f1 (ppm)

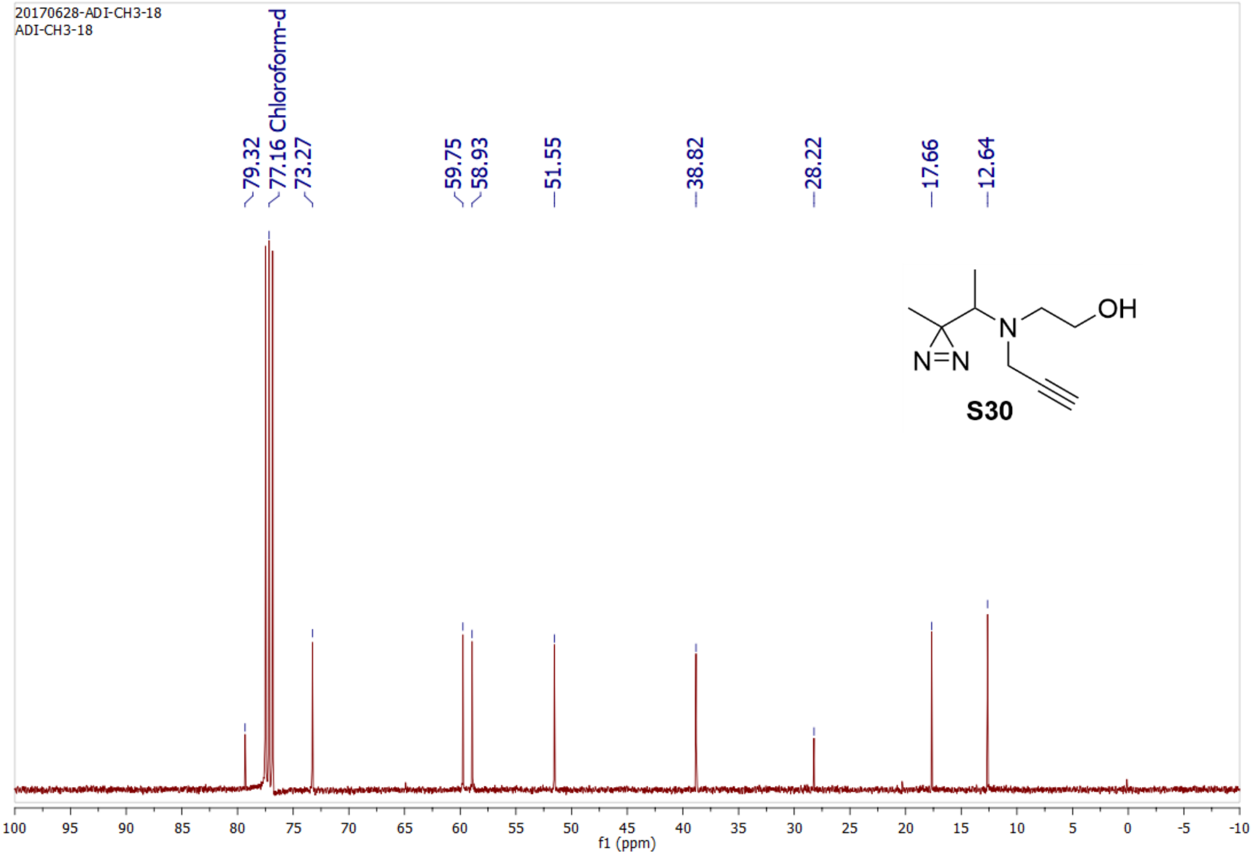
20170628-ADI-CH3-18
ADI-CH3-18
79.32
77.16 Chloroform-d
73.27
59.75
58.93
51.55
38.82
28.22
17.66
12.64
OH
N
N=N
S30
f1 (ppm)

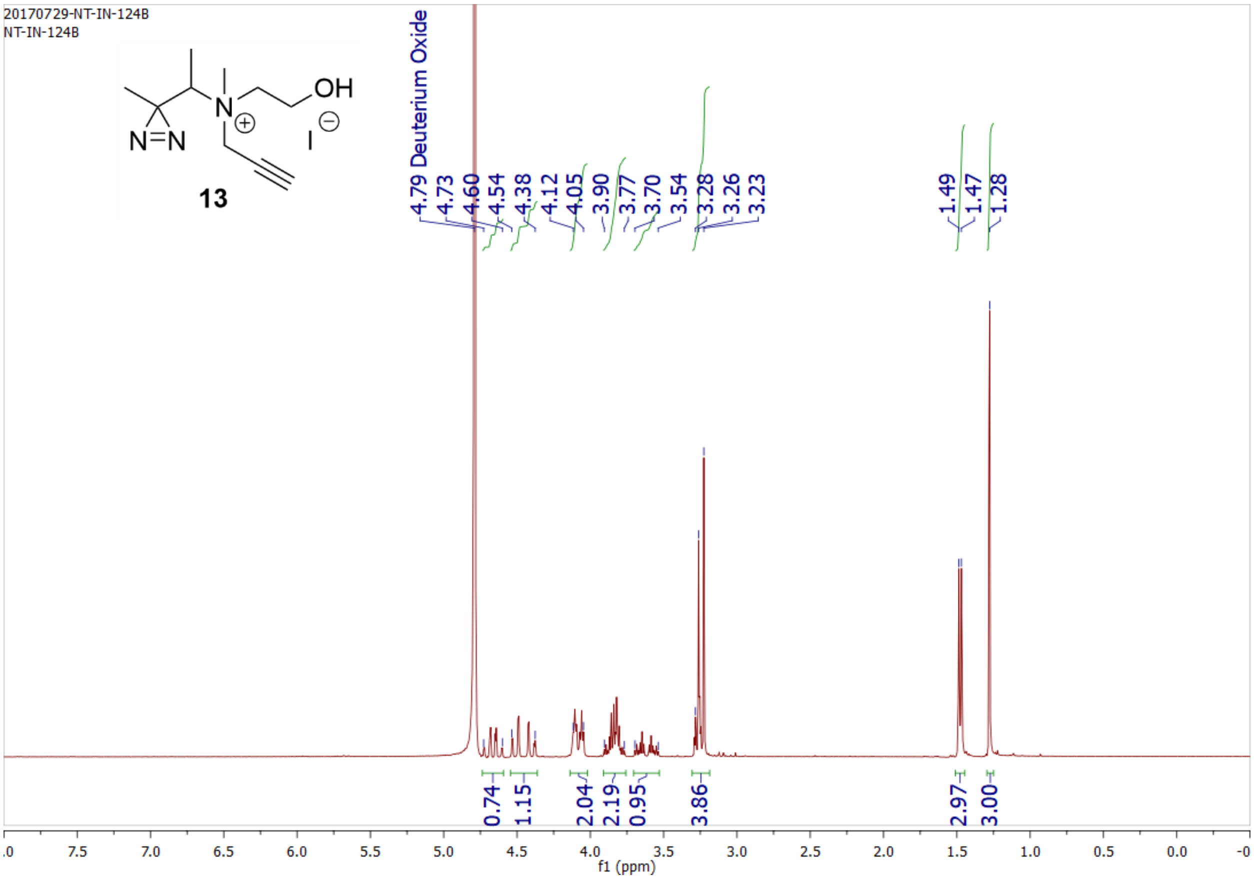
20170729-NT-IN-124B
NT-IN-124B
OH
N
I
N=N
13
4.79 Deuterium Oxide
4.73
4.60
4.54
4.38
4.12
4.05
3.90
3.77
3.70
3.54
3.28
3.26
3.23
1.49
1.47
1.28
0.74
1.15
2.04
2.19
0.95
3.86
2.97
3.00
f1 (ppm)

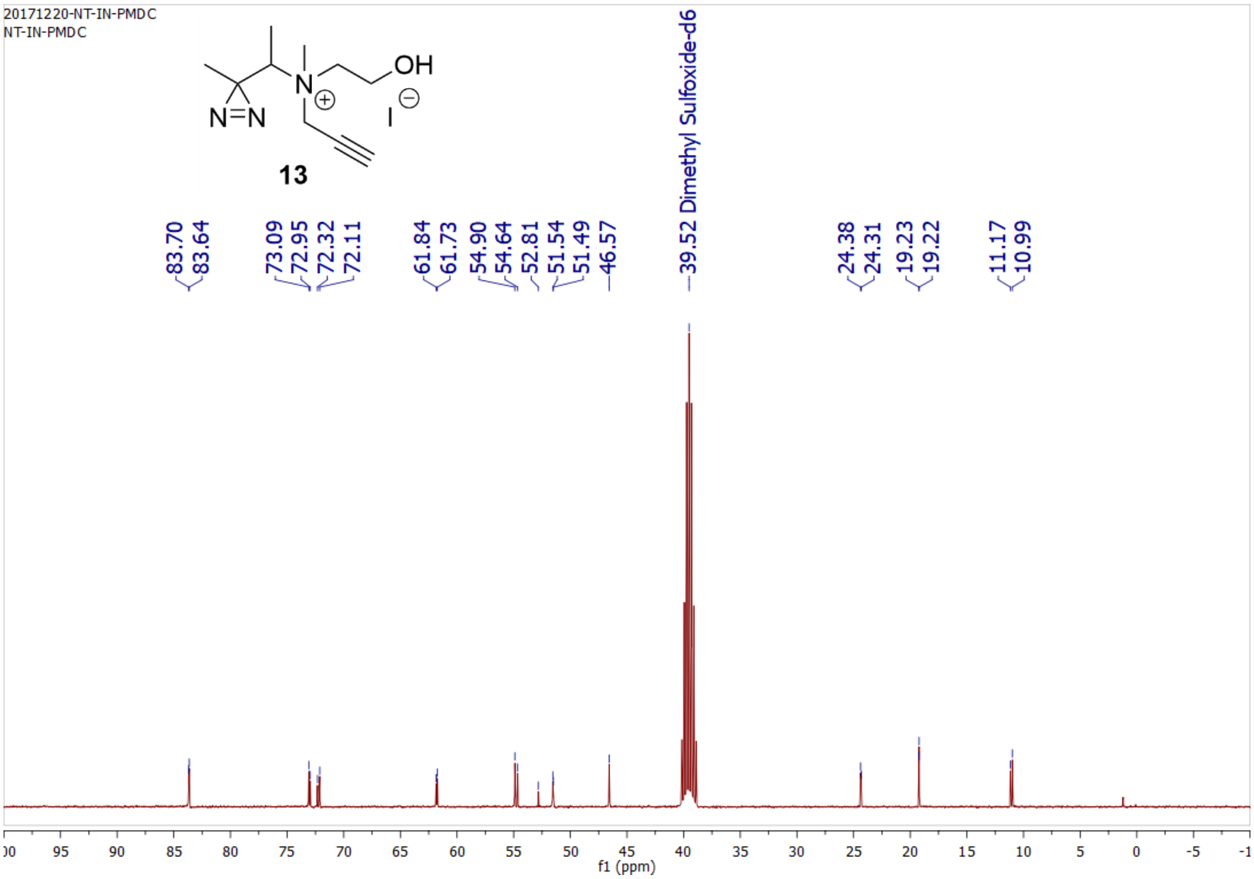
20171220-NT-IN-PMDC
NT-IN-PMDC
OH
N
I
N=N
13
39.52 Dimethyl Sulfoxide-d6
83.70
83.64
73.09
72.95
72.32
72.11
61.84
61.73
54.90
54.64
52.81
51.54
51.49
46.57
24.38
24.31
19.23
19.22
11.17
10.99
f1 (ppm)

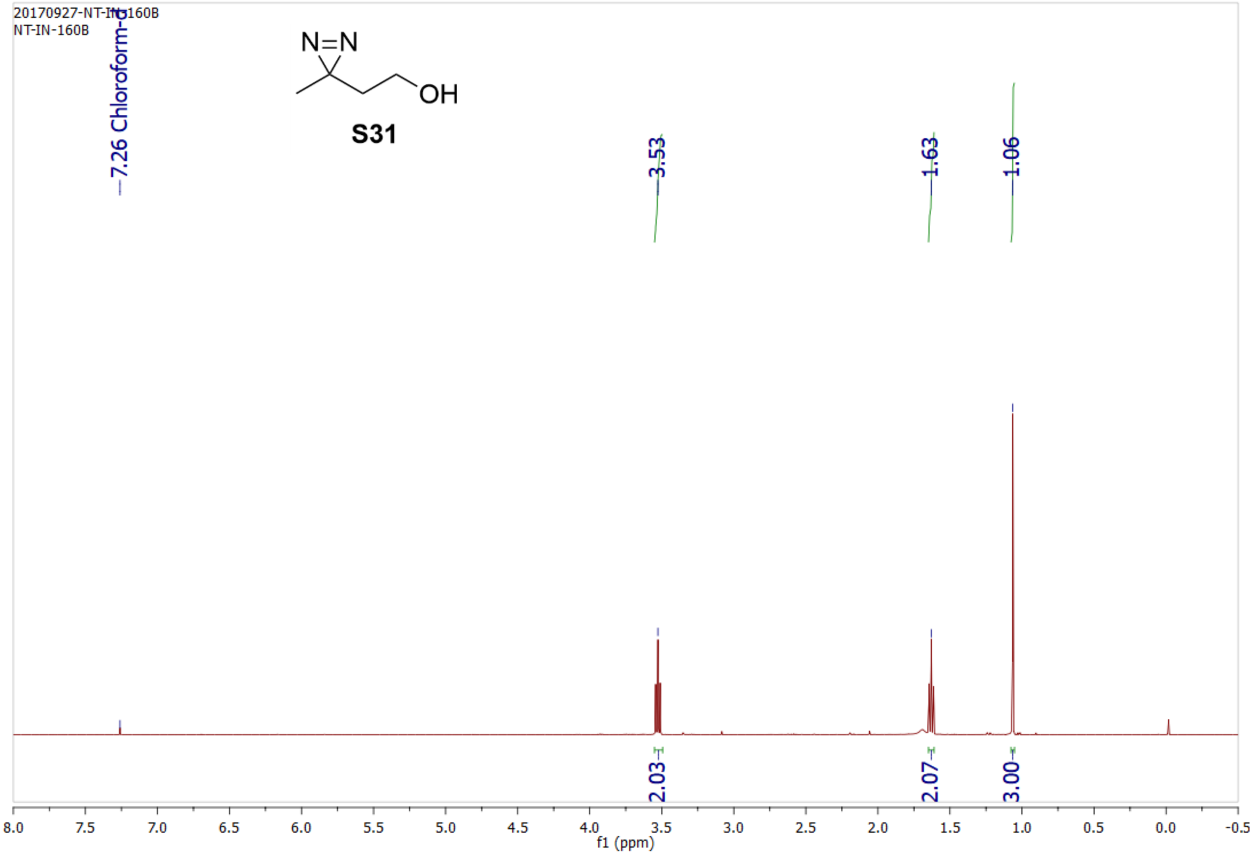
20170927-NT-IN-160B
NT-IN-160B
N=N
OH
S31
7.26 Chloroform-d
3.53
1.63
1.06
2.03
2.07
3.00
8.0 7.5 7.0 6.5 6.0 5.5 5.0 4.5 4.0 3.5 3.0 2.5 2.0 1.5 1.0 0.5 0.0 -0.5
f1 (ppm)

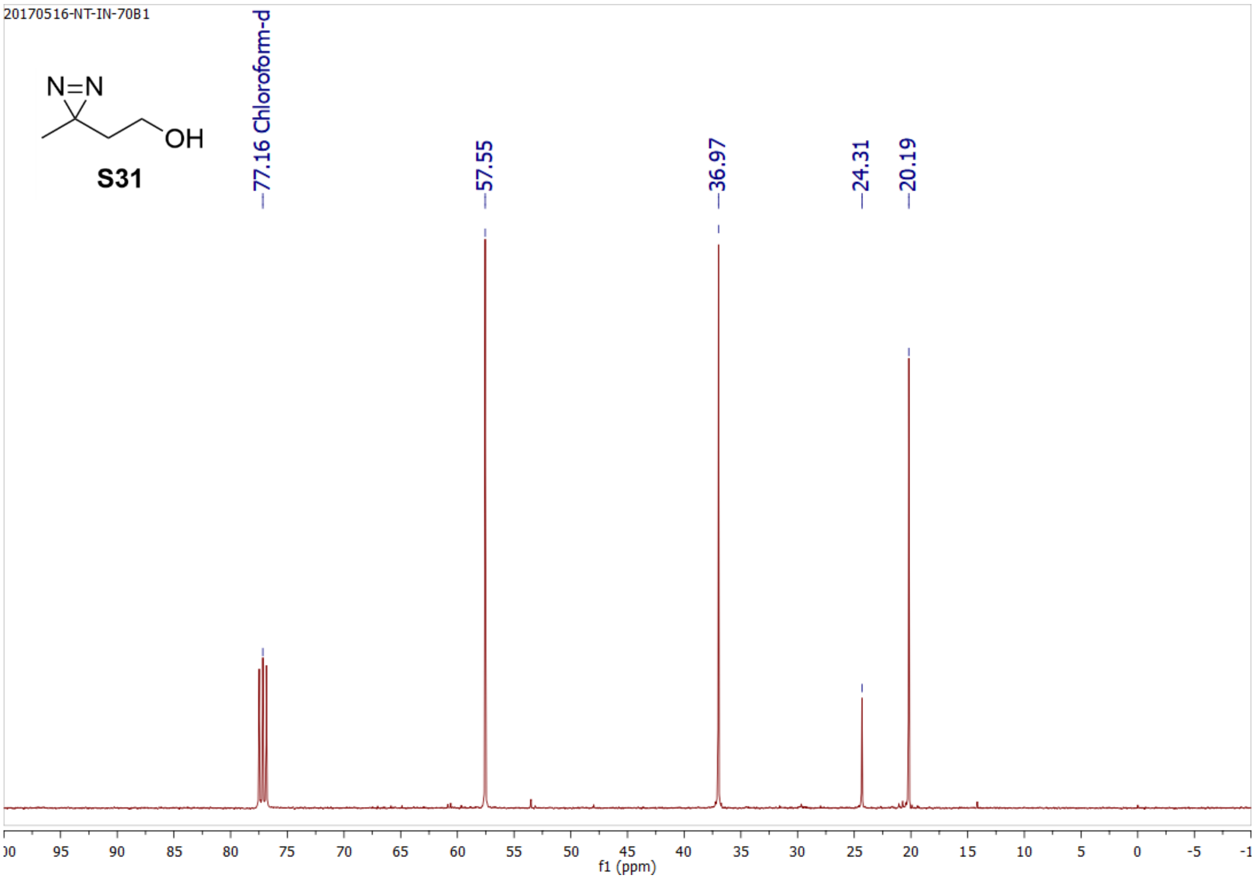
20170516-NT-IN-70B1
N=N
OH
S31
77.16 Chloroform-d
57.55
36.97
24.31
20.19
95 90 85 80 75 70 65 60 55 50 45 40 35 30 25 20 15 10 5 0 -5
f1 (ppm)

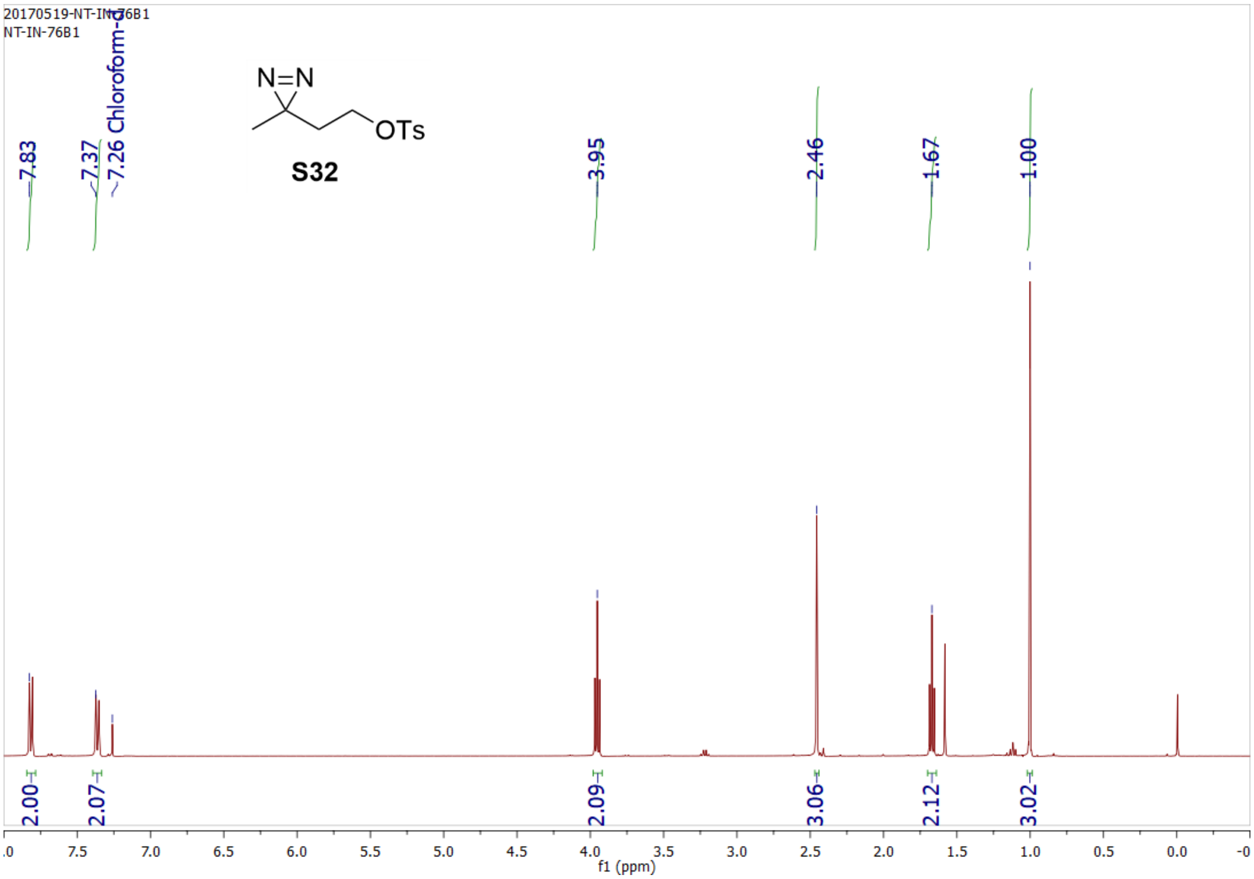
20170519-NT-IN-76B1
NT-IN-76B1
7.26 Chloroform-d
7.83
7.37
N=N
OTs
S32
3.95
2.46
1.67
1.00
2.00
2.07
2.09
3.06
2.12
3.02
f1 (ppm)

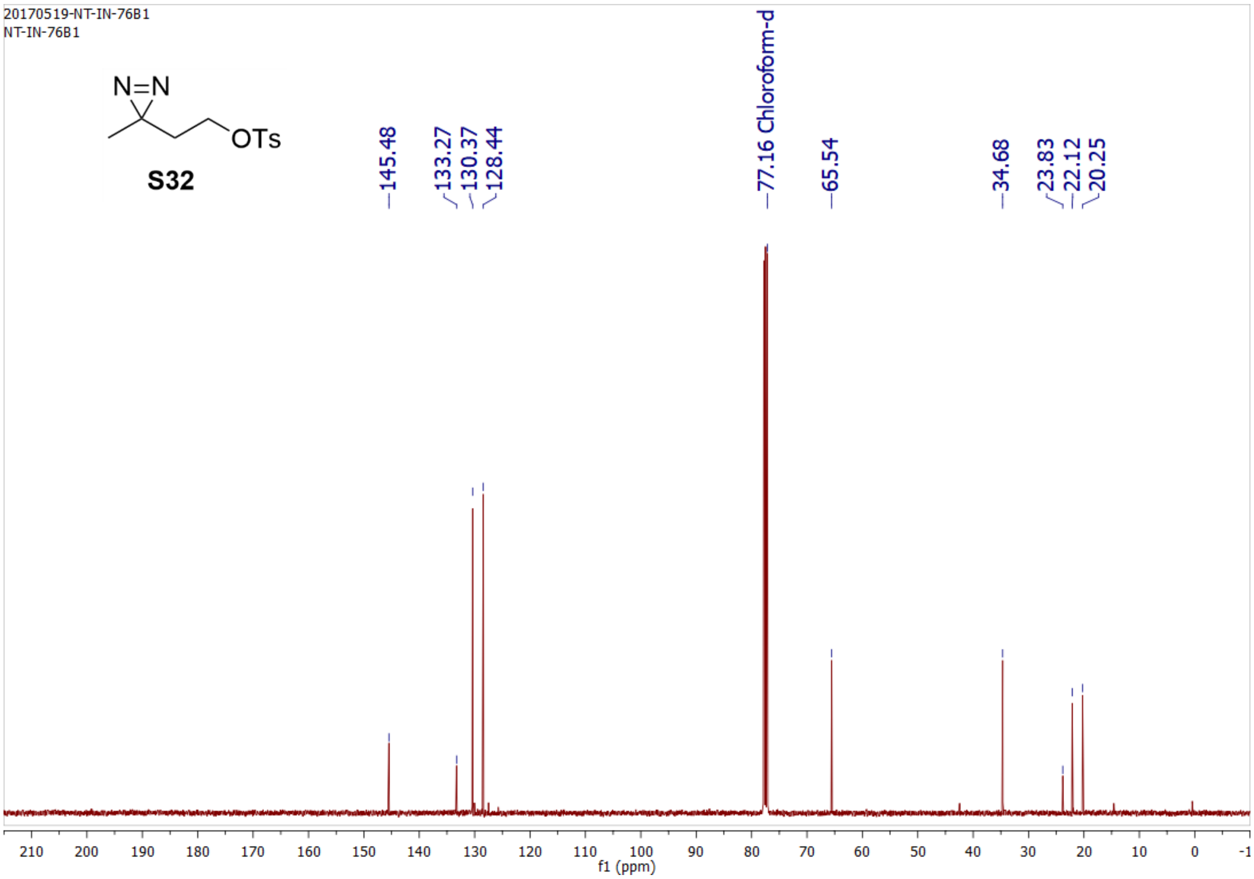
20170519-NT-IN-76B1
NT-IN-76B1
N=N
OTs
S32
145.48
133.27
130.37
128.44
77.16 Chloroform-d
65.54
34.68
23.83
22.12
20.25
f1 (ppm)

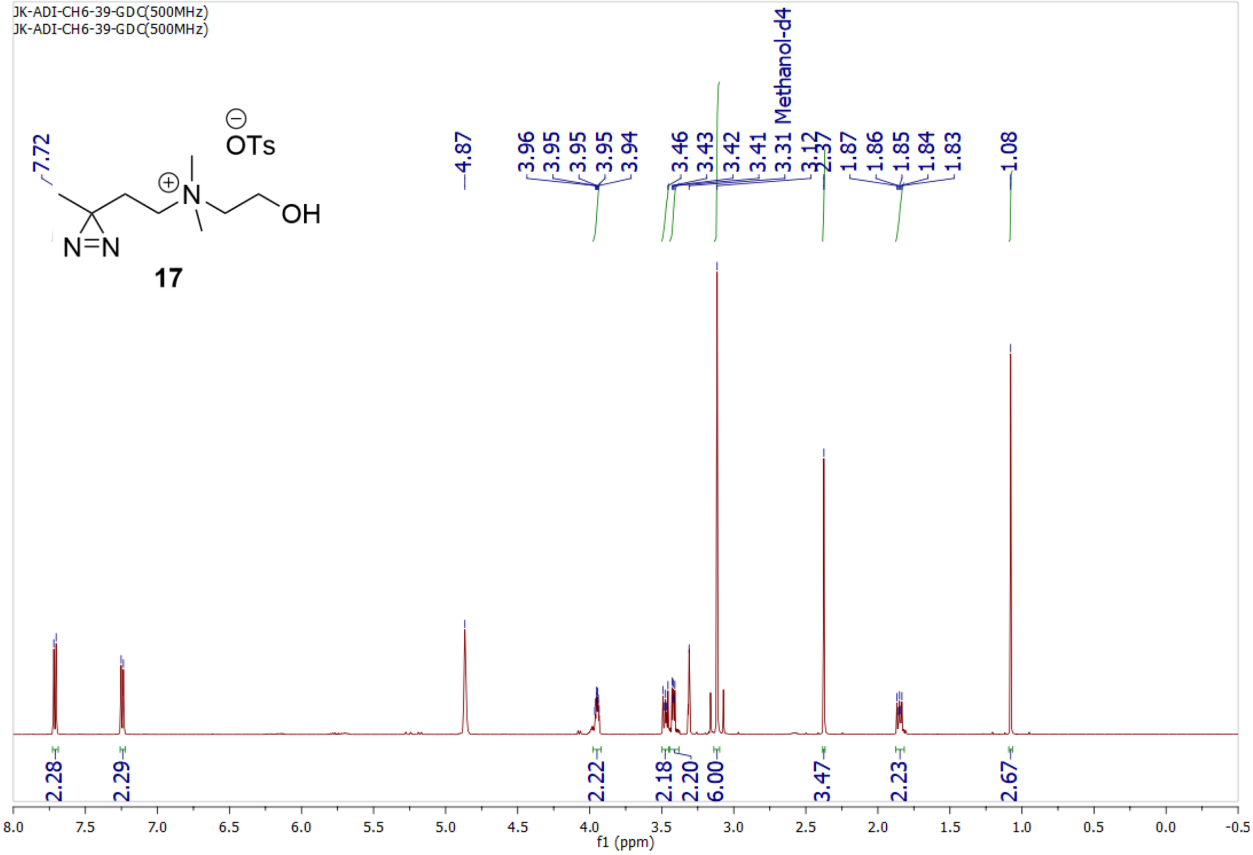
JK-ADI-CH6-39-GDC(500MHz)
JK-ADI-CH6-39-GDC(500MHz)
OTs
OH
N=N
17
7.72
4.87
3.96 3.95 3.95 3.95 3.94
3.46 3.43 3.42 3.41 3.31 Methanol-d4 3.12 2.37
1.87 1.86 1.85 1.84 1.83
1.08
2.28 2.29 2.22 2.18 2.20 6.00 3.47 2.23 2.67
f1 (ppm)

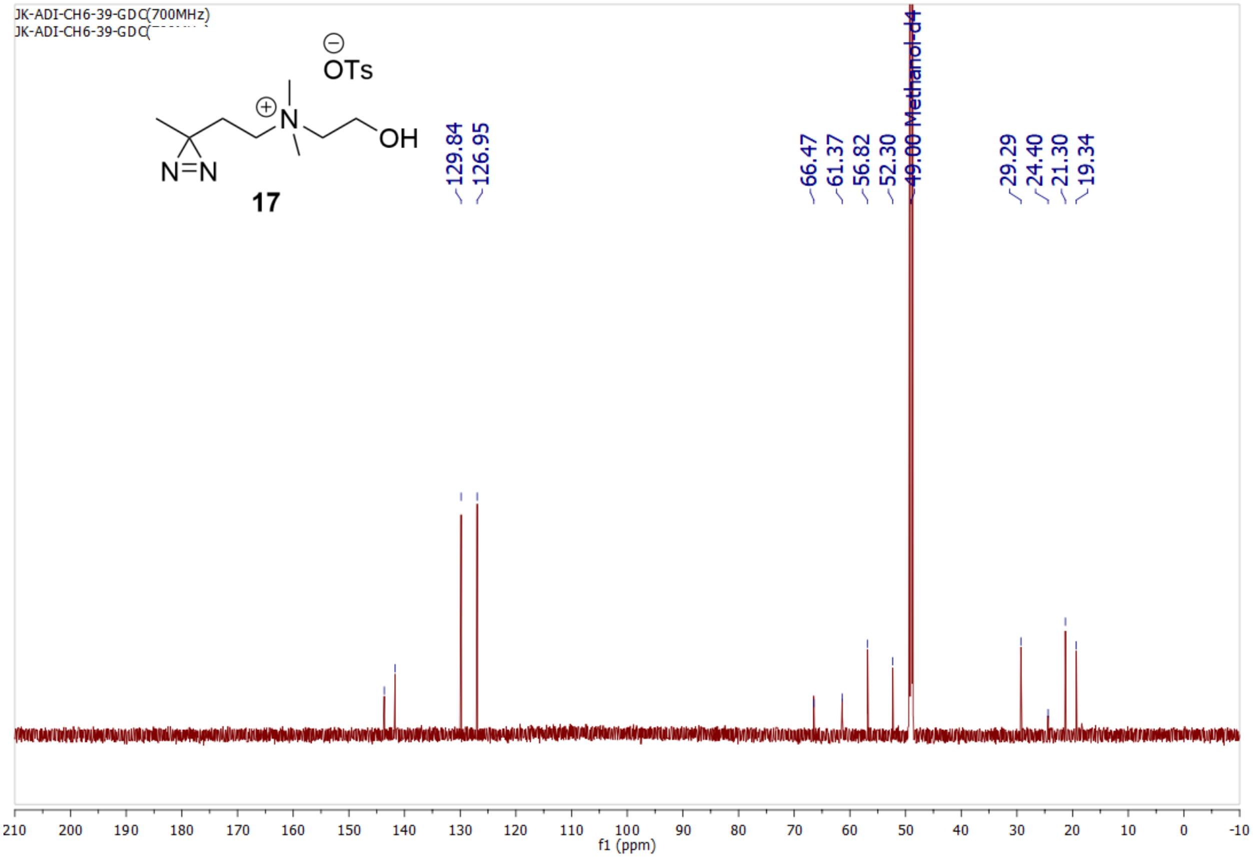
JK-ADI-CH6-39-GDC(700MHz)
JK-ADI-CH6-39-GDC
OTs
OH
N=N
17
129.84
126.95
66.47
61.37
56.82
52.30
49.00 Methanol-d4
29.29
24.40
21.30
19.34
f1 (ppm)

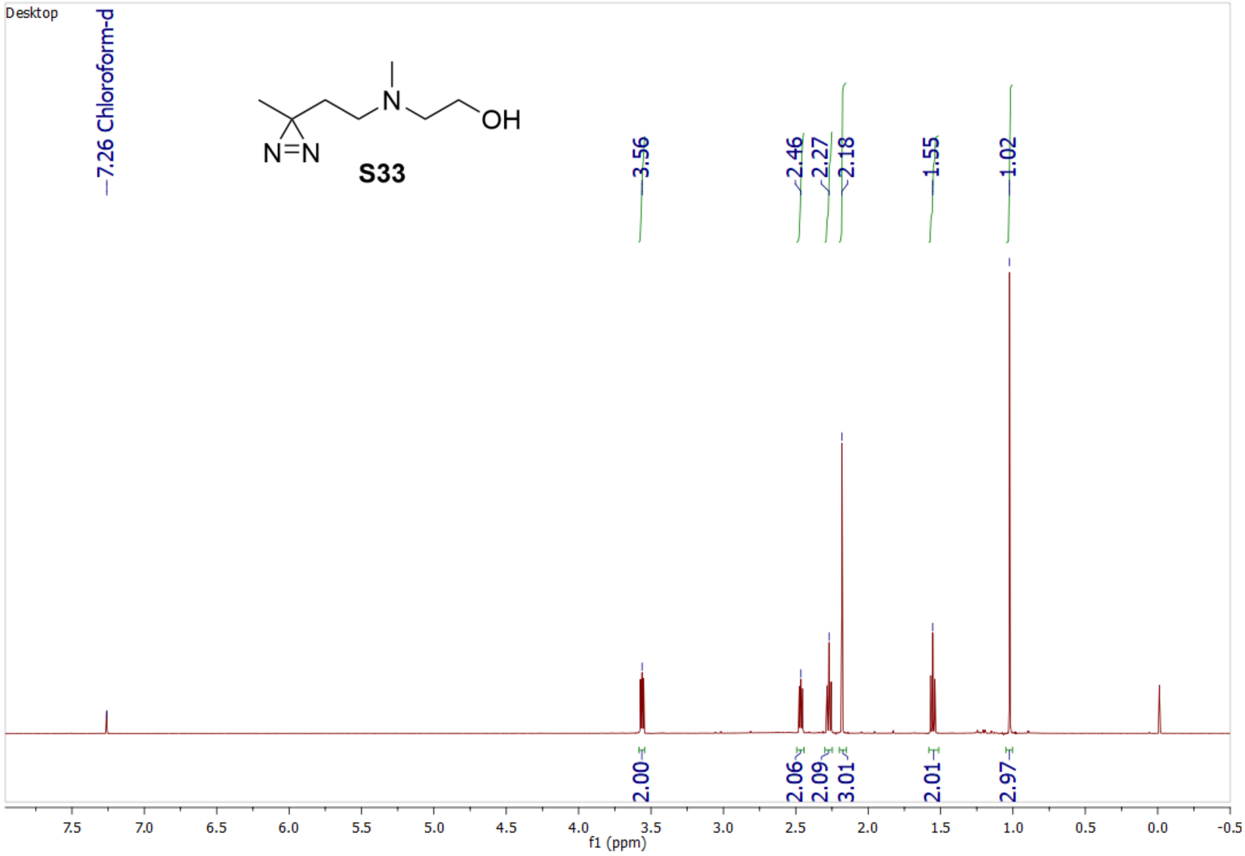
Desktop
7.26 Chloroform-d
N=N
N
OH
S33
3.56
2.46
2.27
2.18
1.55
1.02
2.00
2.06
2.09
3.01
2.01
2.97
7.5 7.0 6.5 6.0 5.5 5.0 4.5 4.0 3.5 3.0 2.5 2.0 1.5 1.0 0.5 0.0 -0.5
f1 (ppm)

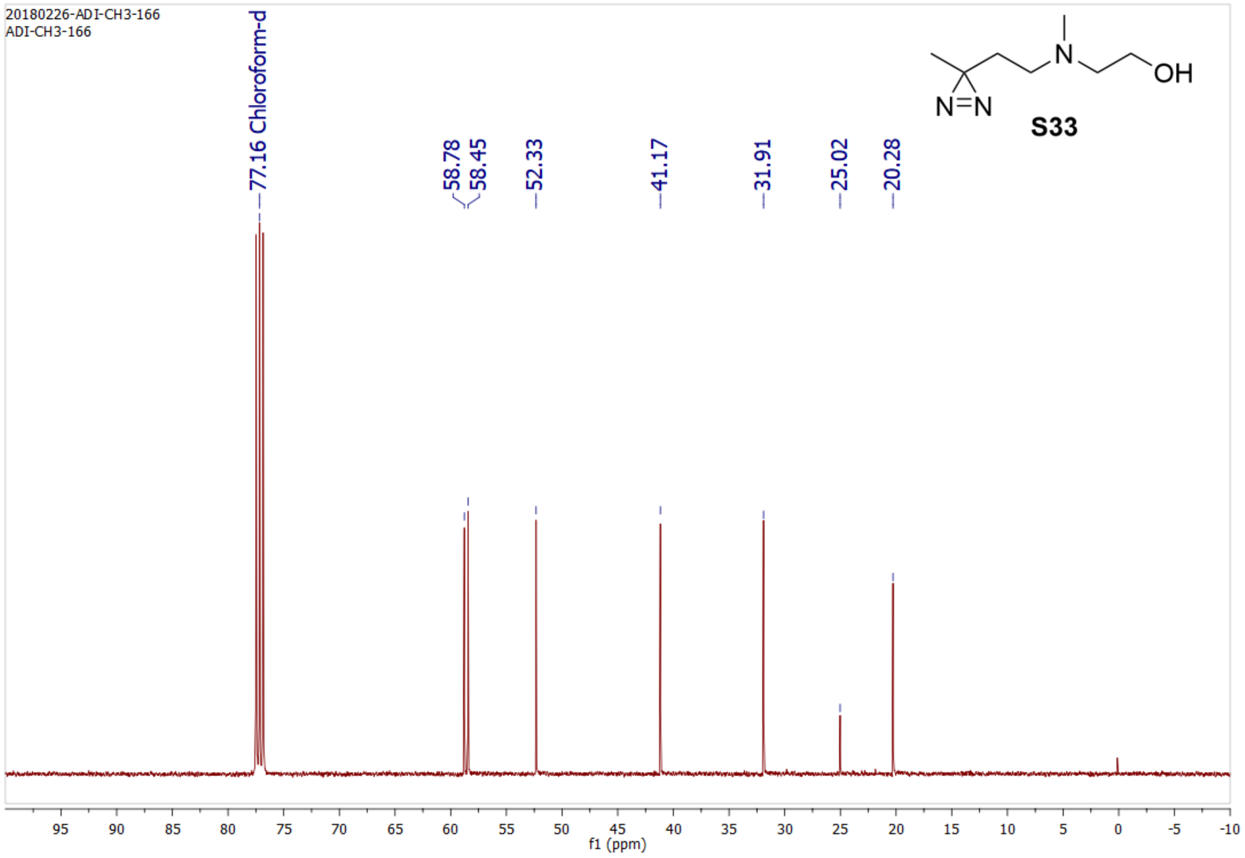
20180226-ADI-CH3-166
ADI-CH3-166
77.16 Chloroform-d
58.78
58.45
52.33
41.17
31.91
25.02
20.28
N=N
N
OH
S33
95 90 85 80 75 70 65 60 55 50 45 40 35 30 25 20 15 10 5 0 -5 -10
f1 (ppm)

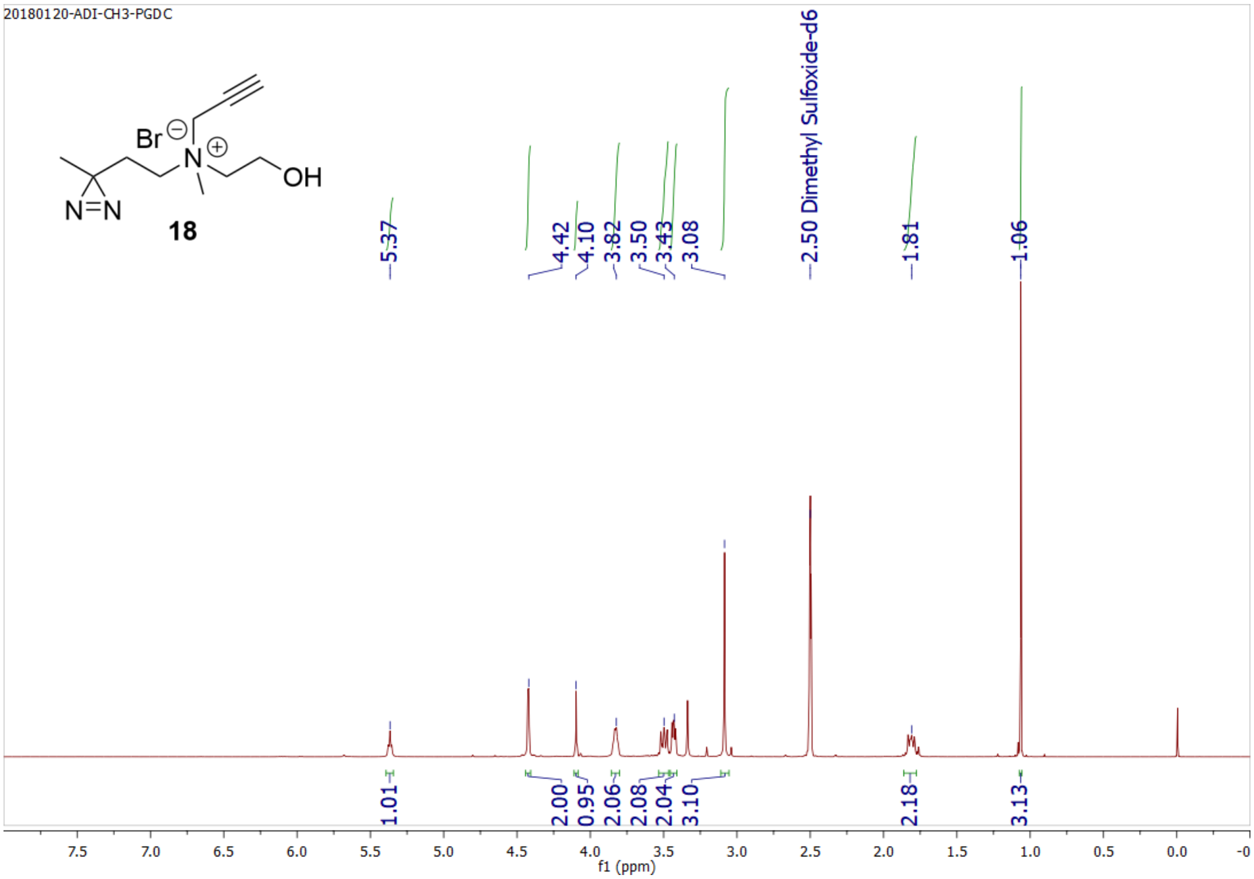
20180120-ADI-CH3-PGDC
Br
N
OH
N=N
18
5.37
4.42
4.10
3.82
3.50
3.43
3.08
2.50 Dimethyl Sulfoxide-d6
1.81
1.06
1.01
2.00
0.95
2.06
2.08
2.04
3.10
2.18
3.13
f1 (ppm)

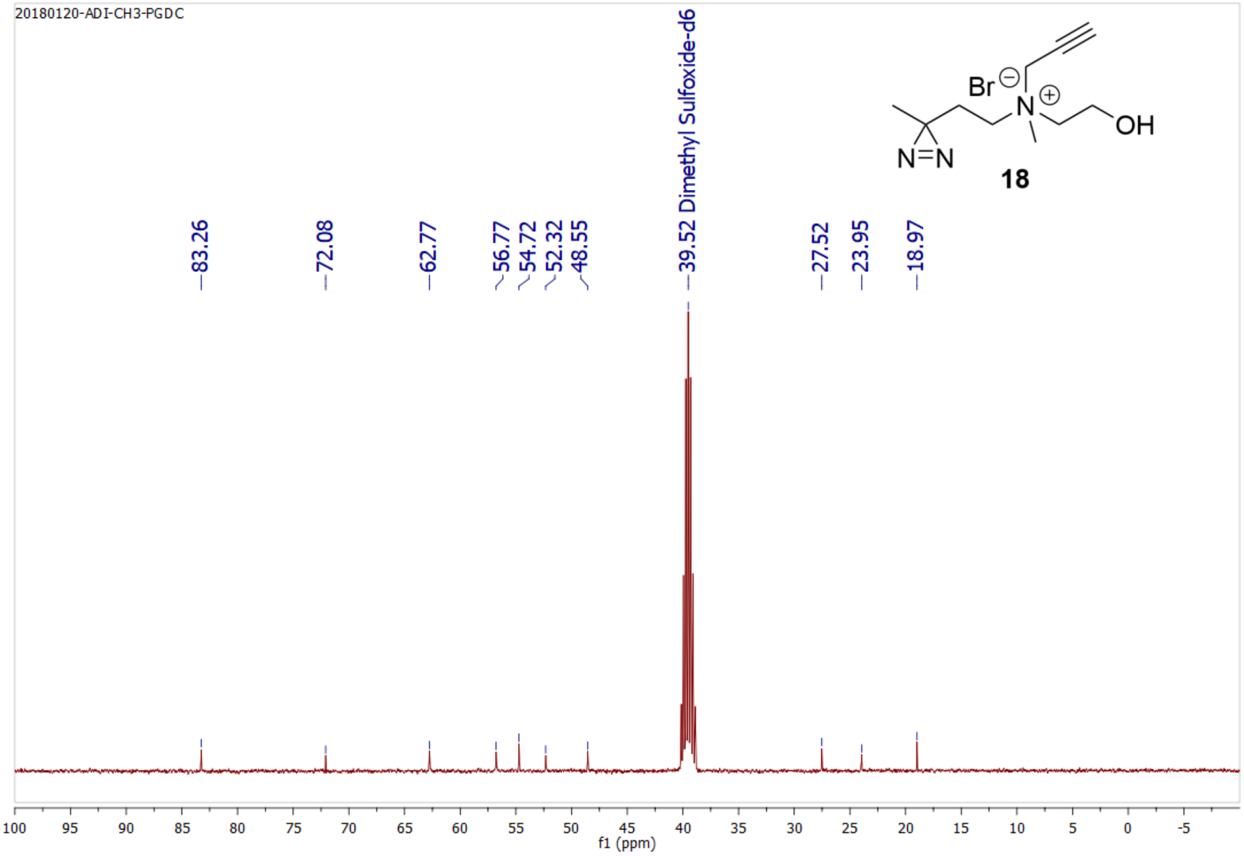
20180120-ADI-CH3-PGDC
Br
N
OH
N=N
18
83.26
72.08
62.77
56.77
54.72
52.32
48.55
39.52 Dimethyl Sulfoxide-d6
27.52
23.95
18.97
f1 (ppm)

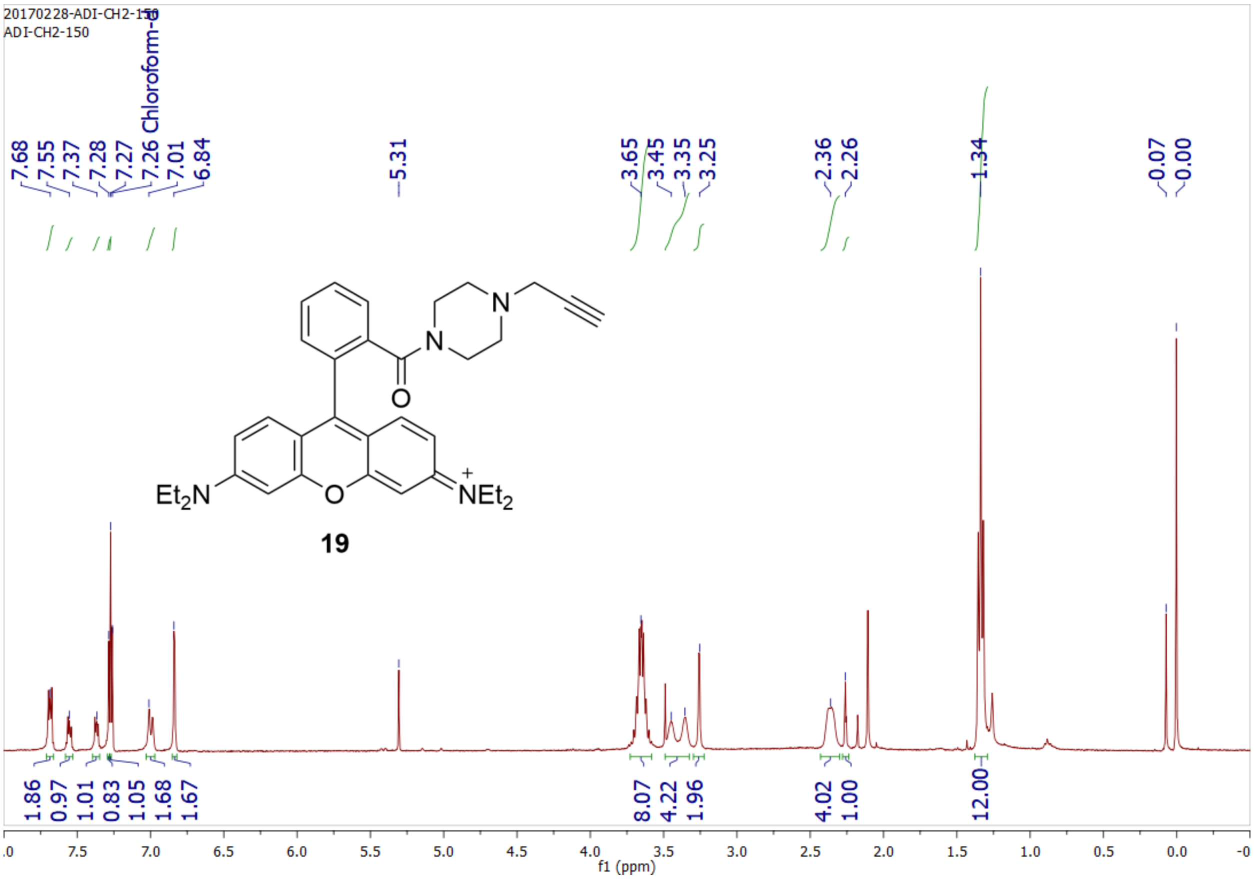
20170228-ADI-CH2-150
ADI-CH2-150
7.68
7.55
7.37
7.28
7.27
7.26 Chloroform-d
7.01
6.84
5.31
3.65
3.45
3.35
3.25
2.36
2.26
1.34
-0.07
-0.00
Et2N
O
N
N
O
NEt2
19
1.86
0.97
1.01
0.83
1.05
1.68
1.67
8.07
4.22
1.96
4.02
1.00
12.00
f1 (ppm)

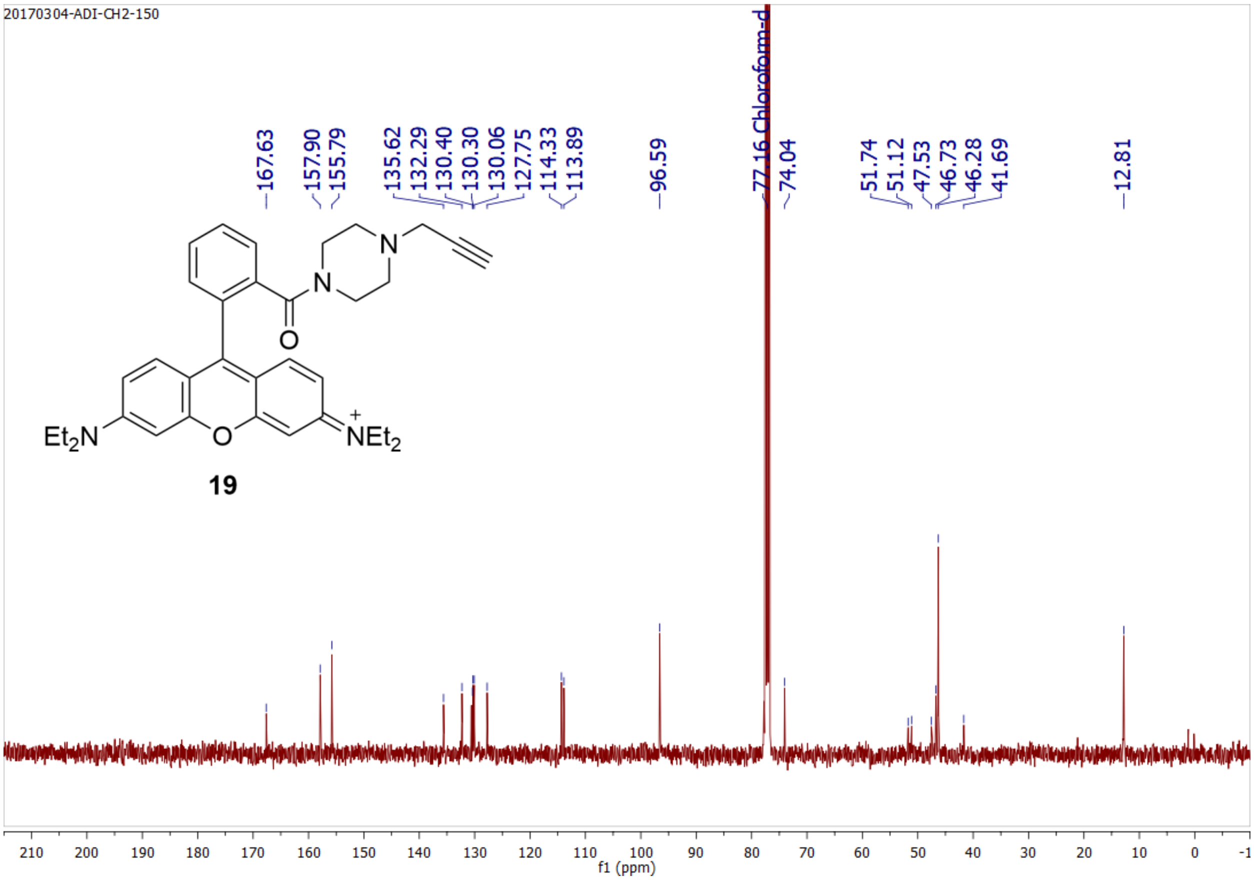
20170304-ADI-CH2-150
167.63
157.90
155.79
135.62
132.29
130.40
130.30
130.06
127.75
114.33
113.89
96.59
77.16 Chloroform-d
74.04
51.74
51.12
47.53
46.73
46.28
41.69
12.81
Et2N
O
N
N
O
NEt2
19
f1 (ppm)
